# Structural and dynamic consequences of inosine, 2′-O-methylation, and N6-methyladenosine modifications in RNA G-quadruplex: A molecular dynamics study

**DOI:** 10.1101/2024.09.20.614213

**Authors:** Nivedita Dutta, Indrajit Deb, Joanna Sarzynska, Ansuman Lahiri

## Abstract

RNA G-quadruplexes (rG4) are stable non-canonical structures, often found to regulate important biological processes such as transcription, translation, splicing, RNA localization, and other steps in gene expression. rG4 structures can also influence DNA-related processes e.g. DNA replication, telomere elongation and homeostasis, and recombination. Due to the involvement of rG4s in such important processes, these structures are potential therapeutic targets against different diseases e.g., viral infection and cancer. Hence, a better understanding of the structure and stability of rG4s and their role in various therapeutically important cellular processes can help in the design of effective therapeutic strategies for targeting such structures. In the present study, we present our findings on the structural and dynamic effect of RNA modifications (inosine; 2′-O-methylation, and m^6^A-methylation) on RNA G-quadruplex structure from molecular dynamics simulations. Additionally, we also report the dynamic and energetic aspects of inter-quadruplex interactions and the influence of non-G purine tetrads neighboring the inter-quadruplex junction on the interactions.

## Introduction

RNA G-quadruplexes (rG4) are composed of stacked planar G-tetrads each of which contains four guanine bases connected with Hoogsteen (HG) base pairs, forming a negatively charged core with O6 atoms and monovalent cations intercalated within the core [1]. While the structural, thermodynamic, and functional aspects of DNA G-quadruplexes have been widely studied in the past decades [2–5], RNA G-quadruplexes have also emerged as the motif of interest in several theoretical and experimental studies in recent years [6]. The occurrence of rG4 structures has been reported in rRNA[7], mRNA[8,9], lncRNA [10,11], and viral genomic RNAs [12–16]. RNA GQs (rG4s) have been suggested to be involved in RNA metabolism and DNA-related processes and such structures often modulate translation, mitochondrial transcription, mRNA 3′-end processing and polyadenylation, translocation of RNA, pre-miRNA splicing and miRNA biogenesis, piRNA processing, RNA-protein interactions and telomere homeostasis [17–19].

G-quadruplexes are commonly found in 5′-UTR regions of several genes (including those of therapeutic importance), e.g. Bcl-2, NRas, MT3-MMP, Zic-1, FMR1, TRF2, and ESR1, etc. and can significantly influence regulation of translation and RNA-protein interactions [20]. G4 structures are also found in mRNA 3′-UTR, e.g. in neurite mRNAs, and are suggested to regulate mRNA subcellular localization [6,21]. Although not very frequently, RNA G4 motifs have also been identified in coding regions of mRNA [18]. Experimental studies revealed the formation of dimeric rG4 structure with two stacked propeller-type parallel rG4 blocks in the human telomeric repeat-containing RNA (TERRA) [22], for which the solution NMR structure was later reported by Martadinata and Phan (2013) (PDB ID: 2M18) [23]. Although RNA G-quadruplexes have been reported to predominantly adopt the parallel conformation, the formation of antiparallel rG4 structure has been observed in the 8-bromo-guanosine modified human telomeric RNA sequence [24].

Metal ions (cations) play an essential role in the formation and stability of G4 structures and such ions are coordinated at the centers of the stacked tetrads forming a channel [6,18]. The trend in stabilization efficiency of the metal ions in the G4 channel has been reported to be: Sr^2+^ > Ba^2+^ > K+ > Na^+^, NH_4_^+^, Rb^+^ [6,17,25];. Interestingly, in vitro G4 motifs have been observed to dimerize, tetramerize, or form further higher-order motifs like G-wires [26–32]. Quadruplexes can multimerize through stacking interactions and also through short loop structures connecting the intramolecular quadruplex monomers and such multimeric GQs present in human telomeric regions are often associated with cancer and neurological disorders [32–35]. Hence such structures can be potential targets of drug ligands and especially anticancer drugs [34]. Such ligands can interact with the multimeric G4 structures via 1:1 binding mode in which one unit of the ligand interacts with each G4 monomer, 2:1 binding mode corresponding to the intercalation of the ligand unit between the consecutive stacked G4 monomer units or via a 2:1+1 binding mode in which, one ligand unit is seated between the consecutive stacked G4 monomers and the other unit interacts with one of the G4 monomer [32]. Li et al. carried out modeling and MD simulations of a dimeric DNA G4 structure and investigated the interaction of the Tel03 ligand with the G4 dimer in both 1:1 and 2:1 binding modes [36].

Guanosine (G) to Inosine (I) substitutions have been reported to be suitable for studying the consequences of modulation of a functional group and loss of a hydrogen bond, due to the structural similarity of the adenosine-derivative inosine with guanosine resulting in the similarity in the electrostatic potentials. Interestingly, experimental studies have suggested that poly(I) can form a four-stranded structure similar to the quadruplex structure formed by poly(G) (reviewed in [37]). Such structures are less stable compared to the standard G-quadruplexes due to a lesser number of H-bonds, i.e. a single H-bond per I-pair compared to two H-bonds in G-pairs within G-tetrads and their stabilities depend more on the ionic interactions [38–40]. The crystal structure of a quadruplex with an I-tetrad reported by Pan et al. revealed the presence of one N1-- -O6 H-bond per I-pair [29]. Substitution of A to I in RNA can lead to the formation of mixed GI tetrads and hence activate latent rG4 structures with the probability of contributing to the regulation of gene expression [41]. The formation of mixed GI-tetrads has also been observed in DNA quadruplex structures [42]. Recently, another study by Zheng and coworkers revealed the formation of (GIG)_4_ and (IGI)_4_ quadruplexes upon G-to-I substitutions within telomeric DNA repetitive sequence [43].

The influence of other modifications e.g. 2′-O-methyl-RNA (N_m_), LNA (locked nucleic acid), and UNA (unlocked nucleic acid) on G-quadruplex forming DNA sequences and the potential anticancer properties of such modified quadruplexes have been explored in earlier studies [44–48]. These chemical modifications can either stabilize or destabilize the structure of the DNA G-quadruplexes and the respective effect has been observed to be dependent on the position(s) of the substitution(s) and also the base orientation of the guanosine residue being substituted [44–48]. Roxo and Pasternak reported 2′-O-methylated RNA to effectively inhibit cancer cells without significant alteration of thermal and folding properties and resistance to certain enzymes [48]. Considering the abundance of 2′-O-methylation in RNA, understanding the effect of this modification in rG4 structures might also help in understanding the therapeutic potential of such modified rG4s.

Another commonly occurring RNA modification N6-methylated adenosine (m^6^A) has been reported to colocalize with G-quadruplexes. The effect of this modification (m^6^dA) on the thermal stability of DNA telomeric and c-Kit1 G-quadruplex structures (within loop regions) has been investigated with the findings suggesting that this modification can stabilize or destabilize the G4 structures depending on the position of the modification but the stability of quadruplexes with m^6^A-tetrads has not been investigated so far [49,50].

In this work, we present a theoretical comparison of the structural, dynamic, and energetic, hydration and ionization properties of I-, G-, and A-tetrads within similar monomeric and dimeric rG4 structures using extensive molecular dynamics simulations to understand the effect of inosine substitutions on the stability of such structural motifs. Additionally, we report the consequences of the presence of 2′-O-methylated I-, G-, and A-tetrads as well as N6-methylated A-tetrad within the monomeric rG4 structures respectively.

## Methods

### MD simulation of RNA G-quadruplex structures

From the crystal structure of the RNA G-quadruplex containing I-tetrad (PDB ID: 2GRB) [29], the monomer with chains A, B, C, and D was extracted and the 5′BrDU residues present in two of the strands were removed using UCSF chimera resulting in 5′GIGGU3′-quadruplex (**Figure 1**). Two other quadruplex systems were prepared by substituting the inosine(I)-tetrad with either guanosine(G)-tetrad or adenosine(A)-tetrad. The I-tetrad, G-tetrad, and A-tetrad at position 2 (i.e. the second tetrad position in the 5′), were further modified to the respective 2′-O-methylated forms, i.e. I_m_-tetrad, G_m_-tetrad and A_m_-tetrad respectively using the *tleap* module of *cpptraj*. Additionally, a similar quadruplex structure with m^6^A-tetrad at position 2 was also built using the tleap module. FF99_*χ*_kol0__*γ*_bsc0_ and FF99_*χ*_OL3__*γ*_bsc0_ parameters were used for inosine and the canonical residues respectively. For the 2′-O-methylated residues, a combination of FF99_*χ*_OL3__*γ*_bsc0_ parameters and specific parameters from modrna08 were used, while for m^6^A, the modrna08 parameters were used. Each system (except for the quadruplex with m^6^A-tetrad) was solvated in a truncated octahedral waterbox of TIP3P water molecules (with an 11 Å distance between any solute atom and the edge of the box) and was neutralized using Na^+^ ions for the first three simulation sets and using K^+^ ions for the fourth simulation set. For the monomer-quadruplexes containing I-, G-, and A-tetrads or their 2′-O-methylated counterparts, no additional salt was added to the systems for the first three sets of simulations while the fourth set of simulation was carried out at 150mM KCl concentration. For the quadruplex with m^6^A-tetrad, although the solvation procedure was the same, the neutralization was carried out using K^+^ ions in each of the three simulation sets and 150mM KCl salt concentration was considered for the simulations.

**Figure 1.**
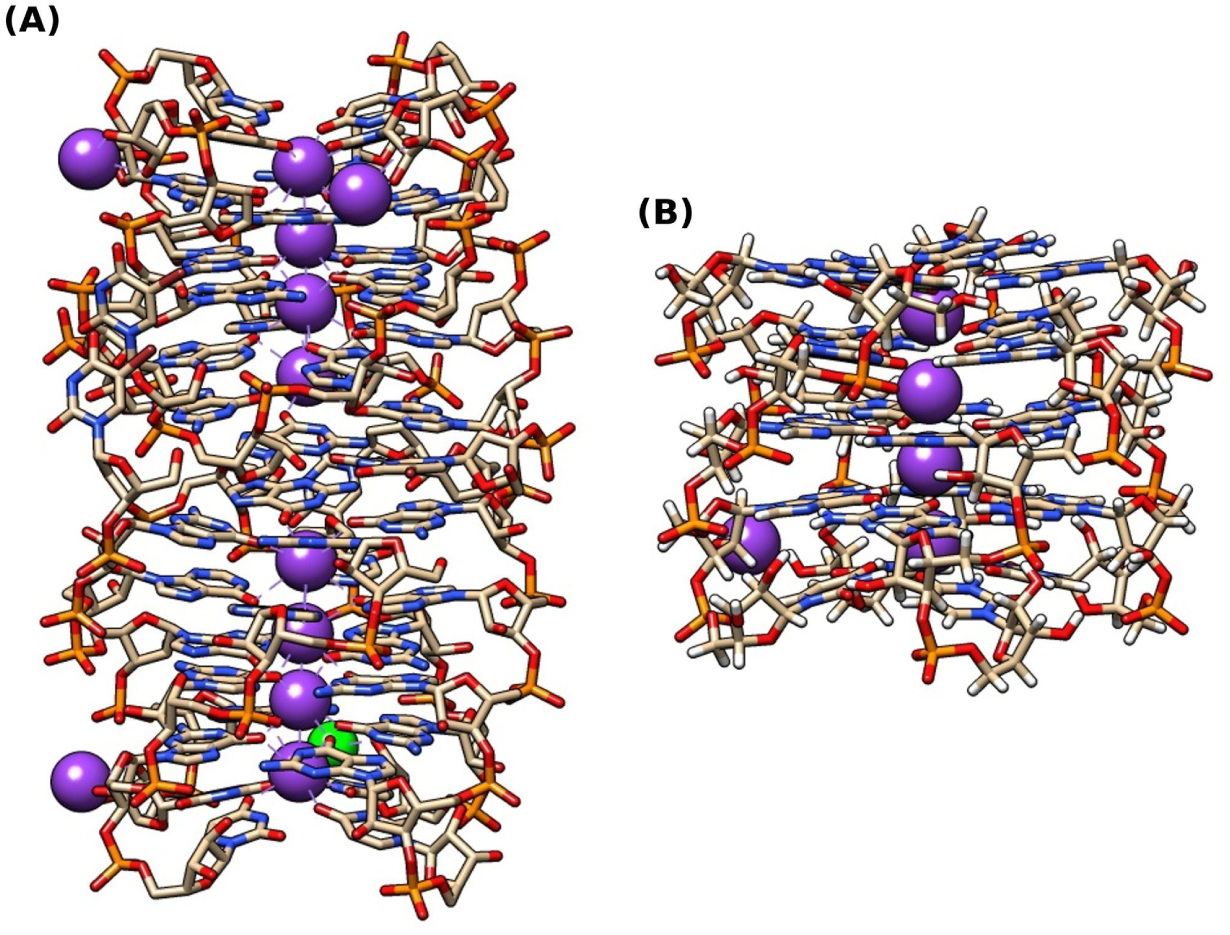
(A) Structure of the RNA quadruplex containing I-tetrad reported by Pan et al. [29] (PDB ID: 2GRB) (The structural water molecules are not shown in the diagram). The monomer with chains A, B, C, and D is shown in the bottom from 5′ to 3′. (B) The isolated monomer (GIGGU) (chains A, B, C, and D with the associated K+ ions) without the BrdU residues and the Sr+ ion present in the original crystal structure of the quadruplex.

Next, for the assessment of the structural and dynamic properties of the quadruplex-dimers, we also carried out simulations of the dimeric forms containing all the chains present in the reported crystal structure (A-H), where each monomer contained either I-tetrad or G-tetrad or A-tetrad at position 2. For each of the three dimer systems under this study, the BrdU5 residues were substituted with U5 using UCSF-chimera, and the Sr^2+^ ion was removed. Each dimeric quadruplex system was solvated in a truncated octahedral water box of TIP3P water molecules having 11 Å distance between any solute atom and the edge of the box and neutralized using K^+^ ions and an additional salt (KCl) concentration of 150 mM was considered.

A three-stage minimization was carried out for each of the monomeric or dimeric quadruplex structures. In the first stage, the quadruplex and the channel ions were kept fixed by the application of a force constant of 25 kcal/mol. In the next stage, only the quadruplex was kept fixed with a reduced restraining force of 5 kcal/mol. In the third stage of minimization, the whole system was minimized without any restraints. In each of these stages, the initial 200 steps of steepest descent followed by 300 steps of conjugate gradient minimization were performed. After minimization, the system was heated from 0 to 300K keeping the quadruplex restrained with a force constant of 15 kcal/mol. Then the system was equilibrated in four consecutive steps each 100 ps with a gradual decrease of the force on the solute from 10, 5, 1, and 0.5 kcal/mol. Next, one set of 1.5 μs simulation and three independent sets of 1 μs simulations yielding a total of 4.5 μs simulation was carried out for each of the monomeric quadruplex systems (except for the monomeric quadruplex system containing m^6^A-tetrad at position 2, for which 3 independent sets of 1.5 μs simulations, i.e. a total of 4.5 μs simulation was carried out). For each dimer system, three independent sets of 1μs simulations and another set of 2 μs, i.e. a total of 5 μs were carried out. All the simulations were carried out using the *pmemd* module of the AMBER18 package [51].

### Analyses of the simulated ensembles

The conformations from the entire simulated trajectories for the monomeric quadruplexes with I-tetrad, G-tetrad, A-tetrad, and their 2′-O-methylated counterparts at position 2 were used for analyses. For the quadruplex with the m^6^A-tetrad, the last 1 μs trajectories (out of 1.5 μs) from each of the three simulation sets were utilized for the analyses. These quadruplexes were primarily analyzed based on RMSD (root mean square deviation), RMSF (root mean square fluctuation), hydrogen bonding energies (for tetrads at position 2) and stacking interaction energies (of the tetrad at position 2 with the neighboring G-tetrads at positions 1 and 3 respectively), interaction energies of the tetrads at position 2 with the nearest channel K^+^ ions (i.e. K22 and K23 in the crystal structure), glycosidic torsion angles and sugar pucker conformation of all residues, backbone torsion angles (for the tetrad at position 2), N9-pyramidalization (for the tetrads at positions 2, 3 and 4) using suitable tools within AmberTools21 [52]. The *‘lie’* tool was utilized for estimating the molecular mechanics-based hydrogen bonding, stacking, and ion-interaction energies. The calculation of Gibbs free energies was carried out using the *MMPBSA.py* program. Different HG base-pair and base-pair step parameters were calculated for the monomeric rG4s using the Curves+ and Canal programs [53].

For the dimeric quadruplexes, the entire trajectories from the first three simulation sets and the last 1 μs of the fourth simulation set were used for the analyses. Similar analyses as those for the monomeric rG4 structures were also performed for the constituent monomeric rG4s within the simulated ensembles of the dimeric quadruplexes. The molecular mechanics-based stacking energies between the 5′-U*(GGGG) pentads (using *‘lie’*) and the dimerization free energies (using *MMPBSA.py)* were also calculated for the dimeric rG4 structures.

To compare the effect of 2′-O-methylation of the G-tetrad, analysis of the fluctuations of some additional parameters including twist angle (ϴ), distances between tetrad-COMs, distance of constituting G/Gm residues from the respective tetrad COMs, angles (ϕ) between normals to the G/G_m_-planes and the tetrad-axis, angles (θ) between G/G_m_-pairs and G/G_m_-triads, were carried out using the VMD software (version 1.9.2) (https://www.ks.uiuc.edu/Research/vmd/) using the scripts obtained from Tsvetkov et al. [54].

## Results and Discussion

### Structural and thermodynamic properties of monomeric quadruplexes

#### Consequences of the presence of inosine and 2′-O-methylation in G-quadruplex structure

From our simulations of the monomeric G-quadruplexes (GIGGU) containing the I-tetrad and those in which the I-tetrad was replaced with G- and A-tetrads respectively, we observed that the overall conformations were similar as reported by Pan and coworkers, [29]. The time series of the RMS deviations from the initial minimized structure (for the first three tetrads from the 5′-direction) suggested overall similar conformations of the quadruplexes Quad-I4, Quad-G4, and Quad-A4 especially in the fourth set (with additional KCl salt) **(Figures 2A; S1A)**. The RMS fluctuations (for each of the residues in the GQ) from the average structure were observed to be similar (except for the 3′ U20 residue of Quad-I4 in simulation set-1, U10 residue of Quad-A4 in simulation set-3 which might be due to the absence of the additional KCl salt) (**Figures 2B; S1B)**. Interestingly, the stacking interaction energies of A-tetrad with G-tetrads at its 5′ and 3′ were observed to be lower compared to the similar interactions of I- and G- tetrads suggesting that A-tetrad may form more stable stacking interactions compared to I- and G-tetrads within the quadruplex structure (**Figures 2C, D; S3; Table S1**). Single G/A base stacking interaction has been previously reported to be energetically more stable compared to G/G base stacking interaction [55].

**Figure 2.**
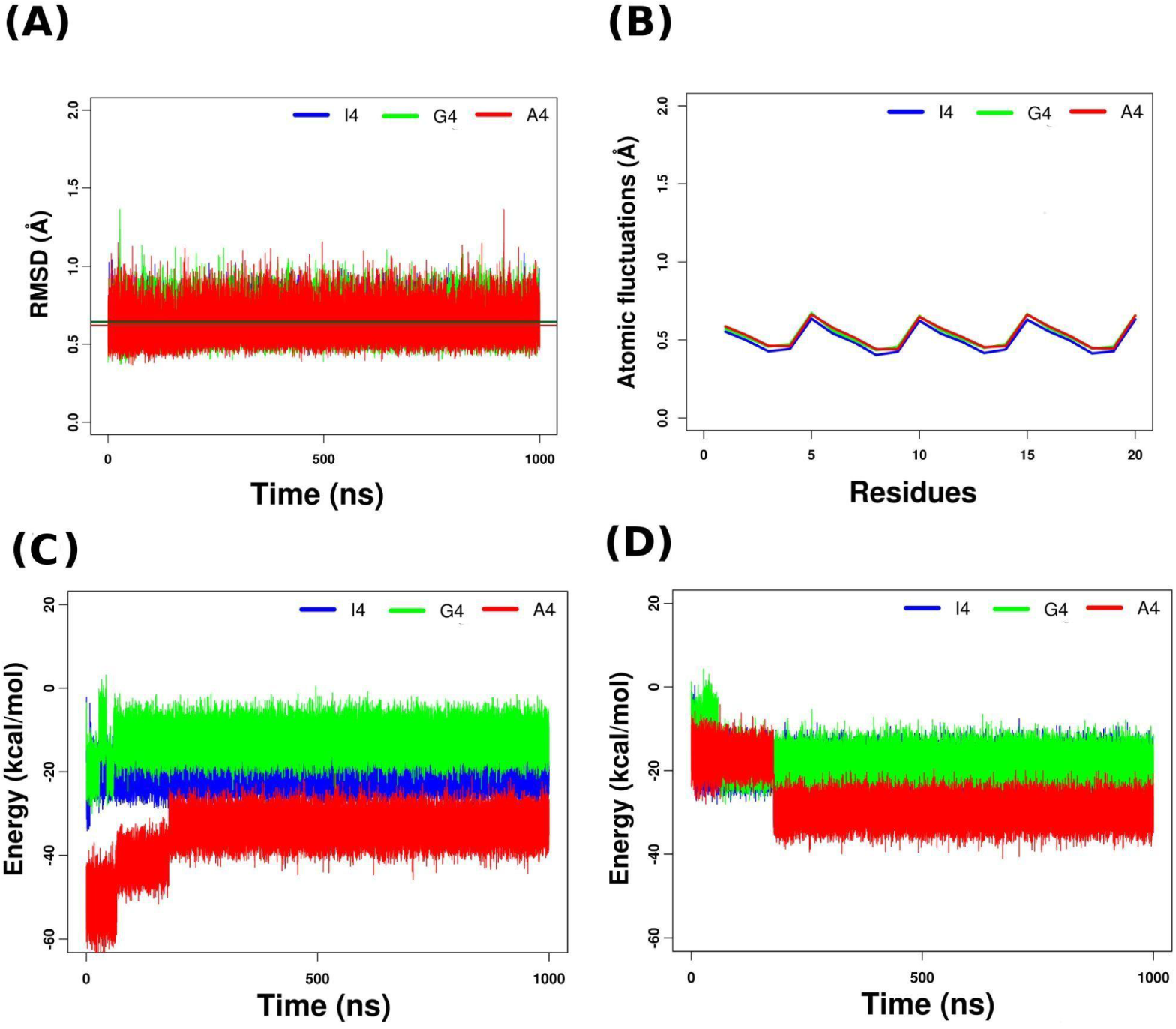
(A) RMSD of heavy atoms of the nucleotides of the three (towards 5′) tetrads; (B) residue-wise RMSF (heavy atoms); (C) stacking interaction energies for the G4(1)-X4(2) tetrad step and (D) that for the X4(1)-G4(2) tetrad step (where X indicates I/G/A) for the quadruplexes containing the I tetrad (Quad-I4), G-tetrad (Quad-G4) and A-tetrad (Quad-A4) at position 2 respectively for the fourth set of simulations.

2′-O-methylation of the I-/G-/A-tetrads at position 2 did not result in significant alteration of the overall conformations of the quadruplexes (**Figures 3A, B; S4; S5**). However, the pattern observed for the RMS fluctuations for the residues in Quad-I_m_4/G_m_4/A_m_4 and those for the residues in Quad-I4/G4/A4 were different (**Figures S2; S5**). The tetrad stacking interaction energies of A_m_-tetrad with G-tetrads at its 5′ and 3′ (especially in simulation set-4, i.e. in the presence of additional KCl salt) indicated enhanced stacking stabilization of the quadruplex in the presence of the A_m_-tetrad compared to I_m_- and G_m_- tetrads (**Figures 3C, D; S6; Table S1**).

**Figure 3.**
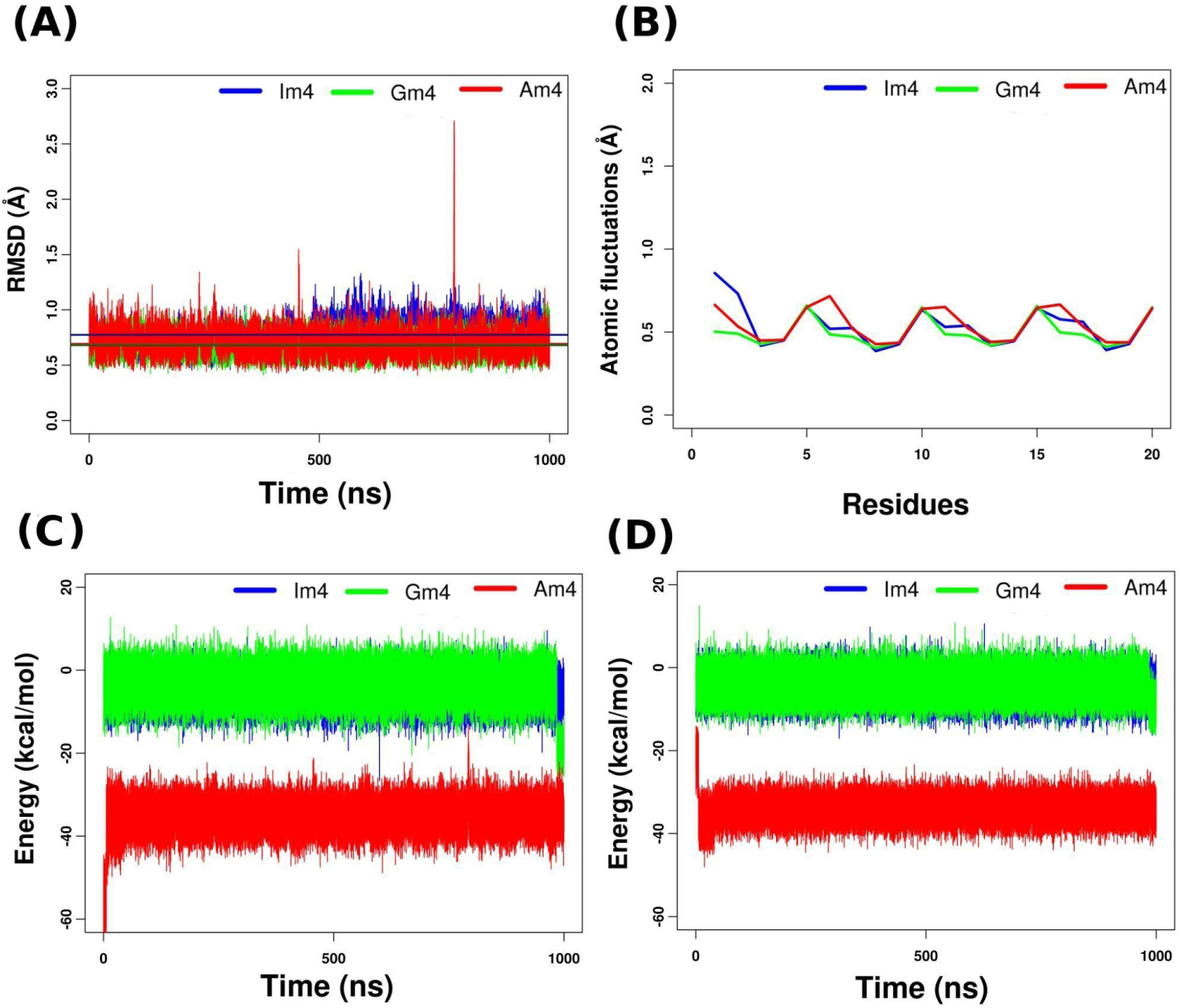
(A) RMSD of heavy atoms of the nucleotides of the three (towards 5′) tetrads; (B) residue-wise RMSF (heavy atoms); (C) stacking interaction energies for the 5′-G4(1)-X4(2)-3′tetrad step and (D) that for the 5′-X4(1)-G4(2)-3′ tetrad step (where X indicates Im/Gm/Am) for the quadruplexes containing the Im tetrad (Quad-Im4), G-tetrad (Quad-Gm4) and A-tetrad (Quad-Am4) at position 2 respectively for the fourth set of simulations.

Each of the residues in the studied quadruplexes was in the *ANTI* base orientation throughout the simulations. Andrałojć et al reported the *C2′-endo* conformation for the residues in the G-tetrad at position 2 for the quadruplexes in their study [56]. However, from our simulations, all the purine tetrads within the quadruplex monomers under this study contained residues in the C3′-endo conformation as reported in the crystal structure of the RNA quadruplex.

For the monomeric quadruplex, the N1(H)-O6 hydrogen-bonded geometry of the I-tetrad observed in the crystal structure was maintained during the simulation for each of the simulation sets (**Table S2**). For the G-tetrad, stable geometries containing N1(H)-O6 and N2(H)-N7 hydrogen bonds were sampled, while the A-tetrad was observed to preferentially adopt the N6(H)-N1 [57,58] geometry (**Figures 4; S7**). The total hydrogen bonding energy for the I-tetrad was observed to be ∼ -32 kcal/mol and those for the G- and A-tetrads were found to be ∼ -54 kcal/mol (except in the first simulation set in which the value was observed to be ∼ -58.4 kcal/mol) and ∼15.3 kcal/mol respectively for each of the simulation sets **(Tables 1, S1)**. The hydrogen bonding energy for the I-tetrad obtained in this study for the N1(H)-O6 geometry is much lower compared to that (-7.4 kcal/mol) obtained for dI-tetrad in the study by Stefl et al. [59] with the N1(H)-N7 geometry suggesting greater stability of the N1(H)-O6 geometry over the N1(H)-N7 geometry. The hydrogen bonding energy for RNA G-tetrad observed in our study was also found to be ∼10 kcal/mol lower compared to that reported for dG-tetrad with similar geometry (i.e. the hydrogen bonding (HG base pairing) interactions were ∼10 kcal/mol more stable for the rG-tetrad than the dG-tetrad) [59]. Earlier studies also suggested the enhanced thermal and thermodynamic stability of RNA G-quadruplex structures compared to their DNA counterparts most probably due to the improved tetrad stacking interactions and additional H-bonding interactions due to the presence of the 2′-OH group in ribose sugar [60–62].

**Figure 4.**
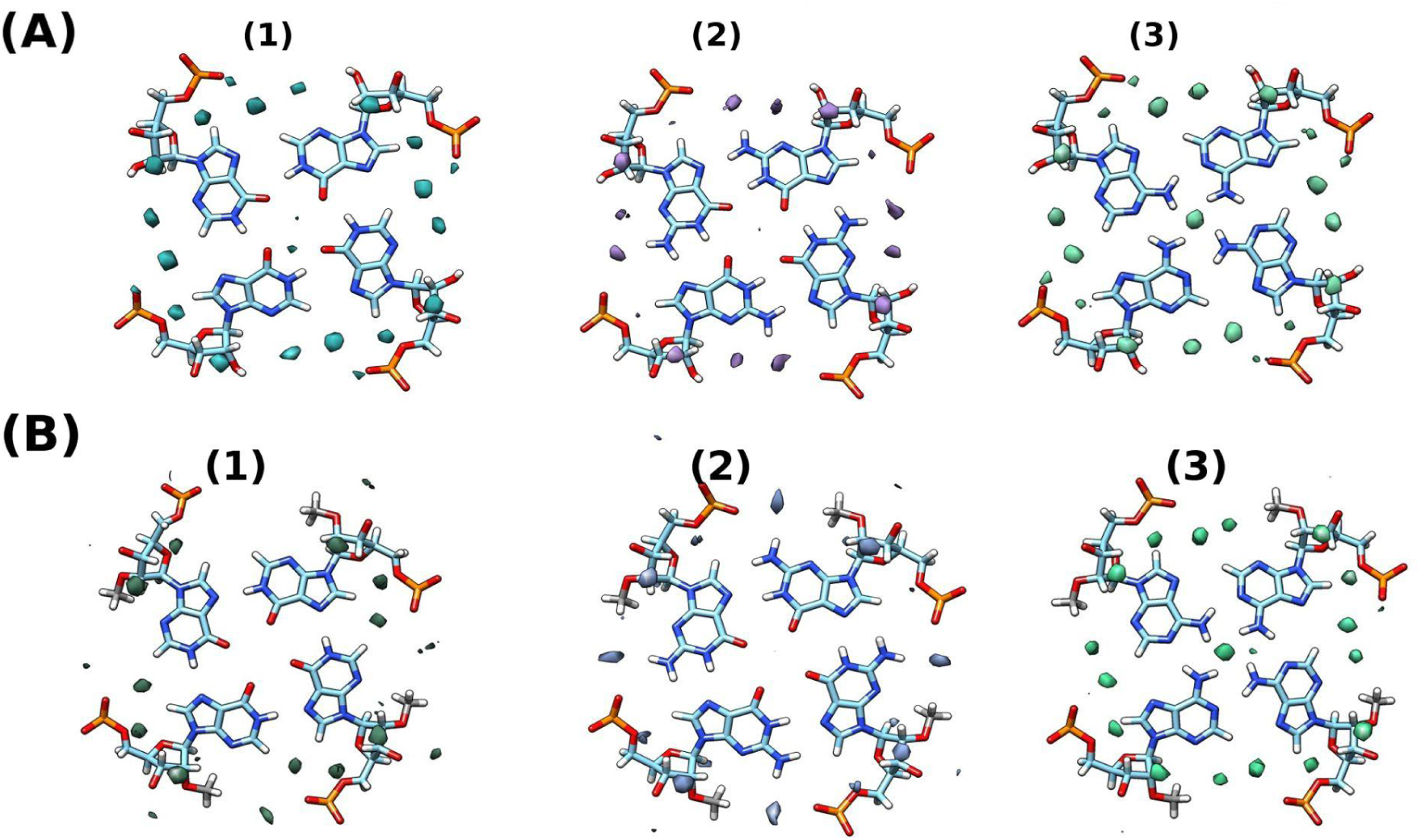
Water occupancy maps for (A) unmethylated tetrads (1) I-tetrad (2) G-tetrad (3) A-tetrad and (B) 2′-O-methylated tetrads (1) I_m_-tetrad (2) G_m_-tetrad (3) A_m_-tetrad respectively for the fourth set of simulations (around the average structures).

For the I_m_-tetrad, while the N1(H)-O6 geometry was predominantly sampled in the first three simulation sets, in the fourth simulation set, the N1(H)-N7 geometry (similar to that reported by Stefl and coworkers for the dI-tetrad [59]) was observed to be preferred (**Figures 4; S7; Table S2**). The N1(H)-N7 geometry was also observed in a lower population of conformers in simulation set-1 as well (**Table S2A**). For the G_m_-tetrad, in the first three simulation sets, we observed similar geometry, i.e. with N1(H)-O6 and N2(H)-N7 hydrogen bonds as was observed for the G-tetrad at the same position within a similar quadruplex. But in simulation set-4, in the presence of KCl, interestingly, a geometry with N1(H)-N7 and N2-H21---N7 hydrogen bonds (with much lower H-bond frequencies) was comparatively predominant over the standard geometry with two hydrogen bonds (**Figure 4; S7; Table S2D**). However, for the A and A_m_-tetrads, similar geometries (N6(H)-N1) were sampled for each simulation set and the total hydrogen bonding energy for the A_m_-tetrad was ∼ -17.3 kcal/mol (in simulation sets 1 and 4 while in simulation sets 2 and 3, the values were ∼ -20.4 kcal/mol and ∼ -18.4 kcal/mol respectively) indicating stabilization of the A-tetrad upon 2′-O methylation. On the other hand, destabilization of tetrad hydrogen bonding interactions was observed for I_m_- and G_m_- tetrads compared to the unmethylated counterparts. For the I_m_-tetrad, the total hydrogen bonding energy was observed to be ∼ -10 kcal/mol in simulation set 1, ∼ 13.8 kcal/mol in simulation sets 2 and 3, and ∼ -0.7 kcal/mol in simulation set 4 suggesting lesser stability of rI_m_-tetrad with N1(H)-N7 geometry compared to dI-tetrad with similar geometry [59] (**Tables 1, S2**). The thermal stabilization of DNA G-quadruplex upon substitution of dG-tetrad with G_m_-tetrad at position 2 has been reported by an experimental study recently [48]. In the present study, for the G_m_-tetrad the total H-bonding energy was found to be -47.2 kcal/mol, -32.8 kcal/mol, -34.8 kcal/mol for the first three simulation sets. In the fourth simulation set, in the presence of KCl, a total H-bonding energy of -12.4 kcal/mol was observed corresponding to the G_m_-tetrad geometry (with unstable N1(H1)---N7 and N2-H21---N7 hydrogen bonds) (Figure 4; Table S2D). Hence in general, the hydrogen bonding interactions were energetically less favorable for the G_m_-tetrad than those for the G-tetrad for the quadruplex sequence context in this study. It is important to note that the H-bonding energies computed here are for the entire trajectory and hence the averaged contributions from different intermediate geometries were also taken into account to examine the overall stability.

Each of the monomeric quadruplex structures contains a reversed U-tetrad at 3′ which might also contribute to the overall stability of the G-quadruplex structure as reported earlier by Andrałojć et al [56]. The uridine residues in the reversed 3′ U-tetrads in the studied quadruplex structures were in the SOUTH sugar pucker conformation and formed a buckled U-tetrad conformation resulting from the interaction between O4 and the K^+^ ion in the channel on the 5′ of the U-tetrad as reported in the experimental structure [29]. The signature H-bond between G-OP2 and 3′U-O2′ was observed to be in general highly stable observed for the monomeric quadruplexes with reversed-3′U-tetrads (**Table S3**). The stable N3(H)-O4 hydrogen-bonded geometry of the reversed U-tetrad was observed throughout the simulation (**Table S2**).

The purine tetrads at position 2 were additionally stabilized by water-bridging interactions (**Figure S7; Table S4**). The hydration pattern for the I-tetrad was observed to be generally similar to the G-tetrad as was reported by Pan et al. [29]. Two bridging water molecules were observed to connect the O2′ (through HO2′), N3, N2 (through H22), and OP2 atoms for each G-G HG pair in the G-tetrad. Similarly, for the I-tetrad, two water molecules connected the O2′ (through HO2′), N3, N7, and OP2 atoms for each I-I HG pair. Pan et al. [63], in their study, observed the presence of three water molecules bridging each A-A pair for the A-tetrad in the N6(H)-N7 geometry (with sodium (Na^+^) ion at the center). In our simulated ensembles, the hydration pattern of A-tetrad involved two water molecules bridging each A-A pair connecting the N3, O2′ (through HO2′), N7, and OP2 atoms for the N6(H)-N1 geometry of the A-tetrad (**Figure S7; Table S4**). 2′-O methylation resulted in an alteration of the hydration pattern for the purine tetrads in the sturdy, especially for the Im and G_m_-tetrads. Only one water molecule was observed to bridge the I_m_-I_m_ (connecting the N7 and OP2 atoms) and G_m_-G_m_ pairs (connecting N2 (through H22) and OP2 atoms). For the A_m_-tetrad, a chain of two water molecules was observed to connect the N3, N7, and OP2 atoms and hence the overall hydration pattern was not very different from that observed for the A-tetrad (**Figure S7, 8**).

The stabilities of the monomeric quadruplexes containing unmethylated and 2′-O-methylation tetrads (at position 2) analyzed using the MM-GBSA approach, showed a similar trend, i.e. Quad-I4 < Quad-A4 < Quad-G4 and Quad-Im4 < Quad-Am4 < Quad-Gm4 respectively for each simulation set (**Tables 2; S5**). There was not much difference between the free energies computed for the Quad-I4 and Quad-G4 with their respective 2′-O-methylated counterparts. But Quad-A4 was observed to be more stable compared to Quad-A_m_4 in this study from the MM-GBSA analysis while in contrast to this observation, the overall H-bonding and stacking interactions were more stabilized for the A_m_-tetrad compared to the unmethylated counterpart (**Table 1, S1**).

**Table 1.**
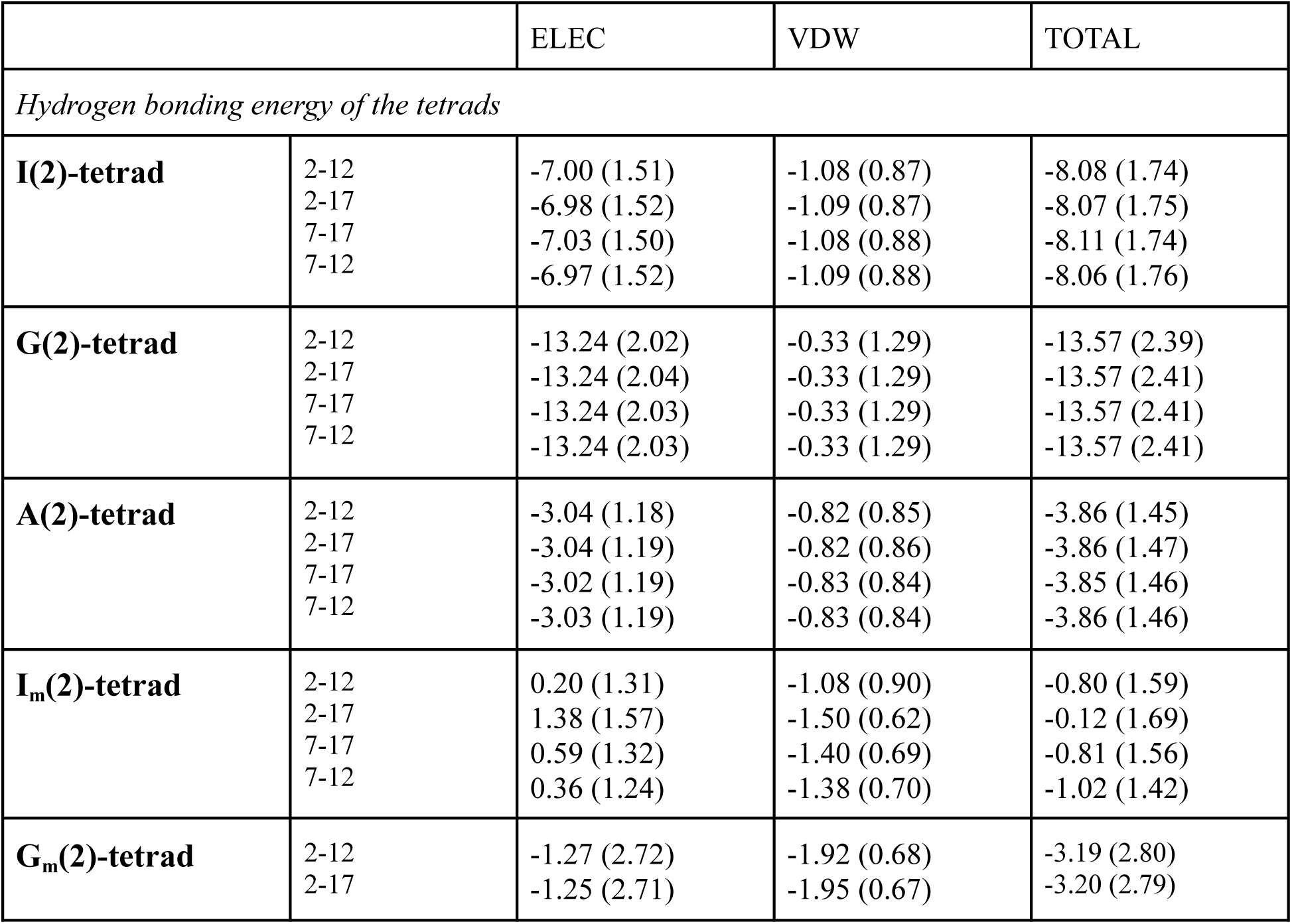
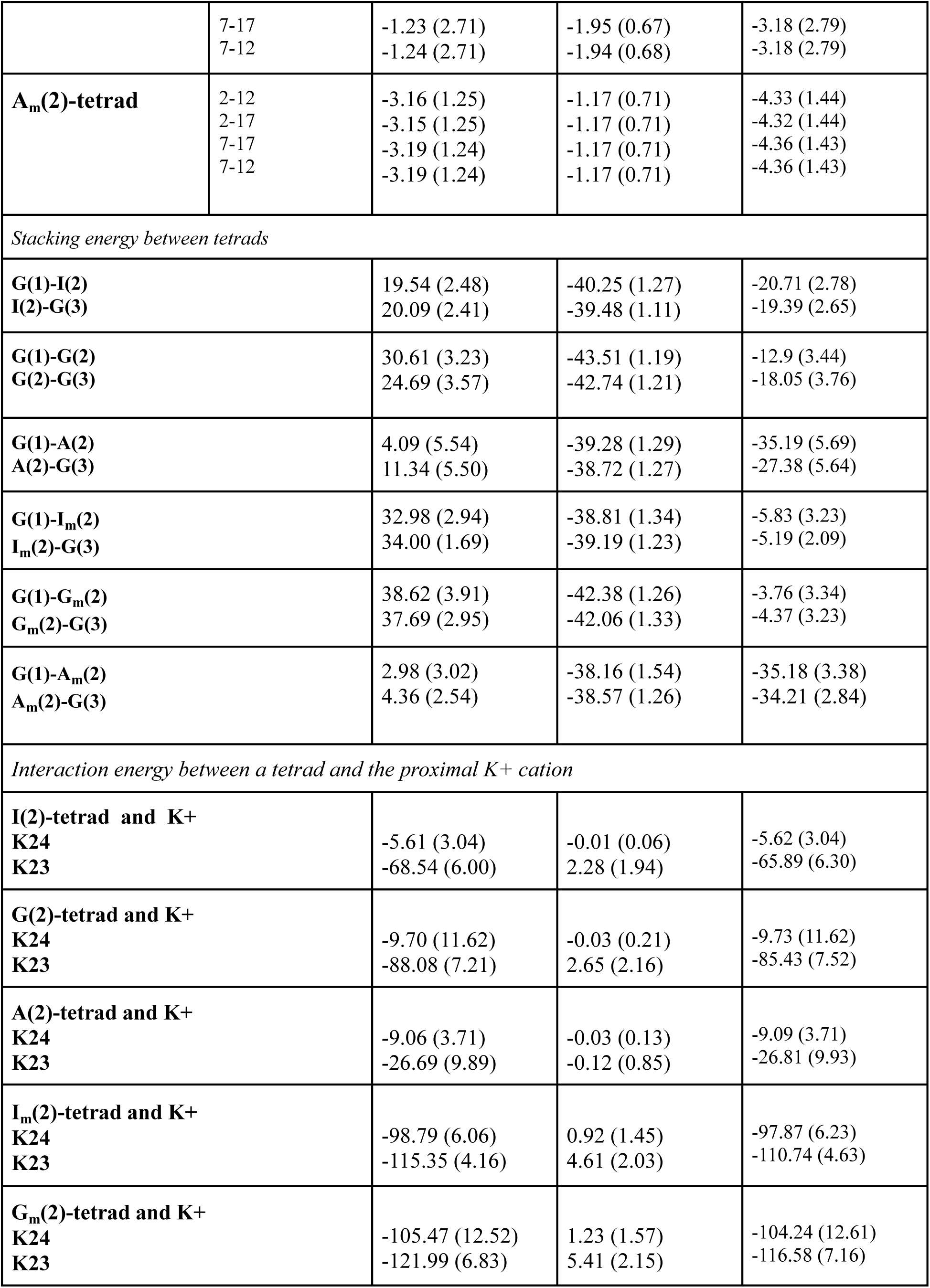
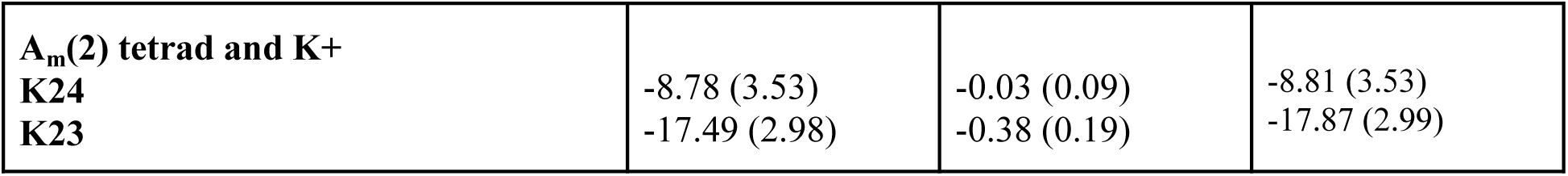
Interaction energies (kcal/mol) for hydrogen bonding, stacking, and ion interactions for the different purine tetrads at position 2 of the monomeric quadruplexes for the fourth set of simulations. Average and standard deviation (within brackets) values are reported.

|  |  | ELEC | VDW | TOTAL |
| --- | --- | --- | --- | --- |
| *Hydrogen bonding energy of the tetrads* |  |  |  |  |
| **I(2)-tetrad** | 2-12 | -7.00 (1.51) | -1.08 (0.87) | -8.08 (1.74) |
|  | 2-17 | -6.98 (1.52) | -1.09 (0.87) | -8.07 (1.75) |
|  | 7-17 | -7.03 (1.50) | -1.08 (0.88) | -8.11 (1.74) |
|  | 7-12 | -6.97 (1.52) | -1.09 (0.88) | -8.06 (1.76) |
| **G(2)-tetrad** | 2-12 | -13.24 (2.02) | -0.33 (1.29) | -13.57 (2.39) |
|  | 2-17 | -13.24 (2.04) | -0.33 (1.29) | -13.57 (2.41) |
|  | 7-17 | -13.24 (2.03) | -0.33 (1.29) | -13.57 (2.41) |
|  | 7-12 | -13.24 (2.03) | -0.33 (1.29) | -13.57 (2.41) |
| **A(2)-tetrad** | 2-12 | -3.04 (1.18) | -0.82 (0.85) | -3.86 (1.45) |
|  | 2-17 | -3.04 (1.19) | -0.82 (0.86) | -3.86 (1.47) |
|  | 7-17 | -3.02 (1.19) | -0.83 (0.84) | -3.85 (1.46) |
|  | 7-12 | -3.03 (1.19) | -0.83 (0.84) | -3.86 (1.46) |
| **$\text{I}_m$(2)-tetrad** | 2-12 | 0.20 (1.31) | -1.08 (0.90) | -0.80 (1.59) |
|  | 2-17 | 1.38 (1.57) | -1.50 (0.62) | -0.12 (1.69) |
|  | 7-17 | 0.59 (1.32) | -1.40 (0.69) | -0.81 (1.56) |
|  | 7-12 | 0.36 (1.24) | -1.38 (0.70) | -1.02 (1.42) |
| **$\text{G}_m$(2)-tetrad** | 2-12 | -1.27 (2.72) | -1.92 (0.68) | -3.19 (2.80) |
|  | 2-17 | -1.25 (2.71) | -1.95 (0.67) | -3.20 (2.79) |
|  | 7-17<br>7-12 | -1.23 (2.71)<br>-1.24 (2.71) | -1.95 (0.67)<br>-1.94 (0.68) | -3.18 (2.79)<br>-3.18 (2.79) |
| **A<sub>m</sub>(2)-tetrad** | 2-12<br>2-17<br>7-17<br>7-12 | -3.16 (1.25)<br>-3.15 (1.25)<br>-3.19 (1.24)<br>-3.19 (1.24) | -1.17 (0.71)<br>-1.17 (0.71)<br>-1.17 (0.71)<br>-1.17 (0.71) | -4.33 (1.44)<br>-4.32 (1.44)<br>-4.36 (1.43)<br>-4.36 (1.43) |
| *Stacking energy between tetrads* |  |  |  |  |
| **G(1)-I(2)**<br>**I(2)-G(3)** |  | 19.54 (2.48)<br>20.09 (2.41) | -40.25 (1.27)<br>-39.48 (1.11) | -20.71 (2.78)<br>-19.39 (2.65) |
| **G(1)-G(2)**<br>**G(2)-G(3)** |  | 30.61 (3.23)<br>24.69 (3.57) | -43.51 (1.19)<br>-42.74 (1.21) | -12.9 (3.44)<br>-18.05 (3.76) |
| **G(1)-A(2)**<br>**A(2)-G(3)** |  | 4.09 (5.54)<br>11.34 (5.50) | -39.28 (1.29)<br>-38.72 (1.27) | -35.19 (5.69)<br>-27.38 (5.64) |
| **G(1)-I<sub>m</sub>(2)**<br>**I<sub>m</sub>(2)-G(3)** |  | 32.98 (2.94)<br>34.00 (1.69) | -38.81 (1.34)<br>-39.19 (1.23) | -5.83 (3.23)<br>-5.19 (2.09) |
| **G(1)-G<sub>m</sub>(2)**<br>**G<sub>m</sub>(2)-G(3)** |  | 38.62 (3.91)<br>37.69 (2.95) | -42.38 (1.26)<br>-42.06 (1.33) | -3.76 (3.34)<br>-4.37 (3.23) |
| **G(1)-A<sub>m</sub>(2)**<br>**A<sub>m</sub>(2)-G(3)** |  | 2.98 (3.02)<br>4.36 (2.54) | -38.16 (1.54)<br>-38.57 (1.26) | -35.18 (3.38)<br>-34.21 (2.84) |
| *Interaction energy between a tetrad and the proximal K<sup>+</sup> cation* |  |  |  |  |
| **I(2)-tetrad and K<sup>+</sup>**<br>**K24**<br>**K23** |  | -5.61 (3.04)<br>-68.54 (6.00) | -0.01 (0.06)<br>2.28 (1.94) | -5.62 (3.04)<br>-65.89 (6.30) |
| **G(2)-tetrad and K<sup>+</sup>**<br>**K24**<br>**K23** |  | -9.70 (11.62)<br>-88.08 (7.21) | -0.03 (0.21)<br>2.65 (2.16) | -9.73 (11.62)<br>-85.43 (7.52) |
| **A(2)-tetrad and K<sup>+</sup>**<br>**K24**<br>**K23** |  | -9.06 (3.71)<br>-26.69 (9.89) | -0.03 (0.13)<br>-0.12 (0.85) | -9.09 (3.71)<br>-26.81 (9.93) |
| **I<sub>m</sub>(2)-tetrad and K<sup>+</sup>**<br>**K24**<br>**K23** |  | -98.79 (6.06)<br>-115.35 (4.16) | 0.92 (1.45)<br>4.61 (2.03) | -97.87 (6.23)<br>-110.74 (4.63) |
| **G<sub>m</sub>(2)-tetrad and K<sup>+</sup>**<br>**K24**<br>**K23** |  | -105.47 (12.52)<br>-121.99 (6.83) | 1.23 (1.57)<br>5.41 (2.15) | -104.24 (12.61)<br>-116.58 (7.16) |

|  |  |  |  |
| --- | --- | --- | --- |
| **A<sub>m</sub>(2) tetrad and K<sup>+</sup>** |  |  |  |
| **K24** | -8.78 (3.53) | -0.03 (0.09) | -8.81 (3.53) |
| **K23** | -17.49 (2.98) | -0.38 (0.19) | -17.87 (2.99) |

**Table 2.** Comparison of Gibbs free energies (obtained from MM-GBSA analysis) of the quadruplexes for the fourth set of simulations. Standard deviations are shown within brackets.

|  | **Quad-I4** | **Quad-G4** | **Quad-A4** | **Quad-Im4** | **Quad-Gm4** | **Quad-Am4** |
| --- | --- | --- | --- | --- | --- | --- |
| G (kcal/mol) | -3207.53<br>(19.25) | -3755.97<br>(19.28) | -3535.52<br>(20.55) | -3192.20<br>(19.20) | -3703.10<br>(19.54) | -3260.27<br>(19.52) |

**Table 4.9.**
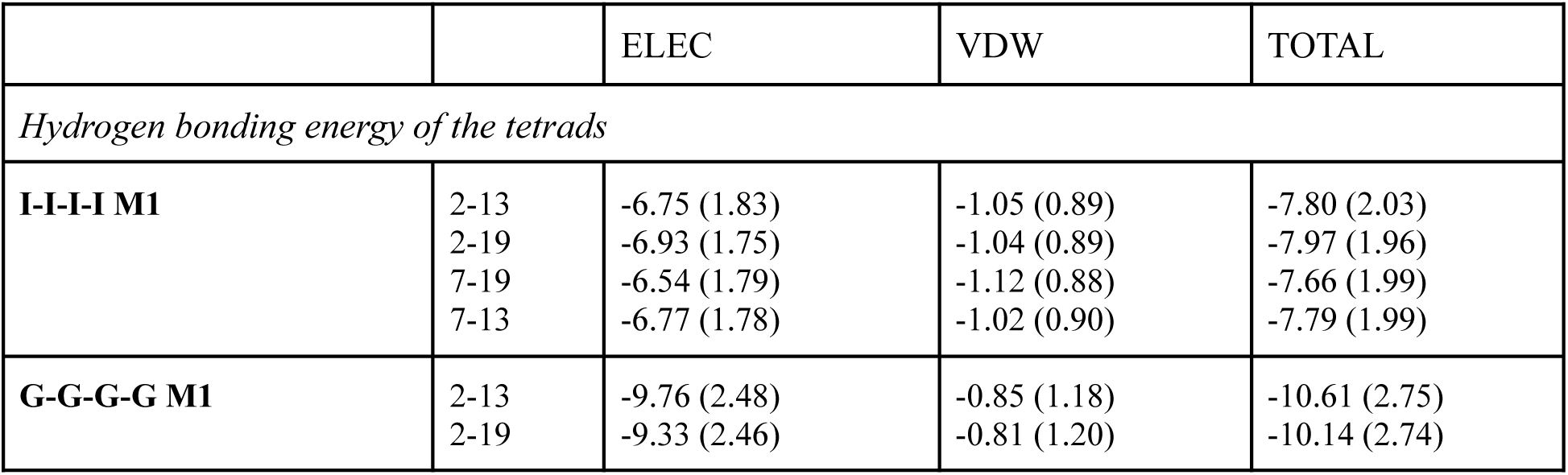
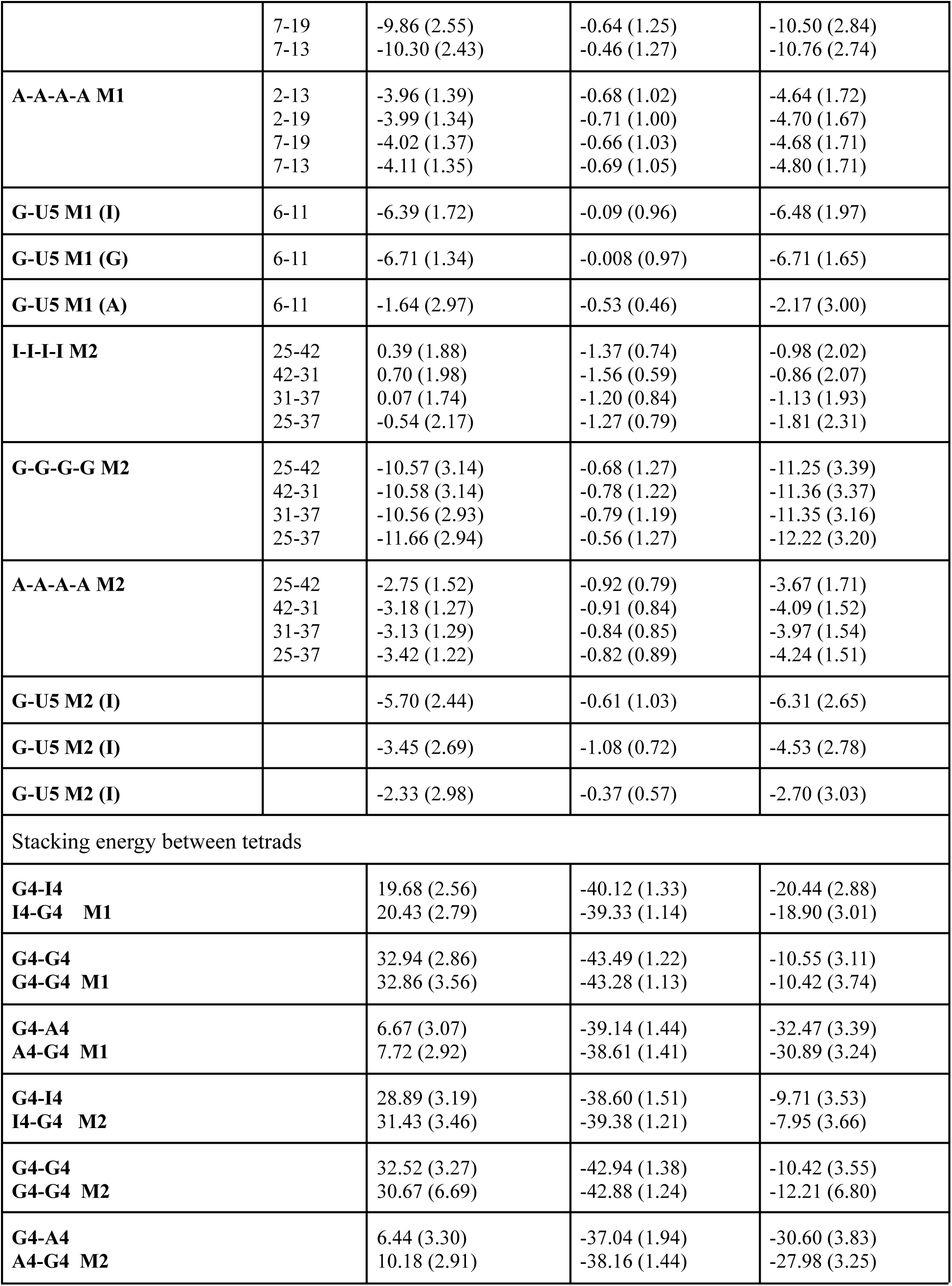
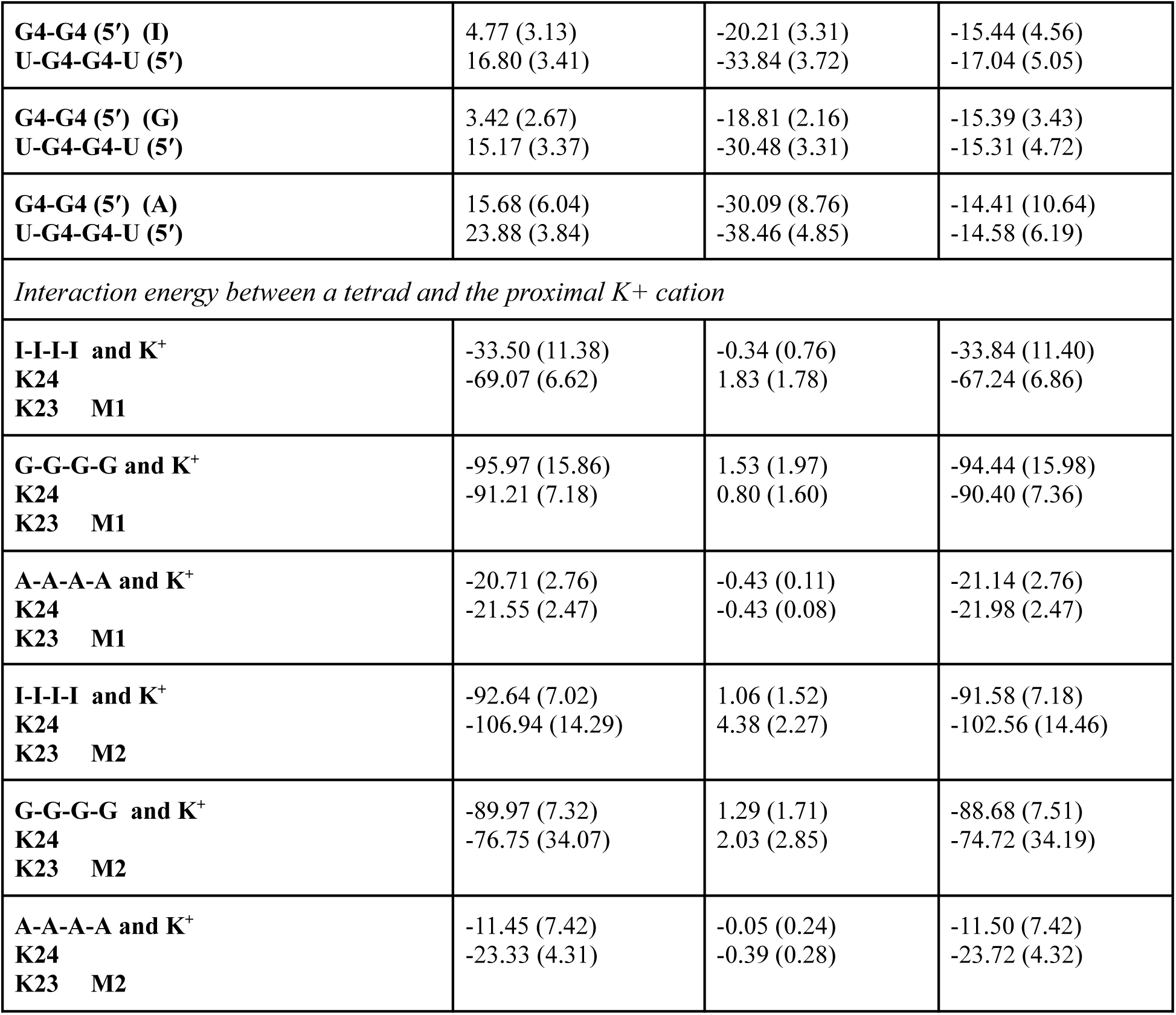
Interaction energies (kcal/mol) for hydrogen bonding, stacking, and ion interactions for the different purine tetrads at position 2 (for each of the monomers 1 and 2 (indicated as M1 and M2)) of the dimeric quadruplexes for the fourth set of simulations. Average and standard deviation (within brackets) values are reported.

|  |  | ELEC | VDW | TOTAL |
| --- | --- | --- | --- | --- |
| *Hydrogen bonding energy of the tetrads* |  |  |  |  |
| **I-I-I-I M1** | 2-13 | -6.75 (1.83) | -1.05 (0.89) | -7.80 (2.03) |
|  | 2-19 | -6.93 (1.75) | -1.04 (0.89) | -7.97 (1.96) |
|  | 7-19 | -6.54 (1.79) | -1.12 (0.88) | -7.66 (1.99) |
|  | 7-13 | -6.77 (1.78) | -1.02 (0.90) | -7.79 (1.99) |
| **G-G-G-G M1** | 2-13 | -9.76 (2.48) | -0.85 (1.18) | -10.61 (2.75) |
|  | 2-19 | -9.33 (2.46) | -0.81 (1.20) | -10.14 (2.74) |
|  | 7-19<br>7-13 | -9.86 (2.55)<br>-10.30 (2.43) | -0.64 (1.25)<br>-0.46 (1.27) | -10.50 (2.84)<br>-10.76 (2.74) |
| **A-A-A-A M1** | 2-13<br>2-19<br>7-19<br>7-13 | -3.96 (1.39)<br>-3.99 (1.34)<br>-4.02 (1.37)<br>-4.11 (1.35) | -0.68 (1.02)<br>-0.71 (1.00)<br>-0.66 (1.03)<br>-0.69 (1.05) | -4.64 (1.72)<br>-4.70 (1.67)<br>-4.68 (1.71)<br>-4.80 (1.71) |
| **G-U5 M1 (I)** | 6-11 | -6.39 (1.72) | -0.09 (0.96) | -6.48 (1.97) |
| **G-U5 M1 (G)** | 6-11 | -6.71 (1.34) | -0.008 (0.97) | -6.71 (1.65) |
| **G-U5 M1 (A)** | 6-11 | -1.64 (2.97) | -0.53 (0.46) | -2.17 (3.00) |
| **I-I-I-I M2** | 25-42<br>42-31<br>31-37<br>25-37 | 0.39 (1.88)<br>0.70 (1.98)<br>0.07 (1.74)<br>-0.54 (2.17) | -1.37 (0.74)<br>-1.56 (0.59)<br>-1.20 (0.84)<br>-1.27 (0.79) | -0.98 (2.02)<br>-0.86 (2.07)<br>-1.13 (1.93)<br>-1.81 (2.31) |
| **G-G-G-G M2** | 25-42<br>42-31<br>31-37<br>25-37 | -10.57 (3.14)<br>-10.58 (3.14)<br>-10.56 (2.93)<br>-11.66 (2.94) | -0.68 (1.27)<br>-0.78 (1.22)<br>-0.79 (1.19)<br>-0.56 (1.27) | -11.25 (3.39)<br>-11.36 (3.37)<br>-11.35 (3.16)<br>-12.22 (3.20) |
| **A-A-A-A M2** | 25-42<br>42-31<br>31-37<br>25-37 | -2.75 (1.52)<br>-3.18 (1.27)<br>-3.13 (1.29)<br>-3.42 (1.22) | -0.92 (0.79)<br>-0.91 (0.84)<br>-0.84 (0.85)<br>-0.82 (0.89) | -3.67 (1.71)<br>-4.09 (1.52)<br>-3.97 (1.54)<br>-4.24 (1.51) |
| **G-U5 M2 (I)** |  | -5.70 (2.44) | -0.61 (1.03) | -6.31 (2.65) |
| **G-U5 M2 (I)** |  | -3.45 (2.69) | -1.08 (0.72) | -4.53 (2.78) |
| **G-U5 M2 (I)** |  | -2.33 (2.98) | -0.37 (0.57) | -2.70 (3.03) |
| Stacking energy between tetrads |  |  |  |  |
| **G4-I4**<br>**I4-G4 M1** |  | 19.68 (2.56)<br>20.43 (2.79) | -40.12 (1.33)<br>-39.33 (1.14) | -20.44 (2.88)<br>-18.90 (3.01) |
| **G4-G4**<br>**G4-G4 M1** |  | 32.94 (2.86)<br>32.86 (3.56) | -43.49 (1.22)<br>-43.28 (1.13) | -10.55 (3.11)<br>-10.42 (3.74) |
| **G4-A4**<br>**A4-G4 M1** |  | 6.67 (3.07)<br>7.72 (2.92) | -39.14 (1.44)<br>-38.61 (1.41) | -32.47 (3.39)<br>-30.89 (3.24) |
| **G4-I4**<br>**I4-G4 M2** |  | 28.89 (3.19)<br>31.43 (3.46) | -38.60 (1.51)<br>-39.38 (1.21) | -9.71 (3.53)<br>-7.95 (3.66) |
| **G4-G4**<br>**G4-G4 M2** |  | 32.52 (3.27)<br>30.67 (6.69) | -42.94 (1.38)<br>-42.88 (1.24) | -10.42 (3.55)<br>-12.21 (6.80) |
| **G4-A4**<br>**A4-G4 M2** |  | 6.44 (3.30)<br>10.18 (2.91) | -37.04 (1.94)<br>-38.16 (1.44) | -30.60 (3.83)<br>-27.98 (3.25) |

|  |  |  |  |
| --- | --- | --- | --- |
| **G4-G4 (5') (I)**<br>**U-G4-G4-U (5')** | 4.77 (3.13)<br>16.80 (3.41) | -20.21 (3.31)<br>-33.84 (3.72) | -15.44 (4.56)<br>-17.04 (5.05) |
| **G4-G4 (5') (G)**<br>**U-G4-G4-U (5')** | 3.42 (2.67)<br>15.17 (3.37) | -18.81 (2.16)<br>-30.48 (3.31) | -15.39 (3.43)<br>-15.31 (4.72) |
| **G4-G4 (5') (A)**<br>**U-G4-G4-U (5')** | 15.68 (6.04)<br>23.88 (3.84) | -30.09 (8.76)<br>-38.46 (4.85) | -14.41 (10.64)<br>-14.58 (6.19) |
| *Interaction energy between a tetrad and the proximal K<sup>+</sup> cation* |  |  |  |
| **I-I-I-I and K<sup>+</sup>**<br>**K24**<br>**K23 M1** | -33.50 (11.38)<br>-69.07 (6.62) | -0.34 (0.76)<br>1.83 (1.78) | -33.84 (11.40)<br>-67.24 (6.86) |
| **G-G-G-G and K<sup>+</sup>**<br>**K24**<br>**K23 M1** | -95.97 (15.86)<br>-91.21 (7.18) | 1.53 (1.97)<br>0.80 (1.60) | -94.44 (15.98)<br>-90.40 (7.36) |
| **A-A-A-A and K<sup>+</sup>**<br>**K24**<br>**K23 M1** | -20.71 (2.76)<br>-21.55 (2.47) | -0.43 (0.11)<br>-0.43 (0.08) | -21.14 (2.76)<br>-21.98 (2.47) |
| **I-I-I-I and K<sup>+</sup>**<br>**K24**<br>**K23 M2** | -92.64 (7.02)<br>-106.94 (14.29) | 1.06 (1.52)<br>4.38 (2.27) | -91.58 (7.18)<br>-102.56 (14.46) |
| **G-G-G-G and K<sup>+</sup>**<br>**K24**<br>**K23 M2** | -89.97 (7.32)<br>-76.75 (34.07) | 1.29 (1.71)<br>2.03 (2.85) | -88.68 (7.51)<br>-74.72 (34.19) |
| **A-A-A-A and K<sup>+</sup>**<br>**K24**<br>**K23 M2** | -11.45 (7.42)<br>-23.33 (4.31) | -0.05 (0.24)<br>-0.39 (0.28) | -11.50 (7.42)<br>-23.72 (4.32) |

For the different simulation sets for each monomeric quadruplex studied, we observed the rearrangement of channel K^+^ ions, and expulsion of channel K^+^ ions vertically from the 3′-end or the 5′-end or horizontally from the side (only in one simulation) (**Figures S9-32**). The rearrangements/expulsions of the channel K^+^ ions could also be realized from the ion interaction energies around the tetrads at position 2(**Table 1, S1**) and the plots of K^+^-O6, K^+^-N6, K^+^-O4 distances for I/G, A and U tetrads respectively. The interaction energies of A-tetrad with the nearest K^+^ ions in the channel were generally low due to the presence of the N(6)H2 group as reported in Pan et al. [29] (**Table 1, S1**). In some simulations, the K^+^ ions from the solvent were exchanged for the channel ion at the 3′-end or the 5′-end (**Figure 5, S20,24,28,32**). The rearrangements of the metal ions in the channel have also been studied earlier by Andrałojć et al [56]. As reported by Andrałojć et al [56], the formation of a K^+^ ion pocket around the reversed 3′-U tetrad (e.g. between G4-O2′ and U5-O2′) was observed in our study as well (**Figures 5, S33)**. The occupancy of the K^+^ ions between G4-O2′ and U5-O2′ was observed to be reduced for the quadruplexes containing the 2′-O-methylated purine tetrads at position 2 (**Figure S33**).

**Figure 5.**
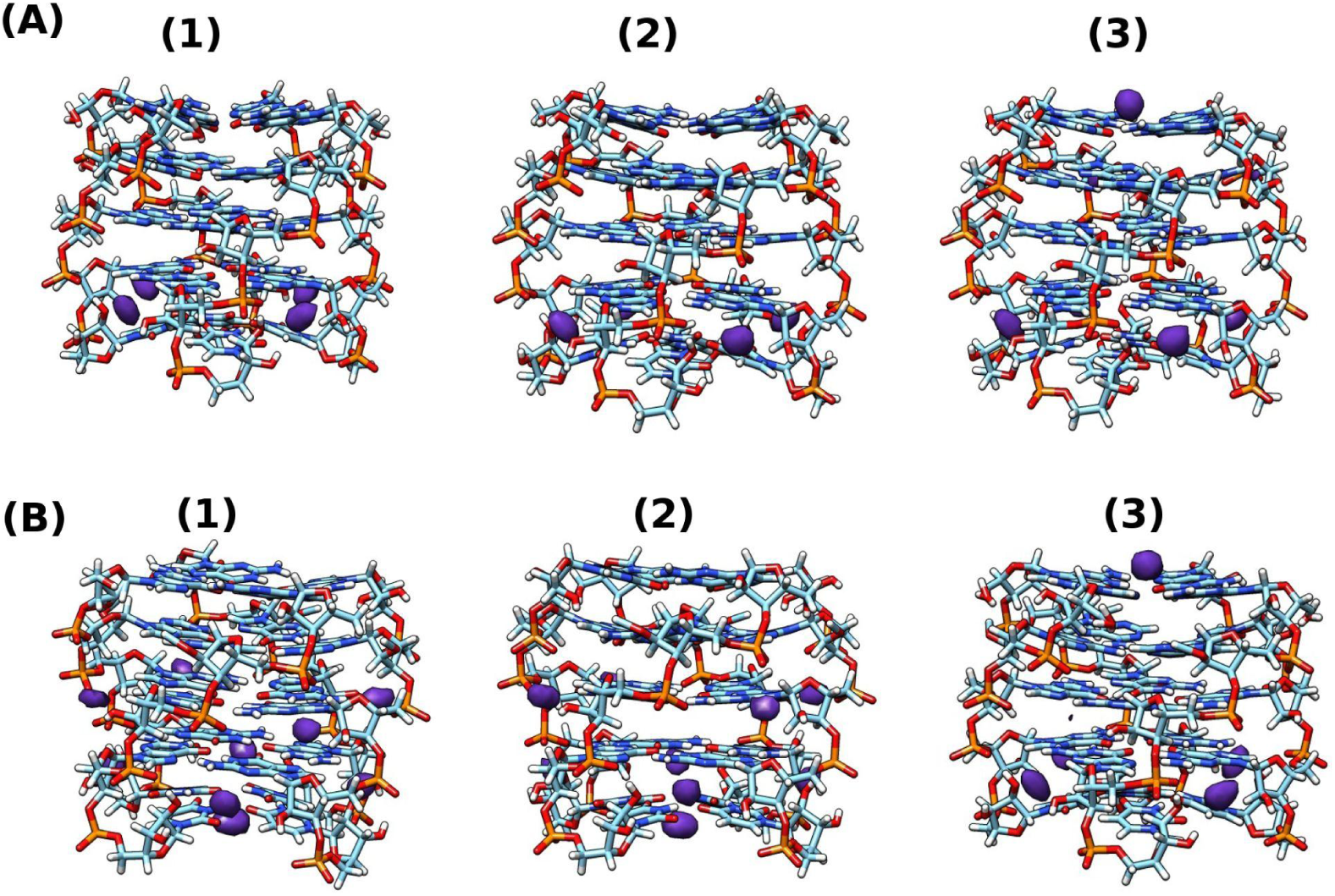
Occupancy maps of K^+^ ions for (A) unmethylated (B) methylated quadruplexes (5′ (top) and 3′ (bottom)) respectively (1) Quad-I/Quad-Im tetrad; (2) Quad-G/Quad-Gm and (3) Quad-A/Quad-Am tetrad. K^+^ ion binding pocket is observed around the reversed U(5)-tetrads and also around the G(4) tetrads for the fourth set of simulations.

Interestingly, we also observed the formation of such an ion pocket between the G-tetrads at positions 3 and 4 (although with much lower densities of K^+^ compared to those around the reversed U-tetrad) for the Quad-I_m_4 and Quad-G_m_4 (**Figures 5, S34)**.

For the assessment of the influence of the A-to-I or G-to-I substitution and the rearrangement of channel K^+^ ions on base non-planarity with G-quadruplex structure, we compared the pyramidalization of the glycosidic nitrogen (N9) of the residues constituting the three middle tetrads-(2-4) within the monomeric quadruplexes. The N9-pyramidalization was measured based on the values of the *κ*′ (C4−N9−C1′−C8 − 180°) improper dihedral angles. From our MD simulations, we observed a similar trend in the N9-pyramidalization for the I(2)-, G(2)- and A(2)-tetrads except for simulation set 1 in which the residues in the G(2) tetrad had much lower extent of N9-pyramidalization (**Table S6; Figures S35-37**). 2′-O-methylation of the tetrad-2 residues resulted in significant increase in the extent of N9-pyramidalization especially for the I-terad (**Table S6; Figures S38-40**). The presence of different (unmethylated/2′-O-methylated tetrads at position 2 of the monomeric quadruplexes also influenced the N9-pyramidalization of the tetrads at position 3 (**Table S6**). The rearrangements and/or expulsions of channel K^+^ ions also resulted in changes in the N9-pyramidalization, e.g. in simulation set 1 for Quad-G4, the K23 channel ion was not involved in the rearrangement but in the other simulation sets, with the expulsion of K24 channel ion, the position of K23 was changed which might have resulted in the difference in the extent of non-planarity of the G(2) tetrad residues between simulation set 1 with the other simulation sets **(Figure S41)**.

Analysis of the six backbone dihedrals (**α, β**, *γ*, δ, ε, and ζ) of the constituent residues of the I-, G-, and A- tetrads at position 2 within monomeric quadruplexes revealed the similarity of the distribution of the backbone dihedral angles (**Figure S42**). The addition of 2′-O-methylation was not observed to alter the distributions of the backbone dihedrals for the residues within G_m_-, and A_m_-tetrads from what was observed for the unmethylated counterparts. However, for the I_m_-tetrad, although the distributions of the backbone dihedrals were similar for the first three simulation sets where I-tetrad with N1(H)-O6 geometry was sampled predominantly, the distributions were altered in the fourth simulation set, suggesting the lower stability and high flexibility of the alternate tetrad geometry (N1(H)-N7) observed for the I_m_-tetrad (**Figure S43**).

The local base pair and base pair step parameters for the HG base pairs within the tetrads of the studied monomeric quadruplexes presented in **Table S7** (data for the first simulation set is shown) indicated the general structural stability of the quadruplexes during the simulations.

As we also observed alteration of the stability of the G-tetrad geometry from the calculation of the hydrogen bonding energies upon 2′-O-methylation of G, to better explore the conformational rearrangements due to the presence of the G_m_-tetrad, we further compared some other parameters following Tsvetkov et al. [54]. The time evolution of the twist angle (ϴ) revealed increased fluctuations of the twist angle (ϴ) between the tetrads 1 and 2 and 2 and 3, respectively, for the monomeric quadruplex containing G_m_-tetrad at position 2 (Quad-G_m_4) compared to the unmodified quadruplex (**Figures S44-45**). Time evolution of the distances between the tetrad-COMs did not indicate a substantial difference in this parameter upon the inclusion of the 2′-O-methylation (**Figures S46-47**). For the description of the integrity of the tetrads, the time evolution of the distances between the tetrad-COM and COM of the constituent individual G/G_m_ residues was plotted for each tetrad (for G/G_m_-tetrads). A substantial difference in the fluctuation of these distances was not observed between Quad-G4 and Quad-G_m_4, particularly in the last three simulation sets (**Figures S48-49**). However, in the first simulation set, the distances between the tetrad-COM and COM of the individual G residues for the first tetrad were observed to fluctuate within a broader range of distance values for both the monomeric quadruplexes (Quad-G4 and Quad-G_m_4). Time evolution plots of the angles (ϕ) between normals to the G/G_m_-planes and the tetrad-axis were generated to estimate the changes in the planarity of the tetrads (**Figures S50-51**). The range of fluctuations of this parameter was observed to be overall similar for the last three simulation sets, while for the first simulation set, the pattern varied especially during the first 750 ns of the simulation for the 5′-G-tetrad indicating alteration of the planarity of this tetrad. The time evolution plots of the angles (θ) between G/G_m_-pairs (for the characterization of lengthwise tetrad bending) and G/G_m_-triads (for the characterization of diagonal tetrad bending) revealed that during the simulations, the lengthwise tetrad bending patterns varied differently for the four simulations sets for a particular quadruplex and also between the quadruplexes (Quad-G4 and Quad-G_m_4) (**Figures 52-55**). The diagonal tetrad bending patterns were overall similar for the Quad-G4 and Quad-G_m_4, especially in the last three simulation sets (**Figures 54-55**).

### Consequences of the presence of m^6^A-methylation in G-quadruplex structure

From the simulations of the monomeric quadruplex containing m^6^A-tetrad at position 2 (Quad-m^6^A4), the N6-H---N3 geometry of the m^6^A-tetrad was observed to be highly predominant (**Figure S56; Table S8-9**). The hydrogen bonding energy of the m^6^A-tetrad was observed to be ∼ -26.4 kcal/mol in each of the simulation sets but the frequency of the N6-H---N3 hydrogen bonding interaction was low (∼ 10%) (**Table S8-9**). The stacking energies corresponding to the interactions of the m^6^A-tetrad with its neighboring G-tetrads were more favorable compared to what was observed for the A-tetrad within a similar quadruplex (**Table S8**). However, the Gibbs free energy for Quad-m^6^A4 was found to be lower than what was observed for Quad-A4. Similar to what was observed for the A-tetrad, the interactions of the m^6^A-tetrad with the channel K^+^ ions were very weak (**Table S8**). Water bridging interactions were not observed between the m^6^A HG-pairs of the m^6^A-tetrad as a result of the N6-methylation (**Figure S56**). The extents of N9-pyramidalization for the tetrads at positions 2, 3 and especially for the tetrad at position 4 were observed to be different from what was observed for Quad-A4 and the other monomeric quadruplexes (**Table S10**).

### Properties of dimeric quadruplexes

#### Consequences of the presence of inosine in inter-(G-)quadruplex interactions

From the simulation of dimeric quadruplexes, we could obtain further ideas about the tetrad conformations, the influence of channel ions in the conformations of the tetrads, and the dynamics and properties of the interaction between the monomer units (through their respective 5′ end U-(G-G-G-G) pentads). The stacking arrangement of the I-tetrad in monomer 1 (in monomer 2 the arrangement is identical) and that of the 5′-pentads are shown in **Figure 6**.

**Figure 6.**
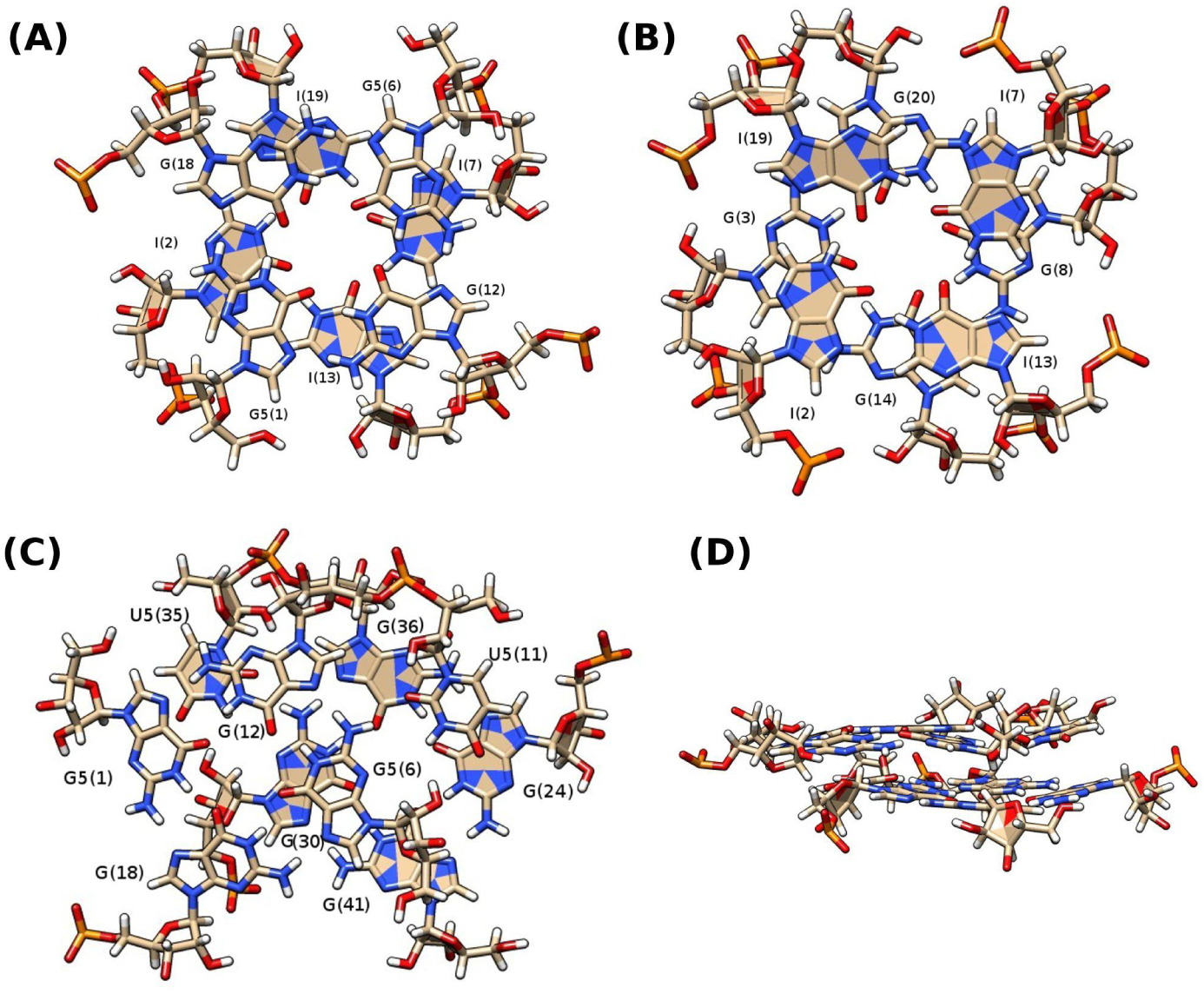
(A) Stacked bases of G-tetrad at position 1 and I-tetrad at position 2 (B) Stacked bases of I-tetrad at position 2 and G-tetrad at position 3 for quadruplex monomer 1; (C) top view and (D) side view of stacked bases of U-(G-G-G-G)-pentads of the two monomers.

The N1(H)-N7 tetrad geometry (with a much less favorable hydrogen bonding interaction energy and low occurrence frequency of the hydrogen bonding (N1-H1---N7) interaction, compared to the N1(H)-O6 geometry) was largely predominant for the I(2)-tetrads in the two quadruplex monomers for the first and second simulation sets. while for the third set, the N1(H)-O6 geometry was the most populated and the N1(H)-N7 geometry was observed to be in a smaller population of conformers for monomer1 and even smaller population for monomer 2. Interestingly, in simulation set 4, N1(H)-O6 geometry was predominant for monomer 1 and the N1(H)-N7 geometry was preferentially sampled for monomer 2 (**Figures 7, S57; Table S11,12**). The differences in the tetrad geometries were reflected in the hydrogen bonding interaction energies (**Table 3, S11**). For the G(2) tetrads, geometry with N1(H)-O6 and N2(H)-N7 hydrogen bonds, was preferentially sampled for each of the monomers (**Figure S57; Table S12**). However, for the dimeric quadruplexes, the hydrogen bonding energies for the G(2) tetrads were observed to be lower compared to what was observed in the isolated monomer which might be due to the low occurrence frequency of the N1(H)-O6 hydrogen bonding interaction (**Table S11,12)**. The N6(H)-N1 geometry was predominant for the A-tetrads in both the monomers in the first, third, and fourth simulation sets, but in the second set, interestingly, the N6(H)-N3 geometry [58,64] was predominantly sampled for the A-tetrad in monomer 2 after the first ∼220 ns of the simulation (**Figure S57**). The hydrogen bonding interaction energies of the A-tetrads in the dimeric quadruplex were similar to what was observed in the isolated monomer. The stacking interaction energies between the tetrad 2 with tetrad 1 and tetrad 3 for each monomer were observed to be similar as was observed in the isolated monomer for the respective quadruplexes (Quad-I4, Quad-G4, and Quad-A4). A(2)-tetrad formed more favorable stacking interactions with the upper and lower G tetrads, compared to those formed with I(2)/G(2)-tetrads in each monomer (**Table S11)**. Highly stable signature hydrogen bond interaction between G-OP2 and 3′U-O2′ was observed in the constituent monomeric quadruplexes with reversed-3′U-tetrads within each of the quadruplex dimers (**Table S13**).

**Figure 7.**
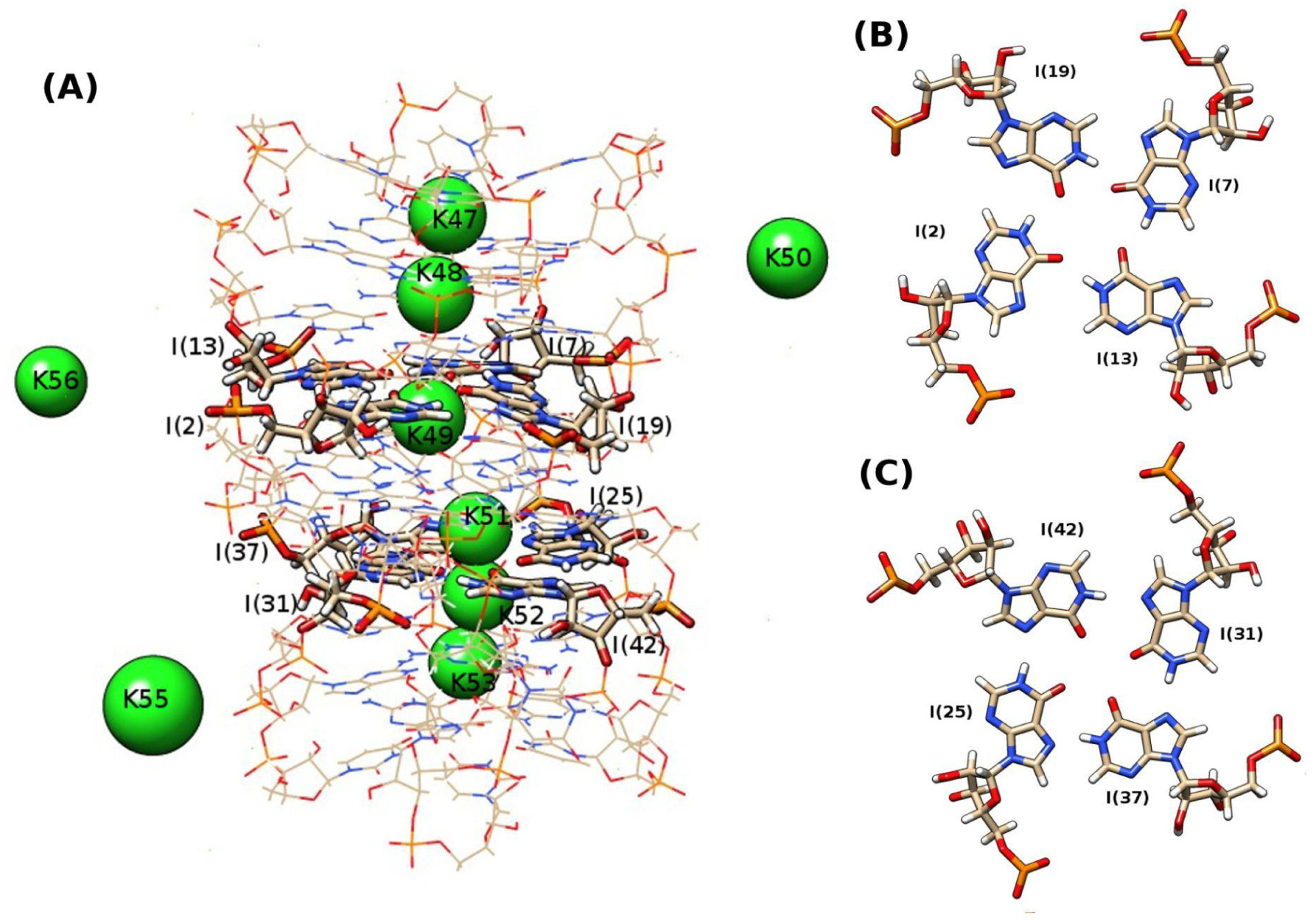
(A) Snapshot of Quad-I4 with monomer 1 (M1) (top) and monomer 2 (M2) (bottom) showing that different arrangements of the channel ions in the two quadruplex monomers might be associated with the differences in the tetrad geometries. (B) N1(H)-O6 tetrad geometry for I-tetrad in monomer 1 and (C) N1(H)-N7 tetrad geometry for I-tetrad in monomer 2.

Different (at position 2 of each monomer) purine tetrad geometries were additionally stabilized by water-bridging interactions as was observed in the isolated monomeric quadruplexes (**Table S14A**). However, the occurrences of the water-bridging interactions between N3 of U and N3 of the closest G residue in the U*(G-G-G-G) pentad [29] were found to be negligible from our simulated ensembles (**Table S14B**).

The channel K^+^ ion rearrangements and expulsion (from the 3′-ends in the case of the dimers as the 5′-ends were involved in the quadruplex-quadruplex interaction) were also observed from the simulation of the quadruplex dimers. The influence of the channel ion rearrangements on the conformational changes in the tetrads was more prominent from the simulation of the dimers. As can be seen from figure 7, the different arrangements of the channel K^+^ ions in the two monomers facilitated the sampling of two different geometries for the I-tetrad. The altered arrangement of the channel K^+^ ions was also observed to facilitate the alternate A-tetrad geometries within the two monomers (**Figure S57**). Formation of a K^+^ ion pocket around the reversed 3′-U tetrads (e.g. between G4-O2′ and U5-O2′) was observed for each of the monomers of the dimeric quadruplexes (**Figures 8, S58**).

**Figure 8.**
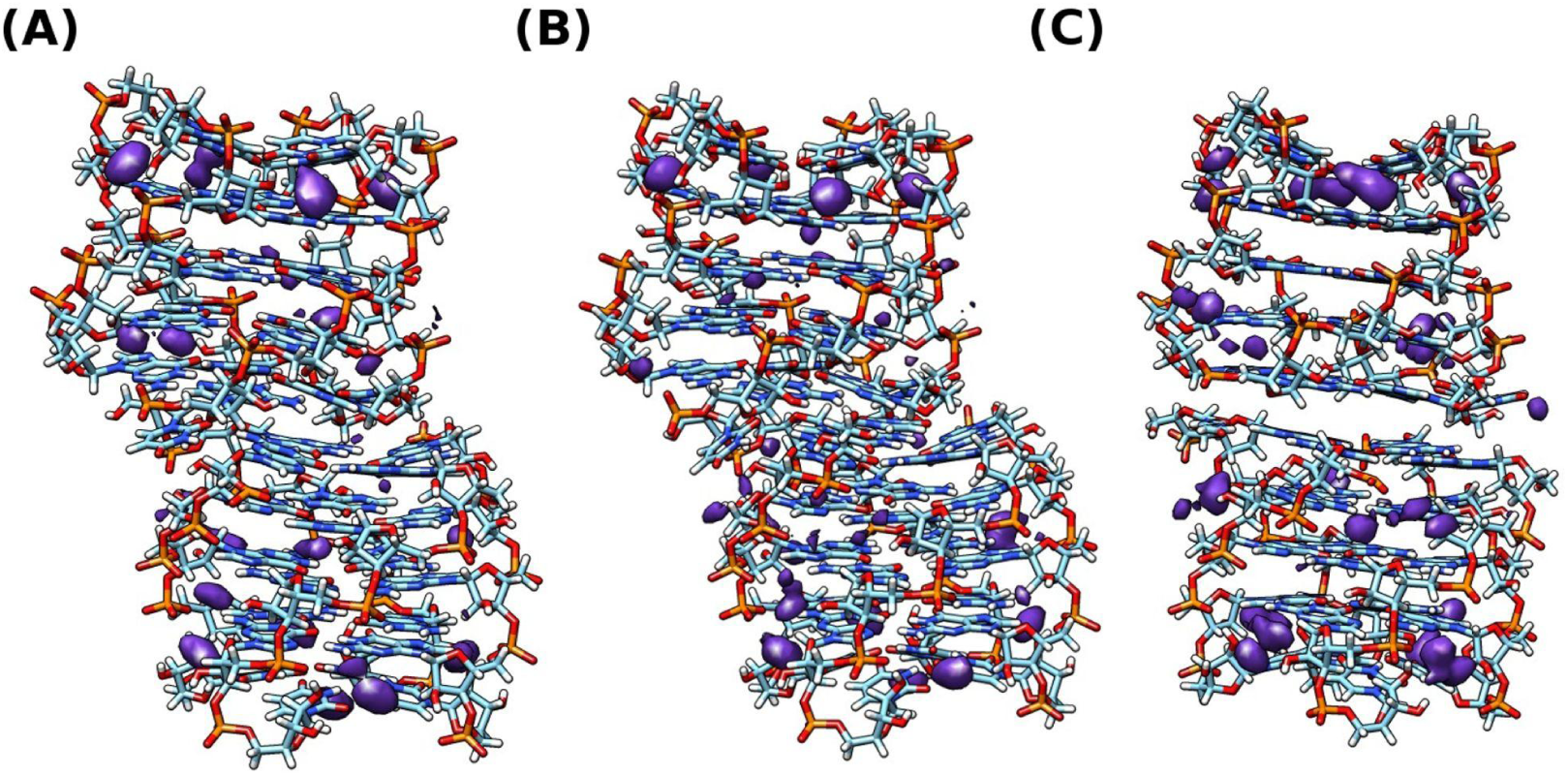
Occupancy maps of K^+^ ions for the quadruplex dimers in this study from the fourth set of simulations (A) Quad-I4 (containing I(2) tetrads); (B) Quad-G4 (containing G(2) tetrads) and (C) Quad-A4 (containing A(2) tetrads). K+ ion binding pocket is observed around the reversed U(5)-tetrads for each monomer.

Comparison of the (total) Gibbs free energies of the monomers and the whole dimers from MM-GBSA analyses revealed the following trend in stabilities: Quad-I4 (dimer/monomers) < Quad-A4 (dimer/monomers) < Quad-G4 (dimer/monomers) (**Table S15A**) and the observations were similar for each simulation set. However, the values of binding free energies (ΔG_bind_) showed more favorable quadruplex-quadruplex interactions for Quad-I4 and Quad-G4 compared to Quad-A4 in the first three simulation sets while in simulation set 4 the interaction for Quad-A4 was found to be comparatively more favorable (**Table S15B**).

## Conclusion

In this study, we have reported the structural and energetic consequences of the presence of the different purine tetrads, i.e. I-, G- or A- tetrad, 2′-O-methylated tetrads (i.e. I_m_-, G_m_-, and A_m_-tetrad) and N6-methylated A-tetrad (i.e. m^6^A-tetrad) respectively at a similar position within the quadruplex structure (monomer) based on molecular dynamic simulations. Besides that, our study also showcases the hydration patterns around the different purine tetrads, the influence of the dynamics of the channel (K^+^) ions on the tetrad geometries, and the presence of ion binding pockets for RNA quadruplex structures. Additionally, the properties of dimeric quadruplexes containing I-, G- or A- tetrads neighboring the 5′ stacked U*(G-G-G-G) pentads within the constituent monomeric quadruplexes and their effect on the inter-quadruplex interaction are also reported. The overall stability of the monomeric and dimeric quadruplexes containing I-tetrad/s further suggests the possibility of the formation of stable RNA quadruplex structures with inosine tetrad (or mixed GI-tetrads) as a result of A to I RNA editing. However, the structural and thermodynamic stability of rG4 structures containing I- or mixed GI-tetrad/s can be expected to depend on the sequence context and position of the modifications. For the quadruplex sequence context studied in the present work, 2′-O methylation of the I- and G-tetrads was not observed to stabilize the tetrads but this modification resulted in slight stabilization of the hydrogen bonding interactions and tetrad-stacking interactions for the the A-tetrad. Interestingly, the N6-methylation of adenosine also stabilized the A-tetrad geometry compared to the unmodified counterpart. However, further experimental and theoretical studies would be required for a better understanding of the structural, thermodynamic, and energetic stabilities of RNA quadruplexes containing I-tetrads, mixed GI-tetrads, or other purine tetrads at different positions within such quadruplexes. Considering the importance of RNA quadruplex structures in several cellular processes and emerging potentials of quadruplex-quadruplex interactions/multimeric quadruplexes as therapeutic targets, our results might be helpful for future studies involving RNA quadruplexes and understanding the functions of purine tetrads other than G-tetrad within quadruplexes.

## Author Contributions

Concept: ND. Design of the study: ND and AL. Calculations and analyses: ND. ND wrote the manuscript. The manuscript has been reviewed by all the authors.

## ACKNOWLEDGEMENTS

N.D. acknowledges support from the DST-INSPIRE Senior Research Fellowship (DST/INSPIRE Fellowship/2018/IF180895). Some of the simulations and calculations were carried out at the Poznan Supercomputing and Networking Center. The authors would like to thank Prof. Dhananjay Bhattacharya for his suggestions on the calculation of structural parameters for the Hoogsteen base pair and base pair steps.

## DATA AVAILABILITY

All data is available. Additional data which support the findings in this study will be freely available upon request from the corresponding author.

## SUPPORTING INFORMATION

### Supporting Tables

**Table S1.** Interaction energies for the different purine tetrads at position 2 (from the 5′-direction) of the studied quadruplex monomers for the first three simulation sets. **Table S2.** Frequencies of the Hoogsteen (HG) H-bonds between the bases which constitute the different tetrads of the studied quadruplex monomers for the four simulation sets. **Table S3.** Frequencies of the signature H-bond between G-OP2 and 3′U-O2′ for the studied quadruplex monomers for the four simulation sets. **Table S4.** Frequencies of the water bridging interactions observed for the different purine tetrads at position 2 of the studied quadruplex monomers for the four simulation sets. **Table S5.** Comparison of (total) Gibbs free energies (from MM-GBSA analysis) of the studied quadruplex monomers for the first three simulation sets. **Table S6.** Comparison of N9-pyramidalization (based on Kappa’ (*κ*’) dihedral angles) for the tetrads at positions-2, 3, and 4 respectively of the studied monomeric quadruplexes for the four simulation sets. **Table S7.** Structural base pair and base pair step parameters for the Hoogsteen (HG) pairs in the different tetrads for the monomeric quadruplexes containing different purine tetrads at position 2 respectively, for the first simulation set. **Table S8.** Interaction energies and for the m^6^A-tetrad (at position 2) and comparison of Gibbs free energies (from MM-GBSA analysis) of the studied quadruplex monomers containing m^6^A-tetrad at position 2 for the three simulation sets. **Table S9.** Frequencies of the Hoogsteen (HG) H-bonds between the bases which constitute the different tetrads of the studied quadruplex monomer containing m^6^A-tetrad at position 2, for the three simulation sets. **Table S10.** Comparison of N9-pyramidalization (based on Kappa’ (*κ*’) dihedral angles) for the tetrads at position 2, 3, and 4 respectively of the studied quadruplex monomer containing m^6^A-tetrad for the three simulation sets. **Table S11.** Interaction energies for the different purine tetrads at position 2 (from the 5′-direction) of the studied quadruplex dimers for the four simulation sets. **Table S12.** Comparison of binding free energies (of the constituent quadruplex monomers) and (total) Gibbs free energies (from MM-GBSA analysis) of the quadruplex dimers and the constituent quadruplex monomers, for the studied quadruplex dimers for the four simulation sets.**Table S13.** Frequencies of the Hoogsteen (HG) hydrogen bonds between the bases within different tetrads of the studied dimeric quadruplexes for the four simulation sets. **Table S14.** Frequencies of the signature H-bond between G-OP2 and 3′U-O2′ within the constituent quadruplex monomers of the studied dimeric quadruplexes for the four simulation sets. **Table S15.** Frequencies of the water bridging interactions observed for the different purine tetrads at position 2 and those of the U-G water bridging (in U*(G-G-G-G) pentad) interactions, within the constituent quadruplex monomers of the studied dimeric quadruplexes for the four simulation sets. **Table S16.** Comparison of N9-pyramidalization (based on Kappa’ (*κ*’) dihedral angles) for the tetrads at position 2 within the constituent quadruplex monomers of the studied dimeric quadruplexes for the four simulation sets.

### Supporting figures

**Figure S1.** RMSD of heavy atoms of the nucleotides of the first three tetrads (in the 5′) for the monomeric quadruplexes Quad-I4, Quad-G4, and Quad-A4 for the four simulation sets. **Figure S2**. RMSF of heavy atoms of the nucleotides for the monomeric quadruplexes Quad-I4, Quad-G4, and Quad-A4 for the four simulation sets. **Figure S3**. Stacking interaction energies for the (A) G4(1)-X4(2) tetrad step and (B) that for the X4(1)-G4(2) tetrad step (where X indicates I/G/A) for the monomeric quadruplexes Quad-I4, Quad-G4, and Quad-A4 for the four simulation sets. **Figure S4**. RMSD of heavy atoms of the nucleotides of the first three tetrads (in the 5′) for the monomeric quadruplexes Quad-I_m_4, Quad-G_m_4, and Quad-A_m_4 for the four simulation sets. **Figure S5**. RMSF of heavy atoms of the nucleotides for the monomeric quadruplexes Quad-I_m_4, Quad-G_m_4, and Quad-A_m_4 for the four simulation sets. **Figure S3**. Stacking interaction energies for the (A) G4(1)-X4(2) tetrad step and (B) that for the X4(1)-G4(2) tetrad step (where X indicates I_m_/G_m_/A_m_) for the monomeric quadruplexes Quad-I4, Quad-G4 and Quad-A4 for the four simulation sets. **Figure S7.** Water occupancy maps for I-, G-, and A-tetrads around the average structures for the first three simulation sets. **Figure S8.** Water occupancy maps for I_m_-, G_m_-, and A_m_-tetrads around the average structures for the first three simulation sets. **Figure S9-32.** Time evolution of distances between channel K^+^ ions and O6 atom of G(1); O6 atom of I/G/I_m_/G_m_(2), N6 atom of A/A_m_(2); O6 atom of G(3), G(4) and O4 atom of U(5) for the monomeric quadruplexes Quad-I4, Quad-G4, Quad-A4 Quad-I_m_4, Quad-G_m_4, and Quad-A_m_4 respectively for the four simulation sets. **Figure S33.** RDF of K^+^ ions around the geometric center of G-O2′ and U-O2′(for G4-U20, G14-U19, G9-U15, G19-U10 for the monomeric quadruplexes for the fourth set of simulations. **Figure S34.** RDF of K^+^ ions around the geometric center of Gn-O2′ and Gn-1-O2′ (for G4-G3, G9-G8, G14-G13, G19-G18 for the monomeric quadruplexes for the fourth set of simulations. **Figure S35-S40.** 2D-scatter plots of *X* vs *k*’ dihedrals for the residues in the tetrads at positions 2,3, and 4 for the monomeric quadruplexes for the fourth simulation set. **Figure S41.** Snapshot of G(2)-tetrad within Quad-G4 showing the different positions of the channel K23 ion in the G4-channel in the first and second simulation sets resulting in different N9-pyramidalization of the G(2) tetrad residues. **Figure S42-S43.** Distribution of the six backbone dihedral angles for the residues in the I-, G-, and A- tetrads at position 2 for the monomeric quadruplexes corresponding to the four simulation sets. **Figure S44-S45.** Time evolution of the twist angles between the tetrads for the monomeric quadruplexes Quad-G4 and Quad-Gm4 respectively corresponding to the four simulation sets. **Figure S46-S47.** Time evolution of the distance between the COMs of the tetrads respectively for the monomeric quadruplexes Quad-G4 and Quad-Gm4 respectively corresponding to the four simulation sets. **Figure S48-S49.** Time evolution of the distance between the tetrad-COM and COM of each of the constituent residues for the monomeric quadruplexes Quad-G4 and Quad-G_m_4 respectively corresponding to the four simulation sets. **Figure S50-S51.** Time evolution of the angles between the normals to the G(/G_m_)-planes in a tetrad and the axis of the tetrad and COM of each of the constituent residues for the monomeric quadruplexes Quad-G4 and Quad-G_m_4 respectively corresponding to the four simulation sets. **Figure S52-S53.** Time evolution of the angles between nominal planes formed by the G(/G_m_)-pairs for the monomeric quadruplexes Quad-G4 and Quad-G_m_4 respectively corresponding to the four simulation sets. **Figure S54-S55.** Time evolution of the angle between nominal planes formed by the G(/G_m_)-triads for the monomeric quadruplexes Quad-G4 and Quad-G_m_4 respectively corresponding to the four simulation sets. **Figure S56.** Water occupancy maps for m^6^A-tetrad respectively around the average structures for the three simulation sets. **Figure S57.** Different geometries observed for the three purine tetrads at position 2 of each monomeric quadruplex within the quadruplex dimer. **Figure S58.** Occupancy maps of K^+^ ions for the quadruplex dimers for the first three simulation sets showing the observed K^+^ ion binding pockets.

