## Supplemental data for "Structural and dynamic consequences of inosine, 2′-O-methylation, and N6-methyladenosine modifications in RNA G-quadruplex: A molecular dynamics study"

**Table S1. (A)** Interaction energies (kcal/mol) for hydrogen bonding, stacking and ion interactions for the different purine tetrads at position 2 of the monomeric quadruplexes for the first set of simulations. Average and standard deviation (within braces) values are reported.

| | | *ELEC* | *VDW* | TOTAL |
|---|---|---|---|---|
| *Hydrogen bonding energy of the tetrads* | | | | |
| *I-I-I-I* | *2-12*<br>*2-17*<br>*7-17*<br>*7-12* | *-6.91 (1.68)*<br>*-6.84 (1.68)*<br>*-6.93 (1.84)*<br>*-6.89 (1.71)* | *-1.11 (0.87)*<br>*-1.09 (0.85)*<br>*-1.11 (0.87)*<br>*-1.10 (0.87)* | *-8.02 (1.89)*<br>*-7.93 (1.88)*<br>*-8.04 (2.04)*<br>*-7.99 (1.92)* |
| *G-G-G-G* | *2-12*<br>*2-17*<br>*7-17*<br>*7-12* | *-14.27 (1.85)*<br>*-14.27 (1.84)*<br>*-14.27 (1.85)*<br>*-14.28 (1.84)* | *-0.41 (1.23)*<br>*-0.42 (1.22)*<br>*-0.41 (1.22)*<br>*-0.41 (1.22)* | *-14.68 (2.22)*<br>*-14.69 (2.21)*<br>*-14.68 (2.22)*<br>*-14.69 (2.21)* |
| *A-A-A-A* | *2-12*<br>*2-17*<br>*7-17*<br>*7-12* | *-3.02 (1.17)*<br>*-3.02 (1.17)*<br>*-3.02 (1.17)*<br>*-3.04 (1.16)* | *-0.87 (0.81)*<br>*-0.87 (0.82)*<br>*-0.87 (0.81)*<br>*-0.87 (0.81)* | *-3.89 (1.42)*<br>*-3.89 (1.43)*<br>*-3.89 (1.42)*<br>*-3.91 (1.41)* |
| *Im-Im-Im-Im* | *2-12*<br>*2-17*<br>*7-17*<br>*7-12* | *-4.74 (3.20)*<br>*-4.71 (3.18)*<br>*-4.76 (3.18)*<br>*-4.74 (3.17)* | *-1.32 (0.79)*<br>*-1.34 (0.78)*<br>*-1.32 (0.79)*<br>*-1.32 (0.78)* | *-6.06 (3.29)*<br>*-6.05 (3.27)*<br>*-6.08 (3.28)*<br>*-6.06 (3.26)* |
| *Gm-Gm-Gm-Gm* | *2-12*<br>*2-17*<br>*7-17*<br>*7-12* | *-11.07 (2.91)*<br>*-10.86 (2.95)*<br>*-11.16 (2.94)*<br>*-11.07 (2.92)* | *-0.84 (1.12)*<br>*-0.94 (1.09)*<br>*-0.80 (1.13)*<br>*-0.83 (1.12)* | *-11.91 (3.11)*<br>*-12.80 (3.14)*<br>*-11.96 (3.15)*<br>*-11.90 (3.12)* |
| *Am-Am-Am-Am* | *2-12*<br>*2-17*<br>*7-17*<br>*7-12* | *-3.19 (1.26)*<br>*-3.19 (1.26)*<br>*-3.22 (1.27)*<br>*-3.22 (1.26)* | *-1.16 (0.71)*<br>*-1.17 (0.71)*<br>*-1.15 (0.72)*<br>*-1.16 (0.72)* | *-4.35 (1.45)*<br>*-4.36 (1.45)*<br>*-4.37 (1.46)*<br>*-4.38 (1.45)* |
| *Stacking energy between tetrads* | | | | |
| *G4-I4*<br>*I4-G4* | | *18.37 (4.44)*<br>*22.44 (4.47)* | *-39.91 (1.40)*<br>*-39.43 (1.15)* | *-21.54 (4.66)*<br>*-16.99 (4.62)* |
| *G4-G4*<br>*G4-G4* | | *16.16 (3.39)*<br>*31.46 (2.80)* | *-41.15 (1.43)*<br>*-43.09(1.22)* | *-24.99 (3.68)*<br>*-11.63 (3.05)* |
| *G4-A4*<br>*A4-G4* | | *5.88 (3.45)*<br>*9.39 (3.77)* | *-39.19 (1.39)*<br>*-38.66 (1.26)* | *-33.31 (3.72)*<br>*-29.27 (3.97)* |
| *G4-Im4* | | *25.11 (5.10)* | *-39.79 (1.42)* | *-14.68 (5.29)* |

| | | | |
|---|---|---|---|
| *Im4-G4* | *24.61 (6.00)* | *-39.28 (1.17)* | *-14.67 (6.11)* |
| *G4-Gm4*<br>*Gm4-G4* | *18.29 (5.96)*<br>*32.38 (3.02)* | *-40.16 (1.76)*<br>*-42.51 (1.26)* | *-21.87 (6.21)*<br>*-10.13 (3.27)* |
| *G4-Am4*<br>*Am4-G4* | *3.26 (3.10)*<br>*3.99 (2.57)* | *-38.19 (1.49)*<br>*-38.42 (1.29)* | *-34.93 (3.44)*<br>*-34.43 (2.88)* |
| *Interaction energy between a tetrad and the proximal K+ cation* | | | |
| *I-I-I-I & K+*<br>*K24*<br>*K23* | <br>*-57.60 (21.94)*<br>*-45.33 (22.72)* | <br>*1.66 (2.05)*<br>*1.06 (2.93)* | <br>*-55.94 (22.04)*<br>*-44.27 (22.91)* |
| *G-G-G-G & K+*<br>*K24*<br>*K23* | <br>*-10.27 (11.38)*<br>*-88.58 (7.03)* | <br>*-0.02 (0.20)*<br>*4.56 (2.24)* | <br>*-10.29 (11.38)*<br>*-84.02 (7.38)* |
| *A-A-A-A and K+*<br>*K24*<br>*K23* | <br>*-11.12 (4.89)*<br>*-23.59 (5.79)* | <br>*-0.04 (0.12)*<br>*-0.31 (0.46)* | <br>*-11.16 (11.38)*<br>*-23.9 (5.81)* |
| *Im-Im-Im-Im & K+*<br>*K24*<br>*K23* | <br>*-79.06 (14.74)*<br>*-58.73 (33.49)* | <br>*2.02 (1.95)*<br>*0.97 (2.51)* | <br>*-77.04 (14.87)*<br>*-57.76 (33.58)* |
| *Gm-Gm-Gm-Gm & K+*<br>*K24*<br>*K23* | <br>*-19.01 (23.42)*<br>*-91.45 (9.74)* | <br>*0.05 (0.49)*<br>*4.83 (2.24)* | <br>*-18.96 (23.42)*<br>*-86.62 (9.99)* |
| *Am-Am-Am-Am and K+*<br>*K24*<br>*K23* | <br>*-9.83 (3.73)*<br>*-17.36 (2.87)* | <br>*-0.04 (0.11)*<br>*-0.38 (0.18)* | <br>*-9.87 (3.73)*<br>*-17.74 (2.88)* |

**(B)** Interaction energies (kcal/mol) for hydrogen bonding, stacking and ion interactions for the different purine tetrads at position 2 of the monomeric quadruplexes for the second set of simulations. Average and standard deviation (within braces) values are reported.

| | | *ELEC* | *VDW* | *TOTAL* |
|---|---|---|---|---|
| *Hydrogen bonding energy of the tetrads* | | | | |
| *I-I-I-I* | *2-12*<br>*2-17*<br>*7-17*<br>*7-12* | *-7.07 (1.50)*<br>*-7.04 (1.49)*<br>*-7.08 (1.49)*<br>*-7.02 (1.49)* | *-1.06 (0.89)*<br>*-1.08 (0.87)*<br>*-1.08 (0.88)*<br>*-1.08 (0.88)* | *-8.13 (1.74)*<br>*-8.12 (1.72)*<br>*-8.16 (1.73)*<br>*-8.10 (1.73)* |
| *G-G-G-G* | *2-12*<br>*2-17*<br>*7-17* | *-13.32 (1.99)*<br>*-13.33 (2.01)*<br>*-13.31 (2.02)* | *-0.32 (1.29)*<br>*-0.31 (1.29)*<br>*-0.31 (1.29)* | *-13.64 (2.37)*<br>*-13.64 (2.39)*<br>*-13.62 (2.39)* |

| | | | | |
|---|---|---|---|---|
| | *7-12* | *-13.33 (1.99)* | *–0.30 (1.30)* | *-13.63 (2.38)* |
| *A-A-A-A* | *2-12*<br>*2-17*<br>*7-17*<br>*7-12* | *-3.06 (1.20)*<br>*-2.92 (1.24)*<br>*-3.04 (1.19)*<br>*-3.05 (1.19)* | *-0.87 (0.81)*<br>*-0.90 (0.79)*<br>*-0.87 (0.82)*<br>*-0.87 (0.82)* | *-3.93 (1.45)*<br>*-3.82 (1.53)*<br>*-3.91 (1.44)*<br>*-3.92 (1.44)* |
| *Im-Im-Im-Im* | *2-12*<br>*2-17*<br>*7-17*<br>*7-12* | *-6.46 (1.47)*<br>*-6.43 (1.48)*<br>*-6.47 (1.47)*<br>*-6.45 (1.47)* | *-1.27 (0.82)*<br>*-1.28 (0.81)*<br>*-1.27 (0.82)*<br>*-1.27 (0.82)* | *-7.73 (1.68)*<br>*-7.71 (1.69)*<br>*-7.74 (1.68)*<br>*-7.72 (1.68)* |
| *Gm-Gm-Gm-Gm* | *2-12*<br>*2-17*<br>*7-17*<br>*7-12* | *-7.08 (3.89)*<br>*-7.06 (3.88)*<br>*-7.08 (3.89)*<br>*-7.04 (3.89)* | *-1.28 (1.09)*<br>*-1.29 (1.10)*<br>*-1.29 (1.11)*<br>*-1.29 (1.09)* | *-8.36 (4.04)*<br>*-8.35 (4.03)*<br>*-8.37 (4.04)*<br>*-8.33 (4.04)* |
| *Am-Am-Am-Am* | *2-12*<br>*2-17*<br>*7-17*<br>*7-12* | *-4.22 (1.38)*<br>*-4.21 (1.39)*<br>*-4.24 (1.38)*<br>*-4.22 (1.38)* | *-0.93 (0.93)*<br>*-0.94 (0.93)*<br>*-0.92 (0.94)*<br>*-0.94 (0.93)* | *-5.15 (1.66)*<br>*-5.15 (1.67)*<br>*-5.16 (1.67)*<br>*-5.16 (1.66)* |
| *Stacking energy between tetrads* | | | | |
| *G4-I4*<br>*I4-G4* | | *19.83 (2.39)*<br>*19.60 (2.26)* | *-40.23 (1.26)*<br>*-39.44 (1.12)* | *-20.40 (2.70)*<br>*-19.84 (2.52)* |
| *G4-G4*<br>*G4-G4* | | *30.02 (3.57)*<br>*24.67 (3.92)* | *-43.41 (1.23)*<br>*-42.67(1.24)* | *-13.39 (3.77)*<br>*-18.00 (4.11)* |
| *G4-A4*<br>*A4-G4* | | *5.87 (4.14)*<br>*9.25 (3.33)* | *-39.01 (1.46)*<br>*-38.58 (1.32)* | *-33.14 (4.39)*<br>*-29.33 (3.58)* |
| *G4-Im4*<br>*Im4-G4* | | *22.92 (2.38)*<br>*21.01 (2.36)* | *-40.21 (1.24)*<br>*-39.34 (1.12)* | *-17.29 (2.68)*<br>*-18.33 (2.61)* |
| *G4-Gm4*<br>*Gm4-G4* | | *32.16 (6.34)*<br>*32.43 (5.01)* | *-42.78 (1.43)*<br>*-42.55 (1.27)* | *-10.62 (6.50)*<br>*-10.12 (5.17)* |
| *G4-Am4*<br>*Am4-G4* | | *4.37 (3.43)*<br>*1.58 (2.89)* | *-38.99 (1.40)*<br>*-38.58 (1.34)* | *-34.62 (3.70)*<br>*-37.00 (3.18)* |
| *Interaction energy between a tetrad and the proximal K+ cation* | | | | |
| *I-I-I-I & K+*<br>*K24*<br>*K23* | | <br>*-5.79 (2.31)*<br>*-67.84 (5.82)* | <br>*-0.01 (0.03)*<br>*2.14 (1.89)* | <br>*-5.80 (2.31)*<br>*-65.70 (6.12)* |
| *G-G-G-G & K+*<br>*K24*<br>*K23* | | <br>*-11.30 (10.71)*<br>*-87.56 (7.47)* | <br>*-0.05 (0.20)*<br>*2.59 (2.22)* | <br>*-11.35 (10.71)*<br>*-84.97 (7.79)* |

| | | | |
|---|---|---|---|
| *A-A-A-A and K+*<br>*K24*<br>*K23* | <br>*-10.62 (4.74)*<br>*-11.63 (8.80)* | <br>*-0.04 (0.13)*<br>*0.04 (0.38)* | <br>*-10.66 (4.74)*<br>*-11.67 (8.81)* |
| *Im-Im-Im-Im & K+*<br>*K24*<br>*K23* | <br>*-74.76 (5.64)*<br>*-36.61 (2.65)* | <br>*2.48 (1.95)*<br>*-0.43 (0.07)* | <br>*-72.78 (5.96)*<br>*-37.04 (2.65)* |
| *Gm-Gm-Gm-Gm & K+*<br>*K24*<br>*K23* | <br>*-82.12 (28.00)*<br>*-90.87 (27.32)* | <br>*1.35 (1.89)*<br>*2.53 (2.91)* | <br>*-80.77 (28.06)*<br>*-88.34 (27.47)* |
| *Am-Am-Am-Am and K+*<br>*K24*<br>*K23* | <br>*-18.46 (2.36)*<br>*-9.28 (3.20)* | <br>*-0.44 (0.08)*<br>*-0.005 (0.086)* | <br>*-18.90 (2.36)*<br>*-9.28 (3.20)* |

**(C)** Interaction energies (kcal/mol) for hydrogen bonding, stacking and ion interactions for the different purine tetrads at position 2 of the monomeric quadruplexes for the third set of simulations. Average and standard deviation (within braces) values are reported.

| | | *ELEC* | *VDW* | *TOTAL* |
|---|---|---|---|---|
| *Hydrogen bonding energy of the tetrads* | | | | |
| *I-I-I-I* | *2-12*<br>*2-17*<br>*7-17*<br>*7-12* | *-7.03 (1.51)*<br>*-7.01 (1.51)*<br>*-7.04 (1.51)*<br>*-7.02 (1.51)* | *-1.08 (0.88)*<br>*-1.10 (0.87)*<br>*-1.07 (0.87)*<br>*-1.09 (0.84)* | *-8.11 (1.75)*<br>*-8.11 (1.74)*<br>*-8.11 (1.74)*<br>*-8.11 (1.73)* |
| *G-G-G-G* | *2-12*<br>*2-17*<br>*7-17*<br>*7-12* | *-13.36 (1.95)*<br>*-13.35 (1.95)*<br>*-13.37 (1.95)*<br>*-13.35 (1.95)* | *-0.29 (1.29)*<br>*-0.29 (1.29)*<br>*-0.29 (1.29)*<br>*-0.30 (1.29)* | *-13.65 (2.34)*<br>*-13.64 (2.34)*<br>*-13.66 (2.34)*<br>*-13.65 (2.34)* |
| *A-A-A-A* | *2-12*<br>*2-17*<br>*7-17*<br>*7-12* | *-3.33 (1.32)*<br>*-3.32 (1.32)*<br>*-3.33 (1.32)*<br>*-3.33 (1.32)* | *-0.81 (0.88)*<br>*-0.81 (0.89)*<br>*-0.82 (0.88)*<br>*-0.82 (0.89)* | *-4.14 (1.59)*<br>*-4.13 (1.59)*<br>*-4.15 (1.59)*<br>*-4.15 (1.59)* |
| *Im-Im-Im-Im* | *2-12*<br>*2-17*<br>*7-17*<br>*7-12* | *-5.74 (2.48)*<br>*-5.70 (2.48)*<br>*-5.75 (2.46)*<br>*-5.69 (2.49)* | *-1.29 (0.81)*<br>*-1.31 (0.80)*<br>*-1.29 (0.80)*<br>*-1.31 (0.79)* | *-7.03 (2.61)*<br>*-7.01 (2.60)*<br>*-7.04 (2.59)*<br>*-7.00 (2.61)* |
| *Gm-Gm-Gm-Gm* | *2-12*<br>*2-17*<br>*7-17*<br>*7-12* | *-7.52 (4.03)*<br>*-7.50 (4.04)*<br>*-7.55 (4.04)*<br>*-7.50 (4.02)* | *-1.21 (1.13)*<br>*-1.21 (1.13)*<br>*-1.21 (1.13)*<br>*-1.22 (1.12)* | *-8.73 (4.18)*<br>*-8.71 (4.19)*<br>*-8.76 (4.19)*<br>*-8.72 (4.17)* |
| *Am-Am-Am-Am* | *2-12* | *-3.54 (1.40)* | *-1.07 (0.80)* | *-4.61 (1.61)* |

| | | | | |
|---|---|---|---|---|
| | 2-17<br>7-17<br>7-12 | -3.56 (1.39)<br>-3.57 (1.39)<br>-3.58 (1.39) | -1.08 (0.81)<br>-1.07 (0.81)<br>-1.07 (0.81) | -4.64 (1.61)<br>-4.64 (1.61)<br>-4.65 (1.61) |
| *Stacking energy between tetrads* | | | | |
| G4-I4<br>I4-G4 | | 19.83 (2.39)<br>19.61 (2.26) | -40.23 (1.26)<br>-39.44 (1.11) | -20.40 (2.70)<br>-19.83 (2.52) |
| G4-G4<br>G4-G4 | | 30.47 (3.01)<br>23.98 (3.00) | -43.48 (1.18)<br>-42.57 (1.22) | -13.01 (3.23)<br>-18.59 (3.24) |
| G4-A4<br>A4-G4 | | 5.89 (4.62)<br>8.74 (3.80) | -39.30 (1.41)<br>-38.70 (1.29) | -33.41 (4.83)<br>-29.96 (4.01) |
| G4-Im4<br>Im4-G4 | | 23.95 (3.76)<br>22.44 (4.50) | -40.07 (1.32)<br>-39.32 (1.14) | -16.12 (3.98)<br>-16.88 (4.64) |
| G4-Gm4<br>Gm4-G4 | | 33.42 (4.47)<br>29.66 (6.16) | -43.14 (1.30)<br>-43.21 (1.16) | -9.72 (4.65)<br>-13.55 (6.27) |
| G4-Am4<br>Am4-G4 | | 3.53 (3.99)<br>3.31 (3.26) | -38.48 (1.56)<br>-38.47 (1.32)) | -34.95 (4.28)<br>-35.16 (3.52) |
| *Interaction energy between a tetrad and the proximal K+ cation* | | | | |
| I-I-I-I & K+<br>K24<br>K23 | | <br>-57.35 (20.79)<br>-43.58 (21.48) | <br>1.56 (1.98)<br>0.97 (2.87) | <br>-55.79 (20.88)<br>-42.61 (21.67) |
| G-G-G-G & K+<br>K24<br>K23 | | <br>-9.13 (5.76)<br>-86.71 (6.65) | <br>-0.02 (0.10)<br>2.37 (2.01) | <br>-9.15 (5.76)<br>-84.34 (6.95) |
| A-A-A-A and K+<br>K24<br>K23 | | <br>-14.23 (6.62)<br>-23.15 (6.67) | <br>-0.17 (0.23)<br>-0.32 (0.48) | <br>-14.40 (6.62)<br>-23.47 (6.68) |
| Im-Im-Im-Im & K+<br>K24<br>K23 | | <br>-77.27 (9.06)<br>-45.46 (23.81) | <br>2.36 (1.96)<br>0.05 (1.54) | <br>-74.91 (9.27)<br>-45.51 (23.86) |
| Gm-Gm-Gm-Gm & K+<br>K24<br>K23 | | <br>-91.25 (14.74)<br>-73.17 (34.66) | <br>1.91 (1.92)<br>1.38 (2.67) | <br>-89.34 (14.86)<br>-71.79 (34.76) |
| Am-Am-Am-Am and K+<br>K24<br>K23 | | <br>-13.70 (5.41)<br>-16.93 (4.34) | <br>-0.22 (0.21)<br>-0.36 (0.34) | <br>-13.92 (5.41)<br>-17.29 (4.35) |

**Table S2. (A)** Frequencies (in %) of the Hoogsteen (HG) hydrogen bonds between the bases within different tetrads of the studied monomeric quadruplexes for the first set of simulation.

| H-bonds | | Quad-I4 | Quad-G4 | Quad-A4 | Quad-Im4 | Quad-Gm4 | Quad-Am4 |
|---|---|---|---|---|---|---|---|
| | *(11)N2-H21- - -N7(1)* | 64.80 | 46.57 | 55.18 | 65.85 | 57.45 | 59.40 |
| *1-11* | *(11)N1-H1- - -O6(1)* | 45.11 | 66.95 | 62.31 | 34.00 | 61.98 | 53.70 |
| *2-12* | *(12)N2-H21- - -N7(2)* | NA | 54.99 | NA | NA | 52.17 | NA |
| | *(12)N1-H1- - -O6(2)* | 75.72 | 76.16 | NA | 52.51 | 57.71 | NA |
| | *(2)N6-H62 - - - N1(12)* | NA | NA | 47.80 | NA | NA | 33.16 |
| | *(12)N1-H1- - -N7(2)* | NA | NA | NA | 7.56 | NA | NA |
| *3-13* | | | | | | | |
| | *(13)N2-H21- - -N7(3)* | 57.65 | 65.27 | 57.86 | 63.16 | 71.97 | 65.08 |
| | *(13)N1-H1- - -O6(3)* | 64.64 | 36.01 | 71.88 | 65.28 | 40.26 | 73.93 |
| *4-14* | | | | | | | |
| | *(14)N2-H21- - -N7(4)* | 63.02 | 66.54 | 69.95 | 66.08 | 63.72 | 69.17 |
| *5-15* | *(14)N1-H1- - -O6(4)* | 37.11 | 43.22 | 34.16 | 43.80 | 59.52 | 32.64 |
| | *(5)N3-H3- - -O4(15)* | 81.89 | 79.13 | 81.41 | 81.59 | 79.49 | 80.86 |
| | *(1)N2-H21- - -N7(16)* | 62.73 | 46.58 | 55.34 | 65.73 | 54.94 | 58.28 |
| *1-16* | *(1)N1-H1- - -O6(16)* | 45.77 | 67.28 | 62.16 | 34.42 | 52.47 | 54.14 |
| *2-17* | *(2)N2-H21- - -N7(17)* | NA | 54.85 | NA | NA | 46.15 | NA |
| | *(2)N1-H1- - -O6(17)* | 74.91 | 76.22 | NA | 51.76 | 57.85 | NA |
| | *(17)N6-H62 - - - N1(2)* | NA | NA | 47.75 | NA | NA | 32.72 |
| | *(2)N1-H1- - -N7(17)* | NA | NA | NA | 7.16 | NA | NA |
| *3-18* | | | | | | | |
| | *(3)N2-H21- - -N7(18)* | 58.68 | 65.27 | 58.20 | 64.16 | 70.94 | 65.48 |
| | *(3)N1-H1- - -O6(18)* | 64.28 | 35.87 | 71.90 | 64.87 | 37.41 | 73.86 |
| *4-19* | | | | | | | |
| | *(4)N2-H21- - -N7(19)* | 62.83 | 66.18 | 69.58 | 66.07 | 64.38 | 69.22 |
| *5-20* | *(4)N1-H1- - -O6(19)* | 24.53 | 43.39 | 33.98 | 43.34 | 59.42 | 32.48 |
| | *(20)N3-H3- - -O4(5)* | 18.70 | 79.09 | 81.25 | 81.80 | 80.59 | 80.76 |

| | | | | | | | |
|---|---|---|---|---|---|---|---|
| | (16)N2-H21- - -N7(6) | 65.93 | 46.65 | 55.27 | 65.44 | 62.26 | 59.00 |
| 6-16 | (16)N1-H1- - -O6(6) | 43.21 | 67.57 | 62.06 | 34.13 | 50.02 | 52.92 |
| 7-17 | (17)N2-H21- - -N7(7) | NA | 55.12 | NA | NA | 51.68 | NA |
| | (17)N1-H1- - -O6(7) | 75.97 | 75.88 | NA | 52.53 | 59.90 | NA |
| | (7)N6-H62 - - - N1(17) | NA | NA | 47.87 | NA | NA | 34.00 |
| | (17)N1-H1- - -N7(7) | NA | NA | NA | 7.54 | NA | NA |
| 8-18 | | | | | | | |
| | (18)N2-H21- - -N7(8) | 52.31 | 65.49 | 57.95 | 63.53 | 71.26 | 65.65 |
| | (18)N1-H1- - -O6(8) | 64.15 | 36.05 | 72.10 | 64.79 | 38.83 | 73.76 |
| 9-19 | | | | | | | |
| | (19)N2-H21- - -N7(9) | 66.99 | 66.41 | 69.77 | 65.80 | 62.49 | 68.87 |
| 10-20 | (19)N1-H1- - -O6(9) | 35.63 | 43.26 | 34.09 | 43.72 | 59.61 | 32.58 |
| | (10)N3-H3- - -O4(20) | 18.71 | 79.02 | 81.40 | 81.55 | 80.49 | 80.67 |
| 6-11 | (6)N2-H21- - -N7(11) | 63.65 | 46.61 | 55.22 | 65.49 | 55.33 | 57.97 |
| | (6)N1-H1- - -O6(11) | 42.86 | 67.49 | 62.23 | 34.20 | 60.35 | 54.44 |
| 7-12 | (7)N2-H21- - -N7(12) | NA | 55.24 | NA | NA | 52.97 | NA |
| | (7)N1-H1- - -O6(12) | 76.30 | 76.27 | NA | 52.33 | 57.95 | NA |
| | (12)N6-H62 - - - N1(7) | NA | NA | 48.12 | NA | NA | 33.55 |
| | (7)N1-H1- - -N7(12) | NA | NA | NA | 7.68 | NA | NA |
| 8-13 | | | | | | | |
| | (8)N2-H21- - -N7(13) | 57.65 | 65.75 | 58.31 | 63.55 | 71.39 | 65.76 |
| | (8)N1-H1- - -O6(13) | 62.62 | 36.07 | 71.63 | 65.09 | 37.55 | 73.65 |
| 9-14 | | | | | | | |
| | (9)N2-H21- - -N7(14) | 64.57 | 66.18 | 69.73 | 65.99 | 62.35 | 68.61 |
| | (9)N1-H1- - -O6(14) | 32.67 | 43.29 | 34.14 | 43.63 | 59.82 | 32.26 |
| 10-15 | | | | | | | |
| | (15)N3-H3- - -O4(10) | 76.34 | 79.22 | 81.21 | 81.53 | 80.41 | 80.68 |

**(B)** Frequencies (in %) of the Hoogsteen (HG) hydrogen bonds between the bases within different tetrads of the studied monomeric quadruplexes for the second set of simulation.

| H-bonds | | Quad-I4 | Quad-G4 | Quad-A4 | Quad-Im4 | Quad-Gm4 | Quad-Am4 |
|---|---|---|---|---|---|---|---|
| | (11)N2-H21- - -N7(1) | 66.90 | 72.70 | 53.43 | 66.78 | 70.77 | 67.84 |
| 1-11 | (11)N1-H1- - -O6(1) | 40.63 | 46.31 | 65.49 | 29.15 | 41.84 | 54.19 |
| 2-12 | (12)N2-H21- - -N7(2) | NA | 70.15 | NA | NA | 47.23 | NA |
| | (12)N1-H1- - -O6(2) | 78.34 | 72.12 | NA | 71.18 | 25.10 | NA |
| | (2)N6-H62 - - - N1(12) | NA | NA | 47.52 | NA | NA | 44.78 |
| 3-13 | | | | | | | |
| | (13)N2-H21- - -N7(3) | 56.85 | 55.08 | 58.06 | 61.41 | 65.74 | 67.84 |
| | (13)N1-H1- - -O6(3) | 73.76 | 71.09 | 73.02 | 75.17 | 64.37 | 67.59 |

| | | | | | | | |
|---|---|---|---|---|---|---|---|
| *4-14* | | | | | | | |
| | *(14)N2-H21- - -N7(4)* | 69.54 | 70.00 | 69.47 | 68.77 | 63.70 | 68.34 |
| *5-15* | *(14)N1-H1- - -O6(4)* | 31.74 | 30.87 | 33.43 | 31.56 | 62.11 | 32.10 |
| | *(5)N3-H3- - -O4(15)* | 81.54 | 81.67 | 81.14 | 81.05 | 83.29 | 80.23 |
| | *(1)N2-H21- - -N7(16)* | 67.19 | 72.56 | 53.03 | 66.70 | 71.05 | 63.28 |
| *1-16* | *(1)N1-H1- - -O6(16)* | 41.07 | 46.02 | 64.38 | 29.11 | 41.38 | 54.67 |
| *2-17* | *(2)N2-H21- - -N7(17)* | NA | 70.36 | NA | NA | 46.97 | NA |
| | *(2)N1-H1- - -O6(17)* | 78.04 | 71.87 | NA | 70.50 | 24.70 | NA |
| | *(17)N6-H62 - - - N1(2)* | NA | NA | 47.20 | NA | NA | 44.44 |
| *3-18* | | | | | | | |
| | *(3)N2-H21- - -N7(18)* | 57.00 | 55.08 | 57.78 | 61.47 | 66.00 | 68.04 |
| *4-19* | *(3)N1-H1- - -O6(18)* | 73.32 | 70.95 | 73.15 | 74.78 | 63.68 | 67.39 |
| | *(4)N2-H21- - -N7(19)* | 69.79 | 70.00 | 70.05 | 68.59 | 63.45 | 68.21 |
| *5-20* | *(4)N1-H1- - -O6(19)* | 31.47 | 31.27 | 32.69 | 31.32 | 61.84 | 32.03 |
| | *(20)N3-H3- - -O4(5)* | 81.81 | 81.39 | 81.32 | 80.86 | 83.36 | 80.36 |
| | *(16)N2-H21- - -N7(6)* | 67.10 | 72.60 | 54.65 | 66.51 | 70.96 | 63.12 |
| *6-16* | *(16)N1-H1- - -O6(6)* | 40.38 | 45.80 | 63.20 | 28.38 | 41.23 | 54.53 |
| *7-17* | *(17)N2-H21- - -N7(7)* | NA | 70.11 | NA | NA | 46.98 | NA |
| | *(17)N1-H1- - -O6(7)* | 78.62 | 72.05 | NA | 71.51 | 24.88 | NA |
| | *(7)N6-H62 - - - N1(17)* | NA | NA | 47.20 | NA | NA | 44.78 |
| *8-18* | | | | | | | |
| | *(18)N2-H21- - -N7(8)* | 56.96 | 55.04 | 57.80 | 61.76 | 66.12 | 67.90 |
| | *(18)N1-H1- - -O6(8)* | 73.04 | 71.42 | 72.08 | 74.49 | 64.08 | 67.65 |
| *9-19* | | | | | | | |
| | *(19)N2-H21- - -N7(9)* | 69.61 | 69.79 | 69.88 | 68.35 | 63.89 | 67.84 |
| *10-20* | *(19)N1-H1- - -O6(9)* | 31.60 | 31.40 | 33.14 | 31.59 | 62.12 | 32.03 |
| | *(10)N3-H3- - -O4(20)* | 81.65 | 81.82 | 80.95 | 81.04 | 83.44 | 80.11 |
| *6-11* | *(6)N2-H21- - -N7(11)* | 67.27 | 72.68 | 53.01 | 66.64 | 70.46 | 62.96 |
| | *(6)N1-H1- - -O6(11)* | 40.54 | 46.60 | 65.31 | 28.67 | 41.40 | 55.16 |
| *7-12* | *(7)N2-H21- - -N7(12)* | NA | 70.37 | NA | NA | 46.92 | NA |
| | *(7)N1-H1- - -O6(12)* | 77.70 | 72.36 | NA | 71.40 | 24.67 | NA |
| | *(12)N6-H62 - - - N1(7)* | NA | NA | 48.04 | NA | NA | 45.49 |
| *8-13* | | | | | | | |
| | *(8)N2-H21- - -N7(13)* | 57.18 | 55.05 | 58.01 | 61.46 | 65.64 | 68.43 |
| | *(8)N1-H1- - -O6(13)* | 73.21 | 71.26 | 72.37 | 74.69 | 64.46 | 67.78 |
| *9-14* | | | | | | | |
| | *(9)N2-H21- - -N7(14)* | 69.71 | 70.18 | 69.61 | 68.51 | 63.46 | 68.28 |

| | | | | | | | |
|---|---|---|---|---|---|---|---|
| *10-15* | *(9)N1-H1- - -O6(14)* | 31.69 | 31.11 | 33.10 | 31.64 | 61.86 | 32.42 |
| | *(15)N3-H3- - -O4(10)* | 81.61 | 81.65 | 81.44 | 80.76 | 83.32 | 80.12 |

**(C)** Frequencies (in %) of the Hoogsteen (HG) hydrogen bonds between the bases within different tetrads of the studied monomeric quadruplexes for the third set of simulation.

| H-bonds | | Quad-I4 | Quad-G4 | Quad-A4 | Quad-Im4 | Quad-Gm4 | Quad-Am4 |
|---|---|---|---|---|---|---|---|
| | *(11)N2-H21- - -N7(1)* | 65.66 | 73.47 | 55.65 | 66.05 | 72.84 | 60.42 |
| *1-11* | *(11)N1-H1- - -O6(1)* | 42.13 | 45.48 | 63.58 | 29.54 | 37.57 | 55.25 |
| *2-12* | *(12)N2-H21- - -N7(2)* | NA | 70.60 | NA | NA | 48.22 | NA |
| | *(12)N1-H1- - -O6(2)* | 77.91 | 72.82 | NA | 63.37 | 30.54 | NA |
| | *(2)N6-H62 - - - N1(12)* | NA | NA | 51.76 | NA | NA | 37.25 |
| *3-13* | | | | | | | |
| | *(13)N2-H21- - -N7(3)* | 58.42 | 54.49 | 58.21 | 62.06 | 63.73 | 66.10 |
| | *(13)N1-H1- - -O6(3)* | 67.31 | 73.80 | 70.37 | 71.79 | 68.68 | 70.92 |
| *4-14* | | | | | | | |
| | *(14)N2-H21- - -N7(4)* | 68.04 | 70.18 | 69.96 | 67.66 | 64.69 | 68.12 |
| *5-15* | *(14)N1-H1- - -O6(4)* | 37.41 | 29.89 | 32.96 | 36.66 | 48.84 | 32.99 |
| | *(5)N3-H3- - -O4(15)* | 81.53 | 81.96 | 81.11 | 80.97 | 82.52 | 80.43 |
| | *(1)N2-H21- - -N7(16)* | 65.88 | 73.53 | 55.84 | 65.43 | 73.05 | 60.75 |
| *1-16* | *(1)N1-H1- - -O6(16)* | 42.07 | 45.35 | 63.29 | 30.17 | 37.76 | 54.45 |
| *2-17* | *(2)N2-H21- - -N7(17)* | NA | 70.39 | NA | NA | 48.01 | NA |
| | *(2)N1-H1- - -O6(17)* | 77.09 | 72.64 | NA | 62.53 | 30.68 | NA |
| | *(17)N6-H62 - - - N1(2)* | NA | NA | 51.49 | NA | NA | 37.24 |
| *3-18* | | | | | | | |
| | *(3)N2-H21- - -N7(18)* | 58.66 | 54.43 | 57.97 | 62.23 | 63.98 | 66.55 |
| | *(3)N1-H1- - -O6(18)* | 67.02 | 73.91 | 70.24 | 71.53 | 67.91 | 70.96 |
| *4-19* | | | | | | | |
| | *(4)N2-H21- - -N7(19)* | 67.97 | 70.27 | 69.80 | 67.63 | 64.45 | 68.74 |
| *5-20* | *(4)N1-H1- - -O6(19)* | 37.14 | 30.00 | 33.34 | 36.40 | 48.71 | 32.55 |
| | *(20)N3-H3- - -O4(5)* | 81.78 | 81.96 | 80.98 | 81.08 | 82.31 | 80.83 |
| | *(16)N2-H21- - -N7(6)* | 65.60 | 73.31 | 55.81 | 65.80 | 73.18 | 60.92 |
| *6-16* | *(16)N1-H1- - -O6(6)* | 41.39 | 45.26 | 63.23 | 29.83 | 37.42 | 53.82 |

| | | | | | | | |
|---|---|---|---|---|---|---|---|
| *7-17* | *(17)N2-H21- - -N7(7)* | NA | 70.52 | NA | NA | 48.36 | NA |
| | *(17)N1-H1- - -O6(7)* | 77.91 | 72.98 | NA | 63.52 | 30.88 | NA |
| | *(7)N6-H62 - - - N1(17)* | NA | NA | 51.32 | NA | NA | 37.74 |
| *8-18* | | | | | | | |
| | *(18)N2-H21- - -N7(8)* | 58.44 | 54.10 | 58.42 | 62.31 | 63.89 | 66.54 |
| | *(18)N1-H1- - -O6(8)* | 66.95 | 74.19 | 70.49 | 71.55 | 67.84 | 71.23 |
| *9-19* | | | | | | | |
| | *(19)N2-H21- - -N7(9)* | 68.04 | 70.15 | 69.13 | 67.52 | 64.19 | 68.23 |
| *10-20* | *(19)N1-H1- - -O6(9)* | 37.24 | 30.04 | 32.80 | 36.40 | 49.01 | 32.79 |
| | *(10)N3-H3- - -O4(20)* | 81.46 | 81.98 | 77.79 | 81.15 | 82.50 | 80.65 |
| *6-11* | *(6)N2-H21- - -N7(11)* | 65.64 | 73.39 | 55.63 | 65.91 | 73.06 | 60.77 |
| | *(6)N1-H1- - -O6(11)* | 42.23 | 45.51 | 63.17 | 29.96 | 37.68 | 54.54 |
| *7-12* | *(7)N2-H21- - -N7(12)* | NA | 70.32 | NA | NA | 47.95 | NA |
| | *(7)N1-H1- - -O6(12)* | 77.37 | 72.68 | NA | 62.92 | 30.26 | NA |
| | *(12)N6-H62 - - - N1(7)* | NA | NA | 51.39 | NA | NA | 37.77 |
| *8-13* | | | | | | | |
| | *(8)N2-H21- - -N7(13)* | 58.40 | 54.70 | 58.39 | 61.86 | 63.69 | 66.57 |
| | *(8)N1-H1- - -O6(13)* | 66.92 | 73.85 | 70.65 | 71.47 | 68.20 | 70.62 |
| *9-14* | | | | | | | |
| | *(9)N2-H21- - -N7(14)* | 67.91 | 70.20 | 69.49 | 67.13 | 64.49 | 68.26 |
| | *(9)N1-H1- - -O6(14)* | 37.31 | 29.91 | 33.47 | 36.41 | 48.97 | 32.54 |
| *10-15* | | | | | | | |
| | *(15)N3-H3- - -O4(10)* | 81.75 | 82.10 | 77.67 | 81.24 | 82.34 | 80.71 |

**(D)** Frequencies (in %) of the Hoogsteen (HG) hydrogen bonds between the bases within different tetrads of the studied monomeric quadruplexes for the fourth set of simulation.

| H-bonds | | Quad-I4 | Quad-G4 | Quad-A4 | Quad-Im4 | Quad-Gm4 | Quad-Am4 |
|---|---|---|---|---|---|---|---|
| | *(11)N2-H21- - -N7(1)* | 66.04 | 71.60 | 55.61 | 57.73 | 71.18 | 60.38 |
| *1-11* | *(11)N1-H1- - -O6(1)* | 42.38 | 46.67 | 59.48 | 52.90 | 47.38 | 51.76 |
| *2-12* | *(12)N2-H21- - -N7(2)* | NA | 70.09 | NA | NA | 8.37 | NA |
| | *(12)N1-H1- - -O6(2)* | 77.73 | 71.75 | NA | NA | 1.14 | NA |
| | *(2)N6-H62 - - - N1(12)* | NA | NA | 46.99 | NA | NA | 32.63 |
| | *(12)N1-H1- - -N7(2)* | NA | NA | NA | 53.63 | 14.11 | NA |
| *3-13* | | | | | | | |
| | *(13)N2-H21- - -N7(3)* | 54.40 | 52.85 | 57.57 | 69.70 | 69.87 | 63.87 |
| | *(13)N1-H1- - -O6(3)* | 71.67 | 70.99 | 65.18 | 31.06 | 30.52 | 72.95 |
| *4-14* | | | | | | | |
| | *(14)N2-H21- - -N7(4)* | 71.44 | 72.06 | 70.47 | 58.74 | 59.05 | 70.53 |
| | *(14)N1-H1- - -O6(4)* | 31.22 | 29.96 | 37.30 | 71.99 | 70.49 | 32.43 |
| *5-15* | | | | | | | |

| | | | | | | | |
|---|---|---|---|---|---|---|---|
| | *(5)N3-H3- - -O4(15)* | 80.67 | 80.52 | 79.85 | 80.46 | 81.97 | 79.76 |
| *1-16* | *(1)N2-H21- - -N7(16)* | 65.62 | 71.54 | 55.30 | 65.32 | 71.01 | 60.26 |
| | *(1)N1-H1- - -O6(16)* | 41.88 | 46.91 | 59.32 | 50.50 | 47.60 | 51.24 |
| *2-17* | *(2)N2-H21- - -N7(17)* | NA | 69.90 | NA | NA | 8.07 | NA |
| | *(2)N1-H1- - -O6(17)* | 77.20 | 71.63 | NA | NA | 1.10 | NA |
| | *(17)N6-H62 - - - N1(2)* | NA | NA | 50.17 | NA | NA | 32.38 |
| | *(2)N1-H1- - -N7(17)* | | | | 23.54 | 14.15 | NA |
| *3-18* | | | | | | | |
| | *(3)N2-H21- - -N7(18)* | 54.88 | 53.47 | 57.39 | 68.08 | 70.01 | 64.19 |
| | *(3)N1-H1- - -O6(18)* | 71.63 | 70.56 | 64.96 | 31.06 | 30.49 | 72.65 |
| *4-19* | | | | | | | |
| | *(4)N2-H21- - -N7(19)* | 71.32 | 71.86 | 70.87 | 61.90 | 59.67 | 70.77 |
| | *(4)N1-H1- - -O6(19)* | 31.06 | 30.00 | 37.15 | 72.23 | 70.22 | 32.17 |
| *5-20* | | | | | | | |
| | *(20)N3-H3- - -O4(5)* | 80.57 | 80.38 | 79.89 | 81.03 | 81.62 | 79.53 |
| *6-16* | *(16)N2-H21- - -N7(6)* | 65.44 | 71.67 | 55.39 | 62.44 | 70.73 | 60.63 |
| | *(16)N1-H1- - -O6(6)* | 41.60 | 47.11 | 59.73 | 48.97 | 47.22 | 50.23 |
| *7-17* | *(17)N2-H21- - -N7(7)* | NA | 69.55 | NA | NA | 8.13 | NA |
| | *(17)N1-H1- - -O6(7)* | 78.26 | 71.98 | NA | NA | 1.14 | NA |
| | *(7)N6-H62 - - - N1(17)* | NA | NA | 49.85 | NA | NA | 33.24 |
| | *(17)N1-H1- - -N7(7)* | NA | NA | NA | 31.95 | 14.33 | NA |
| *8-18* | | | | | | | |
| | *(18)N2-H21- - -N7(8)* | 55.11 | 52.88 | 56.77 | 69.81 | 70.28 | 63.99 |
| | *(18)N1-H1- - -O6(8)* | 71.40 | 70.70 | 65.09 | 31.13 | 30.64 | 72.63 |
| *9-19* | | | | | | | |
| | *(19)N2-H21- - -N7(9)* | 71.67 | 71.70 | 70.95 | 61.98 | 59.52 | 70.33 |
| | *(19)N1-H1- - -O6(9)* | 31.04 | 29.75 | 37.50 | 71.52 | 70.26 | 32.34 |
| *10-20* | | | | | | | |
| | *(10)N3-H3- - -O4(20)* | 80.45 | 80.53 | 79.97 | 82.01 | 81.72 | 79.84 |
| *6-11* | *(6)N2-H21- - -N7(11)* | 65.40 | 71.51 | 55.41 | 61.38 | 71.11 | 60.15 |
| | *(6)N1-H1- - -O6(11)* | 42.02 | 46.87 | 59.62 | 53.29 | 47.13 | 51.36 |
| *7-12* | *(7)N2-H21- - -N7(12)* | NA | 69.59 | NA | NA | 8.07 | NA |
| | *(7)N1-H1- - -O6(12)* | 77.50 | 71.88 | NA | NA | 1.11 | NA |
| | *(12)N6-H62 - - - N1(7)* | NA | NA | 49.85 | NA | NA | 33.38 |
| | *(7)N1-H1- - -N7(12)* | NA | NA | NA | 33.45 | 14.40 | NA |
| *8-13* | | | | | | | |
| | *(8)N2-H21- - -N7(13)* | 55.33 | 52.89 | 56.42 | 69.50 | 69.98 | 64.17 |
| | *(8)N1-H1- - -O6(13)* | 71.23 | 70.60 | 64.87 | 30.48 | 30.57 | 72.86 |
| *9-14* | | | | | | | |
| | *(9)N2-H21- - -N7(14)* | 71.12 | 71.71 | 70.93 | 61.88 | 59.48 | 70.21 |
| | *(9)N1-H1- - -O6(14)* | 30.65 | 29.88 | 37.05 | 71.36 | 70.36 | 32.16 |
| *10-15* | | | | | | | |

| | | | | | | | |
|---|---|---|---|---|---|---|---|
| | *(15)N3-H3- - -O4(10)* | 80.66 | 80.20 | 79.92 | 81.34 | 81.63 | 79.74 |

**Table S3.** Frequencies (in %) of the signature hydrogen bond between G-OP2 and 3′U-O2′ observed for the quadruplexes with reversed-3′U-tetrads for the studied monomeric quadruplexes for the four sets of simulations.

| Simulation set-1 | H-bonds | Quad-I4 | Quad-G4 | Quad-A4 | Quad-Im4 | Quad-Gm4 | Quad-Am4 |
|---|---|---|---|---|---|---|---|
| *G4-U5*<br>*G9-U10*<br>*G14-U15*<br>*G19-U20* | (4)OP2- - - HO2′-O2′(5)<br>(9)OP2- - - HO2′-O2′(10)<br>(14)OP2- - - HO2′-O2′(15)<br>(19)OP2- - - HO2′-O2′(20) | 97.78<br>98.54<br>98.03<br>22.56 | 98.15<br>98.24<br>97.22<br>97.95 | 98.46<br>98.41<br>98.36<br>98.46 | 97.72<br>97.87<br>97.79<br>98.09 | 97.35<br>97.70<br>96.84<br>97.68 | 98.47<br>98.51<br>98.38<br>98.45 |

| Simulation set-2 | H-bonds | Quad-I4 | Quad-G4 | Quad-A4 | Quad-Im4 | Quad-Gm4 | Quad-Am4 |
|---|---|---|---|---|---|---|---|
| *G4-U5*<br>*G9-U10*<br>*G14-U15*<br>*G19-U20* | (4)OP2- - - HO2′-O2′(5)<br>(9)OP2- - - HO2′-O2′(10)<br>(14)OP2- - - HO2′-O2′(15)<br>(19)OP2- - - HO2′-O2′(20) | 98.43<br>98.44<br>98.28<br>98.26 | 98.25<br>98.45<br>97.95<br>98.07 | 97.94<br>98.43<br>98.36<br>98.70 | 98.22<br>98.28<br>98.24<br>98.23 | 97.27<br>97.41<br>97.26<br>97.20 | 98.52<br>98.43<br>98.37<br>98.30 |

| Simulation set-3 | H-bonds | Quad-I4 | Quad-G4 | Quad-A4 | Quad-Im4 | Quad-Gm4 | Quad-Am4 |
|---|---|---|---|---|---|---|---|
| *G4-U5*<br>*G9-U10*<br>*G14-U15*<br>*G19-U20* | (4)OP2- - - HO2′-O2′(5)<br>(9)OP2- - - HO2′-O2′(10)<br>(14)OP2- - -H O2′-O2′(15)<br>(19)OP2- - - HO2′-O2′(20) | 98.39<br>98.33<br>98.40<br>98.48 | 98.17<br>98.64<br>98.38<br>98.65 | 98.59<br>94.33<br>98.33<br>98.29 | 98.12<br>98.20<br>98.43<br>98.26 | 97.57<br>97.37<br>97.55<br>97.82 | 98.28<br>98.52<br>98.13<br>98.33 |

| Simulation set-4 | H-bonds | Quad-I4 | Quad-G4 | Quad-A4 | Quad-Im4 | Quad-Gm4 | Quad-Am4 |
|---|---|---|---|---|---|---|---|
| *G4-U5*<br>*G9-U10*<br>*G14-U15*<br>*G19-U20* | (4)OP2- - - HO2′-O2′(5)<br>(9)OP2- - - HO2′-O2′(10)<br>(14)OP2- - - HO2′-O2′(15)<br>(19)OP2- - - HO2′-O2′(20) | 98.84<br>98.70<br>98.77<br>98.84 | 98.86<br>98.72<br>98.76<br>98.71 | 98.81<br>98.91<br>98.76<br>98.65 | 96.46<br>97.56<br>97.26<br>97.58 | 97.59<br>97.76<br>97.48<br>97.60 | 98.77<br>98.61<br>98.71<br>98.73 |

**Table S4.** Frequencies (in %) of the water bridging interactions observed for the I4/G4/A4/Im4/Gm4/Am4 tetrads within the studied monomeric quadruplexes for the four sets of simulations.

| I4 | N3(2)-HO2′(2) | N3(7)-HO2′(7) | N3(12)-HO2′(12) | N3(17)-HO2′(17) |
|---|---|---|---|---|

| | | | | |
|---|---|---|---|---|
| Simulation set-1<br>Simulation set-2<br>Simulation set-3<br>Simulation set-4 | 7.7<br>8.2<br>8.6<br>8.5 | 8.9<br>8.4<br>8.5<br>8.3 | 7.9<br>8.5<br>8.5<br>8.5 | 7.3<br>8.4<br>8.4<br>8.2 |
| | N7(2)-OP2(2) | N7(7)-OP2(7) | N7(12)-OP2(12) | N7(17)-OP2(17) |
| Simulation set-1<br>Simulation set-2<br>Simulation set-3<br>Simulation set-4 | 2.6<br>2.7<br>3.0<br>2.6 | 2.3<br>2.4<br>2.7<br>2.4 | 2.7<br>2.3<br>2.7<br>2.4 | 2.7<br>2.5<br>2.8<br>2.5 |
| G4 | N3(2)-HO2′(2) | N3(7)-HO2′(7) | N3(12)-HO2′(12) | N3(17)-HO2′(17) |
| Simulation set-1<br>Simulation set-2<br>Simulation set-3<br>Simulation set-4 | 13.7<br>12.6<br>12.4<br>12.3 | 13.6<br>12.7<br>12.6<br>12.04 | 13.4<br>12.6<br>12.3<br>11.9 | 13.5<br>12.6<br>12.8<br>11.9 |
| | (N2-)H22(12)-OP2(2) | (N2-)H22(17)-OP2(7) | (N2-)H22(7)-OP2(12) | (N2-)H22(2)-OP2(17) |
| Simulation set-1<br>Simulation set-2<br>Simulation set-3<br>Simulation set-4 | 28.73<br>37.36<br>37.18<br>39.34 | 28.41<br>37.16<br>37.26<br>39.53 | 28.53<br>37.13<br>37.21<br>39.11 | 28.62<br>37.38<br>37.17<br>39.19 |
| | N7(2)-OP2(2) | N7(7)-OP2(7) | N7(12)-OP2(12) | N7(17)-OP2(17) |
| Simulation set-1<br>Simulation set-2<br>Simulation set-3<br>Simulation set-4 | 1.1<br><0.2<br><0.2<br><0.2 | 1.1<br><0.2<br><0.2<br><0.2 | 1.1<br><0.2<br><0.2<br><0.2 | 1.2<br><0.2<br><0.2<br><0.2 |
| A4 | N3(2)-HO2′(2) | N3(7)-HO2′(7) | N3(12)-HO2′(12) | N3(17)-HO2′(17) |
| Simulation set-1<br>Simulation set-2<br>Simulation set-3<br>Simulation set-4 | 10.1<br>8.91<br>10.8<br>9.6 | 10.1<br>10.3<br>10.7<br>9.7 | 10.3<br>10.5<br>11.0<br>9.5 | 10.2<br>10.3<br>10.8<br>9.9 |
| | N7(2)-OP2(2) | N7(7)-OP2(7) | N7(12)-OP2(12) | N7(17)-OP2(17) |
| Simulation set-1<br>Simulation set-2<br>Simulation set-3<br>Simulation set-4 | 15.1<br>15.0<br>13.0<br>15.2 | 15.3<br>15.4<br>13.2<br>15.3 | 15.3<br>14.9<br>13.2<br>15.2 | 15.4<br>14.7<br>12.9<br>15.2 |
| Im4 | N7(2)-OP2(2) | N7(7)-OP2(7) | N7(12)-OP2(12) | N7(17)-OP2(17) |
| Simulation set-1<br>Simulation set-2<br>Simulation set-3<br>Simulation set-4 | 1.8<br>1.6<br>1.4<br><0.1 | 1.7<br>1.5<br>1.3<br><0.1 | 1.6<br>1.4<br>1.3<br><0.1 | 1.8<br>1.5<br>1.4<br><0.1 |
| Gm4 | H22(12)-OP2(2) | H22(17)-OP2(7) | H22(7)-OP2(12) | H22(2)-OP2(17) |

| | | | | |
|---|---|---|---|---|
| Simulation set-1<br>Simulation set-2<br>Simulation set-3<br>Simulation set-4 | 7.52<br>22.52<br>22.61<br>17.33 | 11.85<br>22.95<br>22.64<br>17.81 | 9.55<br>22.35<br>22.91<br>17.86 | 15.41<br>22.92<br>22.93<br>17.78 |
| Am4 | N7(2)-OP2(2) | N7(7)-OP2(7) | N7(12)-OP2(12) | N7(17)-OP2(17) |
| Simulation set-1<br>Simulation set-2<br>Simulation set-3<br>Simulation set-4 | 9.6<br>4.2<br>8.1<br>9.1 | 8.9<br>4.2<br>7.3<br>8.5 | 9.0<br>3.9<br>7.0<br>8.3 | 9.3<br>4.3<br>8.5<br>8.8 |

**Table S5.** Comparison of Gibbs free energies (obtained from MM-GBSA analysis) of the quadruplexes for the first three sets of simulations. Standard eviation values are reported within braces.

| *Free energy (G) (Kcal/mol)* | *I4* | *G4* | *A4* | *Im4* | *Gm4* | *Am4* |
|---|---|---|---|---|---|---|
| *Simulation set-1* | *-3204.72 (19.49)* | *-3757.90 (19.77)* | *-3540.58 (19.68)* | *-3202.76 (19.79)* | *-3715.93 (20.02)* | *-3263.26 (19.43)* |
| *Simulation set-2* | *-3210.16 (19.16)* | *-3759.09 (19.40)* | *-3540.07 (19.53)* | *-3206.27 (19.21)* | *-3713.56 (19.72)* | *-3269.77 (19.45)* |
| *Simulation set-3* | *-3209.69 (19.13)* | *-3759.76 (19.49)* | *-3541.49 (19.75)* | *-3204.50 (19.56)* | *-3712.26 (19.56)* | *-3265.44 (19.93)* |

**Table S6.** Comparison of N9-pyramidalization (based on the average and standard deviation values for ***κ***' dihedral angles (in degrees)) for the tetrads at positions 2, 3 and 4, respectively, (from the 5′-direction) of the studied monomeric quadruplexes. Values observed in the crystal structure (PDB ID:2GRB) are indicated in blue.

| TETRAD- 2 | I4 | G4 | A4 | Im4 | Gm4 | Am4 |
|---|---|---|---|---|---|---|
| 2 | 3.23 (10.65)<br>3.82 (10.67)<br>3.57 (10.65)<br>4.02 (10.58)<br>**-1.46** | 0.66 (10.22)<br>3.29 (10.17)<br>3.35 (10.19)<br>3.59 (10.19) | 3.54 (10.44)<br>2.38 (10.85)<br>4.00 (10.57)<br>4.40 (10.47) | 10.76 (9.56)<br>10.42 (9.59)<br>10.55 (9.56)<br>12.67 (9.28) | 4.97 (9.85)<br>7.72 (9.75)<br>7.81 (9.74)<br>7.78 (9.69) | 7.69 (10.01)<br>9.04 (10.32)<br>8.21 (10.12)<br>7.82 (9.92) |
| 7 | 3.72 (10.69)<br>3.76 (10.58)<br>3.46 (10.64)<br>4.04 (10.62)<br>**-2.27** | 0.64 (10.20)<br>3.28 (10.18)<br>3.42 (10.16)<br>3.59 (10.16) | 3.67 (10.47)<br>3.81 (10.50)<br>3.96 (10.60)<br>4.42 (10.57) | 10.76 (9.56)<br>10.51 (9.59)<br>10.53 (9.59)<br>11.55 (9.41) | 5.29 (9.88)<br>7.67 (9.78)<br>7.85 (9.76)<br>7.75 (9.69) | 7.68 (10.01)<br>9.17 (10.34)<br>8.29 (10.17)<br>7.93 (10.01) |

| | | | | | | |
|---|---|---|---|---|---|---|
| 12 | 3.47 (10.67)<br>3.82 (10.59)<br>3.50 (10.65)<br>3.91 (10.58)<br>**-1.52** | 0.54 (10.17)<br>3.30 (10.18)<br>3.32 (10.19)<br>3.57 (10.16) | 3.65 (10.46)<br>3.65 (10.51)<br>3.97 (10.57)<br>4.44 (10.53) | 10.69 (9.55)<br>10.46 (9.57)<br>10.49 (9.57)<br>11.99 (9.36) | 4.61 (9.85)<br>7.62 (9.80)<br>7.78 (9.77)<br>7.73 (9.71) | 7.56 (10.03)<br>9.17 (10.33)<br>8.15 (10.20)<br>7.84 (10.02) |
| 17 | 2.84 (10.73)<br>3.78 (10.59)<br>3.51 (10.68)<br>3.93 (10.56)<br>**-1.36** | 0.65 (10.18)<br>3.32 (10.15)<br>3.39 (10.19)<br>3.48 (10.19) | 3.68 (10.50)<br>3.61 (10.52)<br>3.96 (10.57)<br>4.44 (10.51) | 10.77 (9.57)<br>10.45 (9.57)<br>10.62 (9.57)<br>11.36 (9.35) | 4.84 (9.90)<br>7.71 (9.78)<br>7.89 (9.75)<br>7.81 (9.72) | 7.59 (10.04)<br>9.03 (10.36)<br>8.13 (10.22)<br>7.90 (9.96) |
| TETRAD 3 | | | | | | |
| 3 | 0.69 (9.95)<br>0.34 (9.88)<br>0.68 (9.95)<br>0.79 (9.86)<br>**1.71** | 2.12 (10.04)<br>0.99 (9.92)<br>0.91 (9.98)<br>1.57 (9.93) | -0.14 (10.00)<br>-0.17 (10.01)<br>0.01 (9.95)<br>0.76 (10.00) | 1.19 (10.05)<br>0.07 (9.85)<br>0.52 (9.92)<br>4.64 (9.76) | 2.34 (10.06)<br>2.65 (10.03)<br>2.24 (9.99)<br>4.38 (9.90) | 0.36 (9.96)<br>0.81 (9.96)<br>0.54 (9.98)<br>0.85 (10.00) |
| 8 | 0.87 (9.97)<br>0.32 (9.88)<br>0.61 (9.96)<br>0.74 (9.87)<br>**-0.09** | 2.13 (10.10)<br>1.08 (9.96)<br>0.96 (9.91)<br>1.56 (9.94) | -0.17 (9.97)<br>-0.26 (9.97)<br>-0.12 (9.98)<br>0.76 (10.02) | 1.14 (10.04)<br>0.07 (9.86)<br>0.48 (9.99)<br>4.76 (9.80) | 2.27 (9.99)<br>2.56 (9.99)<br>2.11 (10.01)<br>4.36 (9.88) | 0.21 (9.99)<br>0.86 (9.92)<br>0.47 (10.01)<br>0.75 (9.96) |
| 13 | 0.41 (9.94)<br>0.32 (9.89)<br>0.69 (9.93)<br>0.75 (9.87)<br>**0.72** | 2.08 (10.08)<br>1.07 (9.97)<br>0.86 (9.93)<br>1.62 (9.93) | -0.06 (9.96)<br>-0.16 (9.98)<br>0.04 (9.97)<br>0.78 (9.98) | 1.10 (10.03)<br>0.04 (9.88)<br>0.43 (9.91)<br>4.95 (9.84) | 2.52 (9.06)<br>2.55 (9.96)<br>2.14 (9.99)<br>4.33 (9.88) | 0.23 (9.96)<br>0.77 (9.97)<br>0.52 (9.99)<br>0.76 (9.94) |
| 18 | -1.46 (10.13)<br>0.26 (9.90)<br>0.66 (9.95)<br>0.72 (9.91)<br>**0.63** | 2.13 (10.10)<br>1.02 (9.97)<br>0.96 (9.95)<br>1.55 (9.94) | -0.12 (10.00)<br>-0.12 (9.98)<br>-0.02 (9.97)<br>0.78 (10.04) | 1.06 (10.01)<br>0.09 (9.87)<br>0.54 (9.97)<br>4.79 (9.79) | 2.47 (10.06)<br>2.58 (9.99)<br>2.12 (10.01)<br>4.46 (9.88) | 0.28 (10.01)<br>0.88 (9.94)<br>0.51 (9.99)<br>0.77 (9.95) |
| TETRAD 4 | | | | | | |
| 4 | -1.85 (9.98)<br>-1.79 (9.95)<br>-2.09 (9.92)<br>-2.67 (10.02)<br>**-2.41** | -2.59 (9.97)<br>-2.31 (10.01)<br>-2.32 (9.96)<br>-3.16 (10.04) | -2.47 (9.95)<br>-2.58 (9.94)<br>-2.66 (9.95)<br>-3.21 (9.99) | -2.81 (9.88)<br>-2.76 (9.92)<br>-2.78 (9.88)<br>-3.19 (9.89) | -2.97 (9.88)<br>-3.05 (9.89)<br>-2.69 (9.91)<br>-3.27 (9.84) | -2.61 (9.93)<br>-2.66 (9.93)<br>-2.51 (9.91)<br>-3.09 (9.98) |
| 9 | -2.69 (9.92)<br>-1.88 (9.95)<br>-2.12 (9.99)<br>-2.67 (9.96)<br>**-0.89** | -2.53 (9.98)<br>-2.36 (9.95)<br>-2.42 (9.98)<br>-3.07 (10.03) | -2.47 (9.95)<br>-2.57 (9.96)<br>-2.66 (9.96)<br>-3.25 (10.01) | -2.85 (9.90)<br>-2.71 (9.87)<br>-2.78 (9.91)<br>-3.26 (9.90) | -2.91 (9.90)<br>-2.97 (9.86)<br>-2.78 (9.88)<br>-3.24 (9.92) | -2.59 (9.94)<br>-2.84 (9.97)<br>-2.54 (9.92)<br>-3.15 (9.98) |
| 14 | -2.26 (9.93)<br>-1.81 (9.96)<br>-2.02 (9.95) | -2.56 (9.95)<br>-2.36 (9.97)<br>-2.34 (9.97) | -2.44 (9.95)<br>-2.36 (9.98)<br>-2.54 (9.97) | -2.91 (9.91)<br>-2.75 (9.90)<br>-2.76 (9.91) | -2.46 (9.87)<br>-3.02 (9.85)<br>-2.73 (9.86) | -2.58 (9.92)<br>-2.66 (9.88)<br>-2.51 (9.92) |

| | | | | | | |
|---|---|---|---|---|---|---|
| | -2.73 (9.97)<br>**-1.03** | -3.17 (9.98) | -3.25 (10.03) | -2.87 (9.91) | -3.21 (9.89) | -3.19 (9.95) |
| 19 | -6.06 (9.94)<br>-1.87 (9.96)<br>-1.96 (9.97)<br>-2.74 (10.02)<br>**0.04** | -2.64 (9.98)<br>-2.38 (9.98)<br>-2.35 (9.94)<br>-3.09 (10.01) | -2.53 (9.96)<br>-2.51 (9.98)<br>-2.49 (9.95)<br>-3.28 (10.04) | -2.91 (1.87)<br>-2.75 (9.92)<br>-2.67 (9.90)<br>-3.22 (9.90) | -2.60 (9.89)<br>-2.98 (9.86)<br>-2.68 (9.90)<br>-3.13 (9.90) | -2.47 (9.92)<br>-2.60 (9.89)<br>-2.51 (9.89)<br>-3.06 (9.98) |

**Table S7. (A)** Structural base pair and base pair step parameters for the Hoogsteen (HG) pairs in the different tetrads within the monomeric quadruplex containing I(2)-tetrad (Quad-I4) for the first simulation set. A, B, C and D represent the chains. Average and standard deviation (within braces) values are reported.

| HG_A_C | shear | stretch | stagger | buckle | propeller | opening |
|---|---|---|---|---|---|---|
| G-G | -3.31 (0.26) | 1.76 (0.30) | 0.04 (0.38) | -11.5 (8.2) | -24.5 (8.0) | -90.6 (3.9) |
| I-I | -3.20 (0.27) | 1.86 (0.33) | -0.09 (0.34) | -14.7 (7.1) | -17.4 (6.5) | -90.8 (4.8) |
| G-G | -3.18 (0.26) | 1.94 (0.31) | 0.01 (0.35) | -6.8 (6.6) | -10.4 (6.5) | -89.9 (2.9) |
| G-G | 3.46 (0.25) | 1.53 (0.30) | 0.13 (0.40) | -5.2 (8.1) | 4.9 (8.1) | -89.6 (2.7) |
| U-U | 2.74 (0.32) | 0.55 (0.34) | 0.12 (0.63) | 19.2 (11.5) | -9.4 (10.9) | 89.4 (6.0) |
| HG_A_D | | | | | | |
| G-G | -3.31 (0.27) | 1.78 (0.30) | 0.01 (0.38) | -10.1 (8.1) | -23.9 (7.7) | 90.7 (3.9) |
| I-I | -3.20 (0.30) | 1.86 (0.34) | 0.13 (0.35) | -12.4 (7.0) | -16.8 (6.7) | 89.8 (5.1) |
| G-G | -3.19 (0.26) | 1.95 (0.32) | 0.18 (0.35) | -2.4 (7.4) | -10.6 (6.5) | 90.2 (2.9) |
| G-G | 3.57 (0.28) | 1.48 (0.33) | 0.15 (0.44) | -10.6 (8.7) | 2.1 (8.2) | 90.8 (2.8) |
| U-U | -1.84 (2.81) | 1.81 (1.72) | 2.69 (2.06) | 33.2 (15.2) | 21.5 (20.3) | -102.2 (14.8) |
| HG_B_C | | | | | | |
| G-G | -3.33 (0.27) | 1.75 (0.30) | -0.0.3 (0.38) | -10.0 (8.1) | -23.8 (7.8) | 90.8 (3.8) |
| I-I | -3.20 (0.28) | 1.86 (0.33) | 0.0.7 (0.34) | -14.0 (7.1) | -17.2 (6.4) | 90.7 (4.8) |
| G-G | -3.19 (0.26) | 1.96 (0.32) | -0.02 (0.34) | -6.5 (6.5) | -9.8 (6.6) | 90.2 (3.0) |
| G-G | 3.50 (0.25) | 1.47 (0.32) | -0.15 (0.41) | -7.5 (8.1) | 6.1 (8.4) | 89.7 (2.7) |
| U-U | 2.87 (0.29) | 0.58 (0.33) | -0.09 (0.60) | 23.3 (11.1) | -6.1 (10.4) | -91.7 (5.8) |
| HG_B_D | | | | | | |

| | | | | | | |
|---|---|---|---|---|---|---|
| G-G | -3.32 (0.27) | 1.77 (0.30) | 0.06 (0.38) | -11.4 (8.1) | -24.4 (7.8) | -91.0 (3.8) |
| I-I | -3.19 (0.30) | 1.87 (0.37) | 0.03 (0.35) | -15.2 (7.2) | -15.7 (6.8) | -90.8 (5.0) |
| G-G | -3.18 (0.26) | 1.93 (0.31) | 0.16 (0.36) | -6.0 (6.2) | -6.3 (6.9) | -89.5 (3.0) |
| G-G | 3.50 (0.25) | 1.43 (0.35) | 0.26 (0.41) | -9.1 (8.0) | 8.0 (8.7) | -89.3 (2.7) |
| U-U | 0.55 (1.66) | -3.67 (2.51) | 2.82 (1.64) | -1.8 (17.7) | 1.6 (11.5) | 70.9 (15.6) |

| HG_A_C | shift | slide | rise | tilt | roll | twist |
|---|---|---|---|---|---|---|
| G-G/I-I | -1.09 (0.33) | 0.17 (0.30) | 3.32 (0.17) | 0.1 (3.6) | -2.2 (4.2) | 33.0 (3.4) |
| I-I/G-G | -0.84 (0.35) | 0.27 (0.34) | 3.11 (0.16) | 0.3 (3.4) | -4.1 (4.1) | 28.4 (3.2) |
| G-G/G-G | -0.88 (0.32) | 0.43 (0.33) | 3.05 (0.16) | 3.5 (3.3) | -4.3 (3.9) | 22.2 (3.2) |
| G-G/U-U | -0.61 (2.35) | -2.96 (0.94) | 0.15 (0.52) | -39.6 (48.1) | 170.0 (13.0) | 2.1 (6.9) |
| HG_A_D | | | | | | |
| G-G/I-I | 1.10 (0.33) | -0.17 (0.30) | 3.28 (0.17) | -0.0 (3.26) | 2.1 (4.2) | 33.1 (3.6) |
| I-I/G-G | 0.95 (0.34) | -0.29 (0.33) | 3.06 (0.17) | -1.0 (3.3) | 3.1 (4.2) | 28.1 (3.4) |
| G-G/G-G | 0.94 (0.36) | -0.56 (0.36) | 2.98 (0.16) | -3.9 (3.2) | 3.0 (3.9) | 22.1 (3.2) |
| G-G/U-U | -3.84 (2.69) | 1.40 (0.99) | -2.61 (1.68) | -28.3 (37.0) | 164.3 (15.5) | -13.0 (11.8) |
| HG_B_C | | | | | | |
| G-G/I-I | 1.13 (0.33) | -0.11 (0.30) | 3.32 (0.17) | -0.0 (3.6) | 2.4 (4.2) | 33.3 (3.4) |
| I-I/G-G | 0.90 (0.34) | -0.31 (0.33) | 3.09 (0.16) | -0.8 (3.3) | 4.4 (4.2) | 28.3 (3.1) |
| G-G/G-G | 0.82 (0.31) | -0.42 (0.33) | 3.03 (0.16) | -3.8 (3.3) | 4.0 (4.0) | 21.9 (3.3) |
| G-G/U-U | 0.74 (2.06) | 3.18 (0.86) | 0.18 (0.49) | 44.3 (44.1) | -168.1 (12.2) | 2.5 (7.0) |
| HG_B_D | | | | | | |
| G-G/I-I | -1.03 (0.34) | 0.12 (0.29) | 3.32 (0.17) | -0.1 (3.6) | -2.6 (4.1) | 33.4 (3.7) |
| I-I/G-G | -0.86 (0.34) | 0.26 (0.35) | 3.08 (0.16) | 0.7 (3.3) | -4.3 (4.2) | 27.9 (3.4) |
| G-G/G-G | -0.79 (0.30) | 0.47 (0.34) | 3.02 (0.16) | 4.5 (3.1) | -3.7 (3.9) | 21.5 (3.4) |
| G-G/U-U | -3.29 (2.94) | -1.13 (1.40) | -0.61 (0.95) | -35.0 (30.2) | 165.9 (11.0) | -19.0 (19.8) |

**(B)** Structural base pair and base pair step parameters for the Hoogsteen (HG) pairs in the different tetrads within the monomeric quadruplex containing G(2)-tetrad (Quad-G4) for the first simulation set. A, B, C and D represent the chains. Average and standard deviation (within braces) values are reported.

| HG_A_C | shear | stretch | stagger | buckle | propeller | opening |
|---|---|---|---|---|---|---|
| G-G | -2.91 (0.41) | 2.27 (0.45) | -0.01 (0.49) | -14.5 (9.9) | -17.9 (8.5) | -90.4 (4.7) |
| G-G | -2.87 (0.26) | 2.29 (0.30) | 0.01 (0.34) | -14.8 (7.0) | -18.9 (6.6) | -90.6 (4.0) |
| G-G | -3.47 (0.21) | 1.57 (0.23) | -0.0.6 (0.35) | -6.6 (6.6) | -6.7 (7.1) | -89.9 (2.8) |
| G-G | 3.43 (0.24) | 1.58 (0.30) | 0.09 (0.42) | -7.6 (8.3) | 3.9 (8.6) | -89.8 (2.7) |
| U-U | 2.74 (0.29) | 0.63 (0.32) | 0.26 (0.62) | 22.7 (11.2) | -10.8 (11.2) | 90.5 (5.6) |
| HG_A_D | | | | | | |
| G-G | -2.91 (0.40) | 2.26 (0.44) | 0.0.2 (0.49) | -14.2 (9.9) | -17.9 (8.5) | 90.3 (4.7) |
| G-G | -2.87 (0.26) | 2.29 (0.30) | -0.01 (0.33) | -14.8 (7.1) | -19.1 (6.5) | 90.7 (3.9) |
| G-G | -3.47 (0.21) | 1.56 (0.24) | 0.06 (0.35) | -6.7 (6.6) | -6.7 (7.0) | 89.9 (2.8) |
| G-G | 3.43 (0.24) | 1.57 (0.30) | -0.09 (0.42) | -7.6 (8.3) | 3.9 (8.7) | 89.8 (2.7) |
| U-U | 2.74 (0.30) | 0.63 (0.32) | -0.25 (0.62) | 22.5 (11.3) | -10.7 (11.3) | -90.4 (5.6) |
| HG_B_C | | | | | | |
| G-G | -2.91 (0.40) | 2.26 (0.44) | 0.01 (0.49) | -14.4 (9.9) | -17.6 (8.6) | 90.3 (4.7) |
| G-G | -2.88 (0.26) | 2.29 (0.30) | -0.00 (0.34) | -14.6 (7.3) | -18.9 (6.5) | 90.6 (3.9) |
| G-G | -3.47 (0.20) | 1.57 (0.23) | 0.06 (0.35) | -6.7 (6.6) | -6.8 (6.9) | 89.9 (2.8) |
| G-G | 3.43 (0.24) | 1.58 (0.30) | -0.09 (0.42) | -7.4 (8.2) | 4.0 (8.6) | 89.8 (2.7) |
| U-U | 2.74 (0.30) | 0.63 (0.32) | -0.25 (0.62) | 22.5 (11.2) | -10.6 (11.4) | -90.5 (5.6) |
| HG_B_D | | | | | | |
| G-G | -2.91 (0.39) | 2.27 (0.44) | -0.01 (0.49) | -14.2 (9.8) | -17.6 (8.5) | -90.4 (4.6) |
| G-G | -2.88 (0.26) | 2.29 (0.30) | -0.00 (0.34) | -14.6 (7.1) | -19.0 (6.5) | -90.7 (3.9) |
| G-G | -3.47 (0.20) | 1.57 (0.23) | -0.06 (0.35) | -6.6 (6.6) | -6.6 (7.0) | -89.9 (2.8) |
| G-G | 3.43 (0.24) | 1.58 (0.30) | 0.09 (0.42) | -7.7 (8.3) | 4.0 (8.7) | -89.8 (2.7) |
| U-U | 2.74 (0.30) | 0.63 (0.32) | 0.27 (0.61) | 22.8 (11.4) | -10.8 (11.3) | 90.5 (5.6) |

| HG_A_C | shift | slide | rise | tilt | roll | twist |
|---|---|---|---|---|---|---|

| | | | | | | |
|---|---|---|---|---|---|---|
| G-G/G-G | -1.00 (0.44) | 0.30 (0.38) | 3.42 (0.19) | -2.4 (3.7) | -2.6 (4.6) | 31.6 (3.5) |
| G-G/G-G | -1.17 (0.27) | 0.51 (0.29) | 3.09 (0.16) | 2.0 (3.2) | -3.5 (3.9) | 23.5 (2.6) |
| G-G/G-G | -0.73 (0.32) | 0.30 (0.34) | 3.10 (0.16) | 2.9 (3.3) | -3.9 (4.0) | 24.5 (2.3) |
| G-G/U-U | -0.55 (2.36) | -2.90 (0.95) | 0.09 (0.50) | -37.8 (48.2) | 170.6 (12.9) | 1.8 (6.8) |
| HG_A_D | | | | | | |
| G-G/G-G | 1.02 (0.43) | -0.30 (0.38) | 3.42 (0.19) | 2.5 (3.7) | 2.7 (4.6) | 31.6 (3.5) |
| G-G/G-G | 1.16 (0.27) | -0.51 (0.29) | 3.09 (0.16) | -2.1 (3.2) | 3.5 (3.9) | 23.4 (2.5) |
| G-G/G-G | 0.73 (0.32) | -0.30 (0.33) | 3.10 (0.16) | -2.8 (3.2) | 3.9 (4.0) | 24.5 (2.4) |
| G-G/U-U | 0.58 (2.34) | 2.91 (0.95) | 0.09 (0.49) | 38.5 (47.8) | -170.4 (12.9) | 1.9 (6.9) |
| HG_B_C | | | | | | |
| G-G/G-G | 1.01 (0.43) | -0.32 (0.38) | 3.42 (0.19) | 2.4 (3.7) | 2.6 (4.6) | 31.6 (3.5) |
| G-G/G-G | 1.16 (0.27) | -0.51 (0.29) | 3.09 (0.16) | -2.1 (3.3) | 3.6 (4.0) | 23.5 (2.6) |
| G-G/G-G | 0.72 (0.32) | -0.31 (0.34) | 3.10 (0.16) | -2.9 (3.2) | 3.9 (4.0) | 24.5 (2.3) |
| G-G/U-U | 0.56 (2.35) | 2.91 (0.95) | 0.08 (0.49) | 38.1 (48.0) | -170.5 (12.9) | 2.0 (6.7) |
| HG_B_D | | | | | | |
| G-G/G-G | -1.02 (0.43) | 0.31 (0.38) | 3.42 (0.19) | -2.5 (3.7) | -2.6 (4.6) | 31.6 (3.4) |
| G-G/G-G | -1.17 (0.27) | 0.51 (0.29) | 3.09 (0.16) | 2.0 (3.2) | -3.6 (3.9) | 23.5 (2.6) |
| G-G/G-G | -0.73 (0.32) | 0.31 (0.33) | 3.10 (0.16) | 2.8 (3.2) | -3.8 (4.0) | 24.5 (2.4) |
| G-G/U-U | -0.55 (2.36) | -2.90 (0.94) | 0.09 (0.50) | -37.9 (48.2) | 170.5 (13.0) | 1.9 (6.7) |

**(C)** Structural base pair and base pair step parameters for the Hoogsteen (HG) pairs in the different tetrads within the monomeric quadruplex containing A(2)-tetrad (Quad-A4) for the first simulation set. A, B, C and D represent the chains. Average and standard deviation (within braces) values are reported.

| HG_A_C | shear | stretch | stagger | buckle | propeller | opening |
|---|---|---|---|---|---|---|
| G-G | -3.13 (0.35) | 2.00 (0.40) | 0.01 (0.45) | -11.2 (9.4) | -18.2 (8.6) | -90.3 (4.0) |
| A-A | -3.13 (0.21) | 2.20 (0.28) | -0.00 (0.42) | -13.1 (7.9) | -18.0 (7.0) | -90.5 (4.7) |
| G-G | -3.09 (0.23) | 2.04 (0.27) | 0.00 (0.35) | -7.0 (6.6) | -11.2 (6.9) | -90.1 (3.1) |
| G-G | 3.50 (0.20) | 1.47 (0.24) | 0.07 (0.38) | -6.1 (7.8) | 3.3 (7.9) | -89.9 (2.6) |

| | | | | | | |
|---|---|---|---|---|---|---|
| U-U | 2.74 (0.27) | 0.63 (0.30) | 0.23 (0.60) | 21.4 (11.1) | -12.2 (10.7) | 90.6 (5.4) |
| HG_A_D | | | | | | |
| G-G | -3.13 (0.35) | 2.00 (0.40) | -0.00 (0.44) | -11.2 (9.3) | -18.4 (8.6) | -90.3 (3.9) |
| A-A | -3.13 (0.22) | 2.21 (0.28) | 0.01 (0.42) | -13.0 (7.8) | -18.0 (7.0) | 90.5 (4.6) |
| G-G | -3.09 (0.23) | 2.04 (0.27) | -0.00 (0.34) | -7.0 (6.6) | -11.1 (6.8) | -90.1 (3.1) |
| G-G | 3.50 (0.20) | 1.47 (0.24) | -0.07 (0.37) | -6.0 (7.7) | 3.3 (7.8) | 89.8 (2.6) |
| U-U | 2.74 (0.27) | 0.63 (0.30) | -0.24 (0.60) | 21.5 (11.1) | -12.4 (10.6) | -90.5 (5.4) |
| HG_B_C | | | | | | |
| G-G | -3.13 (0.35) | 2.00 (0.40) | -0.00 (0.44) | -11.0 (9.3) | -18.3 (8.6) | 90.3 (3.9) |
| A-A | -3.13 (0.22) | 2.21 (0.28) | 0.01 (0.41) | -13.0 (7.9) | -18.0 (7.1) | 90.6 (4.7) |
| G-G | -3.09 (0.23) | 2.04 (0.26) | -0.01 (0.35) | -7.1 (6.6) | -11.1 (6.9) | 90.1 (3.0) |
| G-G | 3.50 (0.20) | 1.47 (0.24) | -0.08 (0.37) | -6.0 (7.8) | 3.3 (7.9) | 89.8 (2.6) |
| U-U | 2.74 (0.27) | 0.62 (0.30) | -0.24 (0.59) | 21.6 (11.1) | -12.3 (10.7) | -90.4 (5.4) |
| HG_B_D | | | | | | |
| G-G | -3.13 (0.34) | 2.00 (0.40) | 0.01 (0.44) | -11.1 (9.4) | -18.4 (8.6) | -90.4 (3.9) |
| A-A | -3.13 (0.21) | 2.21 (0.28) | -0.01 (0.42) | -13.1 (7.9) | -18.0 (7.1) | -90.4 (4.7) |
| G-G | -3.09 (0.23) | 2.04 (0.26) | -0.00 (0.35) | -7.0 (6.6) | -11.1 (6.8) | -90.0 (3.0) |
| G-G | 3.50 (0.20) | 1.47 (0.24) | 0.07 (0.37) | -6.1 (7.7) | 3.3 (7.9) | -89.8 (2.6) |
| U-U | 2.74 (0.27) | 0.63 (0.29) | 0.24 (0.60) | 21.6 (11.2) | -12.3 (10.7) | 90.4 (5.4) |

| HG_A_C | shift | slide | rise | tilt | roll | twist |
|---|---|---|---|---|---|---|
| G-G/A-A | -0.96 (0.38) | 0.29 (0.34) | 3.38 (0.19) | -1.7 (3.6) | -2.2 (4.6) | 32.1 (3.2) |
| A-A/G-G | -0.98 (0.30) | -0.29 (0.30) | 3.16 (0.17) | 0.5 (3.6) | -3.4 (4.5) | 27.8 (2.5) |
| G-G/G-G | -0.92 (0.29) | 0.49 (0.30) | 3.03 (0.16) | 3.7 (3.2) | -3.8 (3.9) | 21.9 (2.5) |
| G-G/U-U | -0.64 (2.28) | -3.00 (0.91) | 0.10 (0.42) | -41.8 (48.1) | 169.4 (12.9) | 2.3 (6.7) |
| HG_A_D | | | | | | |
| G-G/A-A | 0.96 (0.38) | -0.29 (0.35) | 3.38 (0.19) | 1.7 (3.6) | 2.2 (4.6) | 32.0 (3.3) |
| A-A/G-G | 0.98 (0.30) | -0.29 (0.30) | 3.15 (0.18) | -0.5 (3.6) | 3.5 (4.5) | 27.7 (2.5) |

| | | | | | | |
|---|---|---|---|---|---|---|
| G-G/G-G | 0.93 (0.29) | -0.49 (0.30) | 3.03 (0.15) | -3.7 (3.2) | 3.9 (3.8) | 21.9 (2.5) |
| G-G/U-U | 0.63 (2.29) | 2.98 (0.91) | 0.10 (0.42) | 41.4 (48.2) | -169.6 (13.0) | 2.2 (6.7) |
| HG_B_C | | | | | | |
| G-G/A-A | 0.96 (0.38) | -0.29 (0.35) | 3.38 (0.19) | 1.7 (3.6) | 2.2 (4.5) | 32.1 (3.2) |
| A-A/G-G | 0.98 (0.30) | -0.29 (0.30) | 3.16 (0.18) | -0.5 (3.6) | 3.5 (4.5) | 27.7 (2.5) |
| G-G/G-G | 0.92 (0.29) | -0.49 (0.30) | 3.03 (0.16) | -3.7 (3.2) | 3.8 (3.9) | 21.9 (2.5) |
| G-G/U-U | 0.63 (2.29) | 2.99 (0.91) | 0.10 (0.42) | 41.7 (48.1) | -169.6 (13.0) | 2.2 (6.6) |
| HG_B_D | | | | | | |
| G-G/A-A | -0.96 (0.38) | 0.29 (0.35) | 3.38 (0.19) | -1.7 (3.6) | -2.2 (4.6) | 32.1 (3.3) |
| A-A/G-G | -0.99 (0.30) | 0.29 (0.30) | 3.16 (0.18) | 0.5 (3.6) | -3.4 (4.5) | 27.7 (2.5) |
| G-G/G-G | -0.92 (0.29) | 0.49 (0.30) | 3.03 (0.15) | 3.7 (3.1) | -3.8 (3.9) | 21.9 (2.5) |
| G-G/U-U | -0.65 (2.28) | -2.99 (0.91) | 0.10 (0.42) | -41.8 (47.9) | 169.5 (12.9) | 2.4 (6.7) |

**(D)** Structural base pair and base pair step parameters for the Hoogsteen (HG) pairs in the different tetrads within the monomeric quadruplex containing Im(2)-tetrad (Quad-Im4) for the first simulation set. A, B, C and D represent the chains. Average and standard deviation (within braces) values are reported.

| HG_A_C | shear | stretch | stagger | buckle | propeller | opening |
|---|---|---|---|---|---|---|
| G-G | -3.43 (0.28) | 1.61 (0.32) | 0.02 (0.37) | -8.9 (7.6) | -22.9 (7.8) | -90.7 (3.7) |
| Im-Im | -3.63 (0.65) | 1.38 (0.78) | -0.19 (0.33) | -17.4 (7.0) | -18.1 (6.1) | -90.8 (4.3) |
| G-G | -3.19 (0.27) | 1.92 (0.34) | -0.04 (0.33) | -8.6 (6.5) | -10.4 (6.5) | -90.0 (2.9) |
| G-G | 3.42 (0.24) | 1.59 (0.30) | 0.07 (0.38) | -6.4 (7.9) | 3.1 (7.7) | -89.8 (2.7) |
| U-U | 2.80 (0.27) | 0.56 (0.31) | 0.25 (0.60) | 20.0 (10.8) | -8.0 (10.5) | 90.1 (5.5) |
| HG_A_D | | | | | | |
| G-G | -3.42 (0.28) | 1.61 (0.32) | -0.03 (0.36) | -9.5 (7.7) | -23.1 (7.7) | 90.6 (3.7) |
| Im-Im | -3.63 (0.64) | 1.38 (0.77) | 0.16 (0.33) | -17.8 (6.8) | -17.8 (6.0) | 90.6 (4.3) |
| G-G | -3.19 (0.27) | 1.92 (0.34) | 0.04 (0.34) | -8.5 (6.4) | -10.1 (6.6) | 90.2 (2.9) |
| G-G | 3.42 (0.24) | 1.59 (0.30) | -0.08 (0.38) | -6.6 (7.9) | 3.5 (7.7) | 89.9 (2.7) |
| U-U | 2.80 (0.28) | 0.57 (0.31) | -0.24 (0.60) | 20.2 (10.7) | -7.7 (10.7) | -90.3 (5.5) |

| | | | | | | |
|---|---|---|---|---|---|---|
| HG_B_C | | | | | | |
| G-G | -3.42 (0.28) | 1.62 (0.32) | -0.02 (0.36) | -9.1 (7.5) | -23.2 (7.6) | 90.8 (3.7) |
| Im-Im | -3.64 (0.65) | 1.38 (0.78) | 0.17 (0.33) | -17.7 (6.9) | -18.1 (6.1) | 90.8 (4.3) |
| G-G | -3.19 (0.27) | 1.92 (0.34) | 0.03 (0.33) | -8.7 (6.4) | -10.1 (6.6) | 90.0 (2.9) |
| G-G | 3.42 (0.24) | 1.59 (0.30) | -0.08 (0.38) | -6.4 (7.9) | 3.6 (7.8) | 89.8 (2.7) |
| U-U | 2.79 (0.27) | 0.57 (0.31) | -0.23 (0.60) | 20.0 (10.6) | -7.6 (10.7) | -90.3 (5.5) |
| HG_B_D | | | | | | |
| G-G | -3.42 (0.28) | 1.62 (0.32) | 0.02 (0.36) | -9.0 (7.5) | -23.3 (7.7) | -90.7 (3.7) |
| Im-Im | -3.63 (0.65) | 1.39 (0.77) | -0.18 (0.33) | -17.7 (7.0) | -18.1 (6.0) | -90.9 (4.2) |
| G-G | -3.20 (0.27) | 1.92 (0.33) | -0.04 (0.33) | -8.5 (6.4) | -10.2 (6.5) | -90.1 (2.9) |
| G-G | 3.42 (0.24) | 1.59 (0.30) | 0.08 (0.38) | -6.5 (7.9) | 3.3 (7.8) | -89.8 (2.7) |
| U-U | 2.80 (0.28) | 0.57 (0.31) | 0.24 (0.60) | 20.2 (10.8) | -7.9 (10.6) | 90.3 (5.5) |

| HG_A_C | shift | slide | rise | tilt | roll | twist |
|---|---|---|---|---|---|---|
| G-G/Im-Im | -1.33 (0.39) | 0.20 (0.35) | 3.47 (0.18) | -0.6 (3.6) | -1.0 (4.1) | 32.3 (6.2) |
| Im-Im/G-G | -0.94 (0.30) | 0.05 (0.43) | 3.09 (0.19) | -0.0 (3.5) | -4.5 (4.1) | 33.2 (5.1) |
| G-G/G-G | -0.80 (0.32) | 0.41 (0.33) | 3.00 (0.15) | 3.5 (3.2) | -4.3 (3.8) | 21.3 (3.3) |
| G-G/U-U | -0.41 (2.47) | -2.87 (0.95) | 0.11 (0.49) | -34.7 (49.8) | 171.8 (13.0) | 1.5 (6.7) |
| HG_A_D | | | | | | |
| G-G/Im-Im | 1.30 (0.39) | -0.20 (0.35) | 3.47 (0.18) | 0.8 (3.6) | 0.9 (4.0) | 32.3 (6.1) |
| Im-Im/G-G | 0.94 (0.30) | -0.06 (0.43) | 3.09 (0.19) | -0.1 (3.5) | 4.4 (4.0) | 33.1 (5.0) |
| G-G/G-G | 0.79 (0.32) | -0.40 (0.34) | 3.00 (0.15) | -3.4 (3.2) | 4.3 (3.7) | 21.3 (3.3) |
| G-G/U-U | 0.45 (2.44) | 2.90 (0.93) | 0.13 (0.50) | 35.7 (49.3) | -171.5 (12.9) | 1.4 (6.7) |
| HG_B_C | | | | | | |
| G-G/Im-Im | 1.32 (0.39) | -0.19 (0.35) | 3.48 (0.18) | 0.7 (3.6) | 0.9 (4.1) | 32.3 (6.2) |
| Im-Im/G-G | 0.93 (0.31) | -0.04 (0.42) | 3.08 (0.19) | 0.0 (3.4) | 4.5 (4.0) | 33.1 (5.1) |
| G-G/G-G | 0.80 (0.32) | -0.39 (0.33) | 3.00 (0.16) | -3.4 (3.2) | 4.2 (3.8) | 21.4 (3.3) |
| G-G/U-U | 0.47 (2.43) | 2.89 (0.93) | 0.13 (0.49) | 35.9 (48.9) | -171.4 (12.8) | 1.4 (6.7) |

| | | | | | | |
|---|---|---|---|---|---|---|
| HG_B_D | | | | | | |
| G-G/Im-Im | -1.32 (0.39) | 0.19 (0.35) | 3.47 (0.17) | -0.7 (3.6) | -0.9 (4.1) | 32.3 (6.1) |
| Im-Im/G-G | -0.92 (0.30) | 0.06 (0.43) | 3.09 (0.19) | 0.1 (3.5) | 4.5 (4.0) | 33.0 (5.0) |
| G-G/G-G | -0.79 (0.32) | 0.40 (0.33) | 3.00 (0.15) | 3.4 (3.1) | -4.3 (3.8) | 21.4 (3.3) |
| G-G/U-U | -0.42 (2.44) | -2.87 (0.94) | 0.12 (0.50) | -34.8 (49.6) | 171.7 (13.0) | 1.4 (6.6) |

**(E)** Structural base pair and base pair step parameters for the Hoogsteen (HG) pairs in the different tetrads within the monomeric quadruplex containing Gm(2)-tetrad (Quad-Gm4) for the first simulation set. A, B, C and D represent the chains. Average and standard deviation (within braces) values are reported.

| HG_A_C | shear | stretch | stagger | buckle | propeller | opening |
|---|---|---|---|---|---|---|
| G-G | -3.01 (0.47) | 1.83 (0.44) | -0.59 (0.85) | 4.3 (19.7) | -10.7 (11.0) | -86.8 (6.8) |
| Gm-Gm | -3.09 (0.41) | 2.17 (0.48) | -0.09 (0.33) | -18.3 (7.1) | -24.9 (7.3) | -91.5 (3.7) |
| G-G | -3.47 (0.20) | 1.54 (0.23) | -0.14 (0.35) | -9.6 (6.8) | -8.4 (7.3) | -90.0 (2.8) |
| G-G | 3.29 (0.23) | 1.78 (0.29) | 0.02 (0.42) | -7.2 (8.5) | 3.7 (8.5) | -89.9 (2.8) |
| U-U | 2.77 (0.28) | 0.59 (0.31) | 0.30 (0.63) | 20.2 (11.0) | -6.2 (11.1) | 89.8 (5.8) |
| HG_A_D | | | | | | |
| G-G | -2.91 (0.54) | 2.12 (0.60) | -0.71 (0.98) | -17.4 (11.7) | -3.4 (18.0) | 90.2 (5.0) |
| Gm-Gm | -3.11 (0.40) | 2.10 (0.47) | 0.04 (0.34) | -16.5 (7.7) | -19.5 (6.8) | 90.3 (3.9) |
| G-G | -3.48 (0.20) | 1.52 (0.23) | 0.08 (0.35) | -6.2 (7.1) | -5.6 (7.2) | 89.9 (2.8) |
| G-G | 3.30 (0.23) | 1.78 (0.29) | -0.10 (0.41) | -9.6 (8.6) | 5.4 (8.6) | 90.0 (2.8) |
| U-U | 2.77 (0.28) | 0.63 (0.31) | -0.28 (0.64) | 22.8 (11.1) | -5.7 (11.4) | -90.7 (5.8) |
| HG_B_C | | | | | | |
| G-G | -3.10 (0.42) | 2.01 (0.46) | 0.09 (0.52) | -9.6 (9.9) | -18.9 (8.8) | 89.9 (4.4) |
| Gm-Gm | -3.09 (0.40) | 2.14 (0.47) | 0.03 (0.33) | -20.1 (7.1) | -20.0 (6.7) | 90.9 (3.6) |
| G-G | -3.48 (0.21) | 1.52 (0.23) | 0.03 (0.35) | -7.5 (6.7) | -3.9 (7.5) | 89.8 (2.8) |
| G-G | 3.29 (0.23) | 1.78 (0.30) | -0.16 (0.42) | -9.0 (8.4) | 6.8 (8.6) | 89.7 (2.8) |
| U-U | 2.75 (0.27) | 0.62 (0.31) | -0.20 (0.64) | 22.2 (10.8) | -3.7 (11.2) | -90.1 (5.8) |
| HG_B_D | | | | | | |

| | | | | | | |
|---|---|---|---|---|---|---|
| G-G | -3.28 (0.37) | 1.83 (0.42) | -0.05 (0.50) | -10.6 (9.8) | -18.3 (8.5) | -90.2 (4.2) |
| Gm-Gm | -3.06 (0.41) | 2.18 (0.49) | -0.04 (0.33) | -19.0 (6.9) | -21.2 (6.5) | -91.1 (3.7) |
| G-G | -3.47 (0.20) | 1.53 (0.23) | -0.08 (0.35) | -8.0 (6.7) | -7.1 (7.2) | -89.9 (2.8) |
| G-G | 3.29 (0.23) | 1.77 (0.29) | 0.08 (0.41) | -8.7 (8.5) | 4.4 (8.7) | -89.7 (2.8) |
| U-U | 2.75 (0.27) | 0.63 (0.31) | 0.27 (0.63) | 22.0 (11.0) | -5.5 (11.3) | 90.3 (5.7) |

| HG_A_C | shift | slide | rise | tilt | roll | twist |
|---|---|---|---|---|---|---|
| G-G/Gm-Gm | -1.33 (0.48) | 0.80 (0.72) | 3.76 (0.27) | -7.0 (5.6) | -4.3 (6.0) | 34.2 (4.3) |
| Gm-Gm/G-G | -1.22 (0.27) | 0.46 (0.32) | 3.04 (0.17) | 2.4 (3.6) | -4.2 (4.1) | 26.1 (3.7) |
| G-G/G-G | -0.57 (0.31) | 0.32 (0.33) | 3.06 (0.15) | 2.7 (3.2) | -3.6 (4.1) | 24.1 (2.4) |
| G-G/U-U | -0.29 (2.59) | -2.67 (0.98) | 0.07 (0.54) | -28.4 (49.1) | 173.4 (12.6) | 1.4 (6.8) |
| HG_A_D | | | | | | |
| G-G/Gm-Gm | 0.53 (0.80) | -0.10 (0.42) | 3.82 (0.30) | 1.9 (4.4) | -5.5 (8.8) | 35.0 (4.0) |
| Gm-Gm/G-G | 1.29 (0.28) | -0.38 (0.32) | 3.02 (0.17) | -2.9 (3.6) | 4.8 (4.1) | 26.0 (3.7) |
| G-G/G-G | 0.61 (0.32) | -0.25 (0.33) | 3.06 (0.15) | -3.2 (3.2) | 4.7 (4.0) | 24.0 (2.4) |
| G-G/U-U | 0.36 (2.51) | 2.75 (0.96) | 0.12 (0.58) | 30.4 (48.2) | 173.4 (12.6) | 1.4 (6.8) |
| HG_B_C | | | | | | |
| G-G/Gm-Gm | 1.27 (0.46) | -0.19 (0.35) | 3.68 (0.21) | 4.9 (4.2) | 0.2 (4.6) | 35.9 (4.1) |
| Gm-Gm/G-G | 1.25 (0.27) | -0.35 (0.31) | 3.03 (0.17) | -1.8 (3.6) | 4.1 (4.1) | 25.7 (3.7) |
| G-G/G-G | 0.62 (0.31) | -0.19 (0.33) | 3.06 (0.15) | -2.9 (3.2) | 5.0 (4.0) | 23.9 (2.4) |
| G-G/U-U | 0.53 (2.40) | 2.80 (0.92) | 0.15 (0.57) | 33.4 (45.9) | -171.9 (12.0) | 1.1 (6.8) |
| HG_B_D | | | | | | |
| G-G/Gm-Gm | -1.18 (0.41) | 0.08 (0.39) | 3.60 (0.20) | -3.7 (3.9) | -1.5 (4.4) | 35.8 (4.0) |
| Gm-Gm/G-G | -1.21 (0.27) | 0.45 (0.33) | 3.02 (0.17) | 2.5 (3.6) | -4.8 (4.1) | 25.8 (3.7) |
| G-G/G-G | -0.53 (0.32) | 0.29 (0.33) | 3.07 (0.15) | 3.1 (3.1) | -4.1 (4.0) | 24.0 (2.4) |
| G-G/U-U | -0.25 (2.59) | -2.67 (0.97) | 0.07 (0.56) | -27.3 (49.1) | 173.6 (12.7) | 1.3 (6.8) |

**(F)** Structural base pair and base pair step parameters for the Hoogsteen (HG) pairs in the different tetrads within the monomeric quadruplex containing Am(2)-tetrad (Quad-Am4) for the first simulation set. A, B, C and D represent the chains. Average and standard deviation (within braces) values are reported.

| HG_A_C | shear | stretch | stagger | buckle | propeller | opening |
|---|---|---|---|---|---|---|
| G-G | -3.24 (0.38) | 1.85 (0.44) | -0.04 (0.50) | -9.5 (10.1) | -15.3 (9.3) | -90.2 (4.3) |
| Am-Am | -3.33 (0.22) | 2.10 (0.30) | -0.08 (0.44) | -15.7 (8.0) | -20.5 (7.4) | -90.7 (4.9) |
| G-G | -3.12 (0.20) | 2.00 (0.24) | -0.05 (0.36) | -8.1 (6.9) | -9.9 (7.0) | -90.0 (3.0) |
| G-G | 3.51 (0.19) | 1.45 (0.22) | 0.07 (0.38) | -6.7 (7.8) | 2.9 (7.9) | -89.8 (2.6) |
| U-U | 2.77 (0.26) | 0.60 (0.29) | 0.26 (0.61) | 21.2 (11.0) | -10.5 (10.8) | 90.2 (5.3) |
| HG_A_D | | | | | | |
| G-G | -3.23 (0.38) | 1.86 (0.45) | 0.01 (0.49) | -10.6 (10.1) | -15.4 (9.3) | 90.0 (4.3) |
| Am-Am | -3.33 (0.22) | 2.09 (0.29) | 0.03 (0.44) | -16.1 (8.1) | -20.0 (7.3) | 90.6 (5.0) |
| G-G | -3.12 (0.20) | 2.00 (0.23) | 0.02 (0.36) | -8.1 (6.9) | -9.3 (7.1) | 90.1 (3.0) |
| G-G | 3.51 (0.19) | 1.45 (0.22) | -0.09 (0.38) | -7.1 (7.8) | 3.6 (7.9) | 89.9 (2.6) |
| U-U | 2.77 (0.27) | 0.61 (0.30) | -0.24 (0.61) | 21.4 (11.1) | -9.8 (10.8) | -90.5 (5.3) |
| HG_B_C | | | | | | |
| G-G | -3.22 (0.38) | 1.86 (0.45) | 0.02 (0.49) | -10.2 (10.1) | -16.0 (9.5) | 90.0 (4.2) |
| Am-Am | -3.33 (0.22) | 2.09 (0.30) | 0.06 (0.44) | -16.4 (8.0) | -20.3 (7.3) | 90.8 (5.0) |
| G-G | -3.12 (0.20) | 2.00 (0.23) | 0.03 (0.36) | -8.3 (6.8) | -9.5 (7.0) | 90.0 (3.0) |
| G-G | 3.51 (0.19) | 1.45 (0.22) | -0.10 (0.38) | -6.8 (7.8) | 3.5 (8,0) | 89.8 (2.6) |
| U-U | 2.77 (0.27) | 0.61 (0.30) | -0.23 (0.60) | 21.1 (11.0) | -9.9 (10.7) | -90.4 (5.3) |
| HG_B_D | | | | | | |
| G-G | -3.25 (0.38) | 1.85 (0.44) | -0.01 (0.49) | -10.1 (10.1) | -15.7 (9.4) | -90.1 (4.3) |
| Am-Am | -3.32 (0.23) | 2.10 (0.30) | -0.04 (0.44) | -16.0 (8.0) | -20.1 (7.3) | -90.9 (5.0) |
| G-G | -3.12 (0.20) | 2.00 (0.23) | -0.03 (0.36) | -8.3 (6.9) | -9.8 (6.9) | -90.0 (3.0) |
| G-G | 3.51 (0.19) | 1.45 (0.22) | 0.09 (0.38) | -6.6 (7.7) | 3.61(8.0) | -89.9 (2.6) |
| U-U | 2.77 (0.27) | 0.61 (0.30) | 0.24 (0.60) | 21.1 (11.0) | -10.2 (10.8) | 90.3 (5.3) |

| HG_A_C | shift | slide | rise | tilt | roll | twist |
|---|---|---|---|---|---|---|

| | | | | | | |
|---|---|---|---|---|---|---|
| G-G/Am-Am | -1.11 (0.42) | 0.35 (0.40) | 3.54 (0.22) | -3.2 (4.1) | -1.5 (5.2) | 34.4 (3.8) |
| Am-Am/G-G | -1.05 (0.29) | 0.21 (0.28) | 3.10 (0.18) | 1.1 (4.0) | -4.1 (4.5) | 28.8 (2.6) |
| G-G/G-G | -0.89 (0.29) | 0.49 (0.31) | 3.01 (0.15) | 3.3 (3.2) | -4.2 (4.0) | 20.9 (2.3) |
| G-G/U-U | -0.46 (2.42) | -2.94 (0.94) | 0.09 (0.41) | -37.5 (49.8) | 171.0 (13.2) | 1.7 (6.7) |
| HG_A_D | | | | | | |
| G-G/Am-Am | 1.03 (0.43) | -0.32 (0.39) | 3.55 (0.22) | 3.3 (4.2) | 1.2 (5.1) | 34.5 (3.7) |
| Am-Am/G-G | 1.03 (0.28) | -0.20 (0.29) | 3.11 (0.18) | -1.2 (4.0) | 3.9 (4.5) | 28.6 (2.6) |
| G-G/G-G | 0.89 (0.28) | -0.47 (0.31) | 3.01 (0.15) | -3.3 (3.2) | 4.2 (3.9) | 20.9 (2.3) |
| G-G/U-U | 0.54 (2.35) | 2.98 (0.92) | 0.10 (0.42) | 39.1 (48.6) | -170.3 (13.0) | 1.8 (6.8) |
| HG_B_C | | | | | | |
| G-G/Am-Am | 1.09 (0.42) | -0.31 (0.38) | 3.55 (0.22) | 3.3 (4.2) | 1.2 (5.1) | 34.4 (3.7) |
| Am-Am/G-G | 1.04 (0.28) | -0.19 (0.28) | 3.10 (0.18) | -1.0 (4.0) | 4.0 (4.5) | 28.7 (2.6) |
| G-G/G-G | 0.89 (0.28) | -0.46 (0.31) | 3.01 (0.15) | -3.2 (3.2) | 4.2 (3.9) | 20.9 (2.3) |
| G-G/U-U | 0.55 (2.35) | 2.98 (0.91) | 0.10 (0.42) | 39.3 (48.5) | -170.3 (12.9) | 1.8 (6.6) |
| HG_B_D | | | | | | |
| G-G/Am-Am | -1.07 (0.42) | 0.30 (0.38) | 3.55 (0.22) | -3.3 (4.1) | -1.3 (5.2) | 34.5 (3.7) |
| Am-Am/G-G | -1.03 (0.28) | 0.20 (0.28) | 3.10 (0.18) | 1.3 (4.0) | -4.2 (4.5) | 28.6 (2.6) |
| G-G/G-G | -0.88 (0.29) | 0.48 (0.30) | 3.01 (0.15) | 3.4 (3.2) | -4.3 (3.9) | 20.9 (2.3) |
| G-G/U-U | -0.47 (2.41) | -2.95 (0.94) | 0.09 (0.42) | -37.8 (49.6) | 170.9 (13.2) | 1.8 (6.7) |

**Table S8. (A)** Interaction energies (kcal/mol) for hydrogen bonding, stacking and ion interactions for the m$^6$A-tetrad at position 2 of the studied monomeric quadruplex for the three simulation sets. Average and standard deviation (within braces) values are reported.

| *Simulation set 1* | | *ELEC* | *VDW* | *TOTAL* |
|---|---|---|---|---|
| *Hydrogen bonding energy of the tetrads* | | | | |
| *m$^6$A-m$^6$A-m$^6$A-m$^6$A* | *2-13*<br>*2-19*<br>*7-19*<br>*7-13* | *-5.42 (1.07)*<br>*-5.39 (1.07)*<br>*-5.42 (1.06)*<br>*-5.42 (1.07)* | *-1.20 (0.60)*<br>*-1.21 (0.59)*<br>*-1.20 (0.59)*<br>*-1.20 (0.60)* | *-6.62 (0.81)*<br>*-6.60 (0.82)*<br>*-6.62 (0.81)*<br>*-6.62 (0.82)* |
| *Stacking energy between tetrads* | | | | |

| | | | |
|---|---|---|---|
| *G4-m⁶A4*<br>*m⁶A4-G4* | *-9.68 (2.37)*<br>*-6.83 (1.70)* | *-38.92 (1.53)*<br>*-36.73 (1.54)* | *-48.61 (2.56)*<br>*-43.56 (1.71)* |
| *Interaction energy between a tetrad and the proximal K+ cation* | | | |
| *m⁶A-m⁶A-m⁶A-m⁶A & K+*<br>*K24*<br>*K23* | *11.65 (3.77)*<br>*11.74 (3.78)* | *-0.007 (0.03)*<br>*-0.007 (0.03)* | *11.65 (3.75)*<br>*11.74 (3.76)* |

| *Simulation set 2* | | *ELEC* | *VDW* | *TOTAL* |
|---|---|---|---|---|
| *Hydrogen bonding energy of the tetrads* | | | | |
| *m⁶A-m⁶A-m⁶A-m⁶A* | *2-13*<br>*2-19*<br>*7-19*<br>*7-13* | *-5.43 (1.07)*<br>*-5.41 (1.06)*<br>*-5.43 (1.06)*<br>*-5.43 (1.06)* | *-1.20 (0.60)*<br>*-1.21 (0.60)*<br>*-1.20 (0.60)*<br>*-1.19 (0.60)* | *-6.62 (0.82)*<br>*-6.61 (0.81)*<br>*-6.63 (0.81)*<br>*-6.62 (0.81)* |
| *Stacking energy between tetrads* | | | | |
| *G4-m⁶A4*<br>*m⁶A4-G4* | | *-9.57 (2.34)*<br>*-6.83 (1.70)* | *-38.94 (1.53)*<br>*-36.73 (1.54)* | *-48.51 (2.56)*<br>*-43.62 (1.69)* |
| *Interaction energy between a tetrad and the proximal K+ cation* | | | | |
| *m⁶A-m⁶A-m⁶A-m⁶A & K+*<br>*K24*<br>*K23* | | *13.78 (4.90)*<br>*12.14 (4.16)* | *-0.39 (0.74)*<br>*-0.01 (0.06)* | *13.39 (4.53)*<br>*12.13 (4.13)* |

| *Simulation set 3* | | *ELEC* | *VDW* | *TOTAL* |
|---|---|---|---|---|
| *Hydrogen bonding energy of the tetrads* | | | | |
| *m⁶A-m⁶A-m⁶A-m⁶A* | *2-13*<br>*2-19*<br>*7-19*<br>*7-13* | *-5.40 (1.07)*<br>*-5.38 (1.07)*<br>*-5.40 (1.07)*<br>*-5.42 (1.07)* | *-1.21 (0.59)*<br>*-1.22 (0.58)*<br>*-1.21 (0.59)*<br>*-1.20 (0.60)* | *-6.61 (0.82)*<br>*-6.60 (0.82)*<br>*-6.62 (0.81)*<br>*-6.61 (0.82)* |
| *Stacking energy between tetrads* | | | | |
| *G4-m⁶A4*<br>*m⁶A4-G4* | | *-9.73 (2.35)*<br>*-6.84 (1.70)* | *-38.91 (1.53)*<br>*-36.66 (1.54)* | *-48.64 (2.55)*<br>*-43.51 (1.72)* |
| *Interaction energy between a tetrad and the proximal K+ cation* | | | | |
| *m⁶A-m⁶A-m⁶A-m⁶A & K+*<br>*K24*<br>*K23* | | *11.74 (3.82)*<br>*11.83 (3.83)* | *-0.008 (0.04)*<br>*-0.007 (0.04)* | *11.74 (3.80)*<br>*11.82 (3.82)* |

**(B)** Comparison of Gibbs free energies (obtained from MM-GBSA analysis) of the quadruplex containing m$^6$A-tetrad at position 2 for the three simulation sets. Average and standard deviation (within braces) values are reported.

| | *Free energy (Kcal/mol)* |
|---|---|
| *Simulation set 1* | *-3281.61 (19.50)* |
| *Simulation set 2* | *-3282.05 (19.30)* |
| *Simulation set 3* | *-3281.76 (19.41)* |

**Table S9.** Frequencies (in %) of the Hoogsteen (HG) hydrogen bonds between the bases within different tetrads of the studied monomeric quadruplex containing m$^6$A-tetrad at position 2 for the three simulation sets.

| H-bonds | | Simulation set 1 | Simulation set 2 | Simulation set 3 |
|---|---|---|---|---|
| *1-11* | *(11)N2-H21- - -N7(1)*<br>*(11)N1-H1- - -O6(1)* | 60.23<br>52.73 | 61.71<br>50.66 | 59.78<br>52.83 |
| *2-12* | *(2)N6-H16 - - - N3(12)* | 10.27 | 10.41 | 9.96 |
| *3-13* | *(13)N2-H21- - -N7(3)*<br>*(13)N1-H1- - -O6(3)* | 57.51<br>65.21 | 59.32<br>62.88 | 55.54<br>66.06 |
| *4-14* | *(14)N2-H21- - -N7(4)*<br>*(14)N1-H1- - -O6(4)* | 65.75<br>37.56 | 65.31<br>34.56 | 66.07<br>35.33 |
| *5-15* | *(5)N3-H3- - -O4(15)* | 76.70 | 76.30 | 76.88 |
| *1-16* | *(1)N2-H21- - -N7(16)*<br>*(1)N1-H1- - -O6(16)* | 60.51<br>51.93 | 61.92<br>50.29 | 60.05<br>52.65 |
| *2-17* | *(17)N6-H16 - - - N3(2)* | 9.91 | 10.22 | 9.60 |
| *3-18* | *(3)N2-H21- - -N7(18)*<br>*(3)N1-H1- - -O6(18)* | 57.27<br>64.57 | 59.09<br>62.39 | 55.85<br>65.61 |
| *4-19* | *(4)N2-H21- - -N7(19)*<br>*(4)N1-H1- - -O6(19)* | 65.03<br>36.35 | 64.78<br>34.96 | 65.52<br>34.96 |
| *5-20* | *(20)N3-H3- - -O4(5)* | 76.82 | 76.76 | 76.93 |

| | | | | |
|---|---|---|---|---|
| *6-16* | *(16)N2-H21- - -N7(6)*<br>*(16)N1-H1- - -O6(6)* | 60.72<br>52.14 | 61.91<br>50.50 | 60.31<br>52.90 |
| *7-17* | *(7)N6-H16 - - - N3(17)* | 10.36 | 10.39 | 9.89 |
| *8-18* | *(18)N2-H21- - -N7(8)*<br>*(18)N1-H1- - -O6(8)* | 57.53<br>65.15 | 59.24<br>62.76 | 56.15<br>65.72 |
| *9-19* | *(19)N2-H21- - -N7(9)*<br>*(19)N1-H1- - -O6(9)* | 65.06<br>37.35 | 65.22<br>34.83 | 65.39<br>34.53 |
| *10-20* | *(10)N3-H3- - -O4(20)* | 76.79 | 76.53 | 76.87 |
| *6-11* | *(6)N2-H21- - -N7(11)*<br>*(6)N1-H1- - -O6(11)* | 60.27<br>52.16 | 61.74<br>50.66 | 60.01<br>53.09 |
| *7-12* | *(12)N6-H16 - - - N3(7)* | 10.42 | 10.46 | 10.07 |
| *8-13* | *(8)N2-H21- - -N7(13)*<br>*(8)N1-H1- - -O6(13)* | 58.08<br>65.23 | 59.17<br>62.70 | 56.04<br>66.01 |
| *9-14* | *(9)N2-H21- - -N7(14)*<br>*(9)N1-H1- - -O6(14)* | 65.43<br>36.73 | 65.61<br>34.97 | 65.66<br>34.68 |
| *10-15* | *(15)N3-H3- - -O4(10)* | 76.59 | 76.25 | 76.95 |

**Table S10.** Comparison of N9-pyramidalization (based on the average and standard deviation values for Kappa' (***κ***') dihedral angles (in degrees)) for the tetrads at position 2, 3 and 4 respectively (from the 5′-direction) of the studied monomeric quadruplex containing m$^6$A-tetrad at position 2 for the three simulation sets.

| TETRAD-2 | Simulation set-1 | Simulation set-2 | Simulation set-3 |
|---|---|---|---|
| 2 | 2.31 (12.09) | 2.40 (12.07) | 2.40 (12.07) |
| 7 | 2.36 (12.11) | 2.49 (12.11) | 2.49 (12.11) |
| 12 | 2.47 (12.17) | 2.54 (12.16) | 2.54 (12.16) |
| 17 | 2.43 (12.14) | 2.47 (12.08) | 2.47 (12.08) |
| TETRAD-3 | | | |
| 3 | -0.97 (10.59) | -1.16 (10.57) | -1.16 (10.56) |
| 8 | -1.04 (10.55) | -1.24 (10.56) | -1.24 (10.56) |
| 13 | -1.03 (10.63) | -1.24 (10..58) | -1.24 (10.58) |

| | | | |
|---|---|---|---|
| 18 | -1.03 (10.57) | -1.19 (10.59) | -1.19 (10.59) |
| TETRAD-4 | | | |
| 4 | -6.16 (10.56) | -6.55 (10.63) | -6.55 (10.63) |
| 9 | -6.40 (10.64) | -6.47 (10.59) | -6.47 (10.59) |
| 14 | -6.55 (10.63) | -6.42 (10.58) | -6.42 (10.58) |
| 19 | -6.32 (10.59) | -6.37 (10.59) | -6.37 (10.59) |

**Table S11. (A)** Interaction energies (kcal/mol) for hydrogen bonding, stacking and ion interactions for the different purine tetrads at position 2 of the dimeric quadruplexes (M1: monomer 1 and M2: monomer 2) for the first set of simulations. Average and standard deviation (within braces) values are reported.

| | | *ELEC* | *VDW* | *TOTAL* |
|---|---|---|---|---|
| *Hydrogen bonding energy of the tetrads* | | | | |
| *I-I-I-I M1* | *2-13*<br>*2-19*<br>*7-19*<br>*7-13* | *0.71 (1.34)*<br>*0.49 (1.40)*<br>*1.65 (1.46)*<br>*-0.27 (1.39)* | *-1.30 (0.76)*<br>*-1.13 (0.86)*<br>*-1.53 (0.57)*<br>*-0.91 (0.98)* | *-0.59 (1.54)*<br>*-0.64 (1.64)*<br>*0.12 (1.57)*<br>*-1.18 (1.70)* |
| *G-G-G-G M1* | *2-13*<br>*2-19*<br>*7-19*<br>*7-13* | *-9.49 (2.33)*<br>*-8.95 (2.25)*<br>*-9.32 (2.39)*<br>*-9.95 (2.26)* | *-0.92 (1.16)*<br>*-0.79 (1.21)*<br>*-0.76 (1.22)*<br>*-0.53 (1.27)* | *-10.41 (2.60)*<br>*-9.74 (2.55)*<br>*-10.08 (2.68)*<br>*-10.48 (2.59)* |
| *A-A-A-A M1* | *2-13*<br>*2-19*<br>*7-19*<br>*7-13* | *-3.89 (1.37)*<br>*-4.13 (1.32)*<br>*-4.06 (1.34)*<br>*-4.09 (1.32)* | *-0.75 (0.97)*<br>*-0.61 (1.01)*<br>*-0.59 (1.06)*<br>*-0.73 (0.99)* | *-4.64 (1.68)*<br>*-4.74 (1.66)*<br>*-4.65 (1.71)*<br>*-4.82 (1.65)* |
| *G-U5 M1 (I)* | *6-11* | *-6.98 (1.29)* | *0.03 (0.99)* | *-6.95 (1.63)* |
| *G-U5 M1 (G)* | *6-11* | *-6.61 (1.44)* | *-0.006 (0.96)* | *-6.62 (1.73)* |
| *G-U5 M1 (A)* | *6-11* | *-6.76 (1.65)* | *0.15 (1.02)* | *-6.61 (1.94)* |
| *I-I-I-I M2* | *25-42*<br>*42-31*<br>*31-37*<br>*25-37* | *0.73 (1.43)*<br>*-0.08 (1.46)*<br>*0.72 (1.56)*<br>*1.36 (1.46)* | *-1.46 (0.64)*<br>*-0.94 (0.98)*<br>*-1.05 (0.89)*<br>*-1.52 (0.59)* | *-0.73 (1.56)*<br>*-1.02 (1.76)*<br>*-0.33 (1.79)*<br>*-0.16 (1.57)* |
| *G-G-G-G M2* | *25-42*<br>*42-31*<br>*31-37*<br>*25-37* | *-8.31 (2.29)*<br>*-8.34 (2.37)*<br>*-8.49 (2.24)*<br>*-9.51 (2.18)* | *-1.09 (1.09)*<br>*-1.10 (1.10)*<br>*-1.06 (1.09)*<br>*-0.87 (1.15)* | *-9.40 (2.54)*<br>*-9.44 (2.61)*<br>*-9.55 (2.49)*<br>*-10.38 (2.46)* |

| | | | | |
|---|---|---|---|---|
| *A-A-A-A M2* | *25-42<br>42-31<br>31-37<br>25-37* | *-3.49 (1.29)<br>-3.16 (1.35)<br>-3.09 (1.44)<br>-3.86 (1.29)* | *-0.41 (1.08)<br>-0.62 (1.03)<br>-0.70 (0.99)<br>-0.43 (1.09)* | *-3.90 (1.68)<br>-3.78 (1.69)<br>-3.79 (1.75)<br>-4.29 (1.69)* |
| *G-U5 M2 (I)* | *30-35* | *-5.90 (2.11)* | *-0.15 (0.91)* | *-6.05 (2.29)* |
| *G-U5 M2 (I)* | *30-35* | *-4.50 (2.54)* | *-0.98 (0.76)* | *-5.18 (2.65)* |
| *G-U5 M2 (I)* | *30-35* | *-1.54 (1.84)* | *-0.61 (0.49)* | *-2.15 (1.90)* |
| *Stacking energy between tetrads* | | | | |
| *G4-I4<br>I4-G4 M1* | | *29.00 (2.67)<br>29.82 (3.37)* | *-39.31 (1.47)<br>-39.51 (1.26)* | *-10.31 (3.05)<br>-9.69 (3.59)* |
| *G4-G4<br>G4-G4 M1* | | *33.36 (2.74)<br>33.58 (2.75)* | *-43.53 (1.20)<br>-43.28 (1.13)* | *-10.17 (2.99)<br>-9.70 (2.97)* |
| *G4-A4<br>A4-G4 M1* | | *6.62 (3.09)<br>7.44 (2.79)* | *-39.44 (1.38)<br>-38.69 (1.48)* | *-32.82 (3.38)<br>-31.25 (3.16)* |
| *G4-I4<br>I4-G4 M2* | | *29.18 (2.74)<br>31.25 (2.72)* | *-39.52 (1.44)<br>-39.76 (1.20)* | *-10.34 (3.09)<br>-8.51 (2.97)* |
| *G4-G4<br>G4-G4 M2* | | *34.58 (2.71)<br>36.80 (2.64)* | *-42.82 (1.43)<br>-43.07 (1.24)* | *-8.24 (3.06)<br>-6.27 (2.92)* |
| *G4-A4<br>A4-G4 M2* | | *19.67 (4.84)<br>-7.33 (5.44)* | *-39.09 (1.36)<br>-38.01 (1.71)* | *-19.42 (5.03)<br>-45.34 (5.70)* |
| *G4-G4 (5') (I)<br>U-G4-G4-U (5')* | | *17.88 (5.86)<br>25.29 (3.78)* | *-36.93 (5.97)<br>-43.55 (2.56)* | *-19.05 (8.36)<br>-18.26 (4.56)* |
| *G4-G4 (5') (G)<br>U-G4-G4-U (5')* | | *3.27 (1.79)<br>15.38 (2.89)* | *-19.02 (1.69)<br>-30.93 (2.77)* | *-15.75 (2.46)<br>-15.61 (4.00)* |
| *G4-G4 (5') (A)<br>U-G4-G4-U (5')* | | *5.84 (2.88)<br>18.81 (3.64)* | *-19.65 (1.82)<br>-32.09 (2.92)* | *-13.81 (3.41)<br>-13.28 (4.67)* |
| *Interaction energy between a tetrad and the proximal K+ cation* | | | | |
| *I-I-I-I & K+<br>K48<br>K49 M1* | | *<br>-106.63 (6.61)<br>-96.55 (5.97)* | *<br>3.37 (2.31)<br>1.16 (1.56)* | *<br>-103.26 (7.00)<br>-95.39 (6.17)* |
| *G-G-G-G & K+<br>K48<br>K49 M1* | | *<br>-100.28 (7.03)<br>-92.64 (6.82)* | *<br>1.87 (2.08)<br>0.90 (1.56)* | *<br>-98.41 (7.33)<br>-91.74 (6.99)* |
| *A-A-A-A and K+<br>K48<br>K49 M1* | | *<br>-8.72 (5.78)<br>-21.86 (2.42)* | *<br>0.002 (0.18)<br>-0.46 (0.07)* | *<br>-8.72 (5.78)<br>-22.32 (2.42)* |

| | | | |
|---|---|---|---|
| *I-I-I-I & K+*<br>*K51*<br>*K52 M2* | <br>*-94.85 (5.61)*<br>*-109.94 (4.57)* | <br>*0.84 (1.38)*<br>*4.38 (2.01)* | <br>*-94.01 (5.78)*<br>*-105.56 (4.99)* |
| *G-G-G-G & K+*<br>*K51*<br>*K52 M2* | <br>*-93.44 (6.48)*<br>*-109.47 (5.42)* | <br>*0.81 (1.47)*<br>*4.66 (2.09)* | <br>*-92.63 (6.64)*<br>*-104.81 (5.81)* |
| *A-A-A-A & K+*<br>*K51*<br>*K52 M2* | <br>*-46.28 (7.55)*<br>*-10.25 (4.86)* | <br>*-1.30 (0.73)*<br>*-0.41 (0.22)* | <br>*-47.58 (7.59)*<br>*-10.65 (4.86)* |

**(B)** Interaction energies (kcal/mol) for hydrogen bonding, stacking and ion interactions for the different purine tetrads at position 2 of the dimeric quadruplexes (M1: monomer 1 and M2: monomer 2) for the second set of simulations. Average and standard deviation (within braces) values are reported.

| | | *ELEC* | *VDW* | *TOTAL* |
|---|---|---|---|---|
| *Hydrogen bonding energy of the tetrads* | | | | |
| *I-I-I-I M1* | *2-13*<br>*2-19*<br>*7-19*<br>*7-13* | *0.60 (1.26)*<br>*-0.36 (1.33)*<br>*2.02 (1.38)*<br>*0.77 (1.36)* | *-1.49 (0.62)*<br>*-0.83 (0.97)*<br>*-1.46 (0.64)*<br>*-1.31 (0.73)* | *-0.89 (1.40)*<br>*-1.19 (1.65)*<br>*0.56 (1.52)*<br>*-0.54 (1.54)* |
| *G-G-G-G M1* | *2-13*<br>*2-19*<br>*7-19*<br>*7-13* | *-8.84 (2.36)*<br>*-8.76 (2.29)*<br>*-8.69 (2.36)*<br>*-9.74 (2.32)* | *-1.14 (1.08)*<br>*-0.99 (1.12)*<br>*-0.85 (1.19)*<br>*-0.64 (1.24)* | *-9.98 (2.59)*<br>*-9.75 (2.55)*<br>*-9.54 (2.64)*<br>*-10.38 (2.63)* |
| *A-A-A-A M1* | *2-13*<br>*2-19*<br>*7-19*<br>*7-13* | *-4.06 (1.36)*<br>*-4.15 (1.33)*<br>*-4.05 (1.35)*<br>*-4.25 (1.34)* | *-0.71 (0.99)*<br>*-0.66 (1.02)*<br>*-0.58 (1.04)*<br>*-0.67 (1.03)* | *-4.77 (1.68)*<br>*-4.81 (1.68)*<br>*-4.63 (1.70)*<br>*-4.92 (1.69)* |
| *G-U5 M1 (I)* | *6-11* | *-6.70 (1.31)* | *-0.03 (0.95)* | *-6.73 (1.62)* |
| *G-U5 M1 (G)* | *6-11* | *-6.62 (1.33)* | *-0.06 (0.95)* | *-6.68 (1.63)* |
| *G-U5 M1 (A)* | *6-11* | *-5.31 (2.46)* | *-0.38 (0.82)* | *-5.69 (2.59)* |
| *I-I-I-I M2* | *25-42*<br>*42-31*<br>*31-37*<br>*25-37* | *0.85 (1.40)*<br>*0.37 (1.53)*<br>*0.01 (1.28)*<br>*-1.24 (1.08)* | *-1.54 (0.63)*<br>*-1.43 (0.66)*<br>*-1.42 (0.67)*<br>*-1.21 (0.80)* | *-0.69 (1.54)*<br>*-1.06 (1.67)*<br>*-1.41 (1.44)*<br>*-2.45 (1.34)* |
| *G-G-G-G M2* | *25-42*<br>*42-31*<br>*31-37*<br>*25-37* | *-9.15 (2.22)*<br>*-9.11 (2.32)*<br>*-9.08 (2.21)*<br>*-10.27 (2.17)* | *-0.78 (1.20)*<br>*-0.89 (1.17)*<br>*-0.94 (1.14)*<br>*-0.61 (1.24)* | *-9.93 (2.52)*<br>*-10.00 (2.59)*<br>*-10.02 (2.49)*<br>*-10.88 (2.50)* |

| | | | | |
|---|---|---|---|---|
| *A-A-A-A M2* | *25-42<br>42-31<br>31-37<br>25-37* | *-3.89 (1.23)<br>-3.28 (1.45)<br>-3.25 (1.61)<br>-3.69 (1.21)* | *-0.64 (0.95)<br>-0.77 (0.85)<br>-0.77 (0.81)<br>-0.76 (0.85)* | *-4.53 (1.55)<br>-4.05 (1.68)<br>-4.02 (1.80)<br>-4.45 (1.48)* |
| *G-U5 M2 (I)* | *30-35* | *-6.20 (2.08)* | *-0.64 (0.98)* | *-6.84 (2.29)* |
| *G-U5 M2 (G)* | *30-35* | *-1.85 (3.42)* | *-1.86 (0.96)* | *-3.71 (3.55)* |
| *G-U5 M2 (A)* | *30-35* | *-0.29 (1.00)* | *-0.24 (0.21)* | *-0.53 (1.02)* |
| *Stacking energy between tetrads* | | | | |
| *G4-I4<br>I4-G4 M1* | | *29.47 (2.83)<br>29.29 (3.09)* | *-39.33 (1.39)<br>-39.89 (1.16)* | *-9.86 (3.15)<br>-10.60 (3.30)* |
| *G4-G4<br>G4-G4 M1* | | *33.43 (2.81)<br>33.23 (2.63)* | *-43.33 (1.21)<br>-43.20 (1.21)* | *-9.90 (3.06)<br>-9.97 (2.89)* |
| *G4-A4<br>A4-G4 M1* | | *6.69 (3.29)<br>7.13 (2.94)* | *-39.05 (1.52)<br>-38.66 (1.39)* | *-32.36 (3.62)<br>-31.53 (3.25)* |
| *G4-I4<br>I4-G4 M2* | | *29.43 (2.79)<br>32.50 (2.68)* | *-38.59 (1.44)<br>-39.40 (1.16)* | *-9.16 (3.14)<br>-6.90 (2.92)* |
| *G4-G4<br>G4-G4 M2* | | *33.64 (2.59)<br>33.53 (2.51)* | *-43.16 (1.32)<br>-43.31 (1.09)* | *-9.52 (2.91)<br>-9.78 (2.74)* |
| *G4-A4<br>A4-G4 M2* | | *16.61 (9.96)<br>20.15 (5.07)* | *-38.34 (1.69)<br>-36.04 (1.56)* | *-21.73 (10.10)<br>-15.89 (5.30)* |
| *G4-G4 (5') (I)<br>U-G4-G4-U (5')* | | *2.22 (1.67)<br>15.13 (2.91)* | *-18.85 (1.57)<br>-32.87 (2.32)* | *-16.63 (2.29)<br>-17.74 (3.72)* |
| *G4-G4 (5') (G)<br>U-G4-G4-U (5')* | | *3.54 (1.98)<br>19.07 (3.72)* | *-18.55 (1.85)<br>-33.17 (2.76)* | *-15.01 (2.71)<br>-14.10 (4.63)* |
| *G4-G4 (5') (A)<br>U-G4-G4-U (5')* | | *6.64 (2.09)<br>19.41 (3.09)* | *-16.48 (2.62)<br>-30.39 (2.72)* | *-9.84 (3.35)<br>-10.98 (4.12)* |
| *Interaction energy between a tetrad and the proximal K+ cation* | | | | |
| *I-I-I-I & K+<br>K24<br>K23 M1* | | *<br>-106.88 (6.97)<br>-95.57 (5.77)* | *<br>3.07 (2.26)<br>1.21 (1.57)* | *<br>-103.81 (7.33)<br>-94.36 (5.97)* |
| *G-G-G-G & K+<br>K24<br>K23 M1* | | *<br>-108.45 (5.47)<br>-88.99 (7.66)* | *<br>4.94 (2.16)<br>0.04 (1.33)* | *<br>-103.51 (5.88)<br>-88.95 (7.77)* |
| *A-A-A-A and K+<br>K24<br>K23 M1* | | *<br>-7.62 (4.30)<br>-21.79 (2.45)* | *<br>0.002 (0.14)<br>-0.44 (0.08)* | *<br>-7.62 (4.32)<br>-22.23 (2.45)* |
| *I-I-I-I & K+* | | | | |

| | | | |
|---|---|---|---|
| *K24*<br>*K23 M2* | *-92.00 (5.75)*<br>*-109.51 (4.49)* | *1.09 (1.46)*<br>*4.82 (2.05)* | *-93.09 (5.93)*<br>*-104.69 (4.94)* |
| *G-G-G-G & K+*<br>*K24*<br>*K23 M2* | <br>*-94.20 (6.18)*<br>*-98.63 (6.12)* | <br>*1.37 (1.61)*<br>*1.17 (1.5)* | <br>*-92.83 (6.39)*<br>*-97.46 (6.30)* |
| *A-A-A-A and K+*<br>*K24*<br>*K23 M2* | <br>*-8.23 (3.99)*<br>*-47.64 (5.38)* | <br>*-0.04 (0.14)*<br>*-1.21 (1.17)* | <br>*-8.27 (3.99)*<br>*-48.85 (5.51)* |

**(C)** Interaction energies (kcal/mol) for hydrogen bonding, stacking and ion interactions for the different purine tetrads at position 2 of the dimeric quadruplexes (M1: monomer 1 and M2: monomer 2) for the third set of simulations. Average and standard deviation (within braces) values are reported.

| | | *ELEC* | *VDW* | *TOTAL* |
|---|---|---|---|---|
| *Hydrogen bonding energy of the tetrads* | | | | |
| *I-I-I-I M1* | *2-13*<br>*2-19*<br>*7-19*<br>*7-13* | *-3.43 (3.49)*<br>*-3.40 (3.73)*<br>*-2.93 (4.35)*<br>*-3.38 (3.68)* | *-1.19 (0.84)*<br>*-1.20 (0.82)*<br>*-1.32 (0.80)*<br>*-1.23 (0.81)* | *-4.62 (3.59)*<br>*-4.60 (3.82)*<br>*-4.25 (4.42)*<br>*-4.61 (3.77)* |
| *G-G-G-G M1* | *2-13*<br>*2-19*<br>*7-19*<br>*7-13* | *-12.46 (2.67)*<br>*-12.35 (2.56)*<br>*-12.61 (2.66)*<br>*-13.01 (2.51)* | *-0.52 (1.26)*<br>*-0.47 (1.27)*<br>*-0.31 (1.33)*<br>*-0.24 (1.32)* | *-12.98 (2.95)*<br>*-12.82 (2.86)*<br>*-12.92 (2.97)*<br>*-13.25 (2.84)* |
| *A-A-A-A M1* | *2-13*<br>*2-19*<br>*7-19*<br>*7-13* | *-3.25 (1.25)*<br>*-3.49 (1.15)*<br>*-2.83 (1.27)*<br>*-2.83 (1.25)* | *-0.77 (0.86)*<br>*-0.82 (0.89)*<br>*-0.94 (0.78)*<br>*-0.96 (0.77)* | *-4.02 (1.52)*<br>*-4.31 (1.45)*<br>*-3.77 (1.49)*<br>*-3.79 (1.47)* |
| *G-U5 M1 (I)* | *6-11* | *-6.31 (1.55)* | *-0.52 (0.81)* | *-6.83 (1.75)* |
| *G-U5 M1 (G)* | *6-11* | *-6.56 (1.29)* | *-0.06 (0.93)* | *-6.62 (1.59)* |
| *G-U5 M1 (A)* | *6-11* | *-1.29 (2.03)* | *-0.37 (0.35)* | *-1.66 (2.06)* |
| *I-I-I-I M2* | *25-42*<br>*42-31*<br>*31-37*<br>*25-37* | *-5.29(2.98)*<br>*-5.82 (3.05)*<br>*-6.20 (2.82)*<br>*-5.59 (2.79)* | *-1.16 (0.84)*<br>*-1.09 (0.87)*<br>*-1.01 (0.91)*<br>*-1.15 (0.90)* | *-6.45 (3.09)*<br>*-6.91 (3.17)*<br>*-7.21 (2.96)*<br>*-6.74 (2.93)* |
| *G-G-G-G M2* | *25-42*<br>*42-31*<br>*31-37*<br>*25-37* | *-12.70 (2.30)*<br>*-12.66 (2.30)*<br>*-12.41 (2.21)*<br>*-13.53 (2.16)* | *-0.33 (1.31)*<br>*-0.55 (1.23)*<br>*-0.54 (1.23)*<br>*-0.24 (1.30)* | *-13.03 (2.65)*<br>*-13.21 (2.61)*<br>*-12.95 (2.53)*<br>*-13.77 (2.52)* |
| *A-A-A-A M2* | *25-42*<br>*42-31* | *-3.01 (1.26)*<br>*-3.13 (1.20)* | *-0.86 (0.82)*<br>*-0.89 (0.83)* | *-3.87 (1.50)*<br>*-4.02 (1.46)* |

| | | | | |
|---|---|---|---|---|
| | *31-37*<br>*25-37* | *-2.85 (1.21)*<br>*-3.30 (1.16)* | *-0.87 (0.80)*<br>*-0.86 (0.86)* | *-3.72 (1.45)*<br>*-4.16 (1.44)* |
| *G-U5 M2 (I)* | | *-6.36 (1.33)* | *-0.31 (0.86)* | *-6.67 (1.58)* |
| *G-U5 M2 (G)* | | *-2.99 (3.38)* | *-1.05 (0.98)* | *-4.04 (3.52)* |
| *G-U5 M2 (A)* | | *-5.69 (2.17)* | *-0.30 (0.85)* | *-5.99 (2.33)* |
| *Stacking energy between tetrads* | | | | |
| *G4-I4*<br>*I4-G4 M1* | | *24.22 (5.18)*<br>*26.18 (6.52)* | *-39.07 (1.49)*<br>*-39.44 (1.16)* | *-14.85 (5.39)*<br>*-13.26 (6.62)* |
| *G4-G4*<br>*G4-G4 M1* | | *30.78 (3.17)*<br>*26.08 (5.49)* | *-43.37 (1.21)*<br>*-42.75 (1.23)* | *-12.59 (3.39)*<br>*-16.67 (5.63)* |
| *G4-A4*<br>*A4-G4 M1* | | *7.59 (2.29)*<br>*9.97 (2.29)* | *-37.36 (2.25)*<br>*-38.43 (1.32)* | *-29.77 (3.21)*<br>*-28.46 (2.64)* |
| *G4-I4*<br>*I4-G4 M2* | | *21.05 (3.95)*<br>*21.49 (4.89)* | *-39.64 (1.40)*<br>*-39.40 (1.15)* | *-18.59 (4.19)*<br>*-17.91 (5.02)* |
| *G4-G4*<br>*G4-G4 M2* | | *30.46 (2.72)*<br>*24.80 (3.70)* | *-43.16 (1.29)*<br>*-42.71 (1.18)* | *-12.70 (3.01)*<br>*-17.91 (3.88)* |
| *G4-A4*<br>*A4-G4 M2* | | *5.59 (3.43)*<br>*9.67 (2.68)* | *-38.36 (1.43)*<br>*-38.64 (1.47)* | *-32.77 (3.72)*<br>*-28.97 (3.06)* |
| *G4-G4 (5') (I)*<br>*U-G4-G4-U (5')* | | *9.09 (3.41)*<br>*20.12 (3.09)* | *-22.26 (3.55)*<br>*-36.87 (3.14)* | *-13.17 (4.92)*<br>*-16.75 (4.40)* |
| *G4-G4 (5') (G)*<br>*U-G4-G4-U (5')* | | *3.43 (1.78)*<br>*15.35 (4.43)* | *-18.17 (2.07)*<br>*-31.82 (3.11)* | *-14.74 (2.73)*<br>*-16.47 (5.41)* |
| *G4-G4 (5') (A)*<br>*U-G4-G4-U (5')* | | *8.98 (3.19)*<br>*21.67 (3.59)* | *-19.60 (2.20)*<br>*-33.56 (3.16)* | *-10.62 (3.88)*<br>*-11.89 (4.78)* |
| *I-I-I-I & K+*<br>*K24*<br>*K23 M1* | | <br>*-71.08 (38.84)*<br>*-80.92 (13.25)* | <br>*2.11 (2.93)*<br>*1.39 (1.68)* | <br>*-68.97 (38.95)*<br>*-79.53 (13.36)* |
| *G-G-G-G & K+*<br>*K24*<br>*K23 M1* | | <br>*-90.68 (10.87)*<br>*-24.02 (32.41)* | <br>*2.68 (2.43)*<br>*0.0012 (0.62)* | <br>*-88.00 (11.14)*<br>*-24.02 (32.42)* |
| *A-A-A-A and K+*<br>*K24*<br>*K23 M1* | | <br>*-7.78 (4.86)*<br>*-8.80 (4.63)* | <br>*0.001 (0.13)*<br>*-0.025 (0.09)* | <br>*-7.78 (4.86)*<br>*-8.82 (4.63)* |
| *I-I-I-I & K+*<br>*K24*<br>*K23 M2* | | <br>*-72.99 (10.23)*<br>*-43.85 (27.88)* | <br>*1.78 (1.79)*<br>*0.31 (0.96)* | <br>*-71.21 (10.38)*<br>*-43.54 (27.89)* |
| *G-G-G-G & K+* | | | | |

| | | | |
|---|---|---|---|
| *K24*<br>*K23 M2* | *-87.24 (6.57)*<br>*-45.39 (17.39)* | *1.92 (1.80)*<br>*-0.27 (0.76)* | *-85.32 (6.81)*<br>*-45.66 (17.41)* |
| *A-A-A-A and K+*<br>*K24*<br>*K23 M2* | <br>*-20.23 (2.06)*<br>*-8.98 (7.88)* | <br>*-0.30 (0.06)*<br>*0.005 (0.29)* | <br>*-20.53 (2.06)*<br>*-8.98 (7.88)* |

**(D)** Interaction energies (kcal/mol) for hydrogen bonding, stacking and ion interactions for the different purine tetrads at position 2 of the dimeric quadruplexes (M1: monomer 1 and M2: monomer 2) for the fourth set of simulations. Average and standard deviation (within braces) values are reported.

| | | *ELEC* | *VDW* | *TOTAL* |
|---|---|---|---|---|
| *Hydrogen bonding energy of the tetrads* | | | | |
| *I-I-I-I M1* | *2-13*<br>*2-19*<br>*7-19*<br>*7-13* | *-6.75 (1.83)*<br>*-6.93 (1.75)*<br>*-6.54 (1.79)*<br>*-6.77 (1.78)* | *-1.05 (0.89)*<br>*-1.04 (0.89)*<br>*-1.12 (0.88)*<br>*-1.02 (0.90)* | *-7.80 (2.03)*<br>*-7.97 (1.96)*<br>*-7.66 (1.99)*<br>*-7.79 (1.99)* |
| *G-G-G-G M1* | *2-13*<br>*2-19*<br>*7-19*<br>*7-13* | *-9.76 (2.48)*<br>*-9.33 (2.46)*<br>*-9.86 (2.55)*<br>*-10.30 (2.43)* | *-0.85 (1.18)*<br>*-0.81 (1.20)*<br>*-0.64 (1.25)*<br>*-0.46 (1.27)* | *-10.61 (2.75)*<br>*-10.14 (2.74)*<br>*-10.50 (2.84)*<br>*-10.76 (2.74)* |
| *A-A-A-A M1* | *2-13*<br>*2-19*<br>*7-19*<br>*7-13* | *-3.96 (1.39)*<br>*-3.99 (1.34)*<br>*-4.02 (1.37)*<br>*-4.11 (1.35)* | *-0.68 (1.02)*<br>*-0.71 (1.00)*<br>*-0.66 (1.03)*<br>*-0.69 (1.05)* | *-4.64 (1.72)*<br>*-4.70 (1.67)*<br>*-4.68 (1.71)*<br>*-4.80 (1.71)* |
| *G-U5 M1 (I)* | *6-11* | *-6.39 (1.72)* | *-0.09 (0.96)* | *-6.48 (1.97)* |
| *G-U5 M1 (G)* | *6-11* | *-6.71 (1.34)* | *-0.008 (0.97)* | *-6.72 (1.65)* |
| *G-U5 M1 (A)* | *6-11* | *-1.64 (2.97)* | *-0.53 (0.46)* | *-2.27 (3.00)* |
| *I-I-I-I M2* | *25-42*<br>*42-31*<br>*31-37*<br>*25-37* | *0.39 (1.88)*<br>*0.70 (1.98)*<br>*0.07 (1.74)*<br>*-0.54 (2.17)* | *-1.37 (0.74)*<br>*-1.56 (0.59)*<br>*-1.20 (0.84)*<br>*-1.27 (0.79)* | *-0.98 (2.02)*<br>*-0.86 (2.07)*<br>*-1.13 (1.93)*<br>*-0.73 (2.31)* |
| *G-G-G-G M2* | *25-42*<br>*42-31*<br>*31-37*<br>*25-37* | *-10.57 (3.14)*<br>*-10.58 (3.14)*<br>*-10.56 (2.93)*<br>*-11.66 (2.94)* | *-0.68 (1.27)*<br>*-0.78 (1.22)*<br>*-0.79 (1.19)*<br>*-0.56 (1.27)* | *-11.25 (3.39)*<br>*-11.36 (3.37)*<br>*-11.35 (3.16)*<br>*-12.22 (3.20)* |
| *A-A-A-A M2* | *25-42*<br>*42-31*<br>*31-37*<br>*25-37* | *-2.75 (1.52)*<br>*-3.18 (1.27)*<br>*-3.13 (1.29)*<br>*-3.42 (1.22)* | *-0.92 (0.79)*<br>*-0.91 (0.84)*<br>*-0.84 (0.85)*<br>*-0.82 (0.89)* | *-3.67 (1.71)*<br>*-4.09 (1.52)*<br>*-3.97 (1.54)*<br>*-4.24 (1.51)* |
| *G-U5 M2 (I)* | | *-5.70 (2.44)* | *-0.61 (1.03)* | *-6.31 (2.65)* |

| | | | | |
|---|---|---|---|---|
| *G-U5 M2 (I)* | | *-3.45 (2.69)* | *-1.08 (0.72)* | *-4.53 (2.78)* |
| *G-U5 M2 (I)* | | *-2.33 (2.98)* | *-0.37 (0.57)* | *-2.67 (3.03)* |
| *Stacking energy between tetrads* | | | | |
| *G4-I4*<br>*I4-G4 M1* | | *19.68 (2.56)*<br>*20.43 (2.79)* | *-40.12 (1.33)*<br>*-39.33 (1.14)* | *-20.44 (2.88)*<br>*-18.90 (3.01)* |
| *G4-G4*<br>*G4-G4 M1* | | *32.94 (2.86)*<br>*32.86 (3.56)* | *-43.49 (1.22)*<br>*-43.28 (1.13)* | *-10.55 (3.11)*<br>*-10.42 (3.74)* |
| *G4-A4*<br>*A4-G4 M1* | | *6.67 (3.07)*<br>*7.72 (2.92)* | *-39.14 (1.44)*<br>*-38.61 (1.41)* | *-32.47 (3.39)*<br>*-30.89 (3.24)* |
| *G4-I4*<br>*I4-G4 M2* | | *28.89 (3.19)*<br>*31.43 (3.46)* | *-38.60 (1.51)*<br>*-39.38 (1.21)* | *-9.71 (3.53)*<br>*-7.95 (3.66)* |
| *G4-G4*<br>*G4-G4 M2* | | *32.52 (3.27)*<br>*30.67 (6.69)* | *-42.94 (1.38)*<br>*-42.88 (1.24)* | *-10.42 (3.55)*<br>*-12.21 (6.80)* |
| *G4-A4*<br>*A4-G4 M2* | | *6.44 (3.30)*<br>*10.18 (2.91)* | *-37.04 (1.94)*<br>*-38.16 (1.44)* | *-30.6 (3.83)*<br>*-27.98 (3.25)* |
| *G4-G4 (5') (I)*<br>*U-G4-G4-U (5')* | | *4.77 (3.13)*<br>*16.80 (3.41)* | *-20.21 (3.31)*<br>*-33.84 (3.72)* | *-15.44 (4.56)*<br>*-17.04 (5.05)* |
| *G4-G4 (5') (G)*<br>*U-G4-G4-U (5')* | | *3.42 (2.67)*<br>*15.17 (3.37)* | *-18.81 (2.16)*<br>*-30.48 (3.31)* | *-15.39 (3.43)*<br>*-15.31 (4.72)* |
| *G4-G4 (5') (A)*<br>*U-G4-G4-U (5')* | | *15.68 (6.04)*<br>*23.88 (3.84)* | *-30.09 (8.76)*<br>*-38.46 (4.85)* | *-14.41 (10.64)*<br>*-14.58 (6.19)* |
| *Interaction energy between a tetrad and the proximal K+ cation* | | | | |
| *I-I-I-I & K+*<br>*K24*<br>*K23 M1* | | <br>*-33.50 (11.38)*<br>*-69.07 (6.62)* | <br>*-0.34 (0.76)*<br>*1.83 (1.78)* | <br>*-33.84 (11.40)*<br>*-67.24 (6.86)* |
| *G-G-G-G & K+*<br>*K24*<br>*K23 M1* | | <br>*-95.97 (15.86)*<br>*-91.21 (7.18)* | <br>*1.53 (1.97)*<br>*0.80 (1.60)* | <br>*-94.44 (15.98)*<br>*-90.41 (7.36)* |
| *A-A-A-A and K+*<br>*K24*<br>*K23 M1* | | <br>*-20.71 (2.76)*<br>*-21.55 (2.47)* | <br>*-0.43 (0.11)*<br>*-0.43 (0.08)* | <br>*-21.14 (2.76)*<br>*-21.98 (2.147)* |
| *I-I-I-I & K+*<br>*K24*<br>*K23 M2* | | <br>*-92.64 (7.02)*<br>*-106.94 (14.29)* | <br>*1.06 (1.52)*<br>*4.38 (2.27)* | <br>*-91.58 (7.18)*<br>*-102.56 (14.47)* |
| *G-G-G-G & K+*<br>*K24*<br>*K23 M2* | | <br>*-89.97 (7.32)*<br>*-76.75 (34.07)* | <br>*1.29 (1.71)*<br>*2.03 (2.85)* | <br>*-88.68 (7.52)*<br>*-74.72 (34.19)* |

| | | | |
|---|---|---|---|
| *A-A-A-A and K+*<br>*K24*<br>*K23 M2* | <br>*-11.45 (7.42)*<br>*-23.33 (4.31)* | <br>*-0.05 (0.24)*<br>*-0.39 (0.28)* | <br>*-11.50 (7.42)*<br>*-23.72 (4.32)* |

**(B)** Comparison of (total) Gibbs free energies (obtained from MM-GBSA analysis) of the dimeric quadruplexes and individual monomers for the four sets of simulations. Average and standard deviation (within braces) values are reported.

| *Free energy (Kcal/mol)* | *System* | *I4* | *G4* | *A4* |
|---|---|---|---|---|
| Simulation set-*1* | *Dimer* | *-7224.99 (32.82)* | *-8331.13 (29.91)* | *-7892.85 (31.25)* |
| | *Monomer 1* | *-3498.42 (20.55)* | *-4063.14 (20.44)* | *-3849.81 (20.81)* |
| | *Monomer 2* | *-3653.17 (21.39)* | *-4202.94 (21.19)* | *-3986.42 (21.16)* |
| Simulation set-*2* | *Dimer* | *-7227.11 (29.83)* | *-8327.72 (29.45)* | *-7888.81 (34.99)* |
| | *Monomer 1* | *-3503.92 (20.14)* | *-4057.96 (20.38)* | *-3846.28 (20.85)* |
| | *Monomer 2* | *-3651.89 (20.92)* | *-4204.58 (20.83)* | *-3991.18 (24.40)* |
| Simulation set-*3* | *Dimer* | *-7227.95 (31.19)* | *-8333.96 (29.94)* | *-7877.87 (31.25)* |
| | *Monomer 1* | *-3506.42 (21.04)* | *-4063.54 (20.28)* | *-3829.02 (20.92)* |
| | *Monomer 2* | *-3653.47 (21.25)* | *-4207.679 (20.97)* | *-3991.74 (21.55)* |
| Simulation set-*4* | *Dimer* | *-7229.75 (29.43)* | *-8343.6016 (28.64)* | *-7891.01 (30.41)* |
| | *Monomer 1* | *-3515.97 (20.35)* | *-4066.6316 (20.05)* | *-3832.75 (20.47)* |
| | *Monomer 2* | *-3650.01 (20.84)* | *-4210.0740 (20.55)* | *-3980.18 (21.74)* |

**Table S12. (A)** Frequencies (in %) of the Hoogsteen (HG) hydrogen bonds between the bases within different tetrads of the studied dimeric quadruplexes for the first set of simulation.

| H-bonds | MONOMER 1 | Quad-I4 | Quad-G4 | Quad-A4 |
|---|---|---|---|---|
| *1-11* | *(12)N2-H21- - -N7(1)*<br>*(12)N1-H1- - -O6(1)* | 39.79<br>76.32 | 62.89<br>68.55 | 59.94<br>58.63 |
| *2-12* | *(13)N2-H21- - -N7(2)*<br>*(13)N1-H1- - -O6(2)*<br>*(2)N6-H62 - - - N1(13)*<br>***(13)N1-H1- - -N7(2)*** | NA<br>- - -<br>NA<br>35.98 | 67.68<br>28.23<br>NA<br>NA | NA<br>NA<br>52.03<br>NA |

| | | | | |
|---|---|---|---|---|
| *3-13* | | | | |
| | *(14)N2-H21- - -N7(3)* | 69.72 | 53.76 | 50.96 |
| | *(14)N1-H1- - -O6(3)* | 45.94 | 62.27 | 62.60 |
| *4-14* | | | | |
| | *(15)N2-H21- - -N7(4)* | 60.68 | 65.07 | 70.79 |
| *5-15* | *(15)N1-H1- - -O6(4)* | 70.83 | 73.03 | 26.93 |
| | *(5)N3-H3- - -O4(16)* | 80.40 | 82.69 | 80.12 |
| | *(1)N2-H21- - -N7(18)* | 42.47 | 62.22 | 61.10 |
| *1-16* | *(1)N1-H1- - -O6(18)* | 68.83 | 58.22 | 68.28 |
| *2-17* | *(2)N2-H21- - -N7(19)* | NA | 73.65 | NA |
| | *(2)N1-H1- - -O6(19)* | - - - | 23.99 | NA |
| | *(19)N6-H62 - - - N1(2)* | NA | NA | 64.34 |
| | ***(2)N1-H1- - -N7(19)*** | 50.84 | NA | NA |
| *3-18* | | | | |
| | *(3)N2-H21- - -N7(20)* | 67.94 | 54.64 | 55.00 |
| | *(3)N1-H1- - -O6(20)* | 47.36 | 65.17 | 69.87 |
| *4-19* | | | | |
| | *(4)N2-H21- - -N7(21)* | 60.95 | 68.69 | 71.74 |
| *5-20* | *(4)N1-H1- - -O6(21)* | 73.91 | 73.00 | 30.54 |
| | *(22)N3-H3- - -O4(5)* | 79.50 | 82.05 | 79.48 |
| | *(18)N2-H21- - -N7(6)* | 61.93 | 57.71 | 56.68 |
| *6-16* | *(18)N1-H1- - -O6(6)* | 75.30 | 70.97 | 76.95 |
| *7-17* | *(19)N2-H21- - -N7(7)* | NA | 68.08 | NA |
| | *(19)N1-H1- - -O6(7)* | - - - | 36.33 | NA |
| | *(7)N6-H62 - - - N1(19)* | NA | NA | 61.00 |
| | ***(19)N1-H1- - -N7(7)*** | 13.50 | NA | NA |
| *8-18* | | | | |
| | *(20)N2-H21- - -N7(8)* | 64.58 | 60.27 | 59.65 |
| | *(20)N1-H1- - -O6(8)* | 49.35 | 62.93 | 64.32 |
| *9-19* | | | | |
| | *(21)N2-H21- - -N7(9)* | 69.27 | 68.93 | 74.21 |
| *10-20* | *(21)N1-H1- - -O6(9)* | 70.75 | 72.42 | 29.60 |
| | *(10)N3-H3- - -O4(22)* | 80.30 | 82.35 | 79.71 |
| *6-11* | *(6)N2-H21- - -N7(12)* | 77.14 | 63.34 | 74.87 |
| | *(6)N1-H1- - -O6(12)* | 69.85 | 73.15 | 55.82 |
| *7-12* | *(7)N2-H21- - -N7(13)* | NA | 78.87 | NA |
| | *(7)N1-H1- - -O6(13)* | - - - | 36.49 | NA |

| | | | | |
|---|---|---|---|---|
| | *(13)N6-H62 - - - N1(7)* | NA | NA | 59.17 |
| | ***(7)N1-H1- - -N7(13)*** | 60.10 | NA | NA |
| *8-13* | | | | |
| | *(8)N2-H21- - -N7(14)* | 71.19 | 63.41 | 59.65 |
| | *(8)N1-H1- - -O6(14)* | 49.35 | 61.35 | 63.65 |
| *9-14* | | | | |
| | *(9)N2-H21- - -N7(15)* | 65.12 | 67.94 | 74.68 |
| | *(9)N1-H1- - -O6(15)* | 67.51 | 73.22 | 28.25 |
| *10-15* | | | | |
| | *(16)N3-H3- - -O4(10)* | 81.07 | 82.66 | 78.87 |
| *11-6* | *(6)N2-H22- - -O2(11)* | 86.99 | 83.34 | 88.83 |
| | *(6)O2'-HO2'- - -O4(11)* | 0.04 | 2.23 | 0.80 |

| H-bonds | MONOMER 2 | Quad-<br>I4 | Quad-<br>G4 | Quad-<br>A4 |
|---|---|---|---|---|
| | *(36)N2-H21- - -N7(24)* | 46.54 | 67.49 | 76.19 |
| *24-36* | *(36)N1-H1- - -O6(24)* | 69.32 | 73.29 | 63.74 |
| | *(37)N2-H21- - -N7(25)* | NA | 72.33 | NA |
| *25-37* | *(37)N1-H1- - -O6(25)* | - - - | 25.30 | NA |
| | *(25)N6-H62 - - - N1(37)* | NA | NA | 70.80 |
| | ***(37)N1-H1- - -N7(25)*** | 20.78 | NA | NA |
| *26-38* | *(38)N2-H21- - -N7(26)* | 62.52 | 60.21 | 57.21 |
| | *(38)N1-H1- - -O6(26)* | 33.68 | 23.91 | 30.76 |
| *27-39* | | | | |
| | *(39)N2-H21- - -N7(27)* | 63.15 | 61.87 | 69.05 |
| | *(39)N1-H1- - -O6(27)* | 71.61 | 70.29 | 31.77 |
| *28-40* | | | | |
| | *(28)N3-H3- - -O4(40)* | 80.45 | 80.93 | 78.19 |
| | *(41)N2-H21- - -N7(30)* | 65.81 | 76.13 | 72.61 |
| *30-41* | *(41)N1-H1- - -O6(30)* | 74.22 | 50.52 | 59.41 |
| | *(42)N2-H21- - -N7(31)* | NA | 62.20 | NA |
| *31-42* | *(42)N1-H1- - -O6(31)* | - - - | 18.48 | NA |
| | *(31)N6-H62 - - - N1(42)* | NA | NA | 62.66 |
| | ***(42)N1-H1- - -N7(31)*** | 57.85 | NA | NA |
| *32-43* | *(43)N2-H21- - -N7(32)* | 71.53 | 58.08 | 61.44 |
| | *(43)N1-H1- - -O6(32)* | 35.21 | 22.76 | 36.82 |

| | | | | |
|---|---|---|---|---|
| *33-44* | *(44)N2-H21- - -N7(33)* | 62.95 | 66.25 | 69.17 |
| | *(44)N1-H1- - -O6(33)* | 68.60 | 71.28 | 35.18 |
| *34-45* | | | | |
| | *(34)N3-H3- - -O4(45)* | 80.60 | 81.04 | 78.90 |
| | *(30)N2-H21- - -N7(36)* | 68.58 | 54.03 | 48.92 |
| *36-30* | *(30)N1-H1- - -O6(36)* | 60.46 | 55.25 | 62.44 |
| | *(31)N2-H21- - -N7(37)* | NA | 58.57 | NA |
| *37-31* | *(31)N1-H1- - -O6(37)* | - - - | 26.84 | NA |
| | *(37)N6-H62 - - - N1(31)* | NA | NA | 55.98 |
| | ***(31)N1-H1- - -N7(37)*** | 54.14 | NA | NA |
| *38-32* | | | | |
| | *(32)N2-H21- - -N7(38)* | 70.00 | 62.17 | 58.00 |
| | *(32)N1-H1- - -O6(38)* | 31.74 | 26.43 | 39.61 |
| *39-33* | | | | |
| | *(33)N2-H21- - -N7(39)* | 62.40 | 67.16 | 70.41 |
| | *(33)N1-H1- - -O6(39)* | 70.66 | 69.18 | 32.54 |
| *40-34* | | | | |
| | *(40)N3-H3- - -O4(34)* | 80.02 | 79.92 | 77.76 |
| *24-41* | *(24)N2-H21- - -N7(41)* | 51.64 | 61.52 | 59.57 |
| | *(24)N1-H1- - -O6(41)* | 62.52 | 48.97 | 46.68 |
| | *(25)N2-H21- - -N7(42)* | NA | 64.87 | NA |
| *25-42* | *(25)N1-H1- - -O6(42)* | - - - | 14.17 | NA |
| | *(42)N6-H62 - - - N1(25)* | NA | NA | 70.22 |
| | ***(25)N1-H1- - -N7(42)*** | 23.37 | NA | NA |
| *26-43* | *(26)N2-H21- - -N7(43)* | 67.94 | 60.36 | 54.50 |
| | *(26)N1-H1- - -O6(43)* | 34.00 | 25.49 | 34.71 |
| *27-44* | *(27)N2-H21- - -N7(44)* | 66.21 | 63.80 | 68.09 |
| | *(27)N1-H1- - -O6(44)* | 70.14 | 73.19 | 33.04 |
| *28-45* | | | | |
| | *(45)N3-H3- - -O4(28)* | 80.95 | 80.33 | 78.17 |
| *29-41* | *(41)N2-H22- - -O2(29)* | - - - | 20.43 | 44.23 |
| | *(41)O2'-HO2'- - -O4(29)* | - - - | 0.98 | 2.27 |

**(B)** Frequencies (in %) of the Hoogsteen (HG) hydrogen bonds between the bases within different tetrads of the studied dimeric quadruplexes for the second set of simulation.

| H-bonds | MONOMER 1 | Quad-I4 | Quad-G4 | Quad-A4 |
|---|---|---|---|---|

| | | | | |
|---|---|---|---|---|
| *1-11* | *(12)N2-H21- - -N7(1)* | 48.19 | 57.58 | 56.22 |
| | *(12)N1-H1- - -O6(1)* | 67.88 | 59.75 | 63.79 |
| *2-12* | *(13)N2-H21- - -N7(2)* | NA | 61.15 | NA |
| | *(13)N1-H1- - -O6(2)* | - - - | 19.68 | NA |
| | *(2)N6-H62 - - - N1(13)* | NA | NA | 55.71 |
| | ***(13)N1-H1- - -N7(2)*** | 20.17 | NA | NA |
| *3-13* | *(14)N2-H21- - -N7(3)* | 66.34 | 61.56 | 53.16 |
| | *(14)N1-H1- - -O6(3)* | 55.37 | 21.58 | 60.52 |
| *4-14* | | | | |
| | *(15)N2-H21- - -N7(4)* | 66.54 | 59.55 | 72.07 |
| | *(15)N1-H1- - -O6(4)* | 73.70 | 72.30 | 28.77 |
| *5-15* | | | | |
| | *(5)N3-H3- - -O4(16)* | 82.83 | 80.40 | 80.08 |
| *1-16* | *(1)N2-H21- - -N7(18)* | 60.41 | 50.62 | 64.60 |
| | *(1)N1-H1- - -O6(18)* | 70.20 | 45.82 | 63.99 |
| *2-17* | *(2)N2-H21- - -N7(19)* | NA | 68.44 | NA |
| | *(2)N1-H1- - -O6(19)* | - - - | 18.76 | NA |
| | *(19)N6-H62 - - - N1(2)* | NA | NA | 61.56 |
| | ***(2)N1-H1- - -N7(19)*** | 71.41 | NA | NA |
| *3-18* | *(3)N2-H21- - -N7(20)* | 68.61 | 52.40 | 52.76 |
| | *(3)N1-H1- - -O6(20)* | 55.31 | 24.61 | 67.91 |
| *4-19* | | | | |
| | *(4)N2-H21- - -N7(21)* | 60.54 | 68.17 | 70.88 |
| | *(4)N1-H1- - -O6(21)* | 71.62 | 70.93 | 28.03 |
| *5-20* | | | | |
| | *(22)N3-H3- - -O4(5)* | 80.98 | 81.15 | 78.80 |
| *6-16* | *(18)N2-H21- - -N7(6)* | 47.96 | 51.66 | 62.59 |
| | *(18)N1-H1- - -O6(6)* | 75.70 | 69.62 | 56.02 |
| *7-17* | *(19)N2-H21- - -N7(7)* | NA | 68.16 | NA |
| | *(19)N1-H1- - -O6(7)* | - - - | 26.64 | NA |
| | *(7)N6-H62 - - - N1(19)* | NA | NA | 61.23 |
| | ***(19)N1-H1- - -N7(7)*** | 26.16 | NA | NA |
| *8-18* | *(20)N2-H21- - -N7(8)* | 56.29 | 61.65 | 58.11 |
| | *(20)N1-H1- - -O6(8)* | 61.75 | 25.87 | 66.67 |
| *9-19* | | | | |
| | *(21)N2-H21- - -N7(9)* | 62.26 | 68.91 | 70.67 |
| | *(21)N1-H1- - -O6(9)* | 73.61 | 69.25 | 31.19 |
| *10-20* | | | | |

| | | | | |
|---|---|---|---|---|
| | *(10)N3-H3- - -O4(22)* | 81.38 | 81.04 | 80.90 |
| *6-11* | *(6)N2-H21- - -N7(12)* | 78.53 | 62.51 | 49.27 |
| | *(6)N1-H1- - -O6(12)* | 64.29 | 68.93 | 66.90 |
| *7-12* | *(7)N2-H21- - -N7(13)* | NA | 76.14 | NA |
| | *(7)N1-H1- - -O6(13)* | - - - | 32.55 | NA |
| | *(13)N6-H62 - - - N1(7)* | NA | NA | 61.66 |
| | ***(7)N1-H1- - -N7(13)*** | 31.55 | NA | NA |
| *8-13* | *(8)N2-H21- - -N7(14)* | 70.38 | 66.24 | 64.70 |
| | *(8)N1-H1- - -O6(14)* | 56.71 | 24.46 | 69.11 |
| *9-14* | *(9)N2-H21- - -N7(15)* | 69.31 | 66.41 | 73.14 |
| | *(9)N1-H1- - -O6(15)* | 71.85 | 68.66 | 29.49 |
| *10-15* | | | | 78.70 |
| | *(16)N3-H3- - -O4(10)* | 82.90 | 80.17 | |
| *11-6* | *(6)N2-H22- - -O2(11)* | 88.56 | 84.85 | 58.28 |
| | *(6)O2'-HO2'- - -O4(11)* | 5.76 | 2.08 | 4.13 |

| H-bonds | MONOMER 2 | Quad-<br>I4 | Quad-<br>G4 | Quad-<br>A4 |
|---|---|---|---|---|
| | *(36)N2-H21- - -N7(24)* | 70.83 | 73.77 | 65.83 |
| *24-36* | *(36)N1-H1- - -O6(24)* | 68.14 | 77.12 | 58.05 |
| | *(37)N2-H21- - -N7(25)* | NA | 77.41 | NA |
| *25-37* | *(37)N1-H1- - -O6(25)* | - - - | 38.04 | NA |
| | *(25)N6-H62 - - - N1(37)* | NA | NA | 13.01 |
| | ***(37)N1-H1- - -N7(25)*** | 52.97 | NA | NA |
| | *(25)N6-H62 - - - N3(37)* | NA | NA | 11.73 |
| *26-38* | | | | |
| | *(38)N2-H21- - -N7(26)* | 67.01 | 56.43 | 65.45 |
| | *(38)N1-H1- - -O6(26)* | 26.40 | 70.70 | 54.04 |
| *27-39* | | | | |
| | *(39)N2-H21- - -N7(27)* | 64.82 | 65.92 | |
| *28-40* | *(39)N1-H1- - -O6(27)* | 70.79 | 71.63 | 66.28 |
| | | | | 33.28 |
| | *(28)N3-H3- - -O4(40)* | 81.77 | 83.52 | 78.17 |
| | *(41)N2-H21- - -N7(30)* | 64.44 | 77.87 | 66.80 |
| *30-41* | *(41)N1-H1- - -O6(30)* | 64.54 | 60.71 | 63.65 |

| | | | | |
|---|---|---|---|---|
| | *(42)N2-H21- - -N7(31)* | NA | 69.83 | NA |
| *31-42* | *(42)N1-H1- - -O6(31)* | - - - | 27.01 | NA |
| | *(31)N6-H62 - - - N1(42)* | NA | NA | 5.19 (H61- 4.82) |
| | ***(42)N1-H1- - -N7(31)*** | 27.09 | NA | NA |
| | *(31)N6-H62 - - - N3(42)* | NA | NA | 9.68 |
| *32-43* | | | | |
| | *(43)N2-H21- - -N7(32)* | 62.30 | 50.20 | 60.08 |
| | *(43)N1-H1- - -O6(32)* | 26.35 | 73.15 | 55.78 |
| *33-44* | | | | |
| | *(44)N2-H21- - -N7(33)* | 60.89 | 70.84 | 65.03 |
| | *(44)N1-H1- - -O6(33)* | 71.57 | 70.78 | 35.76 |
| *34-45* | | | | |
| | *(34)N3-H3- - -O4(45)* | 81.72 | 82.54 | 77.85 |
| | *(30)N2-H21- - -N7(36)* | 73.56 | 58.64 | 69.36 |
| *36-30* | *(30)N1-H1- - -O6(36)* | 61.90 | 68.90 | 50.10 |
| | *(31)N2-H21- - -N7(37)* | NA | 60.86 | NA |
| *37-31* | *(31)N1-H1- - -O6(37)* | - - - | 36.02 | NA |
| | *(37)N6-H62 - - - N1(31)* | NA | NA | 7.13 |
| | ***(31)N1-H1- - -N7(37)*** | 20.50 | NA | NA |
| | *(37)N6-H62 - - - N3(31)* | NA | NA | 11.21 |
| *38-32* | | | | |
| | *(32)N2-H21- - -N7(38)* | 67.72 | 59.68 | 66.80 |
| | *(32)N1-H1- - -O6(38)* | 31.56 | 72.25 | 56.14 |
| *39-33* | | | | |
| | *(33)N2-H21- - -N7(39)* | 59.60 | 69.27 | 67.39 |
| *40-34* | *(33)N1-H1- - -O6(39)* | 71.83 | 75.47 | 31.89 |
| | *(40)N3-H3- - -O4(34)* | 81.61 | 83.94 | 77.81 |
| *24-41* | *(24)N2-H21- - -N7(41)* | 70.89 | 63.56 | 59.68 |
| | *(24)N1-H1- - -O6(41)* | 50.47 | 54.24 | 61.54 |
| | *(25)N2-H21- - -N7(42)* | NA | 74.81 | NA |
| *25-42* | *(25)N1-H1- - -O6(42)* | - - - | 22.94 | NA |
| | *(42)N6-H62 - - - N1(25)* | NA | NA | 11.95 (H61- 1.62) |
| | ***(25)N1-H1- - -N7(42)*** | 20.83 | NA | NA |
| | *(42)N6-H61 - - - N3(25)* | NA | NA | 18.16 |
| *26-43* | | | | |
| | *(26)N2-H21- - -N7(43)* | 67.59 | 55.55 | 68.58 |
| | *(26)N1-H1- - -O6(43)* | 29.00 | 70.73 | 53.66 |
| *27-44* | | | | |
| | *(27)N2-H21- - -N7(44)* | 59.42 | 68.66 | 65.64 |
| | *(27)N1-H1- - -O6(44)* | 72.36 | 77.09 | 32.92 |
| *28-45* | | | | |

| | | | | |
|---|---|---|---|---|
| | *(45)N3-H3- - -O4(28)* | 80.59 | 84.09 | 77.12 |
| *29-41* | *(41)N2-H22- - -O2(29)* | - - - | 0.11 | 3.06 |
| | *(41)O2'-HO2'- - -O4(29)* | - - - | - - - | 0.38 |

**(C)** Frequencies (in %) of the Hoogsteen (HG) hydrogen bonds between the bases within different tetrads of the studied dimeric quadruplexes for the third set of simulation.

| H-bonds | MONOMER 1 | Quad-I4 | Quad-G4 | Quad-A4 |
|---|---|---|---|---|
| | *(12)N2-H21- - -N7(1)* | 66.83 | 66.34 | 23.78 |
| *1-11* | *(12)N1-H1- - -O6(1)* | 60.58 | 63.32 | 11.80 |
| *2-12* | *(13)N2-H21- - -N7(2)* | NA | 67.39 | NA |
| | *(13)N1-H1- - -O6(2)* | 39.58 | 61.35 | NA |
| | *(2)N6-H62 - - - N1(13)* | NA | NA | 53.65 |
| | ***(13)N1-H1- - -N7(2)*** | 19.25 | NA | NA |
| *3-13* | | | | |
| | *(14)N2-H21- - -N7(3)* | 57.23 | 52.58 | 47.23 |
| | *(14)N1-H1- - -O6(3)* | 51.10 | 61.16 | 72.48 |
| *4-14* | | | | |
| | *(15)N2-H21- - -N7(4)* | 68.07 | 67.74 | 72.58 |
| *5-15* | *(15)N1-H1- - -O6(4)* | 50.36 | 36.93 | 30.93 |
| | *(5)N3-H3- - -O4(16)* | 81.42 | 80.17 | 81.43 |
| | *(1)N2-H21- - -N7(18)* | 56.57 | 61.36 | 28.08 |
| *1-16* | *(1)N1-H1- - -O6(18)* | 63.71 | 60.84 | 86.82 |
| *2-17* | *(2)N2-H21- - -N7(19)* | NA | 71.34 | NA |
| | *(2)N1-H1- - -O6(19)* | - - - | 61.92 | NA |
| | *(19)N6-H62 - - - N1(2)* | NA | NA | 56.46 |
| | ***(2)N1-H1- - -N7(19)*** | 19.89 | NA | NA |
| *3-18* | | | | |
| | *(3)N2-H21- - -N7(20)* | 59.19 | 51.03 | 66.17 |
| | *(3)N1-H1- - -O6(20)* | 55.12 | 68.85 | 71.93 |
| *4-19* | | | | |
| | *(4)N2-H21- - -N7(21)* | 67.67 | 72.36 | 73.01 |
| *5-20* | *(4)N1-H1- - -O6(21)* | 50.61 | 37.24 | 27.63 |
| | *(22)N3-H3- - -O4(5)* | 80.50 | 80.70 | 78.74 |
| | *(18)N2-H21- - -N7(6)* | 57.79 | 60.08 | 24.92 |
| *6-16* | *(18)N1-H1- - -O6(6)* | 67.83 | 71.28 | 77.74 |

| | | | | |
|---|---|---|---|---|
| *7-17* | *(19)N2-H21- - -N7(7)* | NA | 69.22 | NA |
| | *(19)N1-H1- - -O6(7)* | 40.41 | 71.28 | NA |
| | *(7)N6-H62 - - - N1(19)* | NA | NA | 41.69 |
| | ***(19)N1-H1- - -N7(7)*** | 6.31 | NA | NA |
| *8-18* | | | | |
| | *(20)N2-H21- - -N7(8)* | 61.19 | 56.21 | 52.95 |
| | *(20)N1-H1- - -O6(8)* | 51.84 | 64.48 | 74.11 |
| *9-19* | | | | |
| | *(21)N2-H21- - -N7(9)* | 67.99 | 74.94 | 68.78 |
| *10-20* | *(21)N1-H1- - -O6(9)* | 51.21 | 35.69 | 33.23 |
| | *(10)N3-H3- - -O4(22)* | 81.61 | 81.03 | |
| *6-11* | *(6)N2-H21- - -N7(12)* | 77.56 | 74.45 | 24.33 |
| | *(6)N1-H1- - -O6(12)* | 77.29 | 64.22 | 66.24 |
| *7-12* | *(7)N2-H21- - -N7(13)* | NA | 71.93 | NA |
| | *(7)N1-H1- - -O6(13)* | 39.02 | 70.91 | NA |
| | *(13)N6-H62 - - - N1(7)* | NA | NA | 40.84 |
| | ***(7)N1-H1- - -N7(13)*** | 15.90 | NA | NA |
| *8-13* | | | | |
| | *(8)N2-H21- - -N7(14)* | 62.28 | 56.73 | 56.06 |
| | *(8)N1-H1- - -O6(14)* | 53.55 | 63.37 | 77.82 |
| *9-14* | | | | |
| | *(9)N2-H21- - -N7(15)* | 67.75 | 73.58 | 74.37 |
| | *(9)N1-H1- - -O6(15)* | 50.44 | 35.43 | 29.72 |
| *10-15* | | | | |
| | *(16)N3-H3- - -O4(10)* | 80.21 | 79.40 | 79.25 |
| *11-6* | *(6)N2-H22- - -O2(11)* | 69.05 | 84.13 | 8.91 |
| | *(6)O2'-HO2'- - -O4(11)* | 14.34 | 1.49 | 1.72 |

| H-bonds | MONOMER 2 | Quad-<br>I4 | Quad-<br>G4 | Quad-<br>A4 |
|---|---|---|---|---|
| | *(36)N2-H21- - -N7(24)* | 67.31 | 77.82 | 65.32 |
| *24-36* | *(36)N1-H1- - -O6(24)* | 66.89 | 80.06 | 58.34 |
| | *(37)N2-H21- - -N7(25)* | NA | 71.61 | NA |
| *25-37* | *(37)N1-H1- - -O6(25)* | 62.57 | 74.02 | NA |
| | *(25)N6-H62 - - - N1(37)* | NA | NA | 53.09 |
| | ***(37)N1-H1- - -N7(25)*** | 5.06 | NA | NA |

| | | | | |
|---|---|---|---|---|
| 26-38 | (38)N2-H21- - -N7(26) | 52.22 | 51.51 | 58.16 |
| | (38)N1-H1- - -O6(26) | 66.90 | 71.37 | 68.58 |
| 27-39 | (39)N2-H21- - -N7(27) | 69.19 | 71.51 | 72.02 |
| | (39)N1-H1- - -O6(27) | 33.56 | 31.79 | 29.80 |
| 28-40 | (28)N3-H3- - -O4(40) | 80.63 | 81.68 | 80.11 |
| 30-41 | (41)N2-H21- - -N7(30) | 56.11 | 79.48 | 69.84 |
| | (41)N1-H1- - -O6(30) | 73.59 | 69.38 | 64.75 |
| 31-42 | (42)N2-H21- - -N7(31) | NA | 68.84 | NA |
| | (42)N1-H1- - -O6(31) | 66.80 | 61.19 | NA |
| | (31)N6-H62 - - - N1(42) | NA | NA | 49.36 |
| | **(42)N1-H1- - -N7(31)** | 4.74 | NA | NA |
| 32-43 | (43)N2-H21- - -N7(32) | 55.73 | 53.51 | 65.53 |
| | (43)N1-H1- - -O6(32) | 67.81 | 76.23 | 71.76 |
| 33-44 | (44)N2-H21- - -N7(33) | 71.57 | 71.45 | 72.64 |
| | (44)N1-H1- - -O6(33) | 36.95 | 33.69 | 30.39 |
| 34-45 | (34)N3-H3- - -O4(45) | 81.65 | 81.62 | 79.91 |
| 36-30 | (30)N2-H21- - -N7(36) | 37.27 | 65.83 | 50.59 |
| | (30)N1-H1- - -O6(36) | 73.13 | 70.81 | 57.44 |
| 37-31 | (31)N2-H21- - -N7(37) | NA | 56.06 | NA |
| | (31)N1-H1- - -O6(37) | 69.94 | 76.04 | NA |
| | (37)N6-H62 - - - N1(31) | NA | NA | 44.89 |
| | **(31)N1-H1- - -N7(37)** | 6.36 | NA | NA |
| 38-32 | (32)N2-H21- - -N7(38) | 58.79 | 56.23 | 52.82 |
| | (32)N1-H1- - -O6(38) | 69.41 | 77.15 | 72.07 |
| 39-33 | (33)N2-H21- - -N7(39) | 72.36 | 75.77 | 72.82 |
| | (33)N1-H1- - -O6(39) | 35.24 | 33.49 | 31.69 |
| 40-34 | (40)N3-H3- - -O4(34) | 80.78 | 80.11 | 79.54 |
| 24-41 | (24)N2-H21- - -N7(41) | 58.70 | 69.70 | 66.33 |
| | (24)N1-H1- - -O6(41) | 62.51 | | 42.66 |
| 25-42 | (25)N2-H21- - -N7(42) | NA | 74.04 | NA |
| | (25)N1-H1- - -O6(42) | 61.26 | 65.32 | NA |
| | (42)N6-H62 - - - N1(25) | NA | NA | 47.43 |

| | | | | |
|---|---|---|---|---|
| | ***(25)N1-H1- - -N7(42)*** | 4.74 | NA | NA |
| *26-43* | *(26)N2-H21- - -N7(43)* | 54.75 | 48.19 | 49.11 |
| | *(26)N1-H1- - -O6(43)* | 69.29 | 74.66 | 75.36 |
| *27-44* | *(27)N2-H21- - -N7(44)* | 67.53 | 71.73 | 71.01 |
| | *(27)N1-H1- - -O6(44)* | 36.70 | 34.04 | 32.37 |
| *28-45* | | | | |
| | *(45)N3-H3- - -O4(28)* | 79.84 | 80.24 | 79.97 |
| *29-41* | *(41)N2-H22- - -O2(29)* | - - - | 14.08 | 68.93 |
| | *(41)O2'-HO2'- - -O4(29)* | - - - | 1.13 | 9.97 |

**(D)** Frequencies (in %) of the Hoogsteen (HG) hydrogen bonds between the bases within different tetrads of the studied dimeric quadruplexes for the fourth set of simulation.

| H-bonds | MONOMER 1 | Quad-I4 | Quad-G4 | Quad-A4 |
|---|---|---|---|---|
| | *(12)N2-H21- - -N7(1)* | 61.40 | 66.36 | 66.64 |
| *1-11* | *(12)N1-H1- - -O6(1)* | 65.73 | 67.42 | 65.18 |
| *2-12* | *(13)N2-H21- - -N7(2)* | NA | 68.95 | NA |
| | *(13)N1-H1- - -O6(2)* | 77.17 | 31.65 | NA |
| | *(2)N6-H62 - - - N1(13)* | NA | NA | 55.14 |
| | ***(13)N1-H1- - -N7(2)*** | 0.39 | NA | NA |
| *3-13* | | | | |
| | *(14)N2-H21- - -N7(3)* | 46.01 | 53.65 | 54.87 |
| | *(14)N1-H1- - -O6(3)* | 72.28 | 66.45 | 62.79 |
| *4-14* | | | | |
| | *(15)N2-H21- - -N7(4)* | 70.62 | 66.76 | 72.51 |
| *5-15* | *(15)N1-H1- - -O6(4)* | 29.06 | 71.01 | 28.14 |
| | *(5)N3-H3- - -O4(16)* | 81.24 | 82.70 | 79.32 |
| | *(1)N2-H21- - -N7(18)* | 57.83 | 64.70 | 60.92 |
| *1-16* | *(1)N1-H1- - -O6(18)* | 66.24 | 53.37 | 70.12 |
| *2-17* | *(2)N2-H21- - -N7(19)* | NA | 71.87 | NA |
| | *(2)N1-H1- - -O6(19)* | 78.54 | 28.98 | NA |
| | *(19)N6-H62 - - - N1(2)* | NA | NA | 58.28 |
| | ***(2)N1-H1- - -N7(19)*** | 0.54 | NA | NA |
| *3-18* | | | | |
| | *(3)N2-H21- - -N7(20)* | 55.73 | 53.35 | 58.23 |
| | *(3)N1-H1- - -O6(20)* | 74.38 | 68.90 | 66.62 |
| *4-19* | | | | |

| | | | | |
|---|---|---|---|---|
| | *(4)N2-H21- - -N7(21)* | 71.39 | 70.82 | 72.08 |
| *5-20* | *(4)N1-H1- - -O6(21)* | 29.81 | 70.54 | 26.48 |
| | *(22)N3-H3- - -O4(5)* | 80.16 | 82.46 | 79.30 |
| | *(18)N2-H21- - -N7(6)* | 65.05 | 50.59 | 61.02 |
| *6-16* | *(18)N1-H1- - -O6(6)* | 70.33 | 73.32 | 65.27 |
| *7-17* | *(19)N2-H21- - -N7(7)* | NA | 71.85 | NA |
| | *(19)N1-H1- - -O6(7)* | 75.74 | 41.86 | NA |
| | *(7)N6-H62 - - - N1(19)* | NA | NA | 57.40 |
| | ***(19)N1-H1- - -N7(7)*** | 0.59 | NA | NA |
| *8-18* | | | | |
| | *(20)N2-H21- - -N7(8)* | 52.08 | 59.35 | 51.46 |
| | *(20)N1-H1- - -O6(8)* | 72.46 | 67.37 | 68.61 |
| *9-19* | | | | |
| | *(21)N2-H21- - -N7(9)* | 73.41 | 70.70 | 71.93 |
| *10-20* | *(21)N1-H1- - -O6(9)* | 30.41 | 70.06 | 29.60 |
| | *(10)N3-H3- - -O4(22)* | 81.04 | 82.91 | 80.44 |
| *6-11* | *(6)N2-H21- - -N7(12)* | 72.97 | 63.45 | 53.07 |
| | *(6)N1-H1- - -O6(12)* | 62.39 | 76.89 | 68.14 |
| *7-12* | *(7)N2-H21- - -N7(13)* | NA | 79.77 | NA |
| | *(7)N1-H1- - -O6(13)* | 78.06 | 40.26 | NA |
| | *(13)N6-H62 - - - N1(7)* | NA | NA | 60.74 |
| | ***(7)N1-H1- - -N7(13)*** | 0.68 | NA | NA |
| *8-13* | | | | |
| | *(8)N2-H21- - -N7(14)* | 55.31 | 62.52 | 67.58 |
| | *(8)N1-H1- - -O6(14)* | 73.92 | 65.22 | 67.41 |
| *9-14* | | | | |
| | *(9)N2-H21- - -N7(15)* | 74.84 | 69.23 | 73.05 |
| | *(9)N1-H1- - -O6(15)* | 27.88 | 69.98 | 26.58 |
| *10-15* | | | | |
| | *(16)N3-H3- - -O4(10)* | 79.98 | 82.90 | 78.88 |
| *11-6* | *(6)N2-H22- - -O2(11)* | 78.40 | 85.78 | 15.30 |
| | *(6)O2'-HO2'- - -O4(11)* | 2.96 | 1.97 | 4.71 |

| H-bonds | MONOMER 2 | Quad-I4 | Quad-G4 | Quad-A4 |
|---|---|---|---|---|
| | *(36)N2-H21- - -N7(24)* | 59.62 | 73.18 | 16.89 |
| *24-36* | *(36)N1-H1- - -O6(24)* | 78.44 | 76.61 | 43.36 |

| | | | | |
|---|---|---|---|---|
| *25-37* | *(37)N2-H21- - -N7(25)* | NA | 70.96 | NA |
| | *(37)N1-H1- - -O6(25)* | 2.72 | 51.28 | NA |
| | *(25)N6-H62 - - - N1(37)* | NA | NA | 54.43 |
| | ***(37)N1-H1- - -N7(25)*** | 41.34 | NA | NA |
| *26-38* | *(38)N2-H21- - -N7(26)* | 63.82 | 50.40 | 57.20 |
| | *(38)N1-H1- - -O6(26)* | 32.17 | 49.45 | 69.16 |
| *27-39* | *(39)N2-H21- - -N7(27)* | 63.86 | 67.55 | 71.75 |
| | *(39)N1-H1- - -O6(27)* | 70.18 | 51.49 | 27.09 |
| *28-40* | *(28)N3-H3- - -O4(40)* | 81.48 | 81.67 | 80.01 |
| *30-41* | *(41)N2-H21- - -N7(30)* | 70.74 | 75.94 | 56.87 |
| | *(41)N1-H1- - -O6(30)* | 72.10 | 59.83 | 50.43 |
| *31-42* | *(42)N2-H21- - -N7(31)* | NA | 66.35 | NA |
| | *(42)N1-H1- - -O6(31)* | 1.87 | 41.85 | NA |
| | *(31)N6-H62 - - - N1(42)* | NA | NA | 47.34 |
| | ***(42)N1-H1- - -N7(31)*** | 17.92 | NA | NA |
| *32-43* | *(43)N2-H21- - -N7(32)* | 64.69 | 56.84 | 58.48 |
| | *(43)N1-H1- - -O6(32)* | 32.81 | 50.53 | 72.91 |
| *33-44* | *(44)N2-H21- - -N7(33)* | 65.84 | 70.05 | 73.76 |
| | *(44)N1-H1- - -O6(33)* | 71.18 | 52.65 | 28.44 |
| *34-45* | *(34)N3-H3- - -O4(45)* | 81.04 | 81.25 | 80.50 |
| *36-30* | *(30)N2-H21- - -N7(36)* | 65.31 | 62.15 | 28.51 |
| | *(30)N1-H1- - -O6(36)* | 68.45 | 59.04 | 70.48 |
| *37-31* | *(31)N2-H21- - -N7(37)* | NA | 57.19 | NA |
| | *(31)N1-H1- - -O6(37)* | 2.40 | 52.46 | NA |
| | *(37)N6-H62 - - - N1(31)* | NA | NA | 48.40 |
| | ***(31)N1-H1- - -N7(37)*** | 33.90 | NA | NA |
| *38-32* | *(32)N2-H21- - -N7(38)* | 68.54 | 61.70 | 55.89 |
| | *(32)N1-H1- - -O6(38)* | 36.99 | 51.53 | 76.05 |
| *39-33* | *(33)N2-H21- - -N7(39)* | 61.61 | 70.50 | 74.17 |
| | *(33)N1-H1- - -O6(39)* | 70.44 | 51.72 | 27.86 |
| *40-34* | *(40)N3-H3- - -O4(34)* | 80.49 | 80.07 | 79.12 |

| | | | | |
|---|---|---|---|---|
| *24-41* | *(24)N2-H21- - -N7(41)*<br>*(24)N1-H1- - -O6(41)* | 58.39<br>59.64 | 62.08<br>57.62 | 15.52<br>39.81 |
| *25-42* | *(25)N2-H21- - -N7(42)*<br>*(25)N1-H1- - -O6(42)*<br>*(42)N6-H62 - - - N1(25)*<br>***(25)N1-H1- - -N7(42)*** | NA<br>2.62<br>NA<br>31.02 | 71.26<br>39.21<br>NA<br>NA | NA<br>NA<br>38.43<br>NA |
| *26-43* | *(26)N2-H21- - -N7(43)*<br>*(26)N1-H1- - -O6(43)* | 65.77<br>37.28 | 55.08<br>52.76 | 48.43<br>74.70 |
| *27-44* | *(27)N2-H21- - -N7(44)*<br>*(27)N1-H1- - -O6(44)* | 60.49<br>72.55 | 68.16<br>54.12 | 73.05<br>27.57 |
| *28-45* | *(45)N3-H3- - -O4(28)* | 80.81 | 80.68 | 78.96 |
| *29-41* | *(41)N2-H22- - -O2(29)*<br>*(41)O2′-HO2′- - -O4(29)* | 32.65<br>2.08 | 4.31<br>0.52 | 47.68<br>2.59 |

**Table S13.** Frequencies (in %) of the signature hydrogen bond between G-OP2 and 3′U-O2′ observed for the quadruplexes with reversed-3′U-tetrads for the studied dimeric quadruplexes for the four sets of simulations.

| Set-1 | H-bonds | Quad-I4 | Quad-G4 | Quad-A4 |
|---|---|---|---|---|
| | Monomer 1 | | | |
| *G4-U5*<br>*G9-U10*<br>*G15-U16*<br>*G21-U22* | (4)OP2- - - HO2′-O2′(5)<br>(9)OP2- - - HO2′-O2′(10)<br>(15)OP2- - - HO2′-O2′(16)<br>(21)OP2- - - HO2′-O2′(22) | 96.87<br>98.06<br>97.01<br>94.45 | 98.12<br>97.75<br>97.62<br>98.00 | 98.86<br>99.13<br>99.01<br>98.59 |
| | Monomer 2 | | | |
| *G27-U28*<br>*G33-U34*<br>*G39-U40*<br>*G44-U45* | (27)OP2- - - HO2′-O2′(28)<br>(33)OP2- - - HO2′-O2′(34)<br>(39)OP2- - - HO2′-O2′(40)<br>(44)OP2- - - HO2′-O2′(45) | 96.15<br>95.17<br>96.30<br>97.86 | 97.89<br>97.94<br>97.19<br>98.13 | 98.79<br>98.83<br>98.91<br>98.55 |

| Set-2 | H-bonds | Quad-I4 | Quad-G4 | Quad-A4 |
|---|---|---|---|---|
| | Monomer 1 | | | |
| *G4-U5*<br>*G9-U10*<br>*G15-U16*<br>*G21-U22* | (4)OP2- - - HO2′-O2′(5)<br>(9)OP2- - - HO2′-O2′(10)<br>(15)OP2- - - HO2′-O2′(16) | 97.74<br>95.70<br>96.95<br>90.41 | 97.66<br>98.08<br>98.12<br>98.48 | 98.85<br>99.01<br>98.75<br>98.81 |

| | | | | |
|---|---|---|---|---|
| | (21)OP2- - - HO2′-O2′(22) | | | |
| | Monomer 2 | | | |
| *G27-U28*<br>*G33-U34*<br>*G39-U40*<br>*G44-U45* | (27)OP2- - - HO2′-O2′(28)<br>(33)OP2- - - HO2′-O2′(34)<br>(39)OP2- - - HO2′-O2′(40)<br>(44)OP2- - - HO2′-O2′(45) | 97.30<br>97.55<br>97.64<br>97.20 | 97.79<br>98.49<br>97.11<br>98.09 | 98.30<br>98.27<br>98.60<br>97.96 |

| Set-3 | H-bonds | Quad-I4 | Quad-G4 | Quad-A4 |
|---|---|---|---|---|
| | Monomer 1 | | | |
| *G4-U5*<br>*G9-U10*<br>*G15-U16*<br>*G21-U22* | (4)OP2- - - HO2′-O2′(5)<br>(9)OP2- - - HO2′-O2′(10)<br>(15)OP2- - - HO2′-O2′(16)<br>(21)OP2- - - HO2′-O2′(22) | 98.24<br>98.63<br>98.15<br>98.28 | 98.40<br>98.90<br>98.42<br>98.63 | 99.03<br>98.82<br>98.67<br>98.85 |
| | Monomer 2 | | | |
| *G27-U28*<br>*G33-U34*<br>*G39-U40*<br>*G44-U45* | (27)OP2- - - HO2′-O2′(28)<br>(33)OP2- - - HO2′-O2′(34)<br>(39)OP2- - - HO2′-O2′(40)<br>(44)OP2- - - HO2′-O2′(45) | 98.32<br>98.79<br>98.48<br>98.36 | 98.72<br>98.84<br>98.77<br>98.55 | 98.75<br>98.94<br>98.79<br>98.83 |

| Set-4 | H-bonds | Quad-I4 | Quad-G4 | Quad-A4 |
|---|---|---|---|---|
| | Monomer 1 | | | |
| *G4-U5*<br>*G9-U10*<br>*G15-U16*<br>*G21-U22* | (4)OP2- - - HO2′-O2′(5)<br>(9)OP2- - - HO2′-O2′(10)<br>(15)OP2- - - HO2′-O2′(16)<br>(21)OP2- - - HO2′-O2′(22) | 98.83<br>98.96<br>98.70<br>98.89 | 98.26<br>98.00<br>97.90<br>98.15 | 98.84<br>98.82<br>98.92<br>98.73 |
| | Monomer 2 | | | |
| *G27-U28*<br>*G33-U34*<br>*G39-U40*<br>*G44-U45* | (27)OP2- - - HO2′-O2′(28)<br>(33)OP2- - - HO2′-O2′(34)<br>(39)OP2- - - HO2′-O2′(40)<br>(44)OP2- - - HO2′-O2′(45) | 97.52<br>97.74<br>96.16<br>97.16 | 98.28<br>98.42<br>97.99<br>98.35 | 98.94<br>98.81<br>98.82<br>98.94 |

**Table S14. (A)** Frequencies (in %) of the water bridging interactions observed for the I4/G4/A4 tetrads within the studied dimeric quadruplexes for the four sets of simulations.

| **I4-monomer1** | N3(2)-HO2′(2) | N3(7)-HO2′(7) | N3(12)-HO2′(13) | N3(17)-HO2′(19) |
|---|---|---|---|---|
| Simulation set-1 | **2.09** | **4.54** | **2.11** | **2.96** |

| | | | | |
|---|---|---|---|---|
| Simulation set-2<br>Simulation set-3<br>Simulation set-4 | **2.98**<br>**9.49**<br>**4.25** | **7.19**<br>**6.69**<br>**8.14** | **7.80**<br>**7.97**<br>**7.83** | **0.34**<br>**6.00**<br>**7.64** |
| | N7(2)-OP2(2) | N7(7)-OP2(7) | N7(13)-OP2(13) | N7(19)-OP2(19) |
| Simulation set-1<br>Simulation set-2<br>Simulation set-3<br>Simulation set-4 | **- - -**<br>**- - -**<br>**0.49**<br>**2.30** | **- - -**<br>**- - -**<br>**1.48**<br>**2.23** | **- - -**<br>**- - -**<br>**1.53**<br>**3.81** | **- - -**<br>**- - -**<br>**1.49**<br>**2.12** |
| **G4-monomer1** | N3(2)-HO2′(2) | N3(7)-HO2′(7) | N3(12)-HO2′(13) | N3(17)-HO2′(19) |
| Simulation set-1<br>Simulation set-2<br>Simulation set-3<br>Simulation set-4 | **5.71**<br>**3.63**<br>**5.03**<br>**4.45** | **17.58**<br>**21.35**<br>**16.11**<br>**16.32** | **8.31**<br>**8.81**<br>**7.58**<br>**7.89** | **24.26**<br>**26.15**<br>**26.84**<br>**33.02** |
| | H22(13)-OP2(2) | H22(7)-OP2(13) | H22(19)-OP2(7) | H22(2)-OP2(19) |
| Simulation set-1<br>Simulation set-2<br>Simulation set-3<br>Simulation set-4 | **41.42**<br>**48.37**<br>**41.85**<br>**39.47** | **42.86**<br>**43.33**<br>**36.48**<br>**39.98** | **50.16**<br>**56.96**<br>**50.22**<br>**48.39** | **38.99**<br>**63.04**<br>**49.90**<br>**46.81** |
| **A4-monomer1** | N3(2)-HO2′(2) | N3(7)-HO2′(7) | N3(12)-HO2′(13) | N3(19)-HO2′(19) |
| Simulation set-1<br>Simulation set-2<br>Simulation set-3<br>Simulation set-4 | **3.63**<br>**1.81**<br>**0.79**<br>**6.7** | **4.46**<br>**5.59**<br>**3.12**<br>**3.0** | **5.11**<br>**4.00**<br>**4.92**<br>**13.31** | **4.31**<br>**5.04**<br>**7.95**<br>**11.61** |
| | N7(2)-OP2(2) | N7(7)-OP2(7) | N7(13)-OP2(13) | N7(19)-OP2(19) |
| Simulation set-1<br>Simulation set-2<br>Simulation set-3<br>Simulation set-4 | **3.53**<br>**3.93**<br>**6.29**<br>**6.58** | **3.81**<br>**4.48**<br>**4.78**<br>**5.39** | **3.67**<br>**3.66**<br>**10.34**<br>**7.59** | **4.19**<br>**4.32**<br>**11.22**<br>**7.11** |

| **I4-monomer2** | N3(25)-HO2′(25) | N3(31)-HO2′(31) | N3(37)-HO2′(37) | N3(42)-HO2′(42) |
|---|---|---|---|---|
| Simulation set-1<br>Simulation set-2<br>Simulation set-3<br>Simulation set-4 | **6.38**<br>**7.33**<br>**6.38**<br>**5.71** | **2.29**<br>**5.73**<br>**10.31**<br>**4.93** | **2.64**<br>**3.48**<br>**5.02**<br>**3.02** | **3.61**<br>**4.91**<br>**5.48**<br>**4.28** |
| | N7(25)-OP2(25) | N7(31)-OP2(31) | N7(37)-OP2(37) | N7(42)-OP2(42) |
| Simulation set-1<br>Simulation set-2<br>Simulation set-3<br>Simulation set-4 | **- - -**<br>**- - -**<br>**0.69**<br>**- - -** | **- - -**<br>**- - -**<br>**5.17**<br>**0.09** | **- - -**<br>**- - -**<br>**3.12**<br>**0.28** | **- - -**<br>**- - -**<br>**2.34**<br>**- - -** |

| **G4-monomer2** | N3(25)-HO2′(25) | N3(31)-HO2′(31) | N3(37)-HO2′(37) | N3(42)-HO2′(42) |
|---|---|---|---|---|
| Simulation set-1<br>Simulation set-2<br>Simulation set-3<br>Simulation set-4 | **11.68**<br>**10.72**<br>**11.72**<br>**11.14** | **19.89**<br>**22.31**<br>**22.09**<br>**26.52** | **3.93**<br>**3.52**<br>**5.16**<br>**2.81** | **6.14**<br>**3.46**<br>**4.31**<br>**3.33** |
| | (N2-)H22(37)-OP2(25) | (N2-)H22(31)-OP2(37) | (N2-)H22(42)-OP2(31) | (N2-)H22(25)-OP2(42) |
| Simulation set-1<br>Simulation set-2<br>Simulation set-3<br>Simulation set-4 | **52.78**<br>**40.63**<br>**33.70**<br>**37.32** | **32.45**<br>**15.06**<br>**24.71**<br>**31.35** | **56.25**<br>**54.38**<br>**45.23**<br>**57.18** | **48.65**<br>**43.82**<br>**38.98**<br>**43.49** |
| **A4-monomer2** | N3(25)-HO2′(25) | N3(31)-HO2′(31) | N3(37)-HO2′(37) | N3(42)-HO2′(42) |
| Simulation set-1<br>Simulation set-2<br>Simulation set-3<br>Simulation set-4 | **7.18**<br>**1.23**<br>**5.84**<br>**9.41** | **5.07**<br>**0.77**<br>**8.16**<br>**9.86** | **2.12**<br>**0.26**<br>**1.30**<br>**0.30** | **5.26**<br>**0.18**<br>**1.97**<br>**6.81** |
| | N7(25)-OP2(25) | N7(31)-OP2(31) | N7(37)-OP2(37) | N7(42)-OP2(42) |
| Simulation set-1<br>Simulation set-2<br>Simulation set-3<br>Simulation set-4 | **2.11**<br>**3.48**<br>**7.28**<br>**13.29** | **3.07**<br>**1.78**<br>**6.01**<br>**12.84** | **4.98**<br>**3.17**<br>**8.51**<br>**16.93** | **3.46**<br>**1.15**<br>**5.86**<br>**11.98** |

**(B)** Frequencies (in %) of the **U-G** water bridging (in U*(G-G-G-G) pentad) interactions within the studied dimeric quadruplexes for the four sets of simulations.

| Monomer 1<br>N3(11)-W- - -H3-N3(6) | I4 | G4 | A4 |
|---|---|---|---|
| Simulation set-1<br>Simulation set-2<br>Simulation set-3<br>Simulation set-4 | 0.002<br><0.001<br><0.001<br>0.04 | 0.003<br><0.001<br><0.001<br><0.001 | <0.001<br>0.07<br>0.08<br>0.02 |
| Monomer 2<br>N3(35)-W-N3(30) | | | |
| Simulation set-1<br>Simulation set-2<br>Simulation set-3<br>Simulation set-4 | 0.004<br><0.001<br><0.001<br><0.001 | <0.001<br><0.001<br>0.004<br>0.002 | 0.009<br>0.012<br>0.09<br>0.03 |

**Table S15. (A)** Comparison of (total) Gibbs free energies (obtained from MM-GBSA analysis) of the dimeric quadruplexes and individual monomers for the four sets of simulations. Average and standard deviation (within braces) values are reported.

| *Free energy (Kcal/mol)* | *System* | *I4* | *G4* | *A4* |
|---|---|---|---|---|
| Simulation set-*1* | *Dimer* | *-7224.99 (32.82)* | *-8331.13 (29.91)* | *-7892.85 (31.25)* |
| | *Monomer 1* | *-3498.42 (20.55)* | *-4063.14 (20.44)* | *-3849.81 (20.81)* |
| | *Monomer 2* | *-3653.17 (21.39)* | *-4202.94 (21.19)* | *-3986.42 (21.16)* |
| Simulation set-*2* | *Dimer* | *-7227.11 (29.83)* | *-8327.72 (29.45)* | *-7888.81 (34.99)* |
| | *Monomer 1* | *-3503.92 (20.14)* | *-4057.96 (20.38)* | *-3846.28 (20.85)* |
| | *Monomer 2* | *-3651.89 (20.92)* | *-4204.58 (20.83)* | *-3991.18 (24.40)* |
| Simulation set-*3* | *Dimer* | *-7227.95 (31.19)* | *-8333.96 (29.94)* | *-7877.87 (31.25)* |
| | *Monomer 1* | *-3506.42 (21.04)* | *-4063.54 (20.28)* | *-3829.02 (20.92)* |
| | *Monomer 2* | *-3653.47 (21.25)* | *-4207.679 (20.97)* | *-3991.74 (21.55)* |
| Simulation set-*4* | *Dimer* | *-7229.75 (29.43)* | *-8343.6016 (28.64)* | *-7891.01 (30.41)* |
| | *Monomer 1* | *-3515.97 (20.35)* | *-4066.6316 (20.05)* | *-3832.75 (20.47)* |
| | *Monomer 2* | *-3650.01 (20.84)* | *-4210.0740 (20.55)* | *-3980.18 (21.74)* |

**(B)** Comparison of binding free energies (obtained from MM-GBSA analysis) of the constituent monomers in the studied dimeric quadruplexes for the four sets/simulation sets of simulations. Average and standard deviation (within braces) values are reported.

| | *ΔG (total) kcal/mol* | *ΔG (QH) kcal/mol* | *ΔS (QH) kcal/mol* |
|---|---|---|---|
| *I4* | *-73.40 (8.74)*<br>*-71.30 (5.36)*<br>*-68.05 (5.70)*<br>*-63.77 (5.33)* | *-42.00*<br>*-29.45*<br>*-36.90*<br>*-24.21* | *-31.40*<br>*-41.85*<br>*-31.14*<br>*-39.56* |
| *G4* | *-65.04 (6.19)*<br>*-65.18 (5.69)*<br>*-62.74 (5.86)*<br>*-66.89* (4.99) | *-27.37*<br>*-26.43*<br>*-23.34*<br>*-28.29* | *-37.67*<br>*-38.76*<br>*-39.40*<br>*-38.59* |
| *A4* | *-56.63 (5.99)*<br>*-51.34 (7.39)*<br>*-57.11 (6.84)*<br>*-78.08 (7.46)* | *-17.59*<br>*-29.41*<br>*-22.32*<br>*-44.49* | *-39.02*<br>*-21.93*<br>*-34.80*<br>*-33.59* |

**Table S16.** Comparison of N9-pyramidalization (based on the average and standard deviation values for Kappa' dihedral angles (in degrees)) for the tetrads at position 2 (from the

5′-direction) within the constituent monomers of the studied dimeric quadruplexes for the four simulation sets. Average and standard deviation (within braces) values are reported.

| Kappa'<br>TETRAD- 2 | I4 | G4 | A4 |
|---|---|---|---|
| Monomer 1 | | | |
| 2 | 3.74 (11.43)<br>9.24 (10.74)<br>8.29 (10.50)<br>4.01 (10.63) | 3.42 (10.28)<br>2.40 (10.38)<br>2.78 (10.26)<br>3.31 (10.31) | 3.57 (10.47)<br>3.52 (10.62)<br>3.70 (10.31)<br>5.69 (10.64) |
| 7 | 10.82 (10.29)<br>12.04 (9.48)<br>6.94 (10.39)<br>5.98 (10.56) | 5.90 (10.45)<br>6.31 (10.32)<br>4.47 (10.44)<br>5.66 (10.40) | 7.40 (10.78)<br>4.13 (10.85)<br>3.13 (10.33)<br>3.84 (10.86) |
| 13 | 2.64 (11.68)<br>11.92 (9.97)<br>7.67 (10.63)<br>5.63 (10.81) | 4.17 (10.56)<br>4.61 (10.61)<br>3.77 (10.35)<br>3.78 (10.54) | 5.73 (10.87)<br>7.17 (10.71)<br>5.98 (10.42)<br>7.01 (10.64) |
| 19 | 11.46 (11.11)<br>15.62 (9.09)<br>5.38 (11.25)<br>3.94 (10.59) | 4.46 (10.89)<br>6.09 (9.92)<br>4.91 (10.03)<br>6.34 (9.90) | 4.35 (10.70)<br>5.26 (10.65)<br>5.87 (10.32)<br>4.52 (10.64) |
| Monomer 2 | | | |
| 25 | 9.06 (11.09)<br>8.96 (10.22)<br>4.65 (11.30)<br>8.80 (10.88) | 3.04 (10.29)<br>3.38 (10.30)<br>2.41 (10.22)<br>2.93 (10.19) | -0.75 (10.37)<br>4.91 (10.19)<br>3.76 (10.38)<br>5.66 (10.34) |
| 31 | 12.99 (10.03)<br>4.52 (10.83)<br>5.59 (10.40)<br>6.89 (10.82) | 3.11 (9.99)<br>4.58 (9.79)<br>3.16 (9.97)<br>3.14 (9.96) | 0.02 (10.31)<br>4.29 (10.04)<br>5.45 (10.23)<br>4.33 (10.51) |
| 37 | 7.93 (11.32)<br>7.87 (10.43)<br>6.83 (10.79)<br>9.78 (10.66) | 5.99 (10.14)<br>5.82 (10.06)<br>5.64 (10.01)<br>5.05 (10.04) | 3.14 (10.25)<br>6.88 (10.20)<br>4.35 (10.46)<br>6.34 (10.44) |
| 42 | 11.12 (10.38)<br>10.15 (10.16)<br>4.37 (10.74)<br>10.31 (10.42) | 4.31 (10.33)<br>3.87 (10.18)<br>3.84 (10.18)<br>3.93 (10.25) | 1.32 (10.34)<br>4.29 (11.09)<br>5.21 (10.42)<br>5.34 (10.42) |

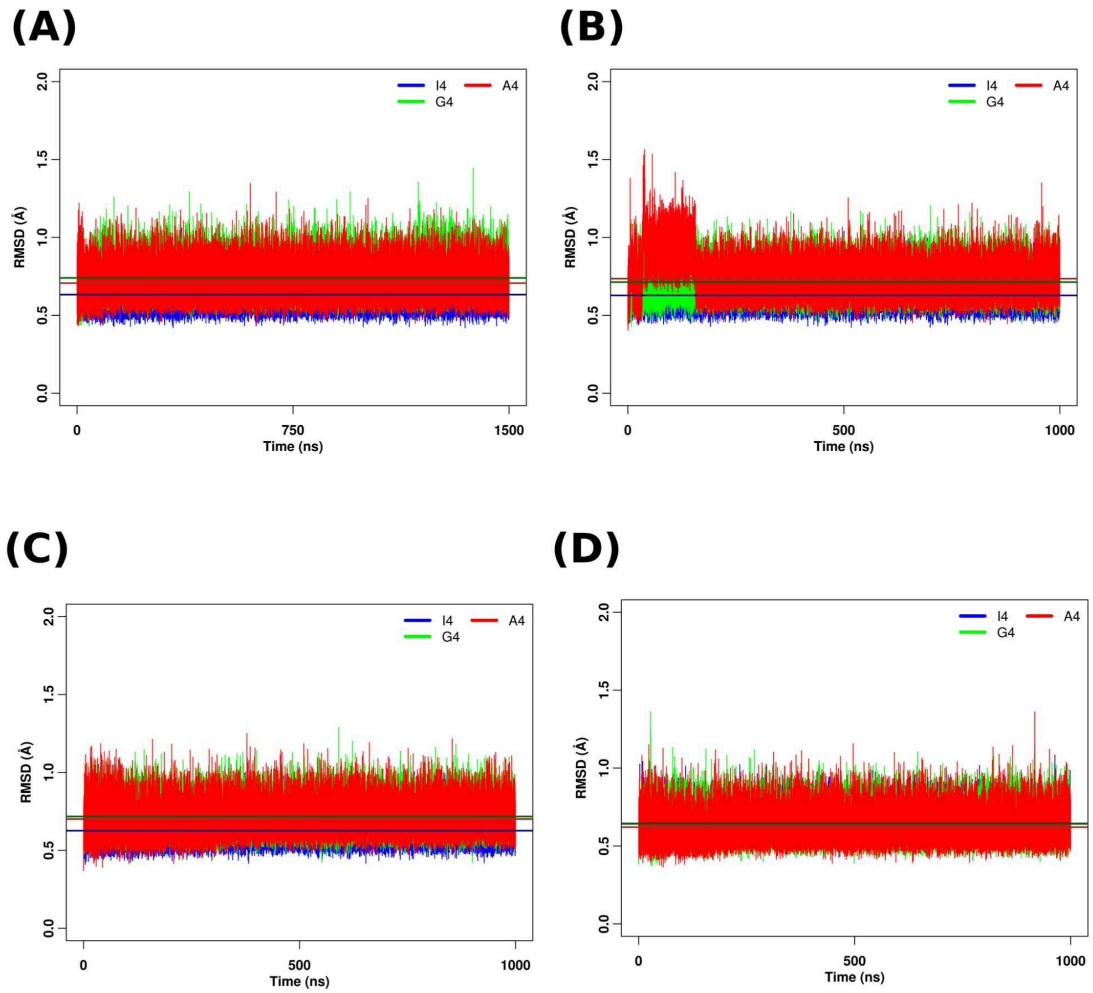


**Figure S1**. RMSD of heavy atoms of the nucleotides of the three (at 5′) tetrads for the monomeric quadruplexes containing the I tetrad (Quad-I4), G-tetrad (Quad-G4) and A-tetrad (Quad-A4) at position 2 respectively for the(A) first (B) second (C) third and (D) fourth simulation sets.

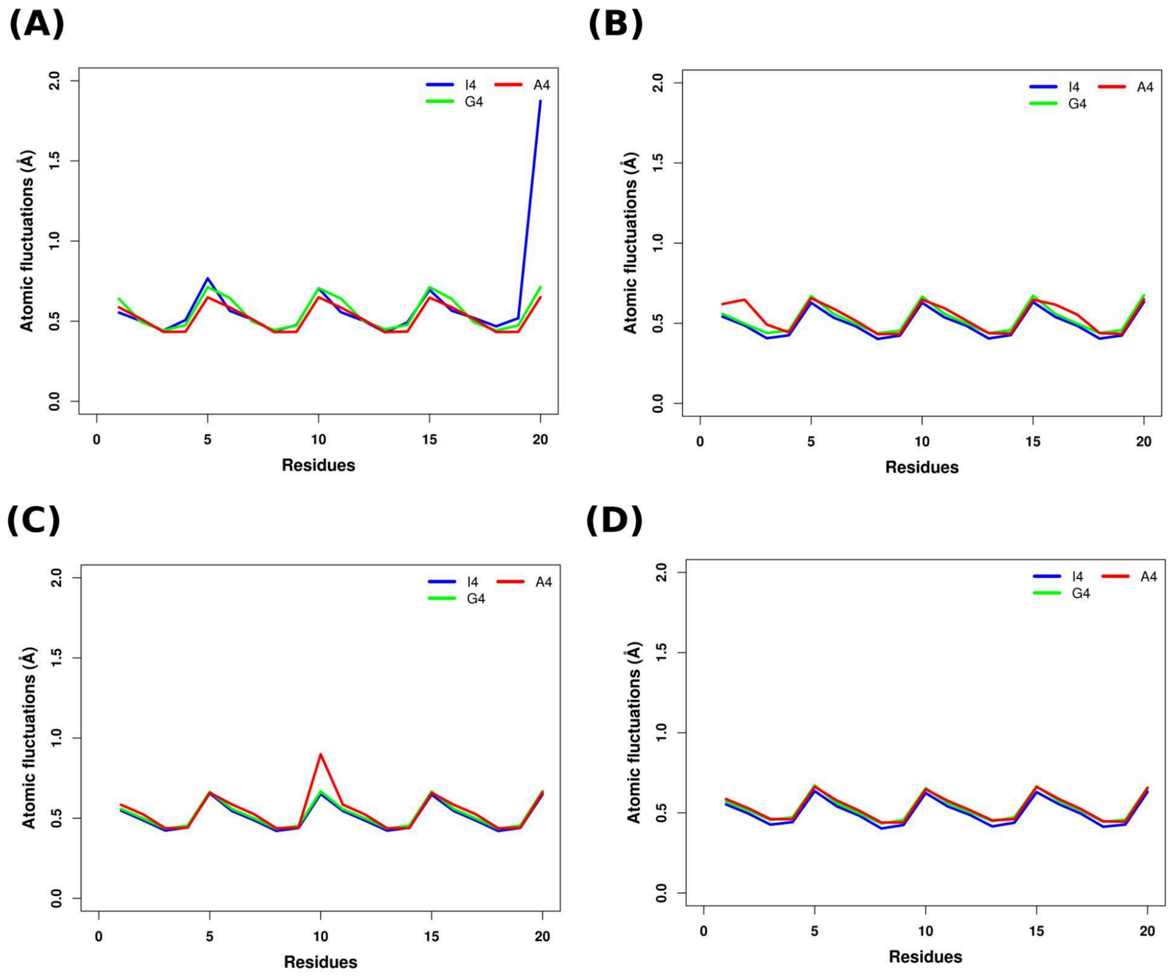


**Figure S2**. RMSF of heavy atoms of the nucleotides for the monomeric quadruplexes containing the I tetrad (Quad-I4), G-tetrad (Quad-G4) and A-tetrad (Quad-A4) at position 2 respectively for the(A) first (B) second (C) third and (D) fourth simulation sets.

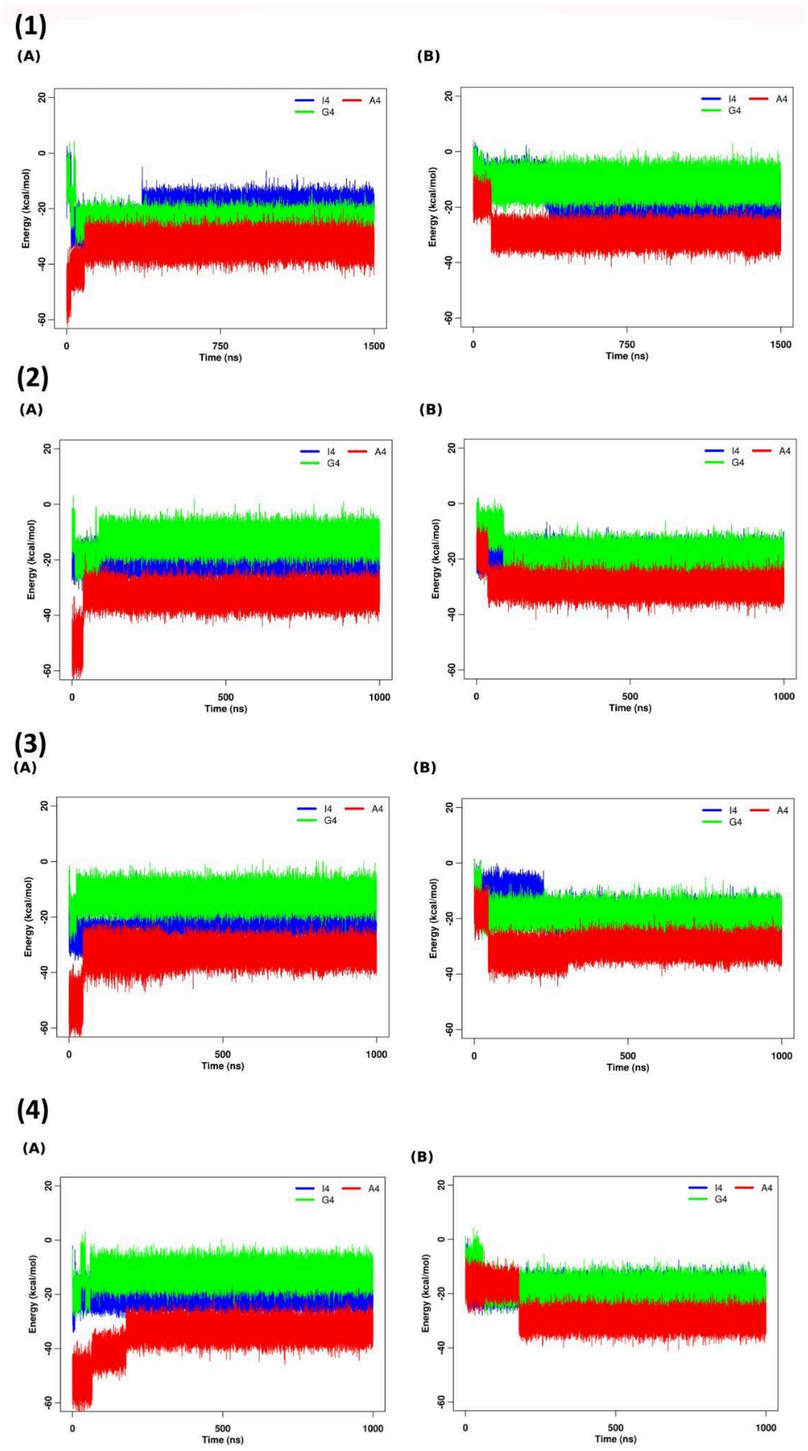


**Figure S3**. Stacking interaction energies for the (A) G4(1)-X4(2) tetrad step and (B) that for the X4(1)-G4(2) tetrad step (where X indicates I/G/A) for the monomeric quadruplexes containing the I tetrad (Quad-I4), G-tetrad (Quad-G4) and A-tetrad (Quad-A4) at position 2 respectively for the (1) first (2) second (3) third and (4) fourth simulation sets.

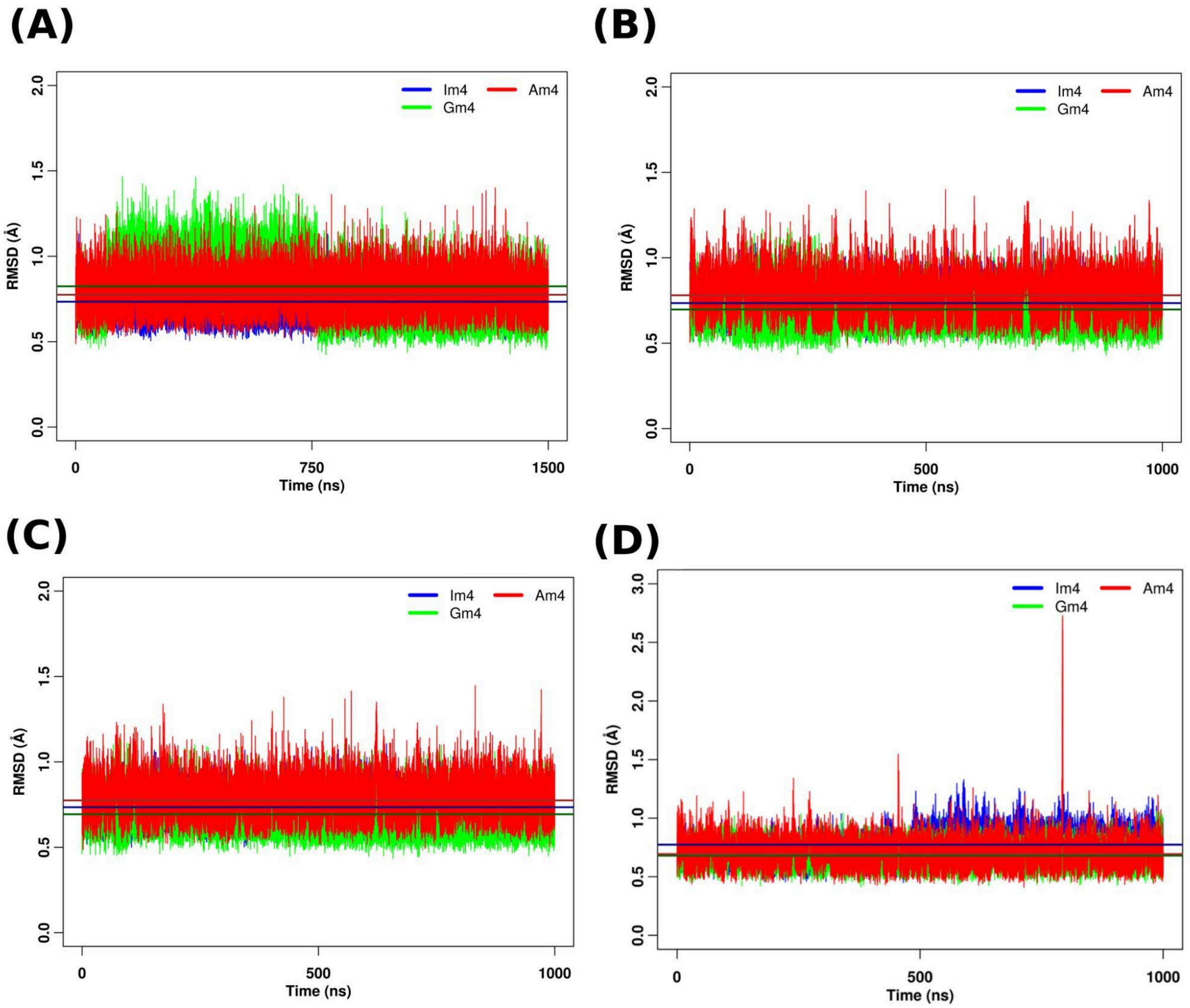


**Figure S4**. RMSD of heavy atoms of the nucleotides of the three (at 5′) tetrad for the quadruplexes containing the I tetrad (Quad-Im4), G-tetrad (Quad-Gm4) and A-tetrad (Quad-Am4) at position 2 respectively for the(A) first (B) second (C) third and (D) fourth simulation sets.

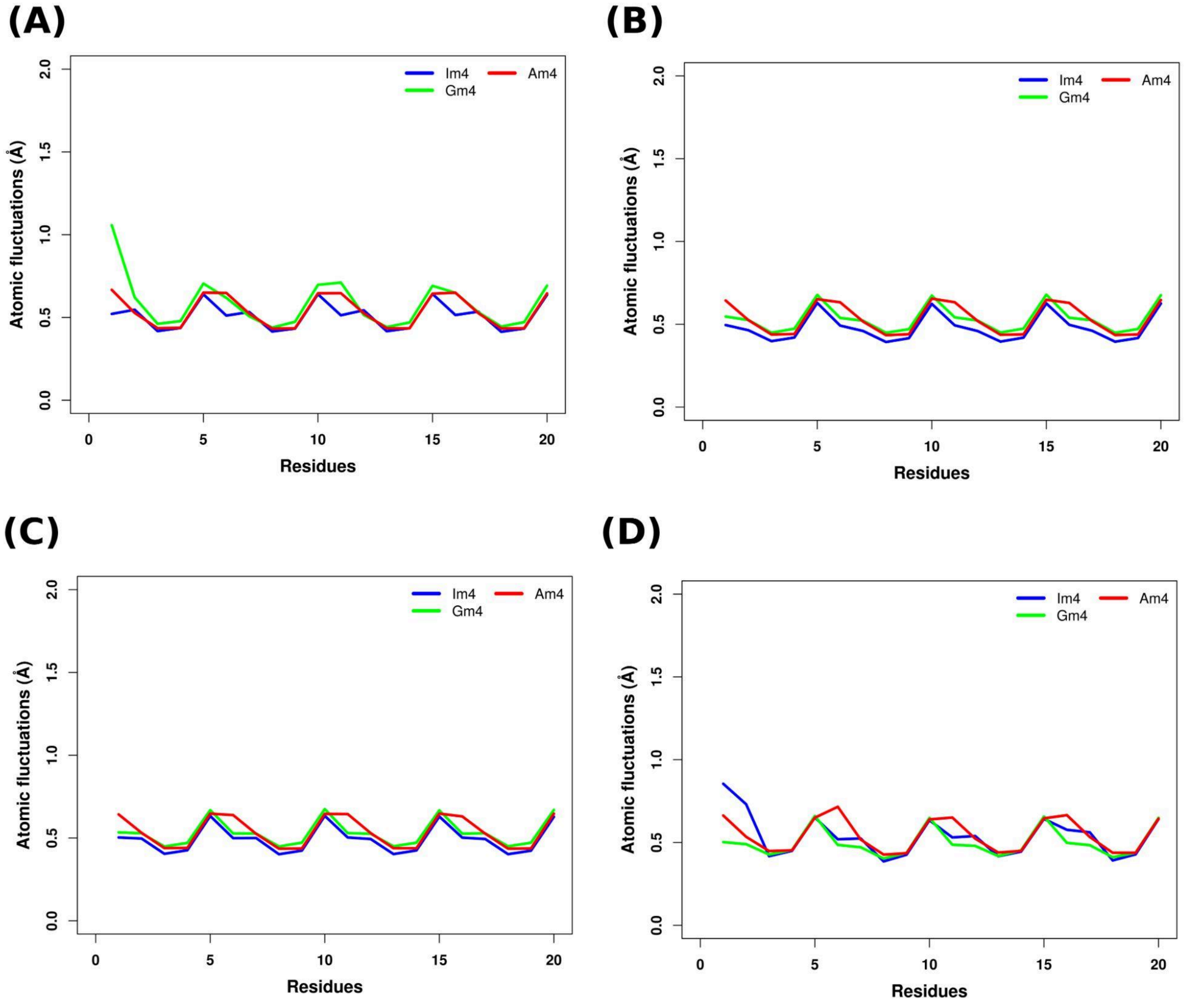


**Figure S5**. RMSF of heavy atoms of the nucleotides for the quadruplexes containing the I tetrad (Quad-I4), G-tetrad (Quad-G4) and A-tetrad (Quad-A4) at position 2 respectively for the(A) first (B) second (C) third and (D) fourth simulation sets.

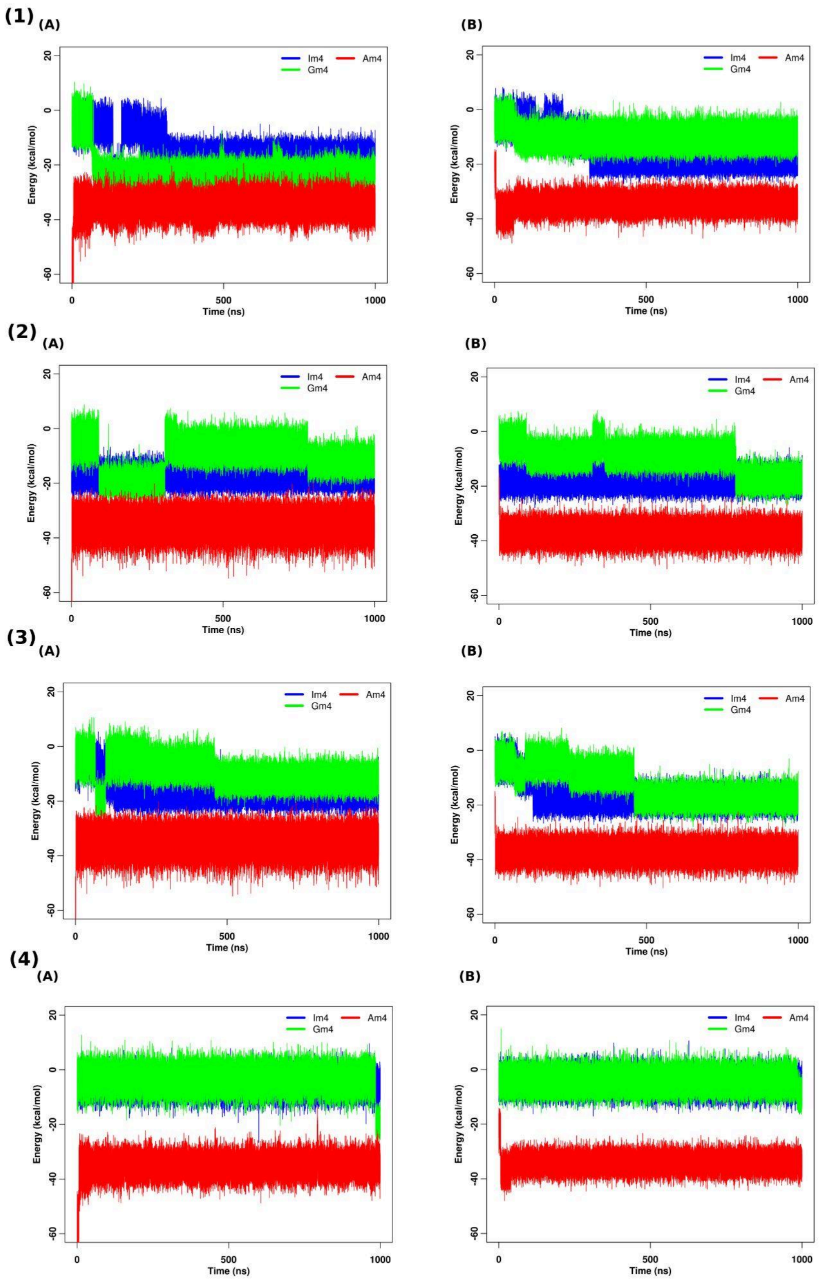


**Figure S6**. Stacking interaction energies for the (A) G4(1)-X4(2) tetrad step and (B) that for the X4(1)-G4(2) tetrad step (where X indicates Im/Gm/Am) for the quadruplexes containing the I tetrad (Quad-Im4), G-tetrad (Quad-Gm4) and A-tetrad (Quad-Am4) at position 2 respectively for the (1) first (2) second (3) third and (4) fourth simulation sets.

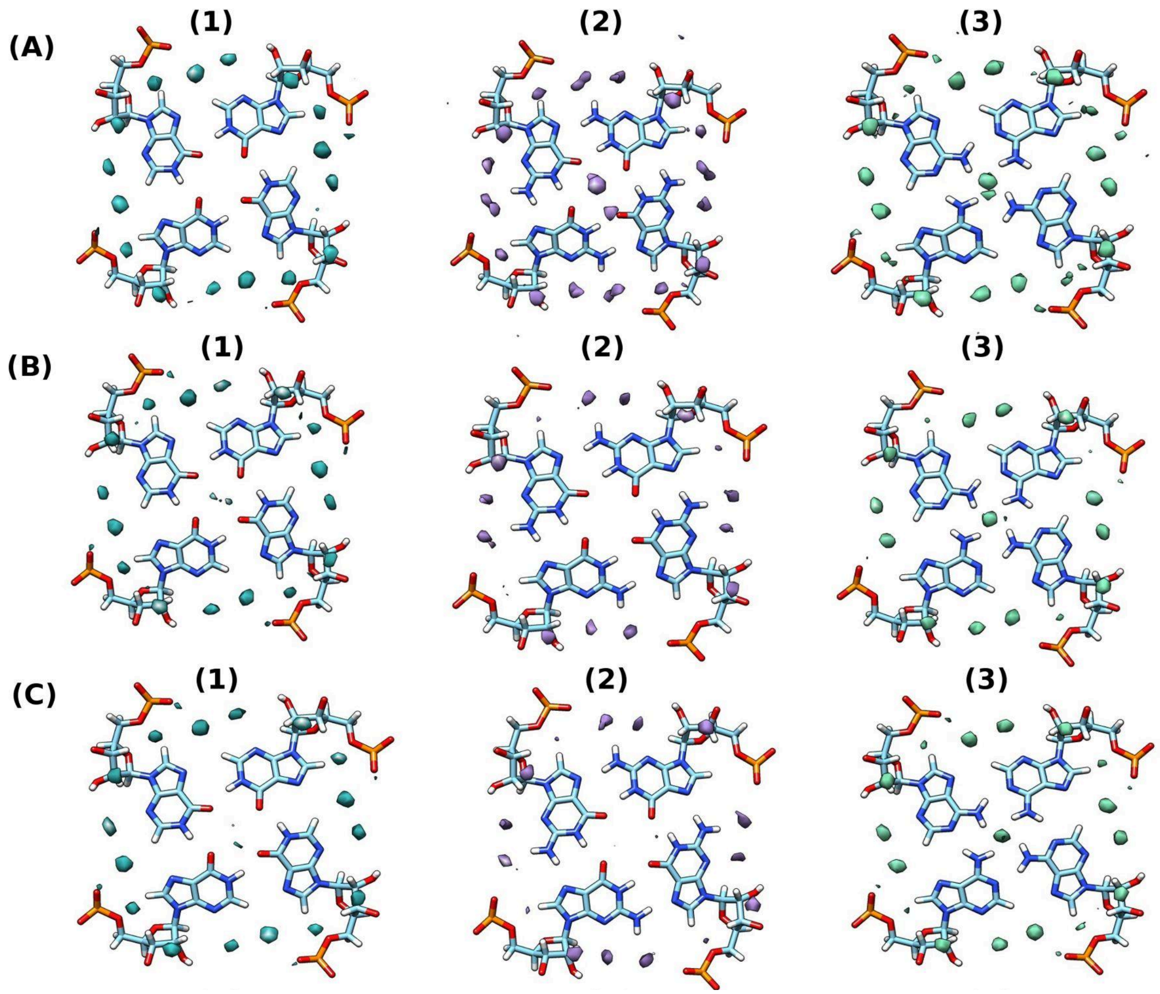


**Figure S7.** Water occupancy maps for (1) I-tetrad (2) G-tetrad (3) A-tetrad respectively for the (A) first (B) second and (C) third simulation sets (around the average structures).

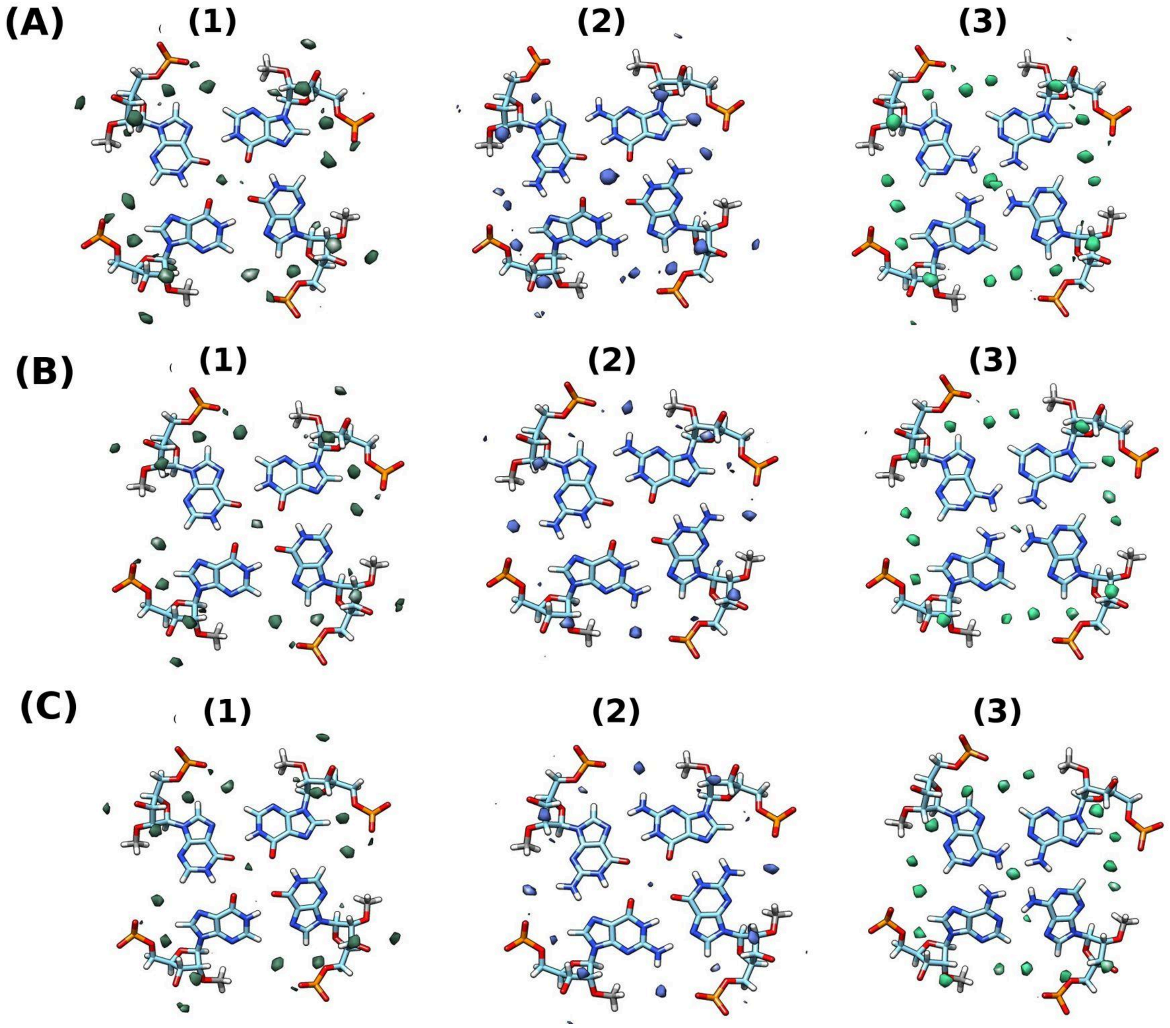


**Figure S8.** Water occupancy maps for (1) Im-tetrad (2) Gm-tetrad (3) Am-tetrad respectively for the (A) first (B) second and (C) third simulation sets (around the average structures).

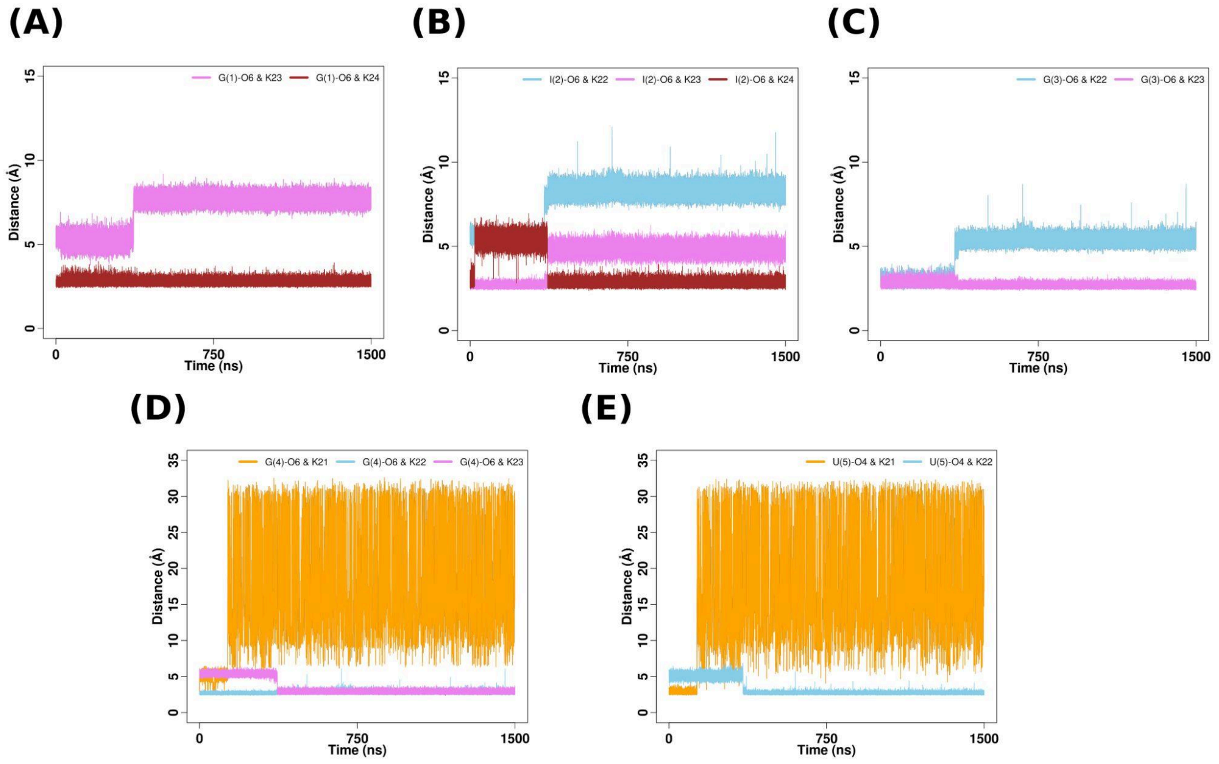


**Figure S9.** Time evolution of distances between (A) G(1)-O6 atom and K+ (23 and 24) (B) I(2)-O6 atom and K+ (22, 23 and 24) (C) G(3)-O6 atom and K+ (22 and 23) (D) G(4)-O6 atom and K+ (21, 22 and 23) and (E) U(5)-O4 atom and K+ (21 and 22) respectively for simulation set-1 for quadruplex containing I-tetrad at position 2.

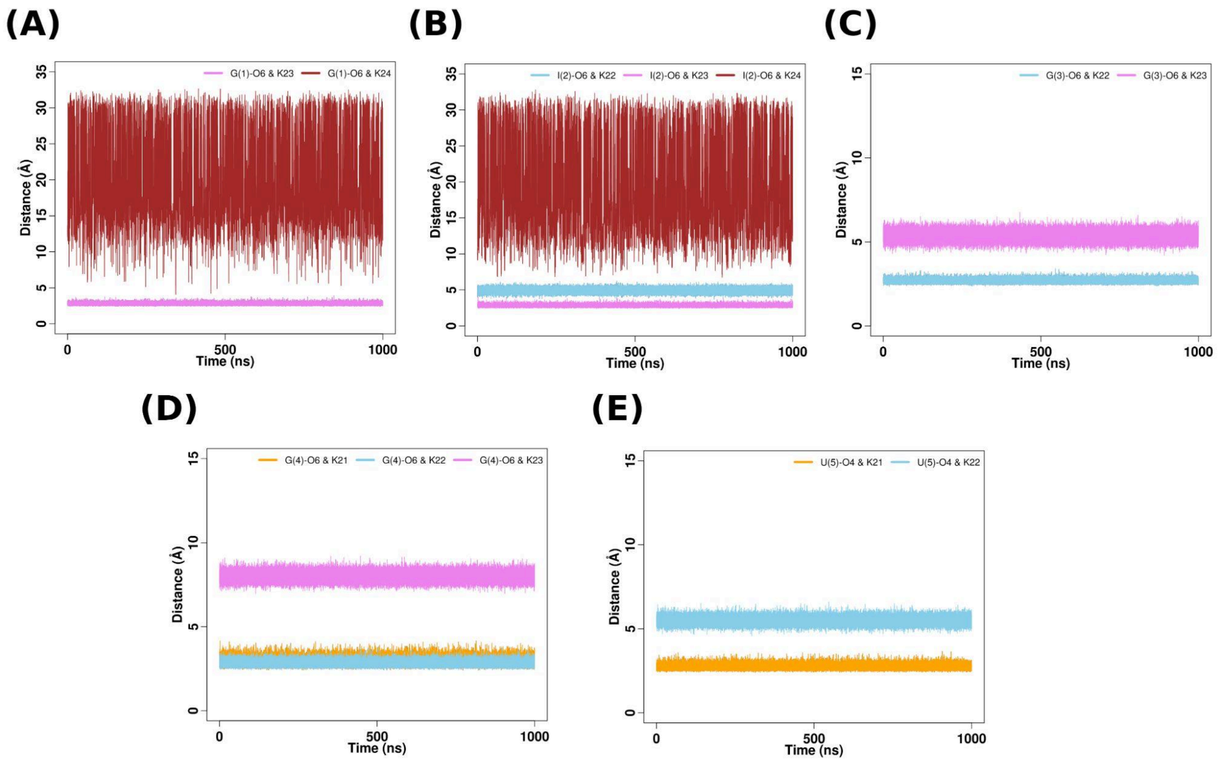


**Figure S10.** Time evolution of distances between (A) G(1)-O6 atom and K+ (23 and 24) (B) I(2)-O6 atom and K+ (22, 23 and 24) (C) G(3)-O6 atom and K+ (22 and 23) (D) G(4)-O6 atom and K+ (21, 22 and 23) and (E) U(5)-O4 atom and K+ (21 and 22) respectively for simulation set-2 for quadruplex containing I-tetrad at position 2.

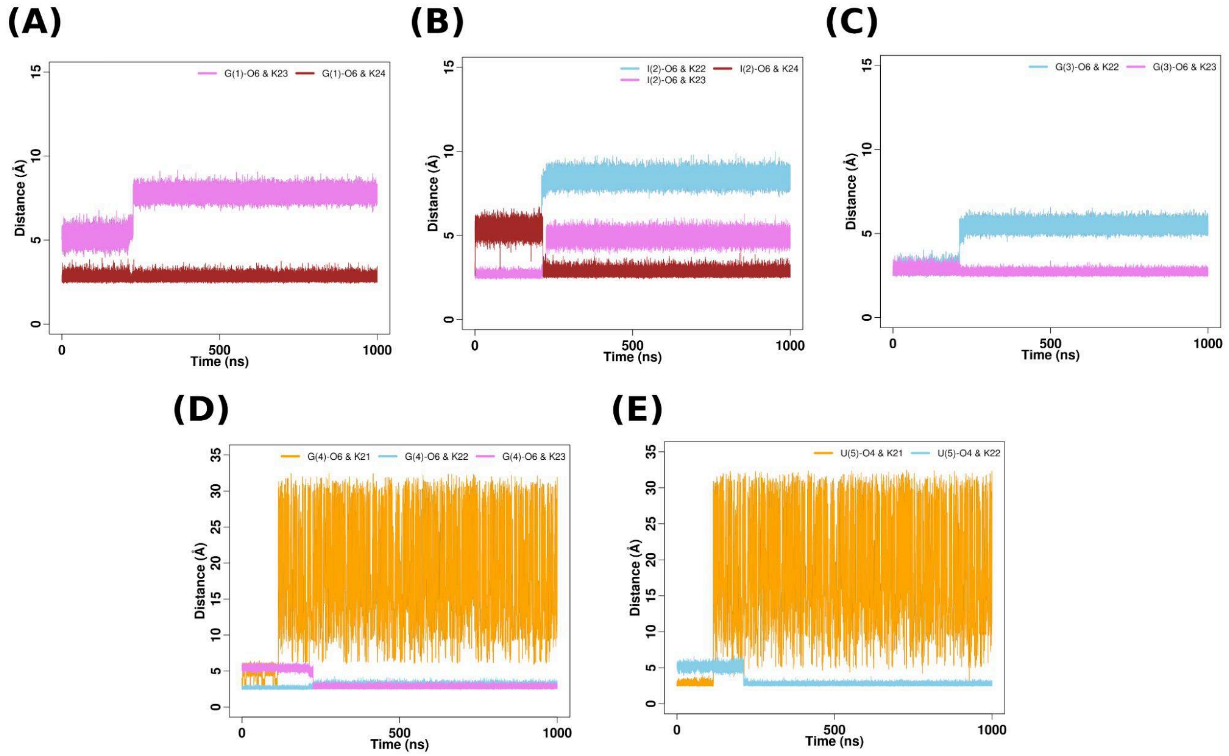


**Figure S11.** Time evolution of distances between (A) G(1)-O6 atom and K+ (23 and 24) (B) I(2)-O6 atom and K+ (22, 23 and 24) (C) G(3)-O6 atom and K+ (22 and 23) (D) G(4)-O6 atom and K+ (21, 22 and 23) and (E) U(5)-O4 atom and K+ (21 and 22) respectively for simulation set-3 for quadruplex containing I-tetrad at position 2.

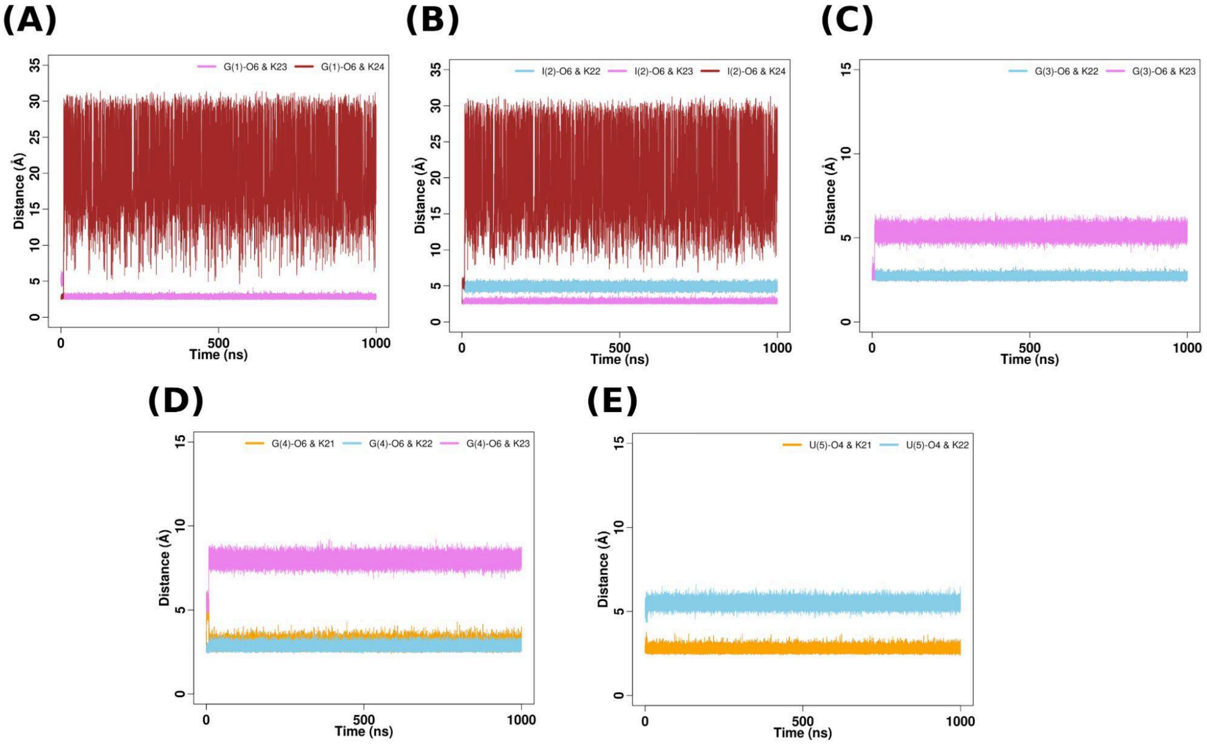


**Figure S12.** Time evolution of distances between (A) G(1)-O6 atom and K+ (23 and 24) (B) I(2)-O6 atom and K+ (22, 23 and 24) (C) G(3)-O6 atom and K+ (22 and 23) (D) G(4)-O6 atom and K+ (21, 22 and 23) and (E) U(5)-O4 atom and K+ (21 and 22) respectively for simulation set-4 for quadruplex containing I-tetrad at position 2.

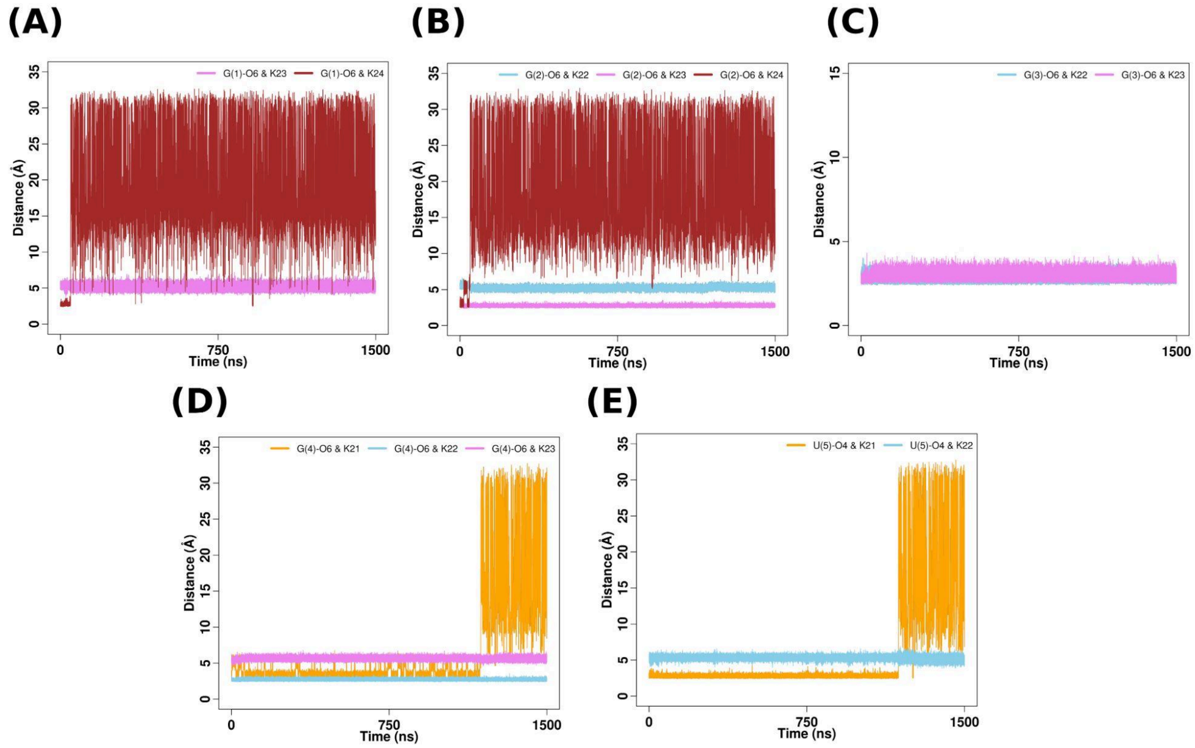


**Figure S13.** Time evolution of distances between (A) G(1)-O6 atom and K+ (23 and 24) (B) G(2)-O6 atom and K+ (22, 23 and 24) (C) G(3)-O6 atom and K+ (22 and 23) (D) G(4)-O6 atom and K+ (21, 22 and 23) and (E) U(5)-O4 atom and K+ (21 and 22) respectively for simulation set-1 for quadruplex containing G-tetrad at position 2.

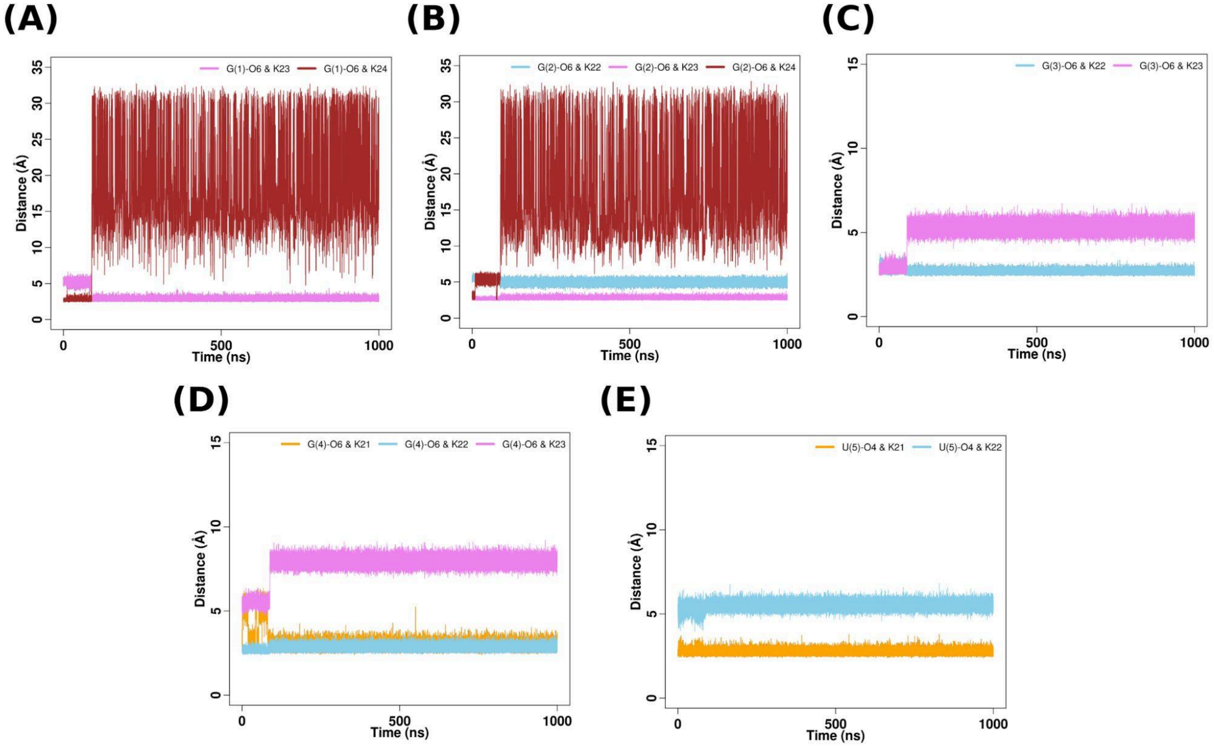


**Figure S14.** Time evolution of distances between (A) G(1)-O6 atom and K+ (23 and 24) (B) G(2)-O6 atom and K+ (22, 23 and 24) (C) G(3)-O6 atom and K+ (22 and 23) (D) G(4)-O6 atom and K+ (21, 22 and 23) and (E) U(5)-O4 atom and K+ (21 and 22) respectively for simulation set-2 for quadruplex containing G-tetrad at position 2.

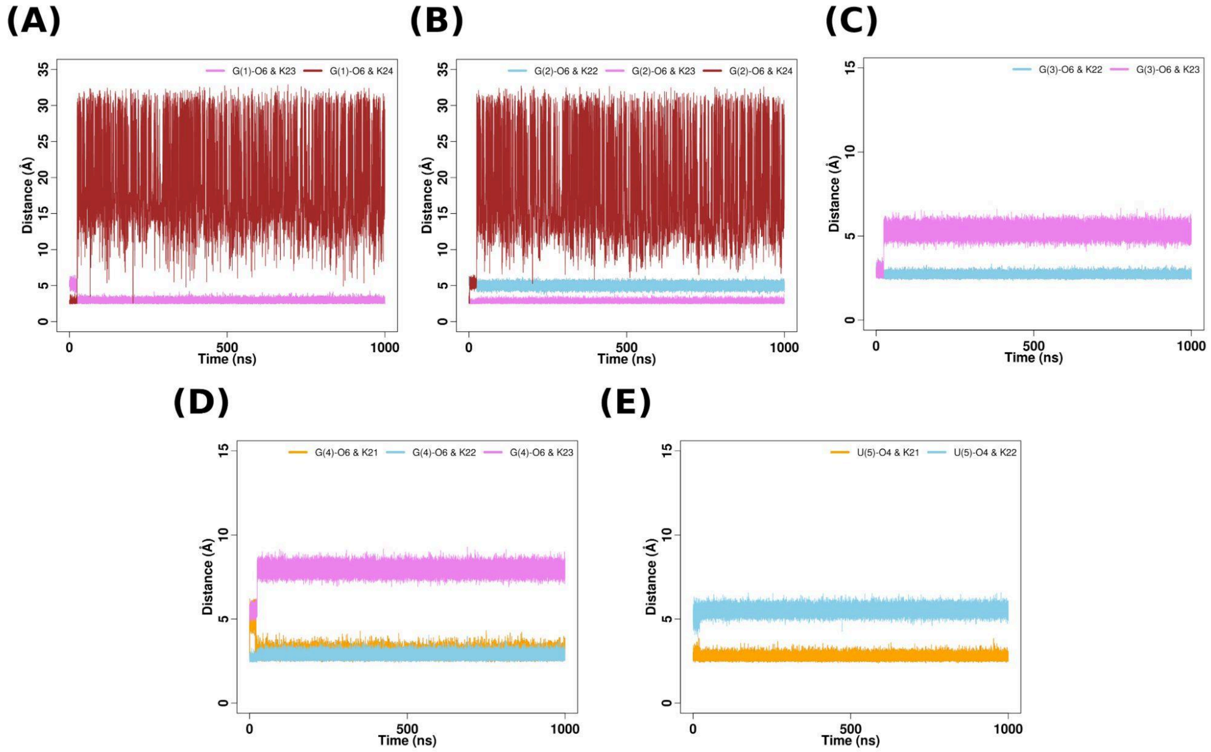


**Figure S15.** Time evolution of distances between (A) G(1)-O6 atom and K+ (23 and 24) (B) G(2)-O6 atom and K+ (22, 23 and 24) (C) G(3)-O6 atom and K+ (22 and 23) (D) G(4)-O6 atom and K+ (21, 22 and 23) and (E) U(5)-O4 atom and K+ (21 and 22) respectively for simulation set-3 for quadruplex containing G-tetrad at position 2.

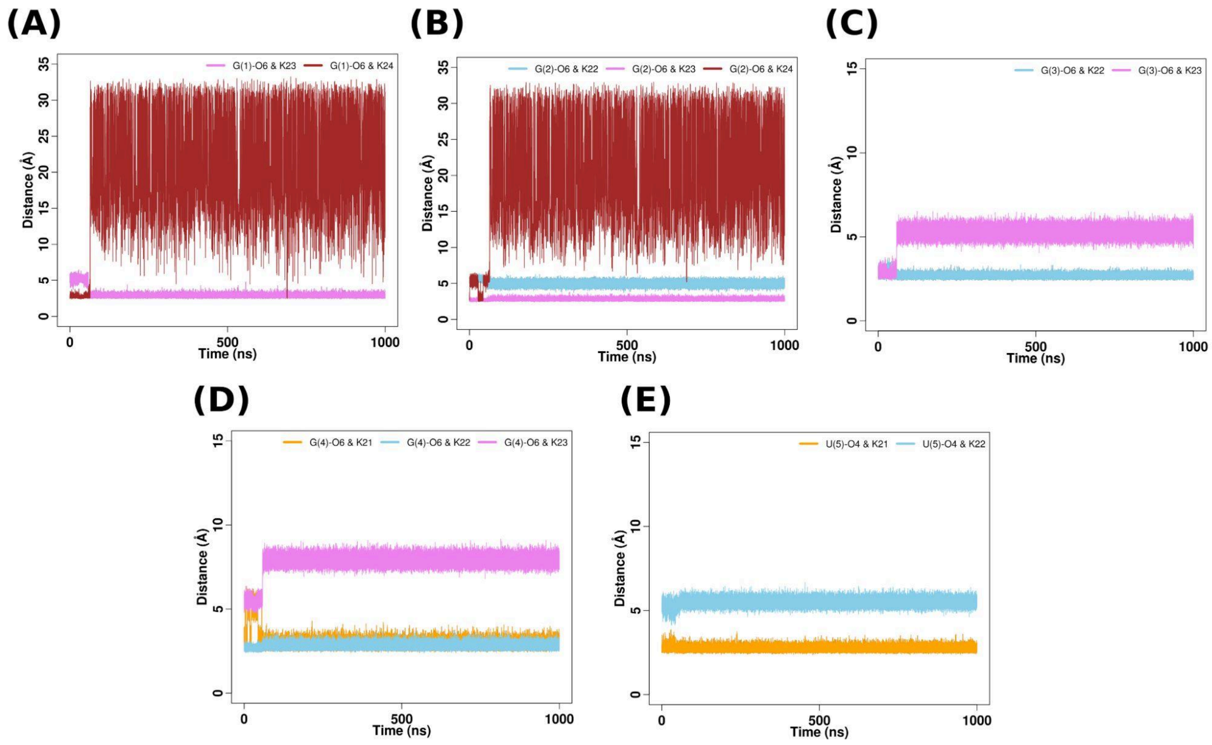


**Figure S16.** Time evolution of distances between (A) G(1)-O6 atom and K+ (23 and 24) (B) G(2)-O6 atom and K+ (22, 23 and 24) (C) G(3)-O6 atom and K+ (22 and 23) (D) G(4)-O6 atom and K+ (21, 22 and 23) and (E) U(5)-O4 atom and K+ (21 and 22) respectively for simulation set-4 for quadruplex containing G-tetrad at position 2.

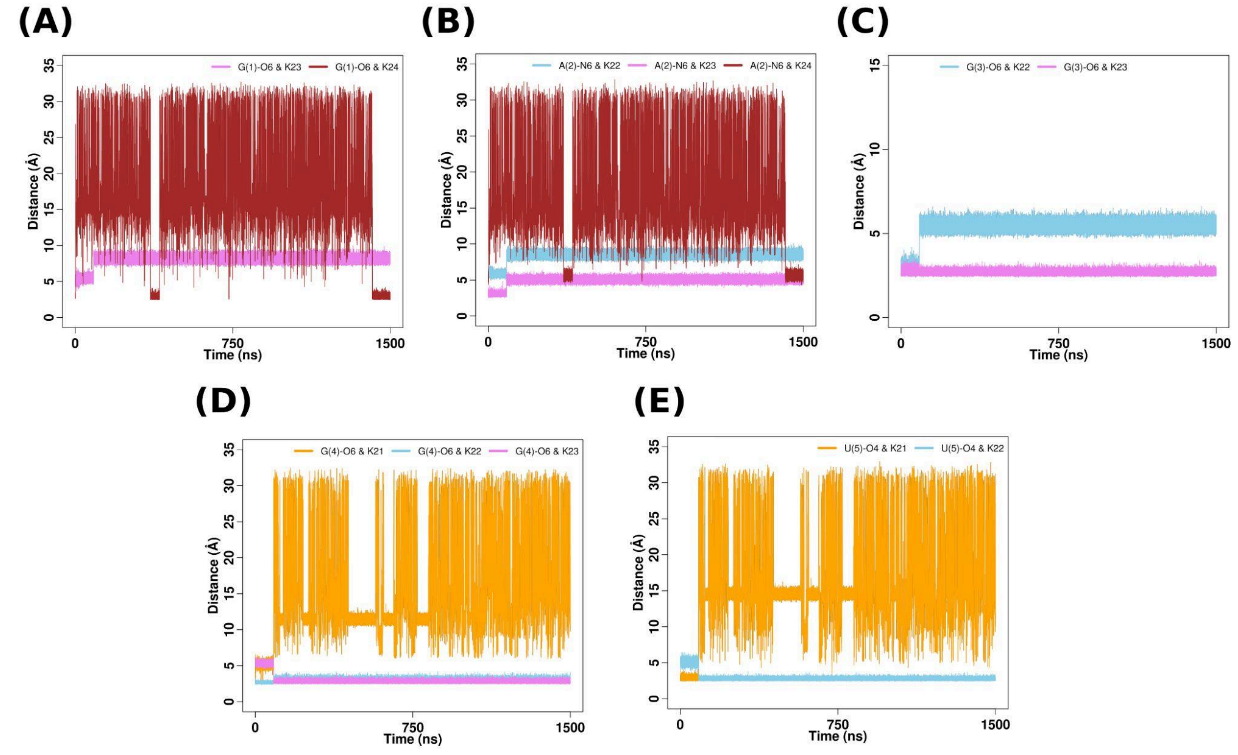


**Figure S17.** Time evolution of distances between (A) G(1)-O6 atom and K+ (23 and 24) (B) A(2)-N6 atom and K+ (22, 23 and 24) (C) G(3)-O6 atom and K+ (21, 22 and 23) (D) G(4)-O6 atom and K+ (21 and 22) and (E) U(5)-O4 atom and K+ (21, 22 and 23) respectively for simulation set-1 for quadruplex containing A-tetrad at position 2.

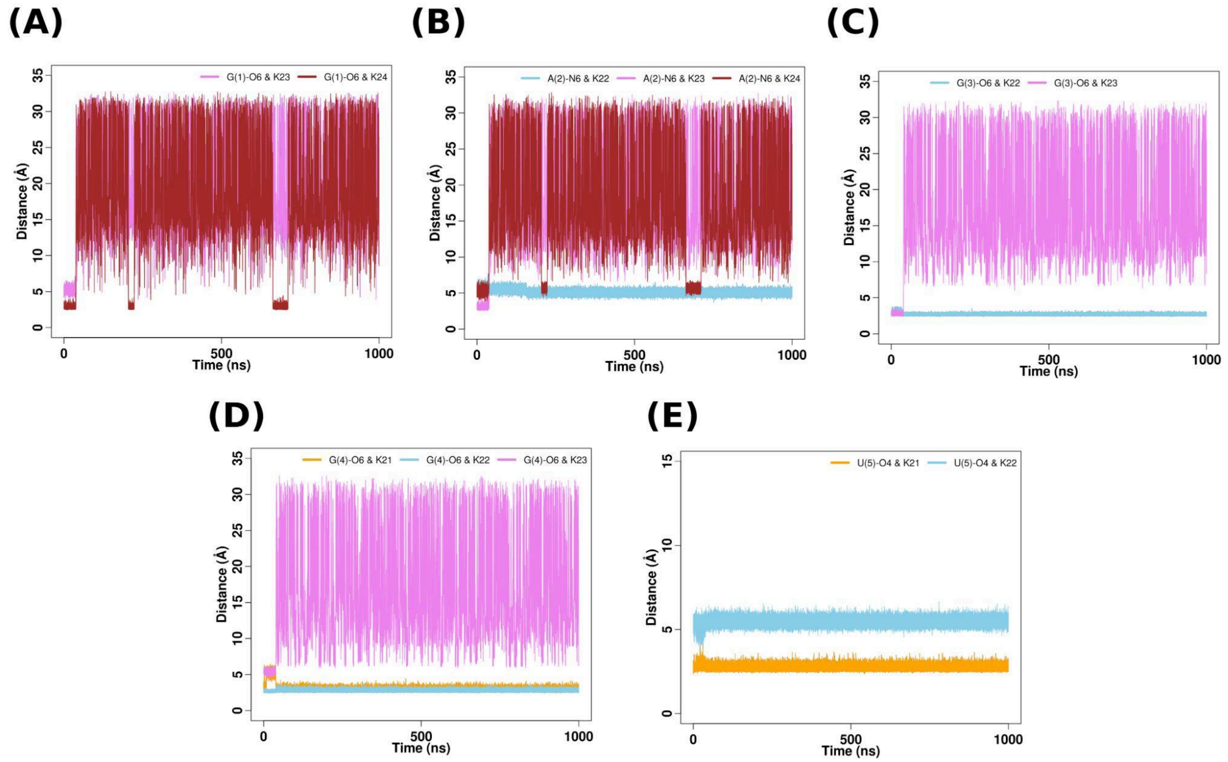


**Figure S18.** Time evolution of distances between (A) G(1)-O6 atom and K+ (23 and 24) (B) A(2)-N6 atom and K+ (22, 23 and 24) (C) G(3)-O6 atom and K+ (21, 22 and 23) (D) G(4)-O6 atom and K+ (21 and 22) and (E) U(5)-O4 atom and K+ (21, 22 and 23) respectively for simulation set-2 for quadruplex containing A-tetrad at position 2.

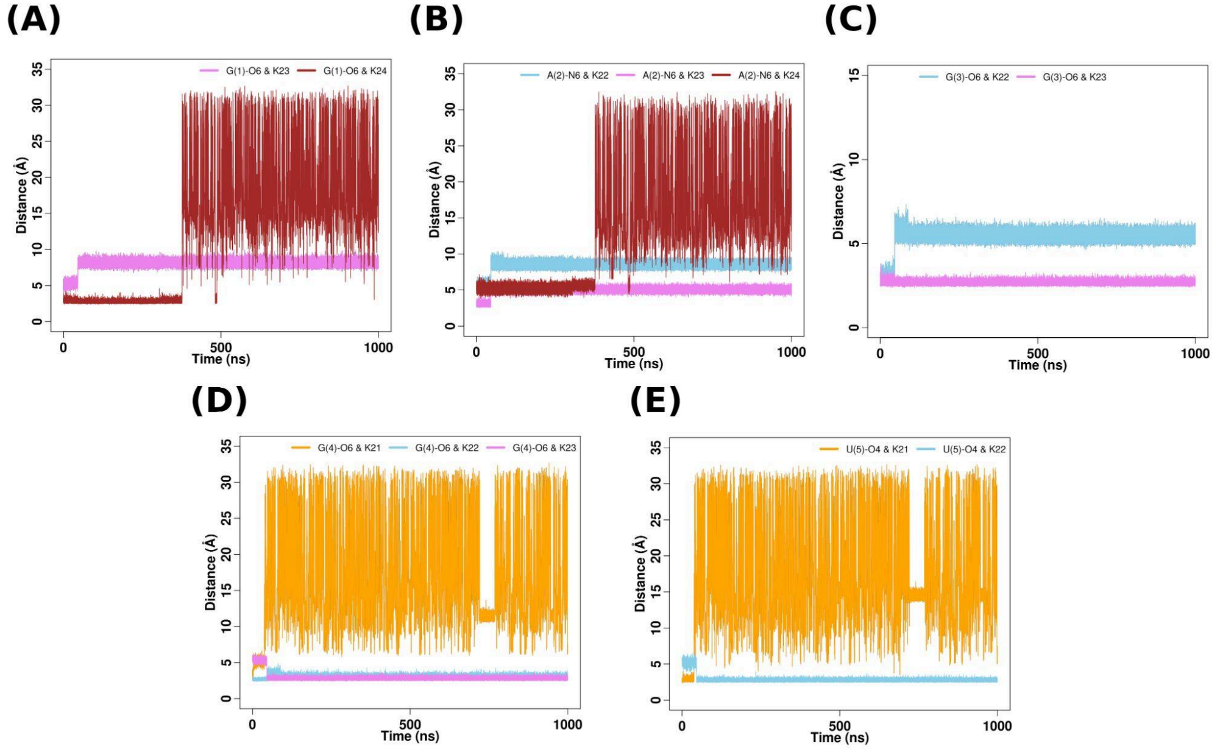


**Figure S19.** Time evolution of distances between (A) G(1)-O6 atom and K+ (23 and 24) (B) A(2)-N6 atom and K+ (22, 23 and 24) (C) G(3)-O6 atom and K+ (21, 22 and 23) (D) G(4)-O6 atom and K+ (21 and 22) and (E) U(5)-O4 atom and K+ (21, 22 and 23) respectively for simulation set-3 for quadruplex containing A-tetrad at position 2.

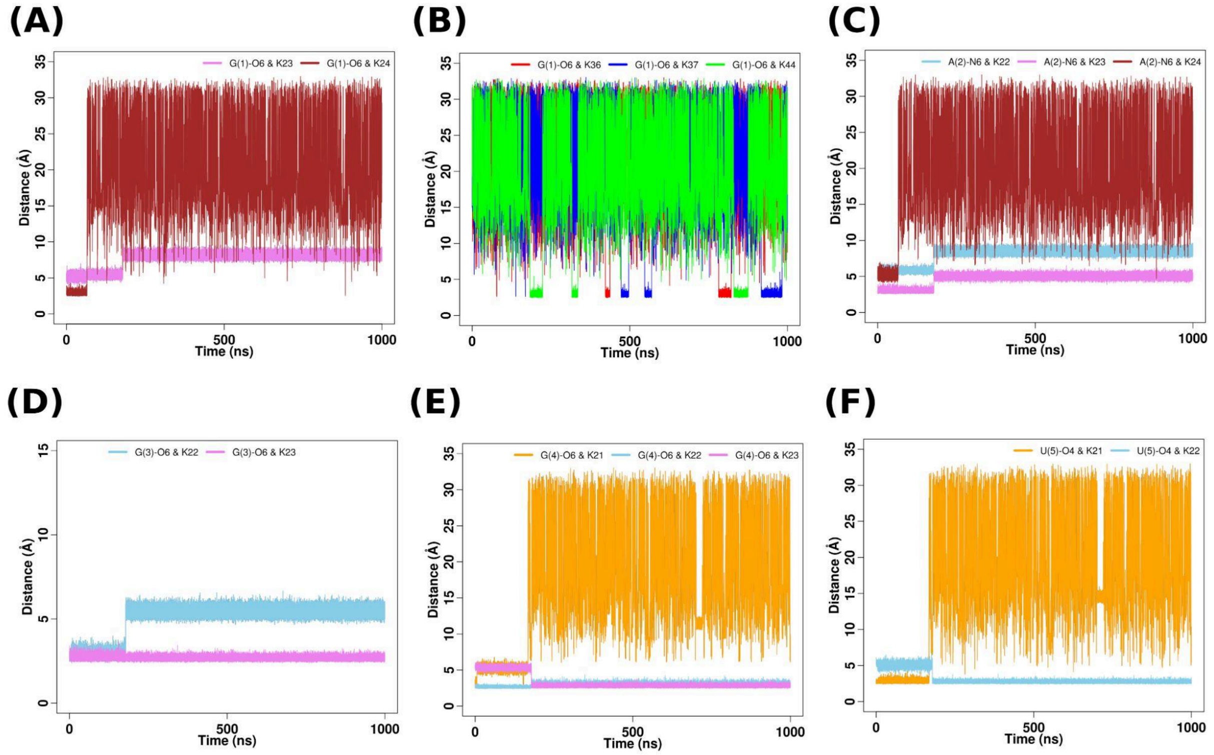


**Figure S20.** Time evolution of distances between (A) G(1)-O6 atom and K+ (23 and 24) (B) G(1)-O6 atom and K+ (36, 37 and 44) (C) A(2)-N6 atom and K+ (22, 23 and 24) (D) G(3)-O6 atom and K+ (21, 22 and 23) (E) G(4)-O6 atom and K+ (21 and 22) and (F) U(5)-O4 atom and K+ (21, 22 and 23) respectively for simulation set-4 for quadruplex containing A-tetrad at position 2.

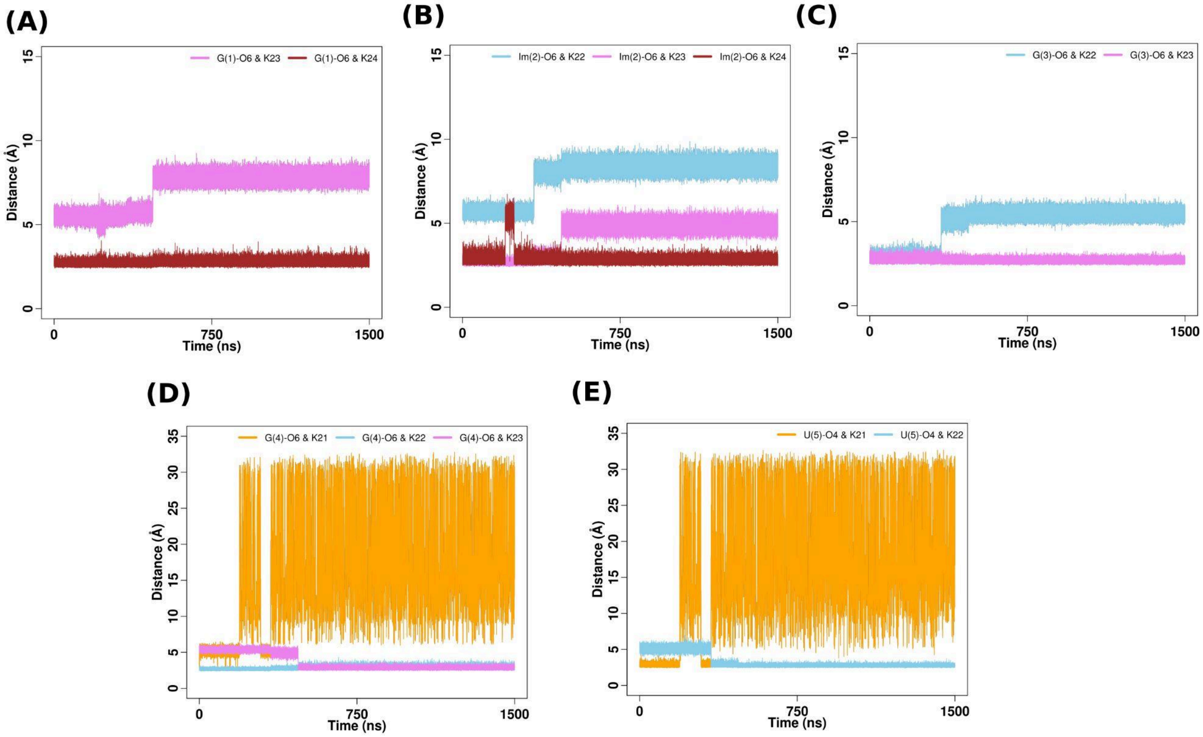


**Figure S21.** Time evolution of distances between (A) G(1)-O6 atom and K+ (23 and 24) (B) Im(2)-O6 atom and K+ (22, 23 and 24) (C) G(3)-O6 atom and K+ (22 and 23) (D) G(4)-O6 atom and K+ (21, 22 and 23) and (E) U(5)-O4 atom and K+ (21 and 22) respectively for simulation set-1 for quadruplex containing Im-tetrad at position 2.

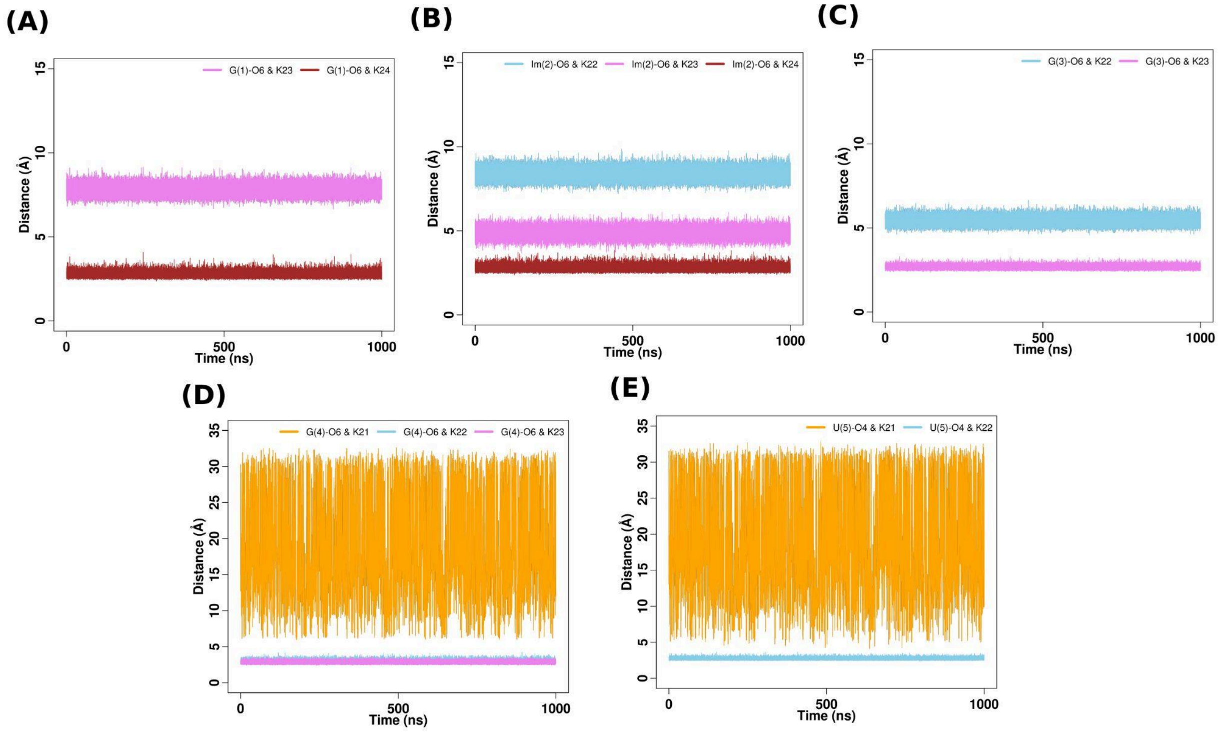


**Figure S22.** Time evolution of distances between (A) G(1)-O6 atom and K+ (23 and 24) (B) Im(2)-O6 atom and K+ (22, 23 and 24) (C) G(3)-O6 atom and K+ (22 and 23) (D) G(4)-O6 atom and K+ (21, 22 and 23) and (E) U(5)-O4 atom and K+ (21 and 22) respectively for simulation set-2 for quadruplex containing Im-tetrad at position 2.

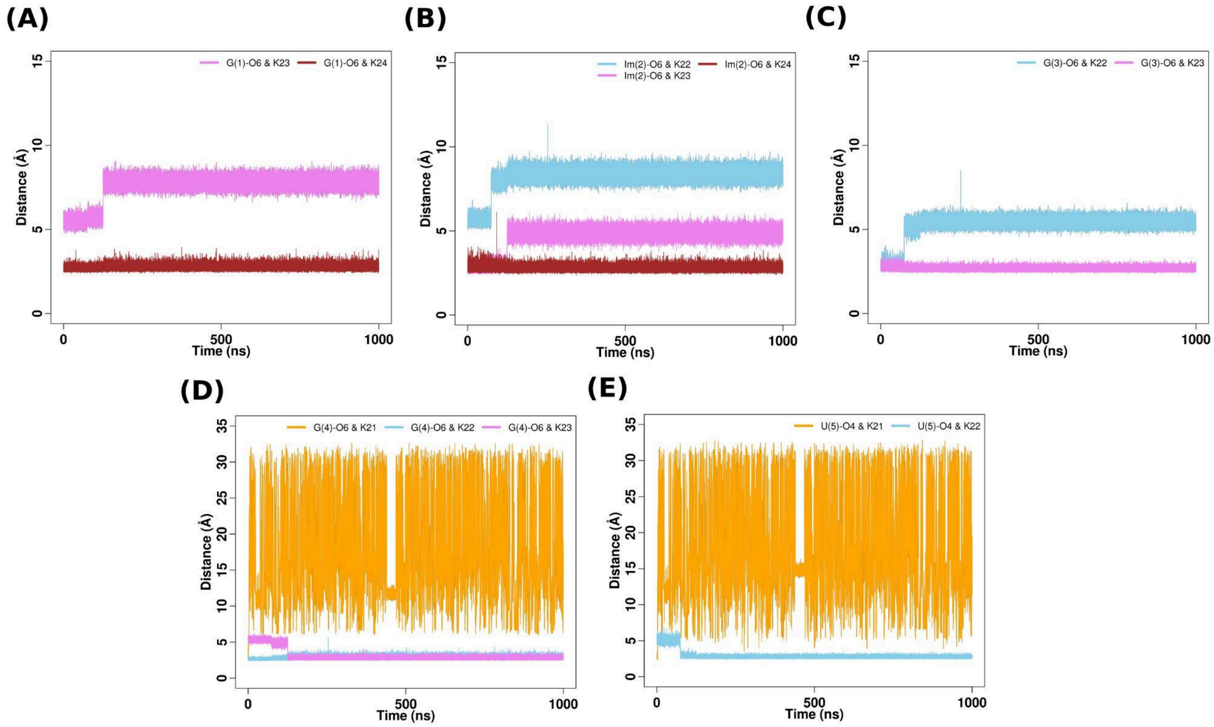


**Figure S23.** Time evolution of distances between (A) G(1)-O6 atom and K+ (23 and 24) (B) Im(2)-O6 atom and K+ (22, 23 and 24) (C) G(3)-O6 atom and K+ (22 and 23) (D) G(4)-O6 atom and K+ (21, 22 and 23) and (E) U(5)-O4 atom and K+ (21 and 22) respectively for simulation set-3 for quadruplex containing Im-tetrad at position 2

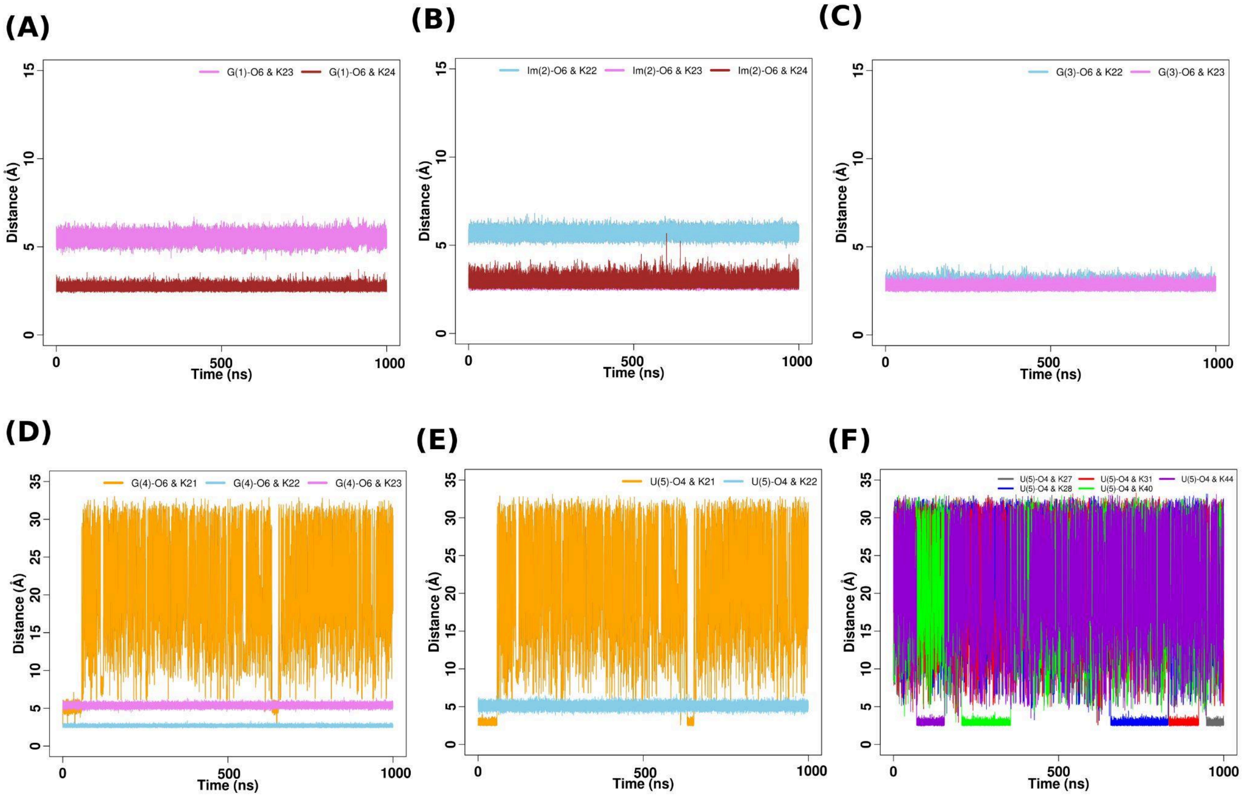


**Figure S24.** Time evolution of distances between (A) G(1)-O6 atom and K+ (23 and 24) (B) Im(2)-O6 atom and K+ (22, 23 and 24) (C) G(3)-O6 atom and K+ (21, 22 and 23) (D) G(4)-O6 atom and K+ (21 and 22), (E) U(5)-O4 atom and K+ (21, 22 and 23) and (F) U(5)-O4 atom and K+ (21, 22 and 23) respectively for simulation set-4 for quadruplex containing Im-tetrad at position 2.

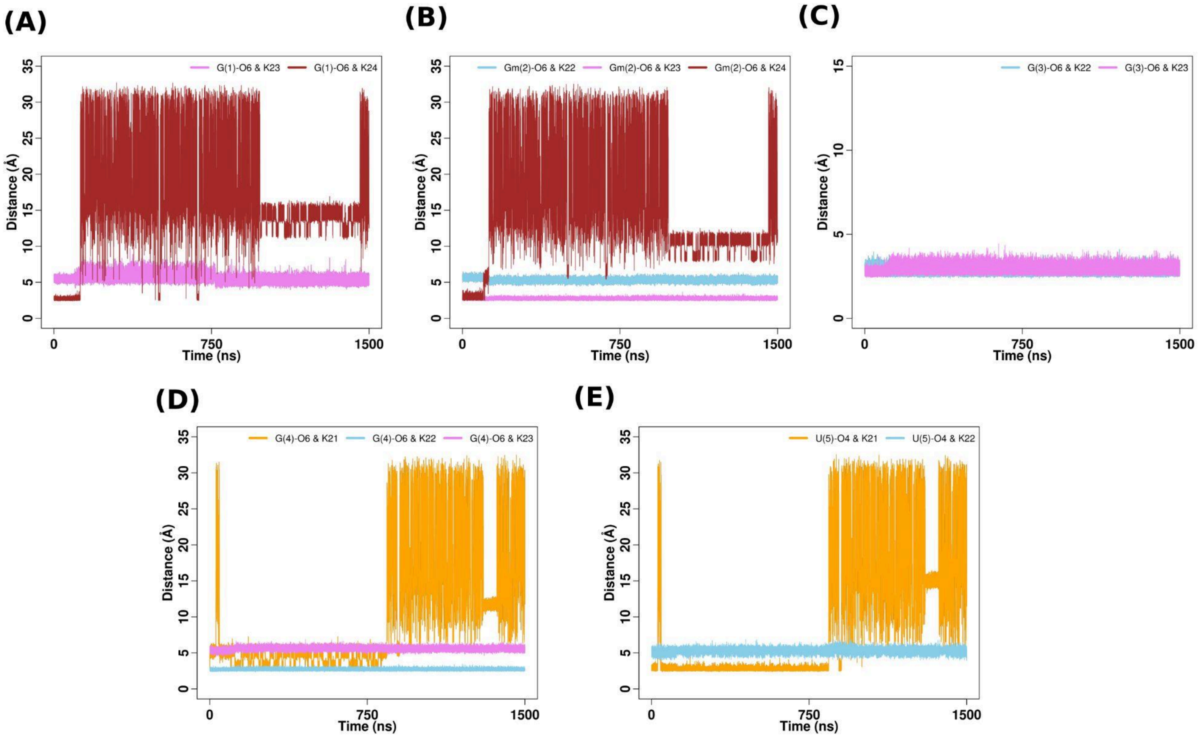


**Figure S25.** Time evolution of distances between (A) G(1)-O6 atom and K+ (23 and 24) (B) Gm(2)-O6 atom and K+ (22, 23 and 24) (C) G(3)-O6 atom and K+ (22 and 23) (D) G(4)-O6 atom and K+ (21, 22 and 23) and (E) U(5)-O4 atom and K+ (21 and 22) respectively for simulation set-1 for quadruplex containing Gm-tetrad at position 2.

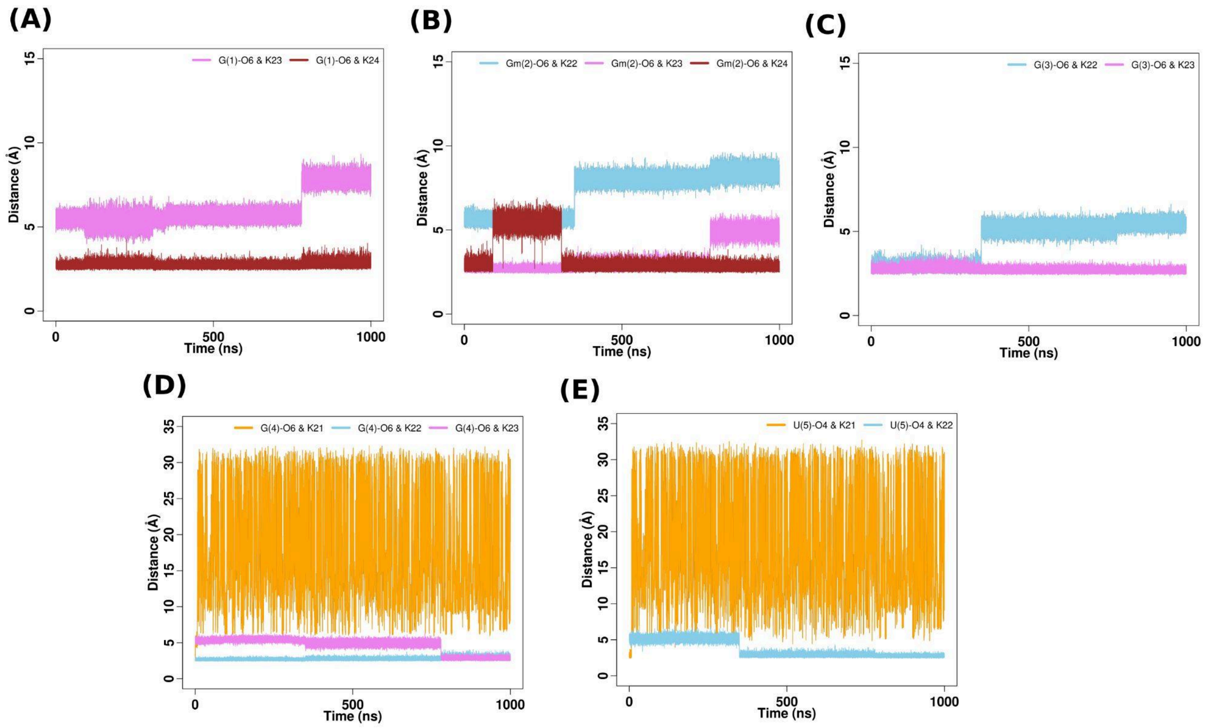


**Figure S26.** Time evolution of distances between (A) G(1)-O6 atom and K+ (23 and 24) (B) Gm(2)-O6 atom and K+ (22, 23 and 24) (C) G(3)-O6 atom and K+ (22 and 23) (D) G(4)-O6 atom and K+ (21, 22 and 23) and (E) U(5)-O4 atom and K+ (21 and 22) respectively for simulation set-2 for quadruplex containing Gm-tetrad at position 2.

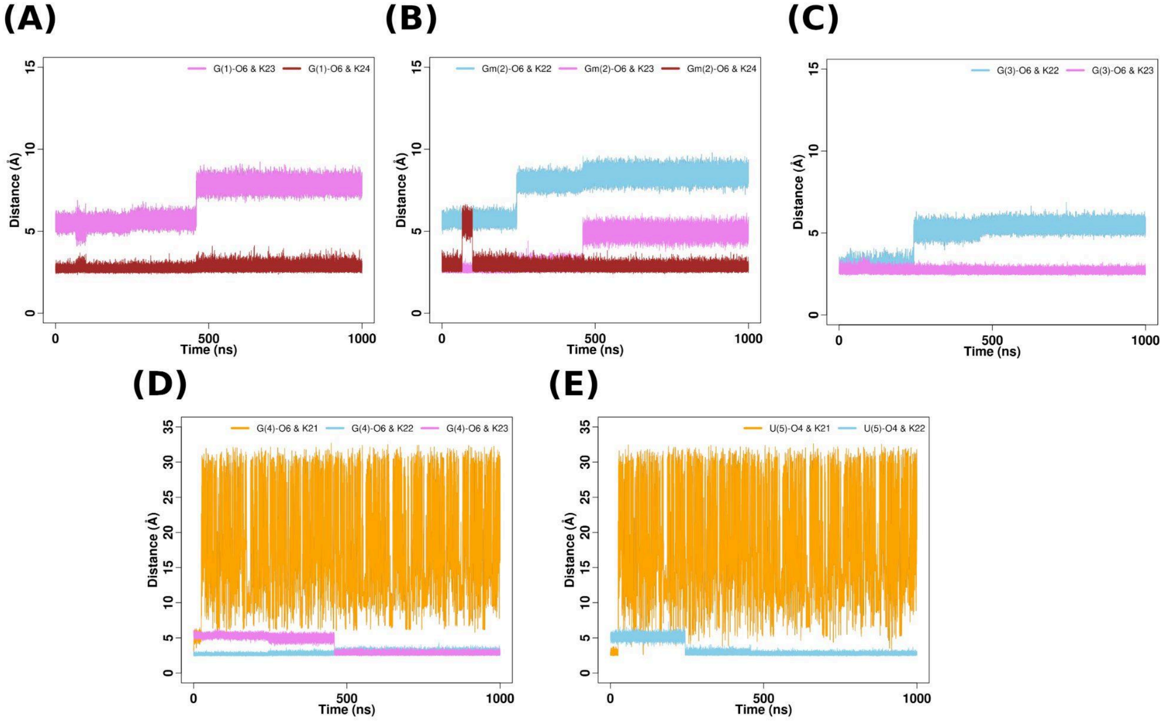


**Figure S27.** Time evolution of distances between (A) G(1)-O6 atom and K+ (23 and 24) (B) Gm(2)-O6 atom and K+ (22, 23 and 24) (C) G(3)-O6 atom and K+ (22 and 23) (D) G(4)-O6 atom and K+ (21, 22 and 23) and (E) U(5)-O4 atom and K+ (21 and 22) respectively for simulation set-3 for quadruplex containing Gm-tetrad at position 2.

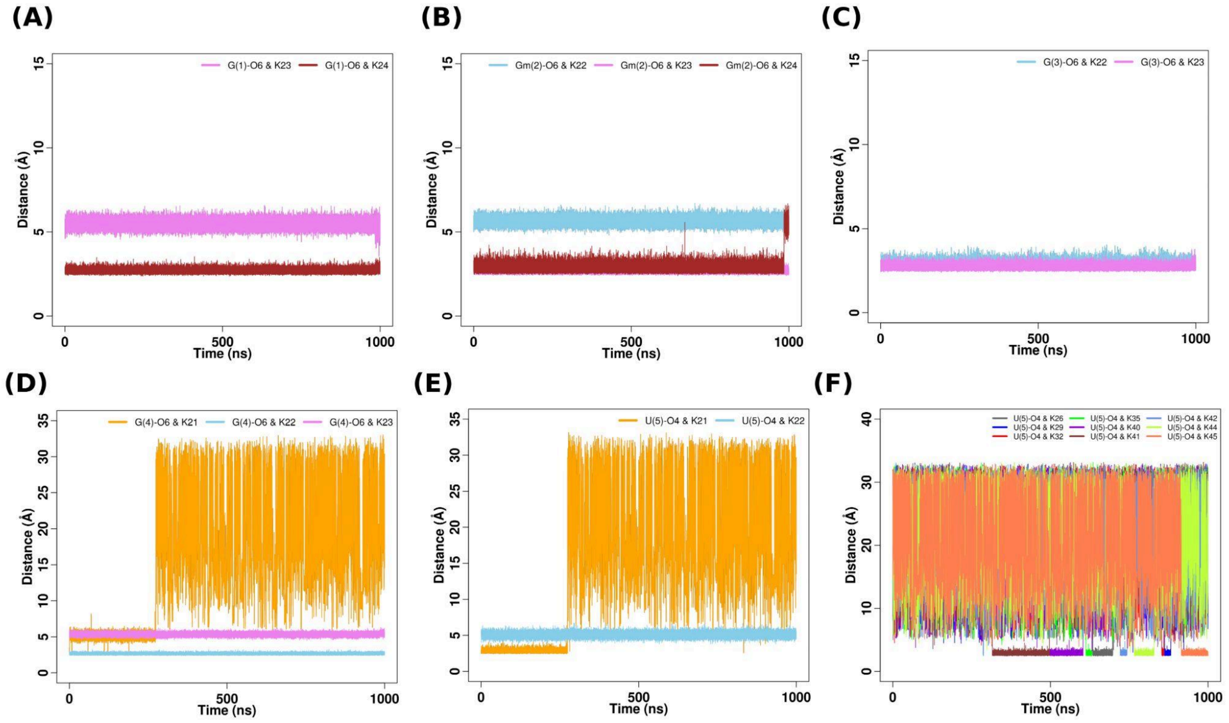


**Figure S28.** Time evolution of distances between (A) G(1)-O6 atom and K+ (23 and 24) (B) Gm(2)-O6 atom and K+ (22, 23 and 24) (C) G(3)-O6 atom and K+ (22 and 23) (D) G(4)-O6 atom and K+ (21, 22 and 23) and (E) U(5)-O4 atom and K+ (21 and 22) (F) U(5)-O4 atom and K+ (26, 29, 32, 35, 40, 41, 42, 44, 45) respectively for simulation set-4 for quadruplex containing Gm-tetrad at position 2.

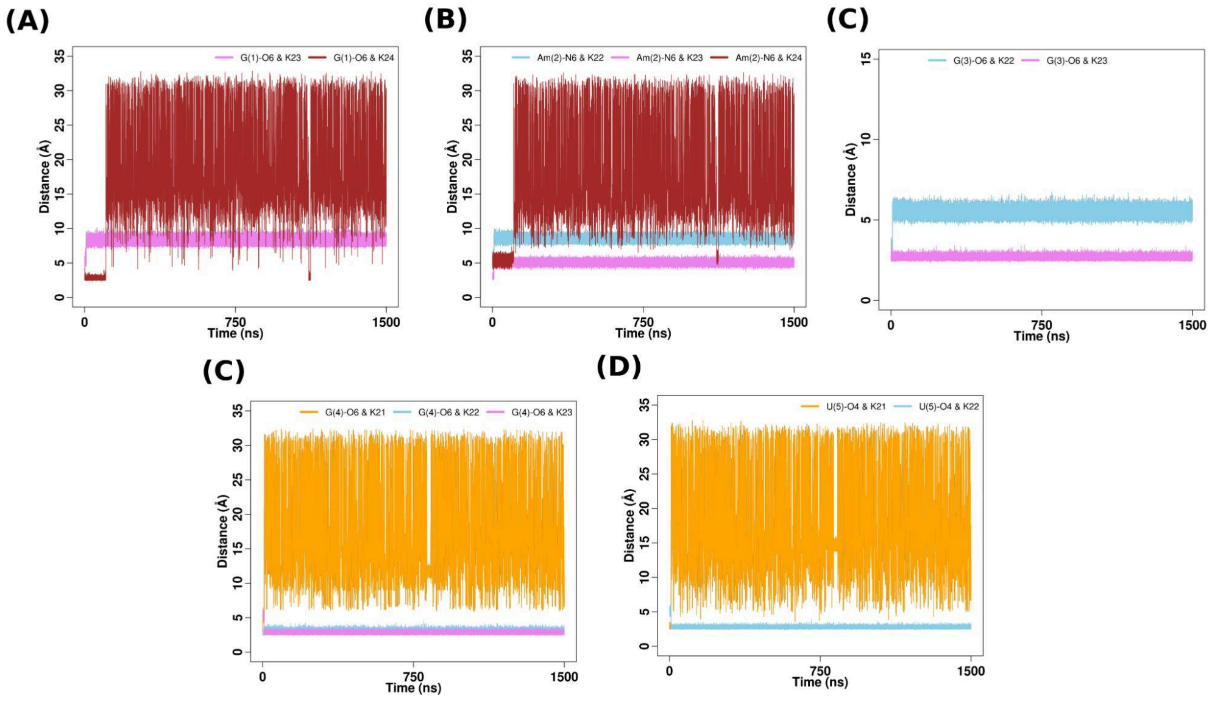


**Figure S29.** Time evolution of distances between (A) G(1)-O6 atom and K+ (23 and 24) (B) Am(2)-N6 atom and K+ (22, 23 and 24) (C) G(3)-O6 atom and K+ (21, 22 and 23) (D) G(4)-O6 atom and K+ (21 and 22) and (E) U(5)-O4 atom and K+ (21, 22 and 23) respectively for simulation set-1 for quadruplex containing Am-tetrad at position 2.

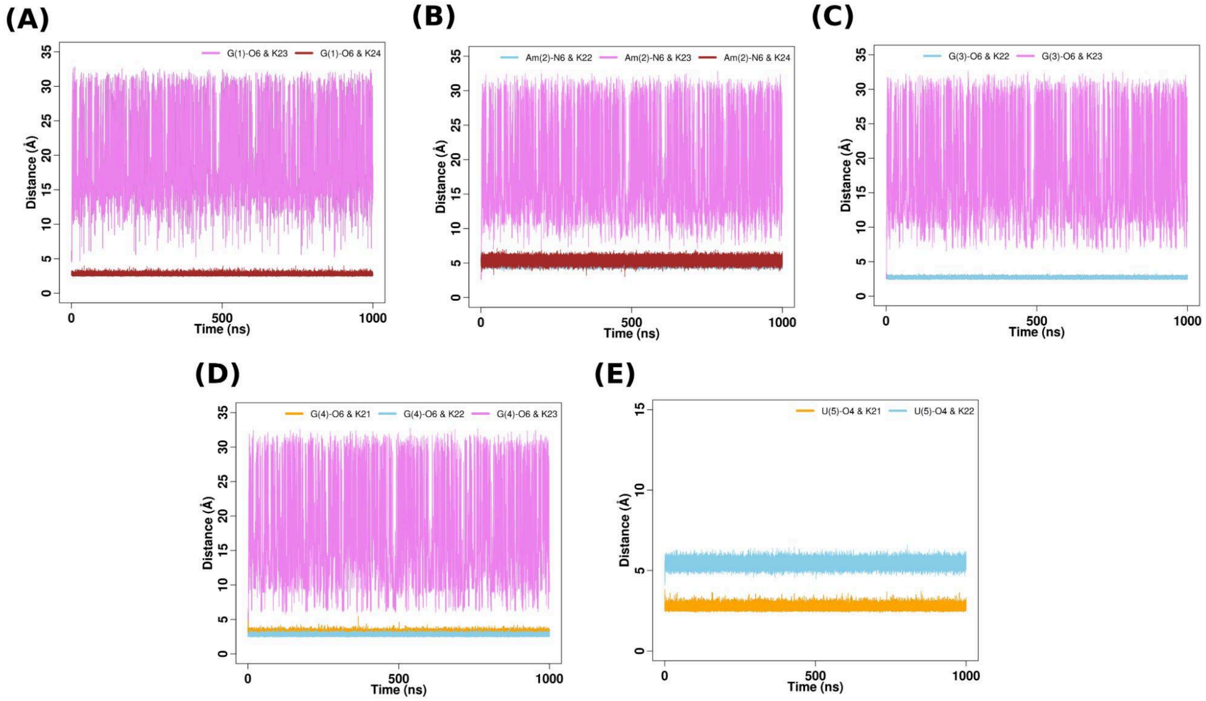


**Figure S30.** Time evolution of distances between (A) G(1)-O6 atom and K+ (23 and 24) (B) Am(2)-N6 atom and K+ (22, 23 and 24) (C) G(3)-O6 atom and K+ (21, 22 and 23) (D) G(4)-O6 atom and K+ (21 and 22) and (E) U(5)-O4 atom and K+ (21, 22 and 23) respectively for simulation set-2 for quadruplex containing Am-tetrad at position 2.

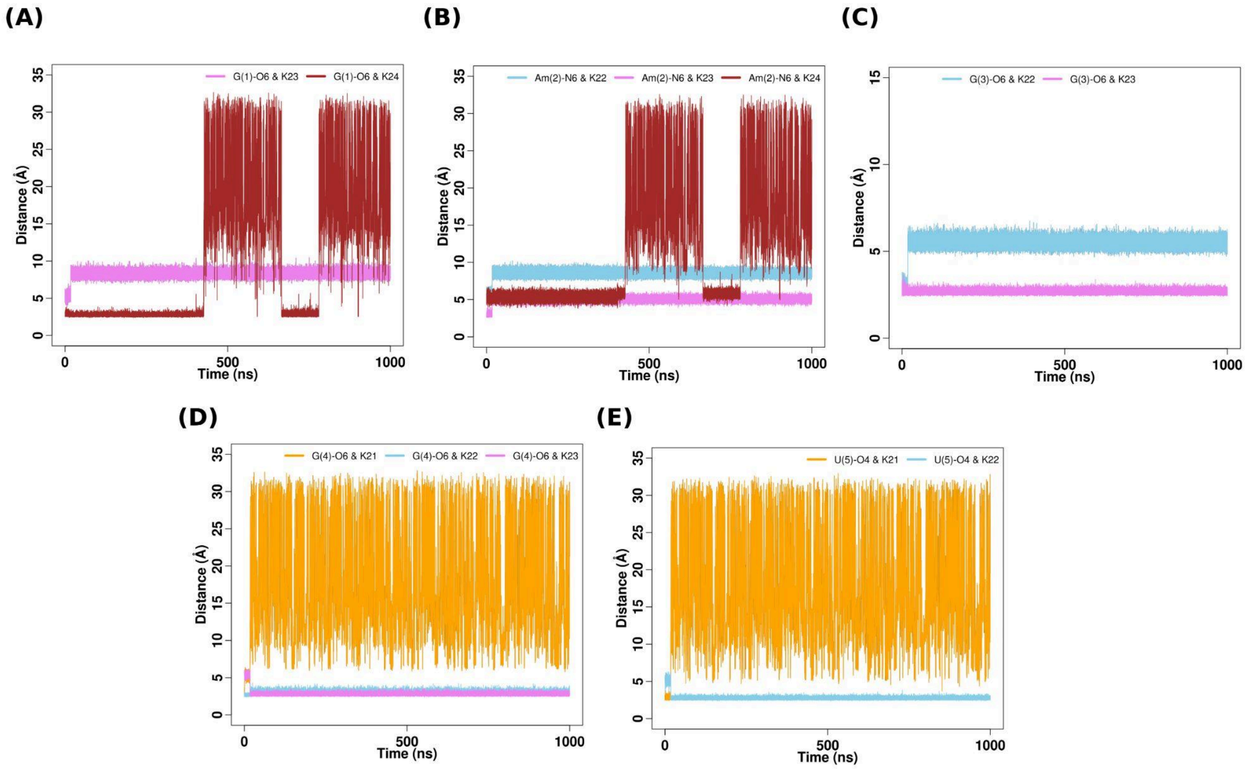


**Figure S31.** Time evolution of distances between (A) G(1)-O6 atom and K+ (23 and 24) (B) Am(2)-N6 atom and K+ (22, 23 and 24) (C) G(3)-O6 atom and K+ (21, 22 and 23) (D) G(4)-O6 atom and K+ (21 and 22) and (E) U(5)-O4 atom and K+ (21, 22 and 23) respectively for simulation set-3 for quadruplex containing Am-tetrad at position 2.

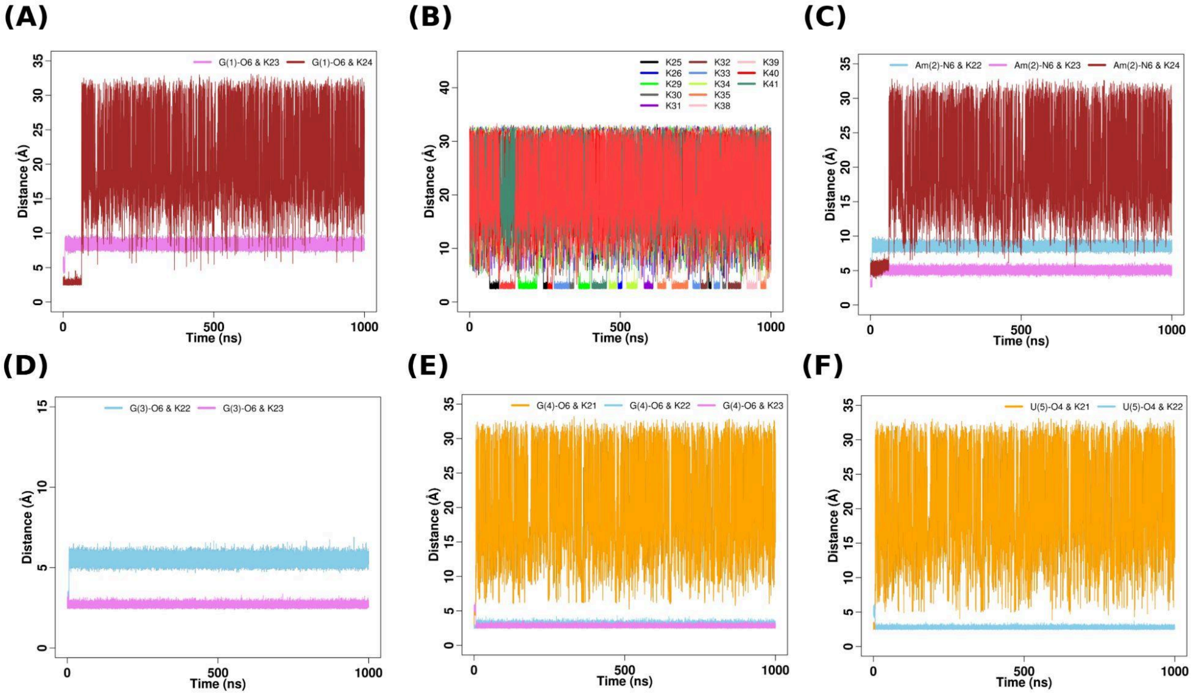


**Figure S32.** Time evolution of distances between (A) G(1)-O6 atom and K+ (23 and 24) (B) G(1)-O6 atom and K(25), K+(26,29-35,38-41) (C) A(2)-N6 atom and K+ (22, 23 and 24) (D) G(3)-O6 atom and K+ (21, 22 and 23) (E) G(4)-O6 atom and K+ (21 and 22) and (F) U(5)-O4 atom and K+ (21, 22 and 23) respectively for simulation set-4 for quadruplex containing Am-tetrad at position 2.

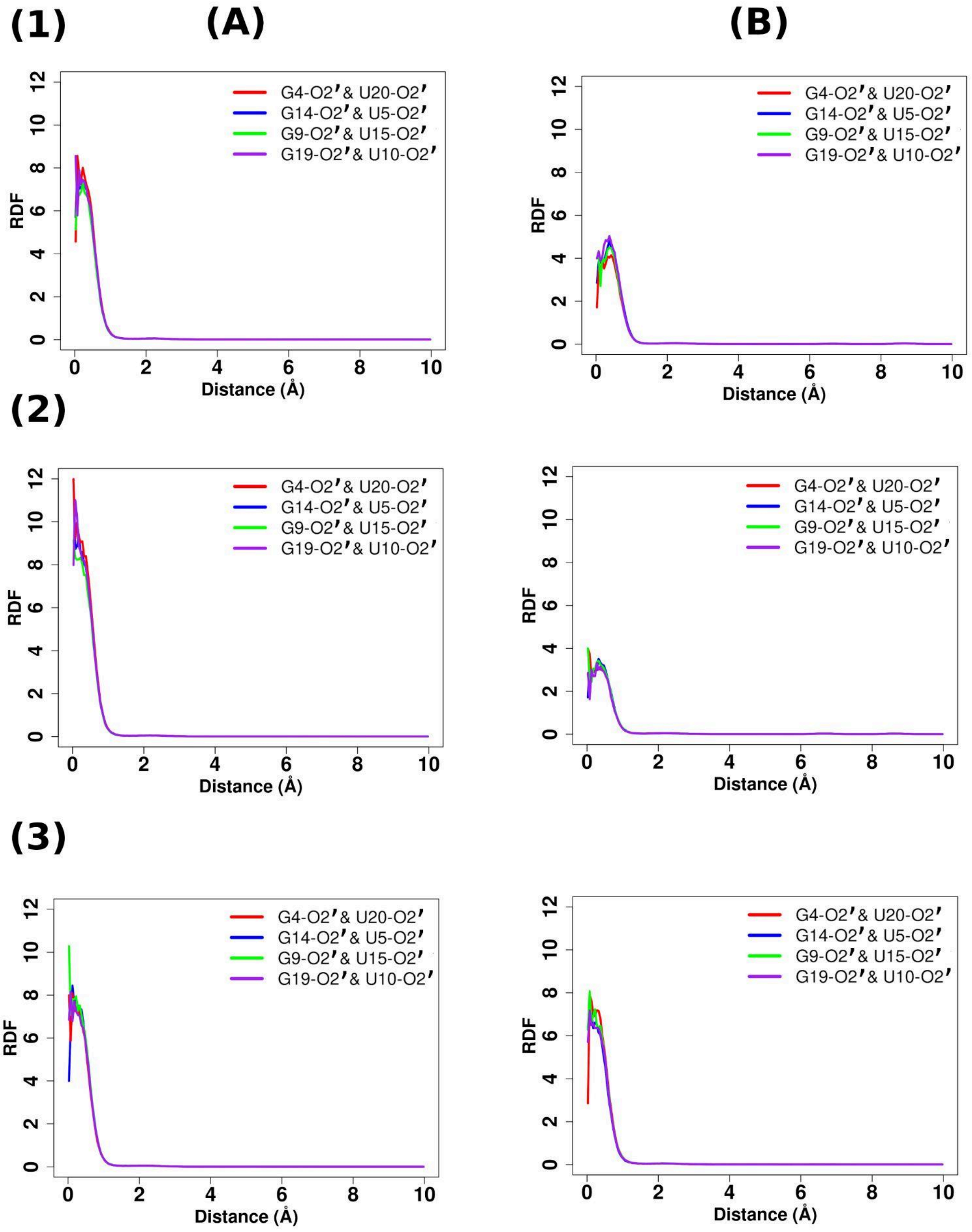


**Figure S33.** RDF of $K^+$ ions around the geometric center of G-O2′ and U-O2′(for G4-U20, G14-U19, G9-U15, G19-U10 for the fourth set of simulations in quadruplexes with (A) unmethylated ((1) Quad-I4 (2) Quad-G4 and (3) Quad-A4) and (B) 2′-O-methylated tetrads ((1) Quad-Im4 (2) Quad-Gm4 and (3) Quad-Am4) at position 2.

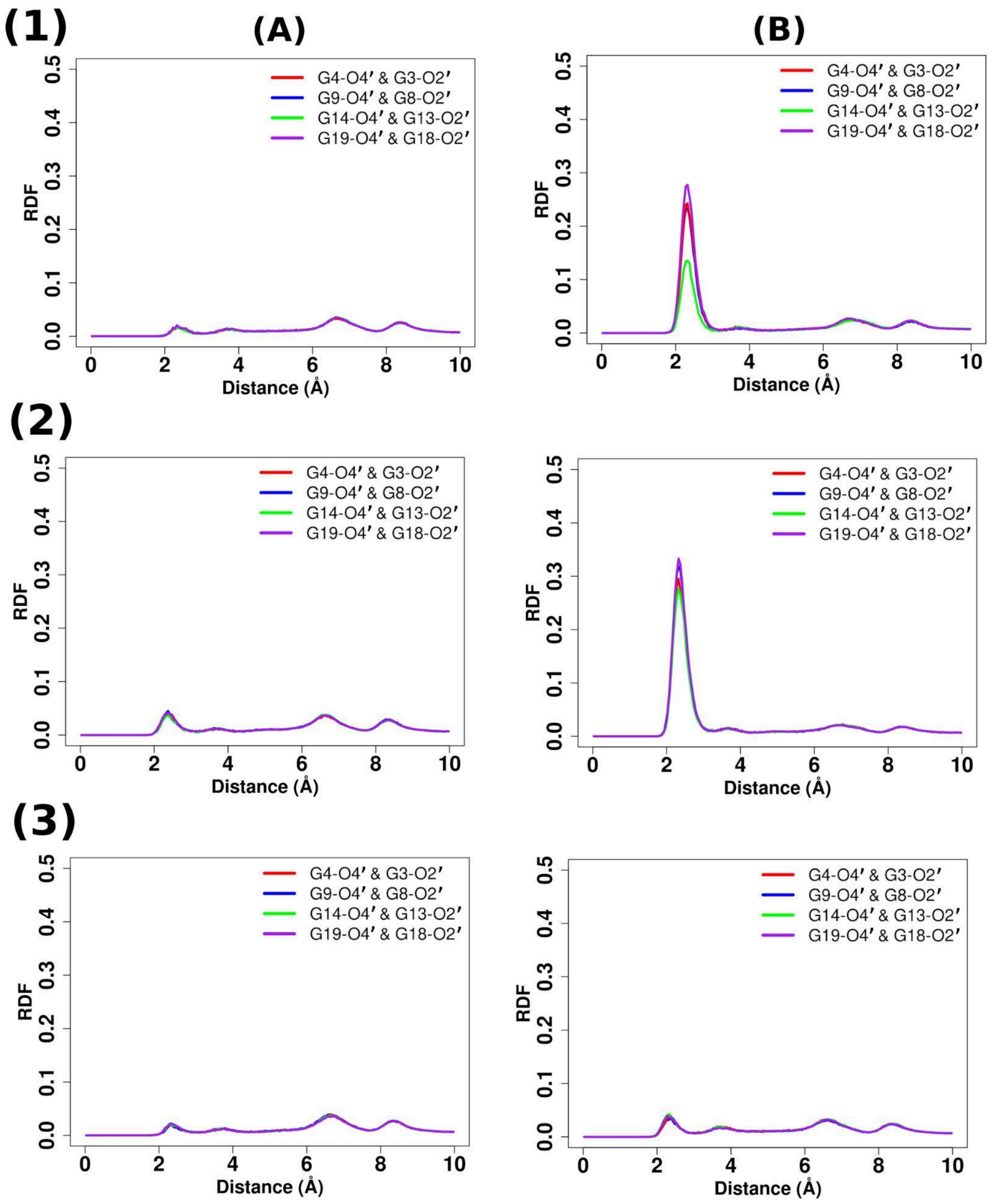


**Figure S34.** RDF of $K^+$ ions around the geometric center of Gn-O2′ and Gn-1-O2′ (for G4-G3, G9-G8, G14-G13, G19-G18 for the fourth set of simulations in quadruplexes with (A) unmethylated ((1) Quad-I4 (2) Quad-G4 and (3) Quad-A4) and (B) 2′-O-methylated tetrads ((1) Quad-Im4 (2) Quad-Gm4 and (3) Quad-Am4) at position 2.

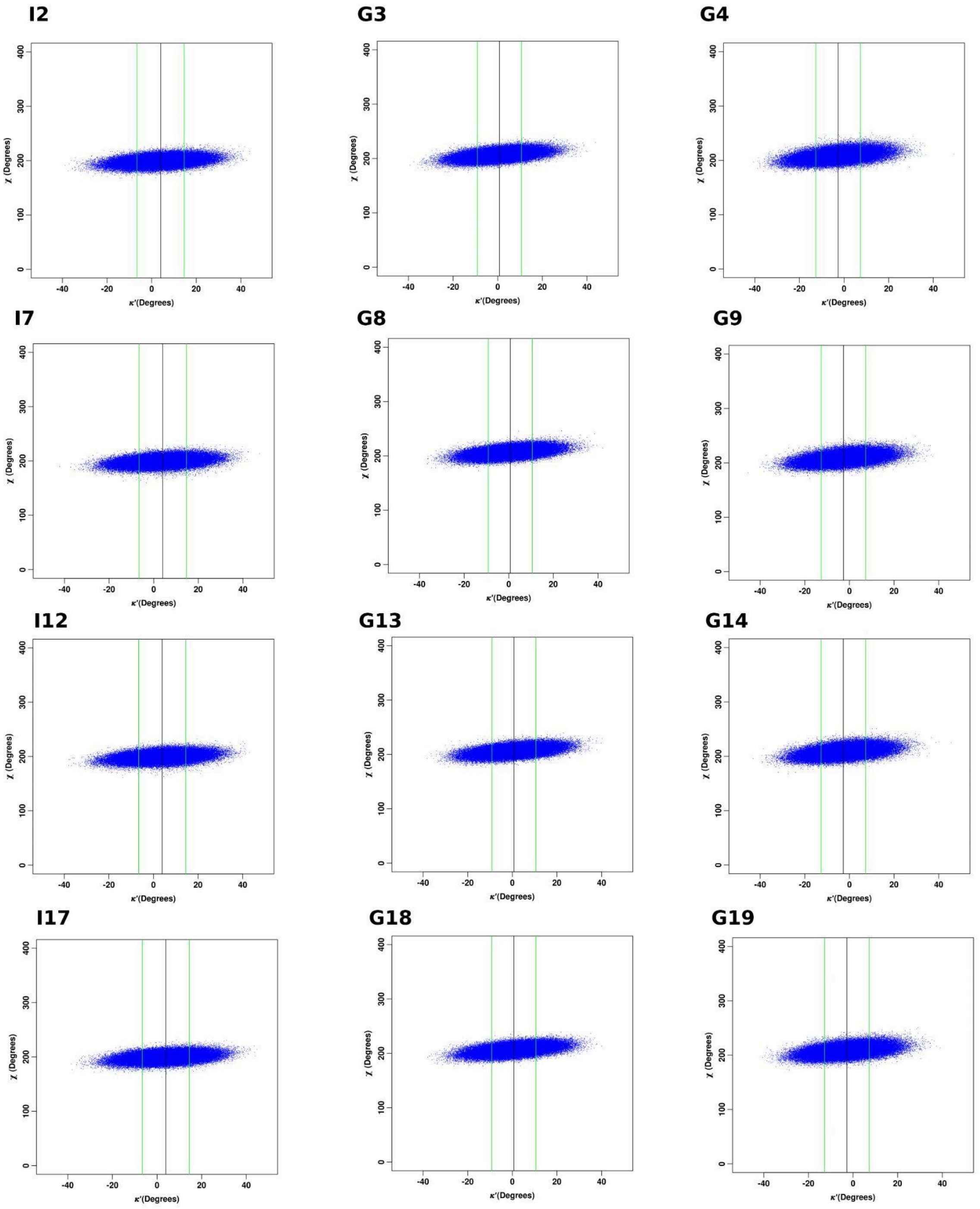


**Figure S35.** 2D-scatter plots of $\chi$ vs $\kappa$' dihedrals for the residues in the tetrads at positions 2,3, and 4 for the monomeric quadruplex (Quad-I4) containing I(2)-tetrad corresponding to the fourth simulation set.

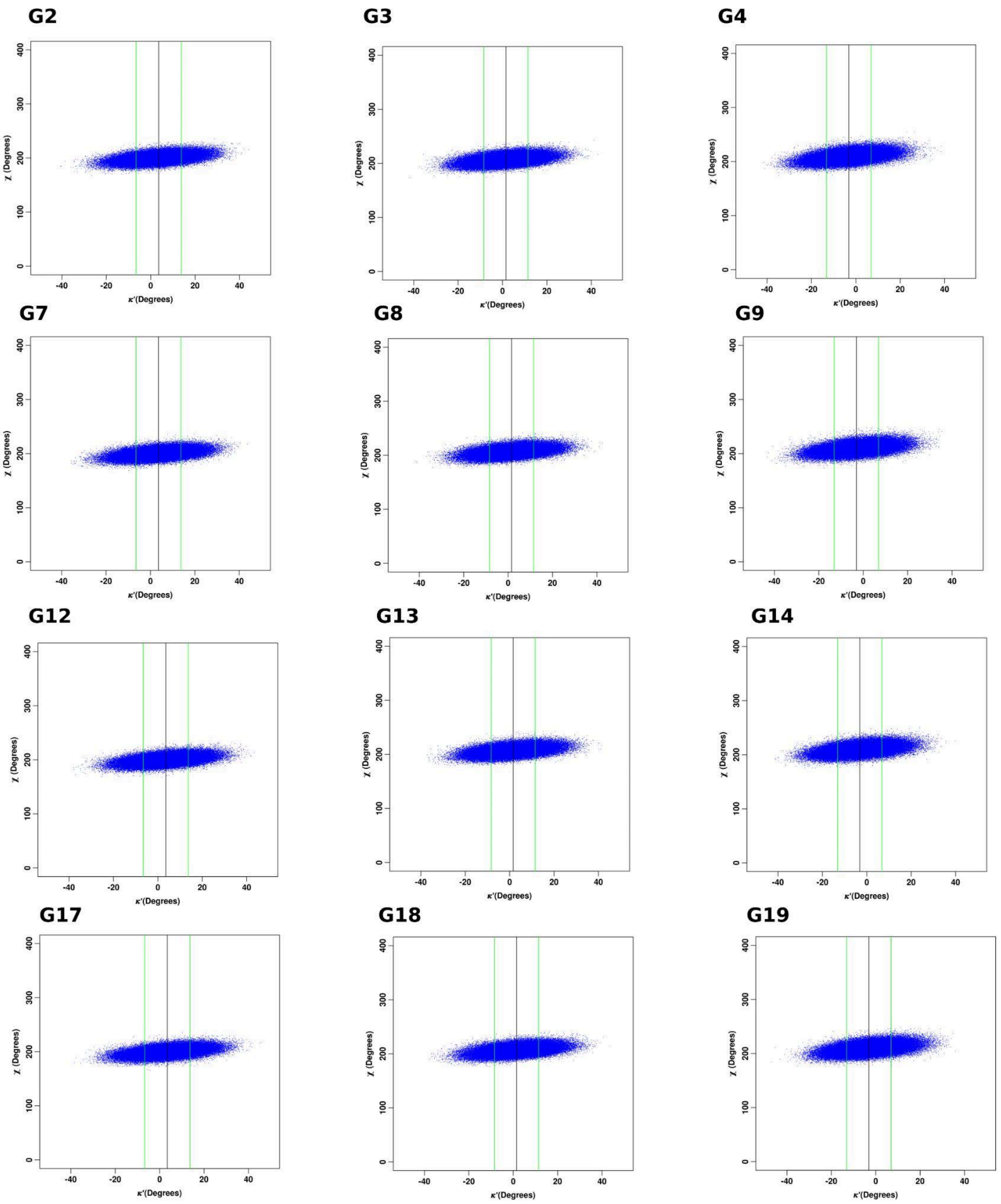


**Figure S36.** 2D-scatter plots of $\chi$ vs $\kappa$’ dihedrals for the residues in the tetrads at positions 2,3, and 4 for the monomeric quadruplex (Quad-G4) containing G(2)-tetrad corresponding to the fourth simulation set.

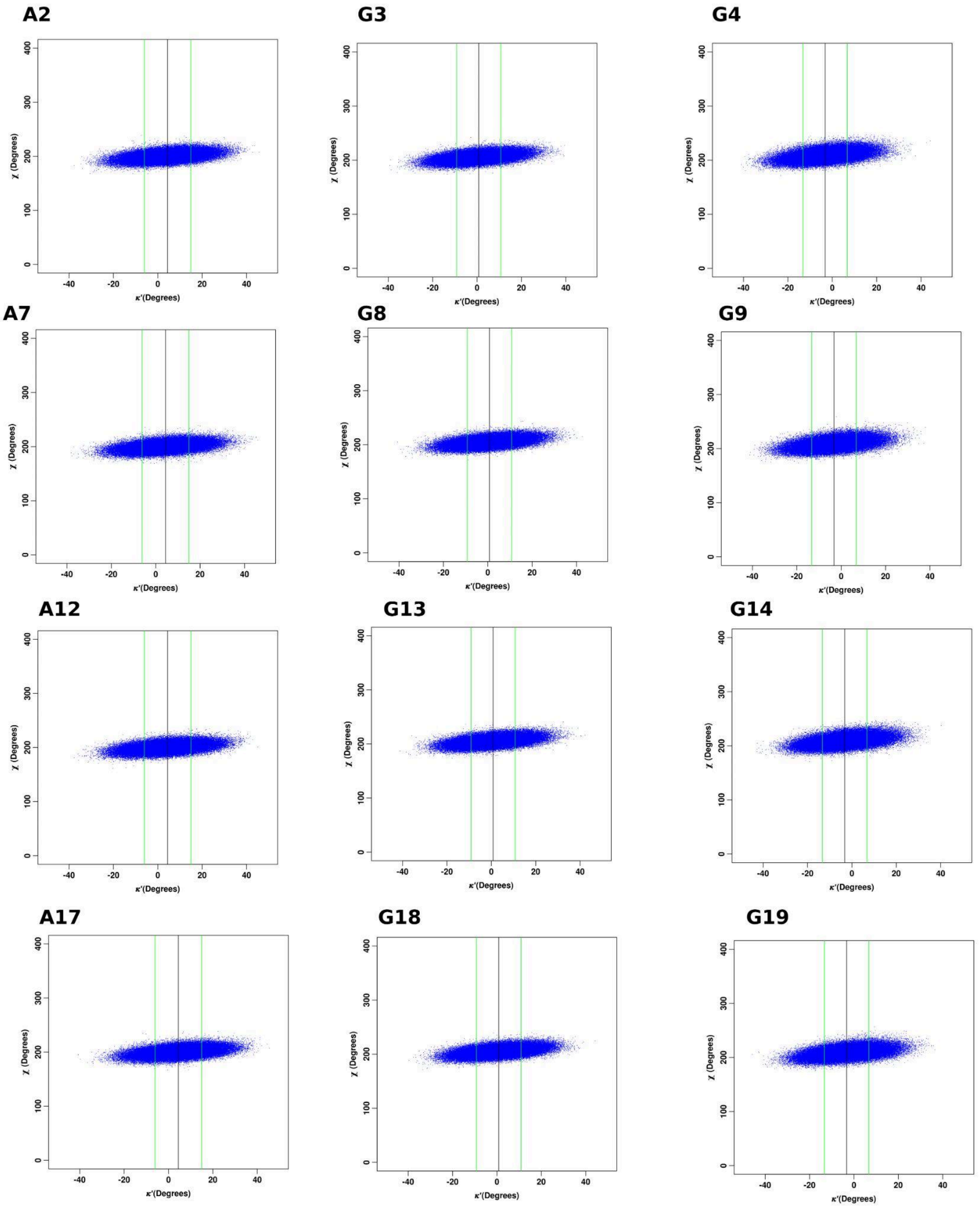


**Figure S37.** 2D-scatter plots of $\boldsymbol{\chi}$ vs $\boldsymbol{\kappa}$’ dihedrals for the residues in the tetrads at positions 2,3, and 4 for the monomeric quadruplex (Quad-A4) containing A(2)-tetrad corresponding to the fourth simulation set.

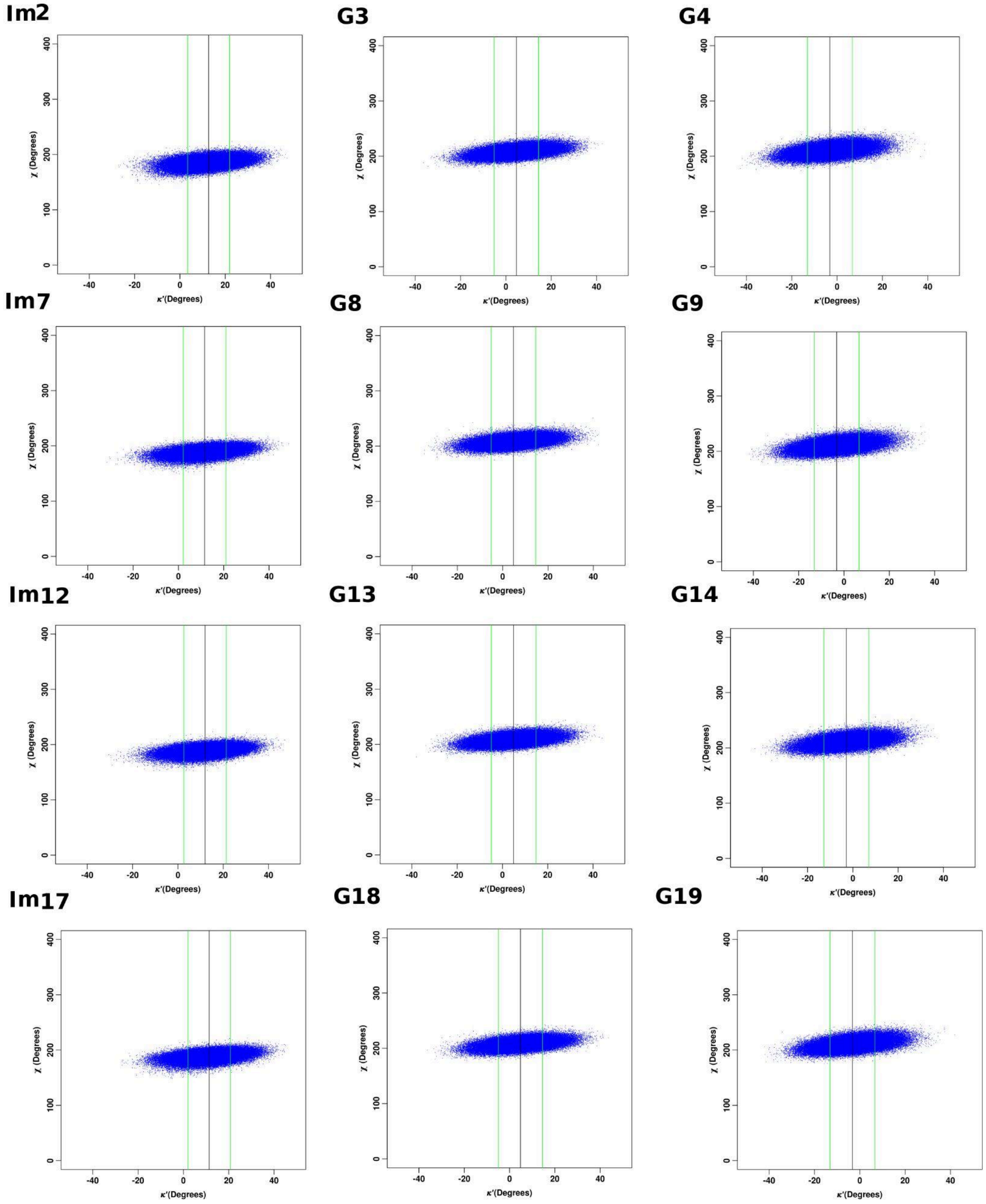


**Figure S38.** 2D-scatter plots of $\chi$ vs $\kappa$’ dihedrals for the residues in the tetrads at positions 2,3, and 4 for the monomeric quadruplex (Quad-Im4) containing Im(2)-tetrad corresponding to the fourth simulation set.

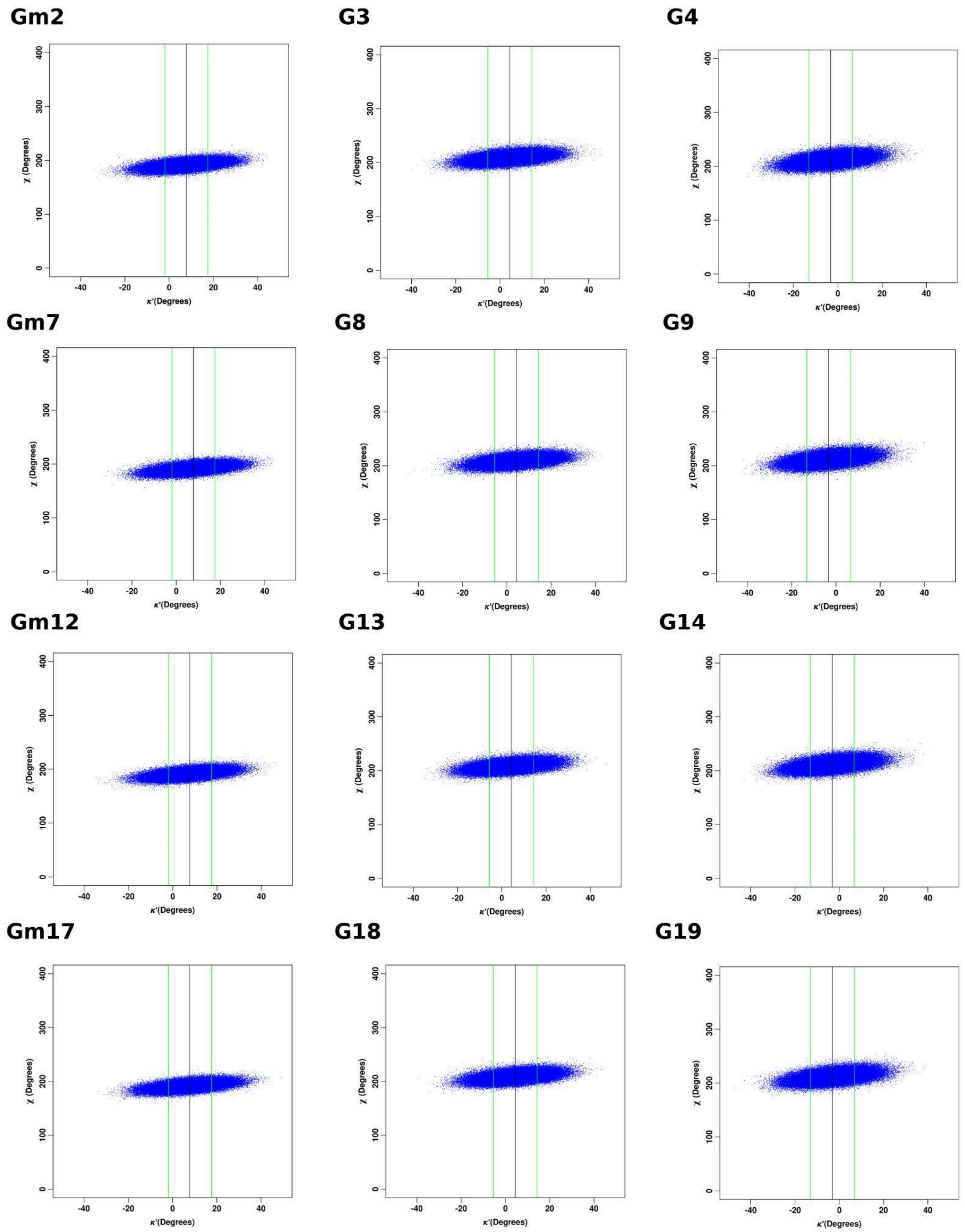


**Figure S39.** 2D-scatter plots of **χ** vs **κ**' dihedrals for the residues in the tetrads at positions 2,3, and 4 for the monomeric quadruplex (Quad-Gm4) containing Gm(2)-tetrad corresponding to the fourth simulation set.

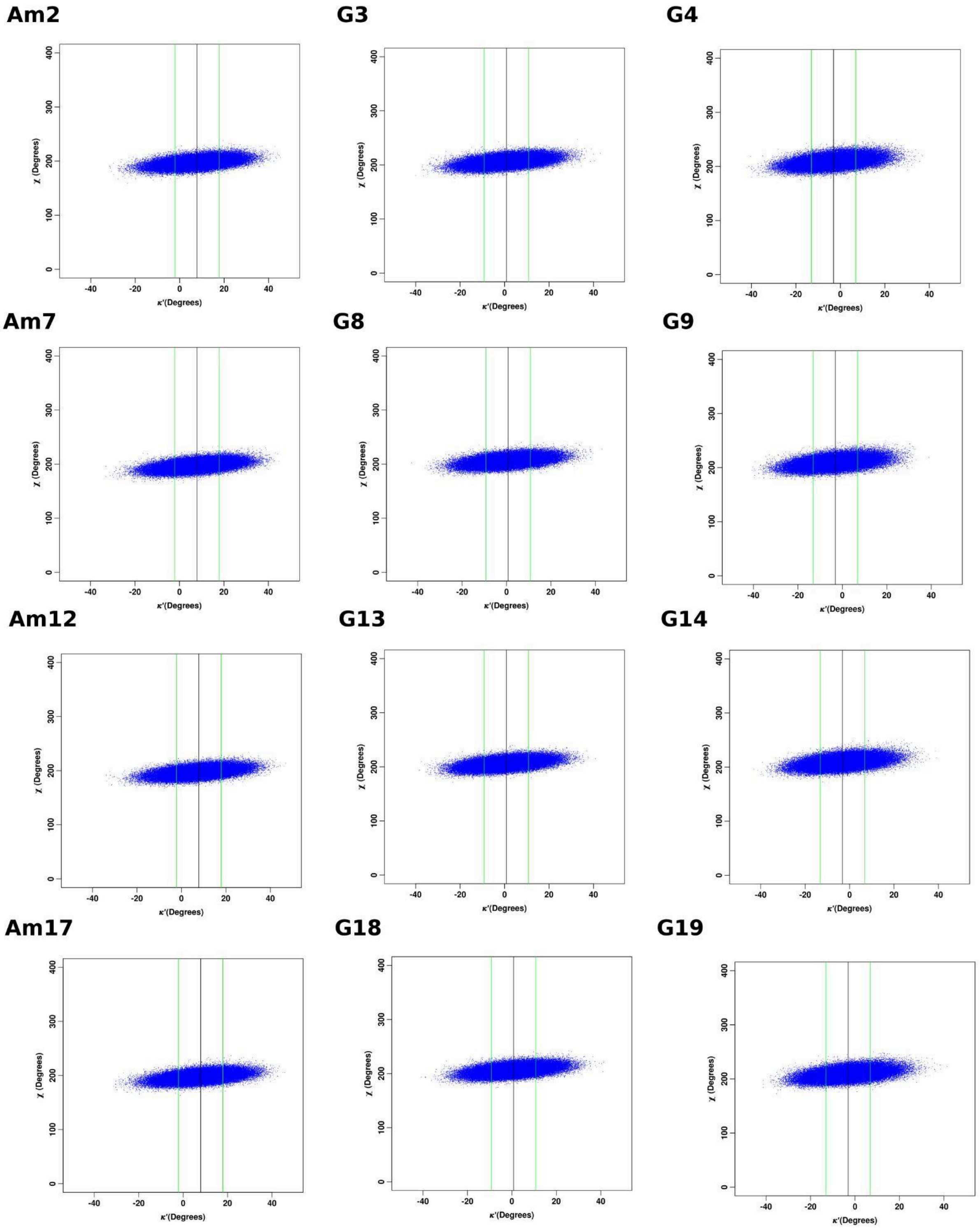


**Figure S40.** 2D-scatter plots of $\boldsymbol{\chi}$ vs $\boldsymbol{\kappa}$' dihedrals for the residues in the tetrads at positions 2,3, and 4 for the monomeric quadruplex (Quad-Am4) containing Am(2)-tetrad corresponding to the fourth simulation set.

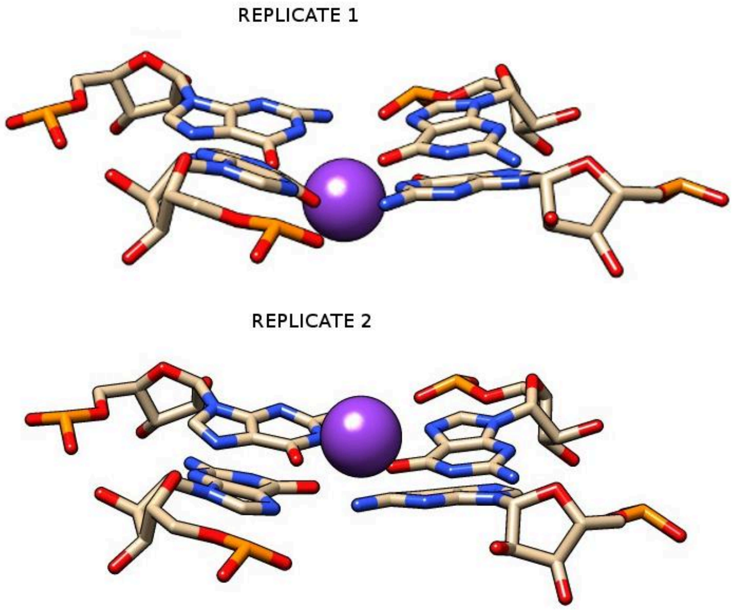


**Figure S41.** Snapshot of G(2)-tetrad within Quad-G4. The different positions of the channel K23 ion in the G4-channel in the first and second simulation sets resulting in different N9-pyramidalization of the G(2) tetrad residues.

*Simulation set-1*

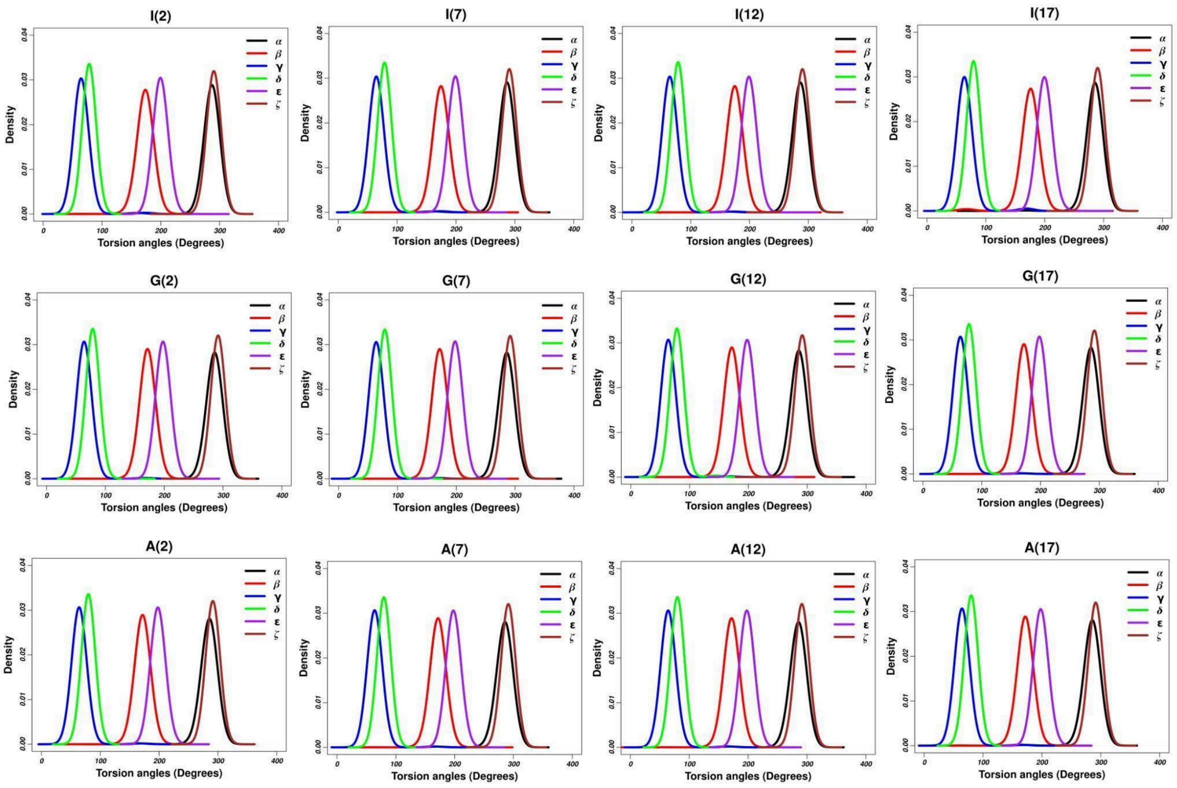


*Simulation set-2*

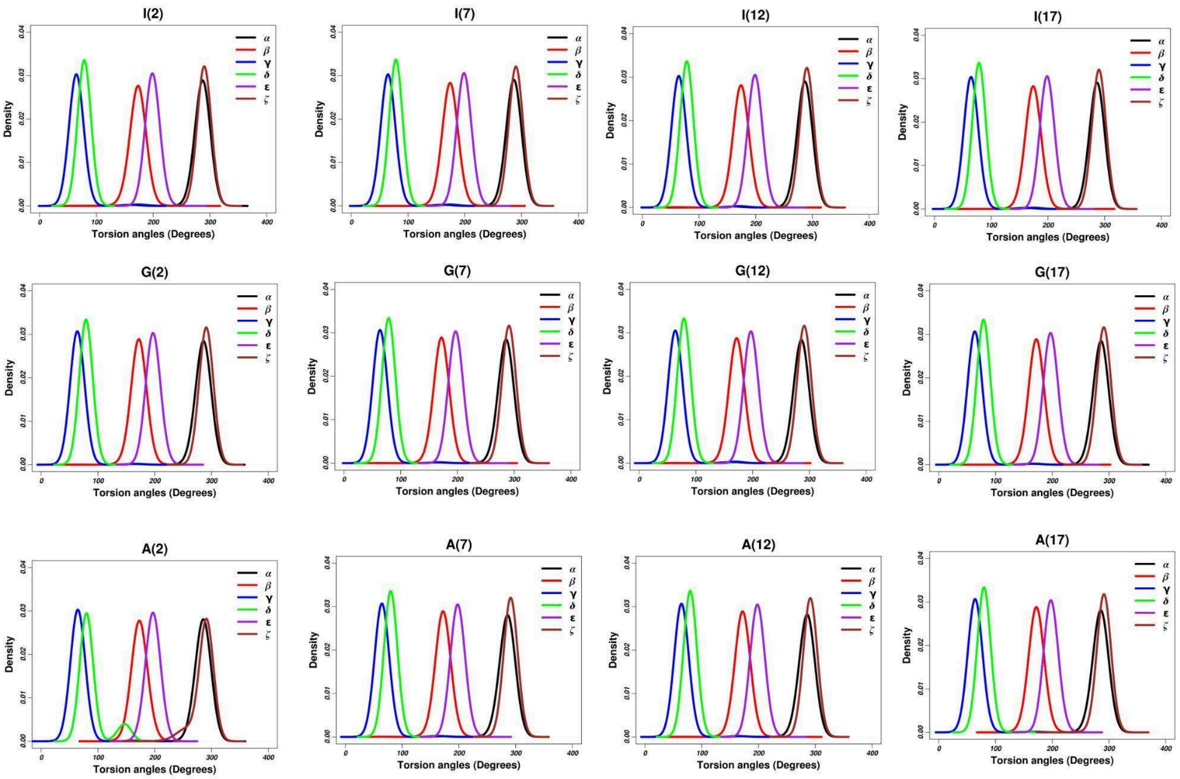

*Simulation set-3*

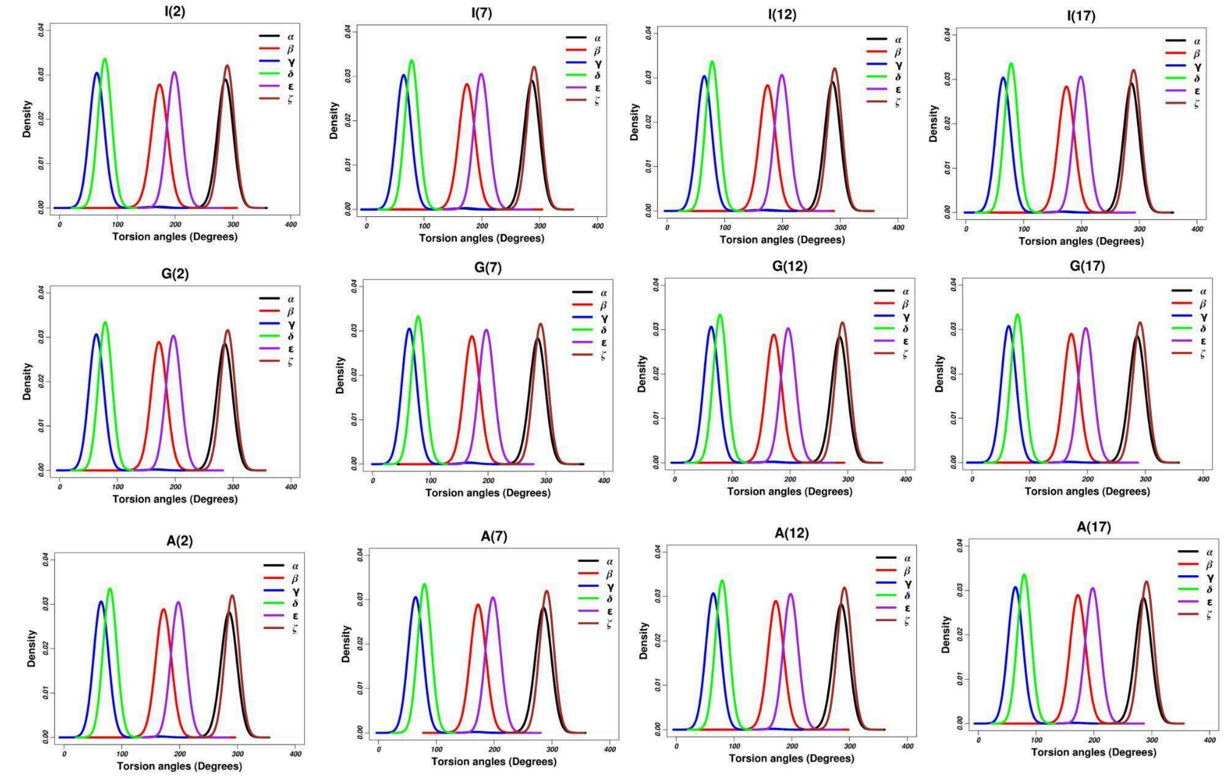


*Simulation set-4*

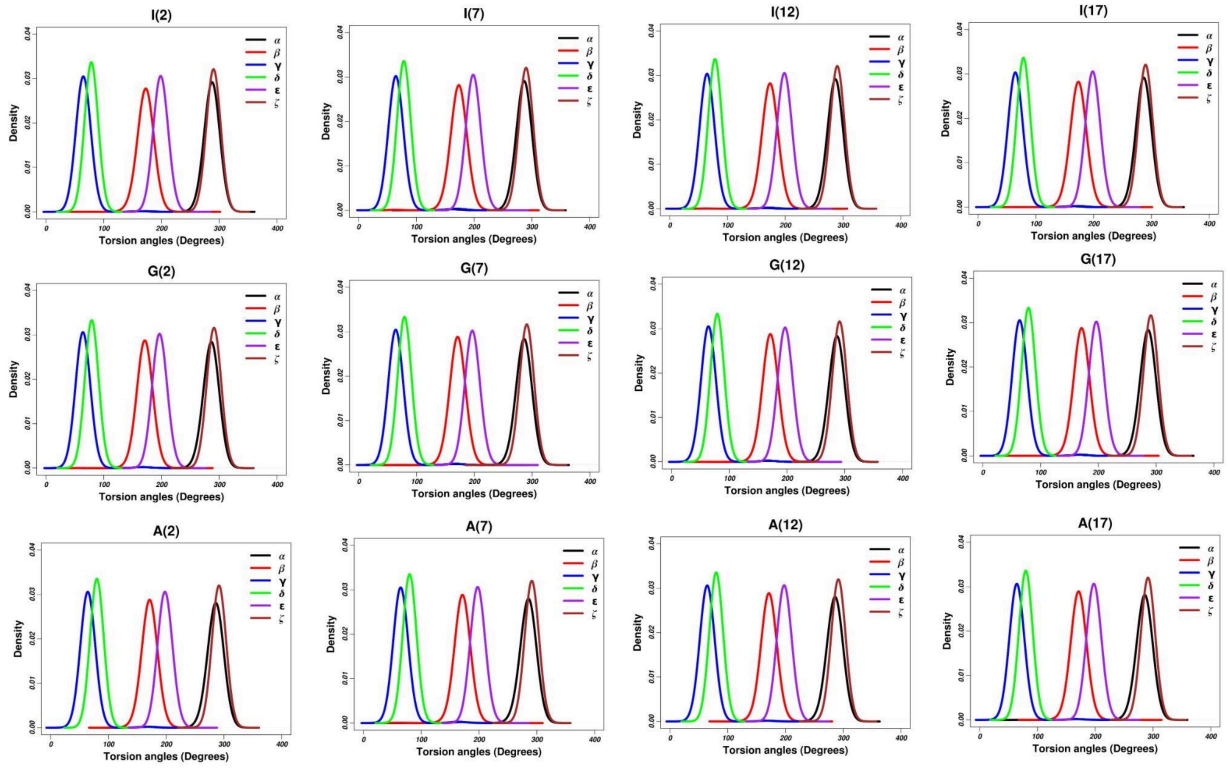

**Figure S42.** Distribution of the six backbone dihedral angles for the residues in the I-,G- and A-tetrads at position 2 for the monomeric quadruplexes (Quad-I4, Quad-G4 and Quad-A4 respectively) corresponding to the four simulation sets.

***Simulation set-1***

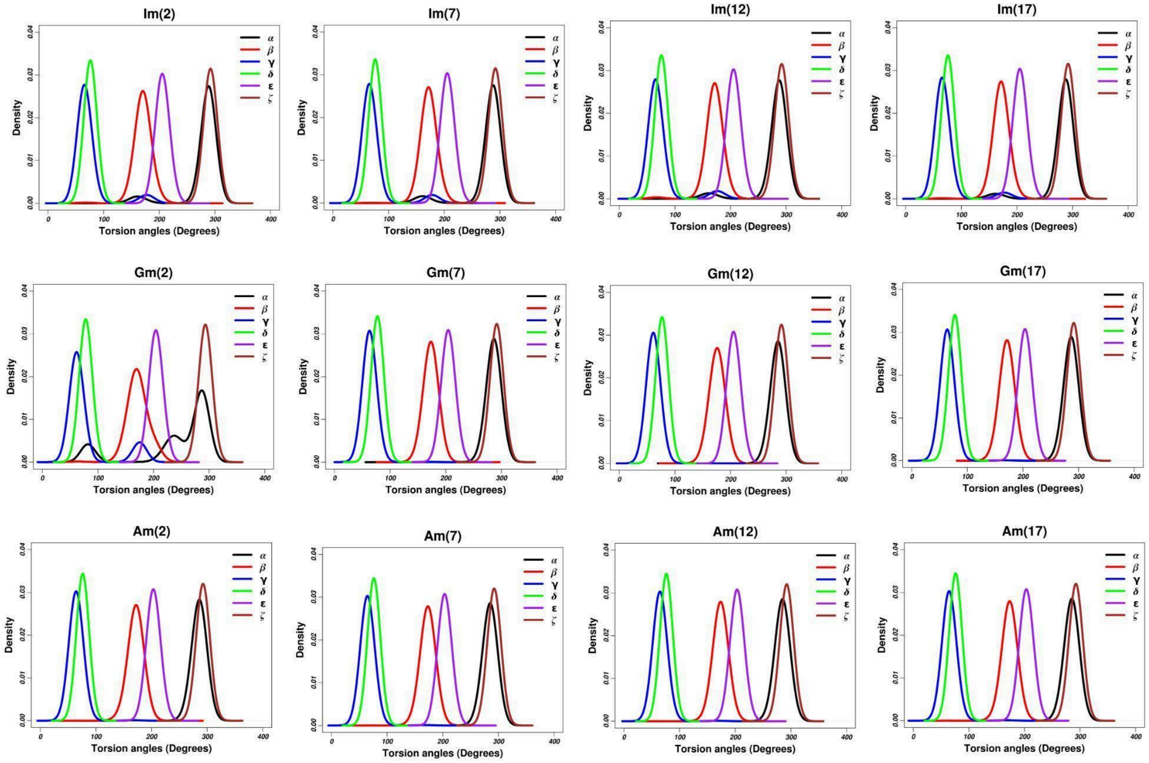


***Simulation set-2***

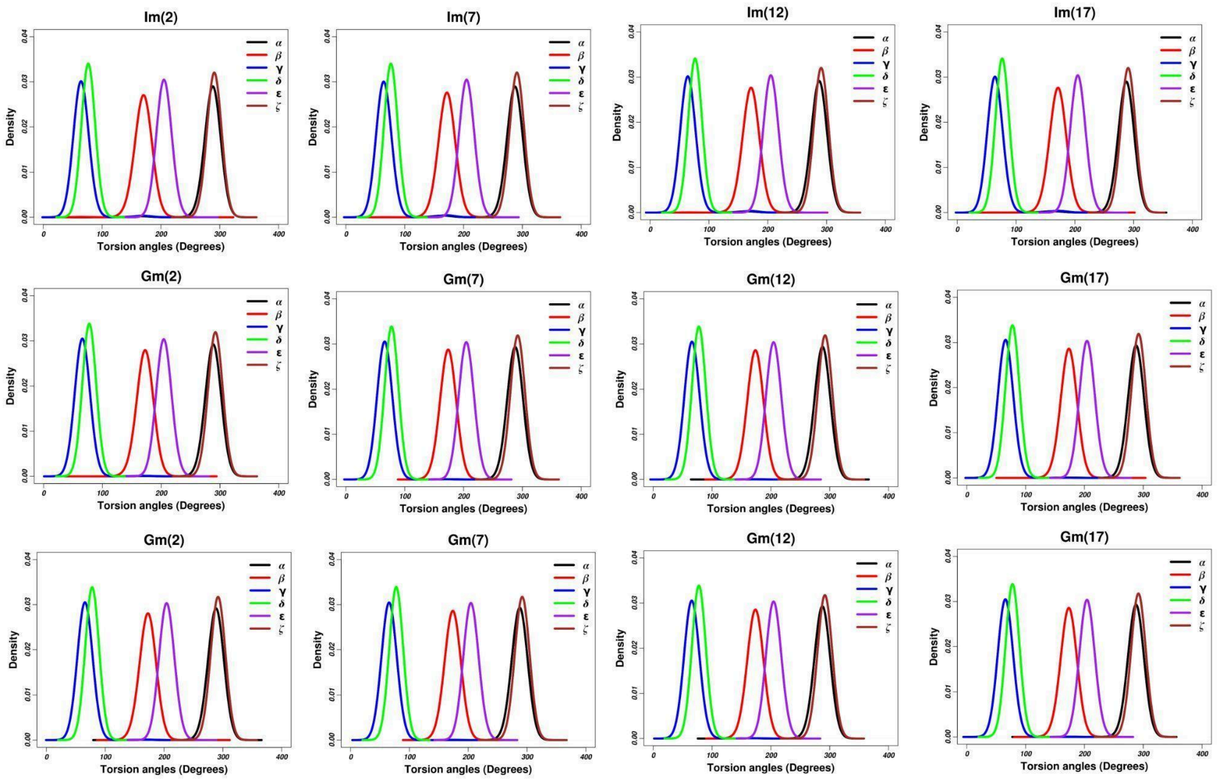


***Simulation set-3***

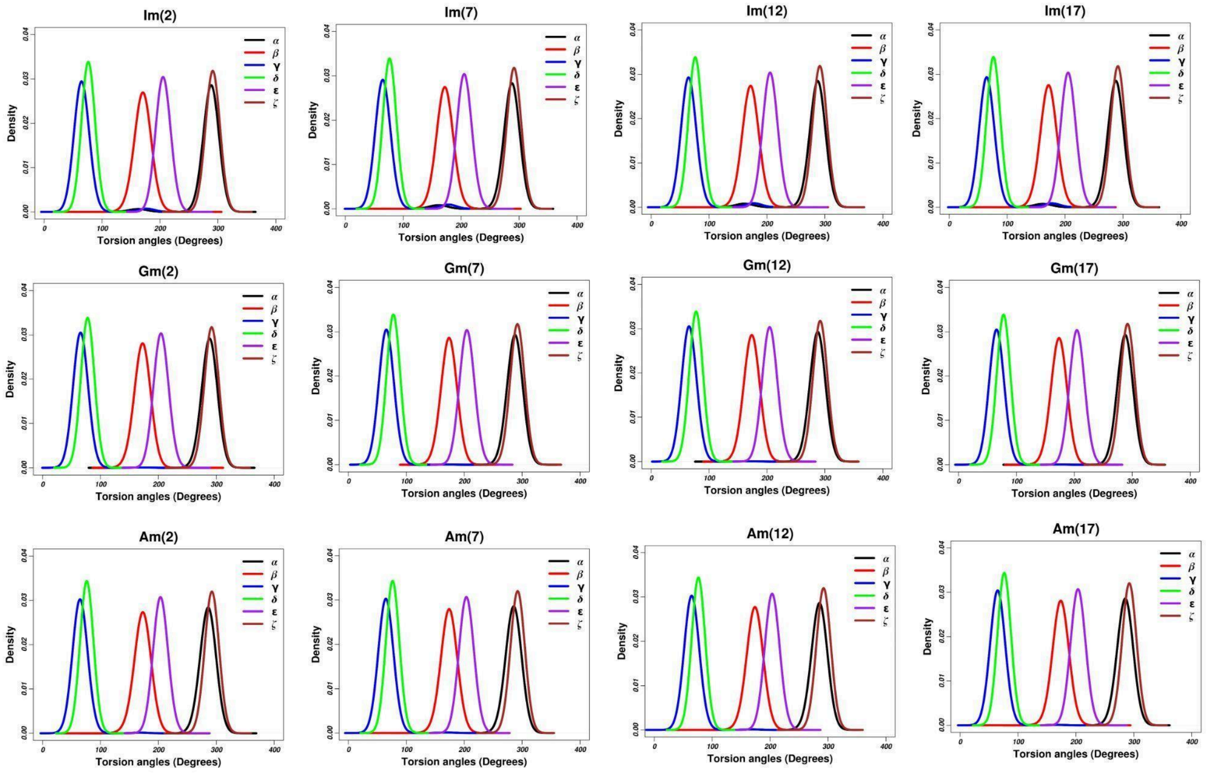


***Simulation set-4***

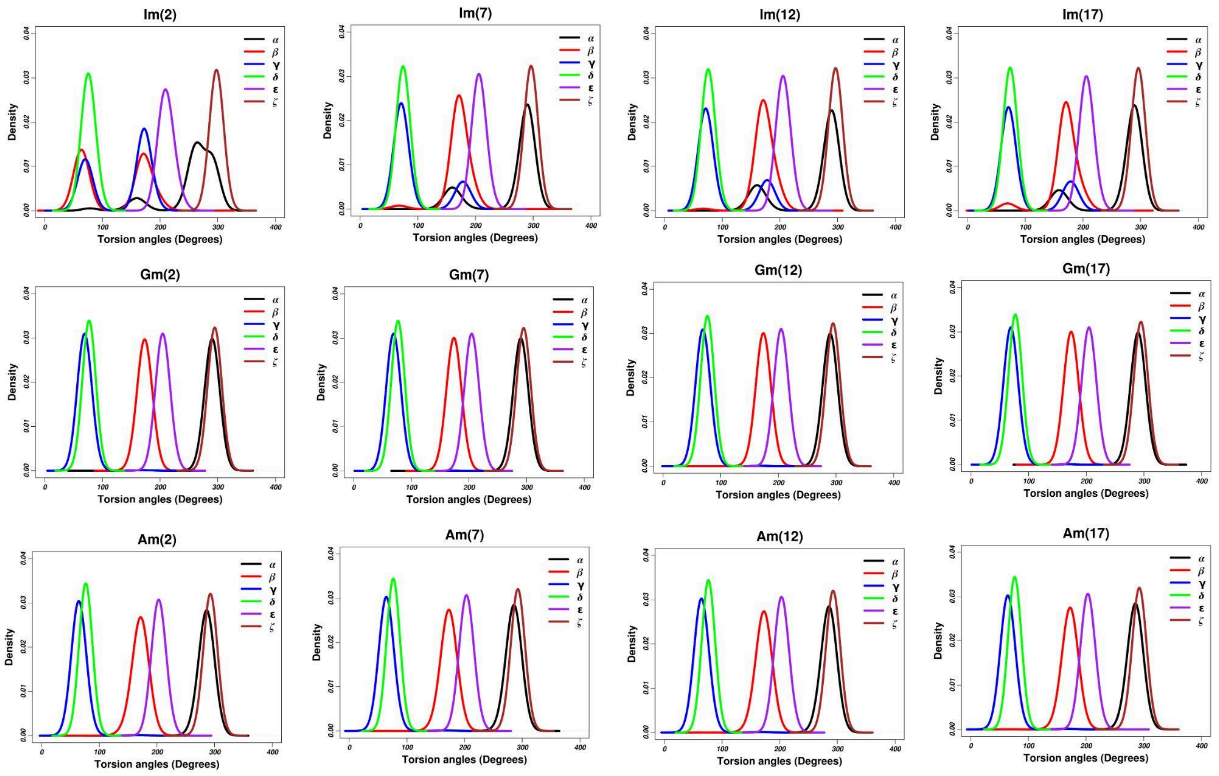


**Figure S43.** Distribution of the six backbone dihedral angles for the residues in the Im-, Gm- and Am- tetrads at position 2 for the monomeric quadruplexes (Quad-Im4, Quad-Gm4 and Quad-A4 respectively) corresponding to the four simulation sets.

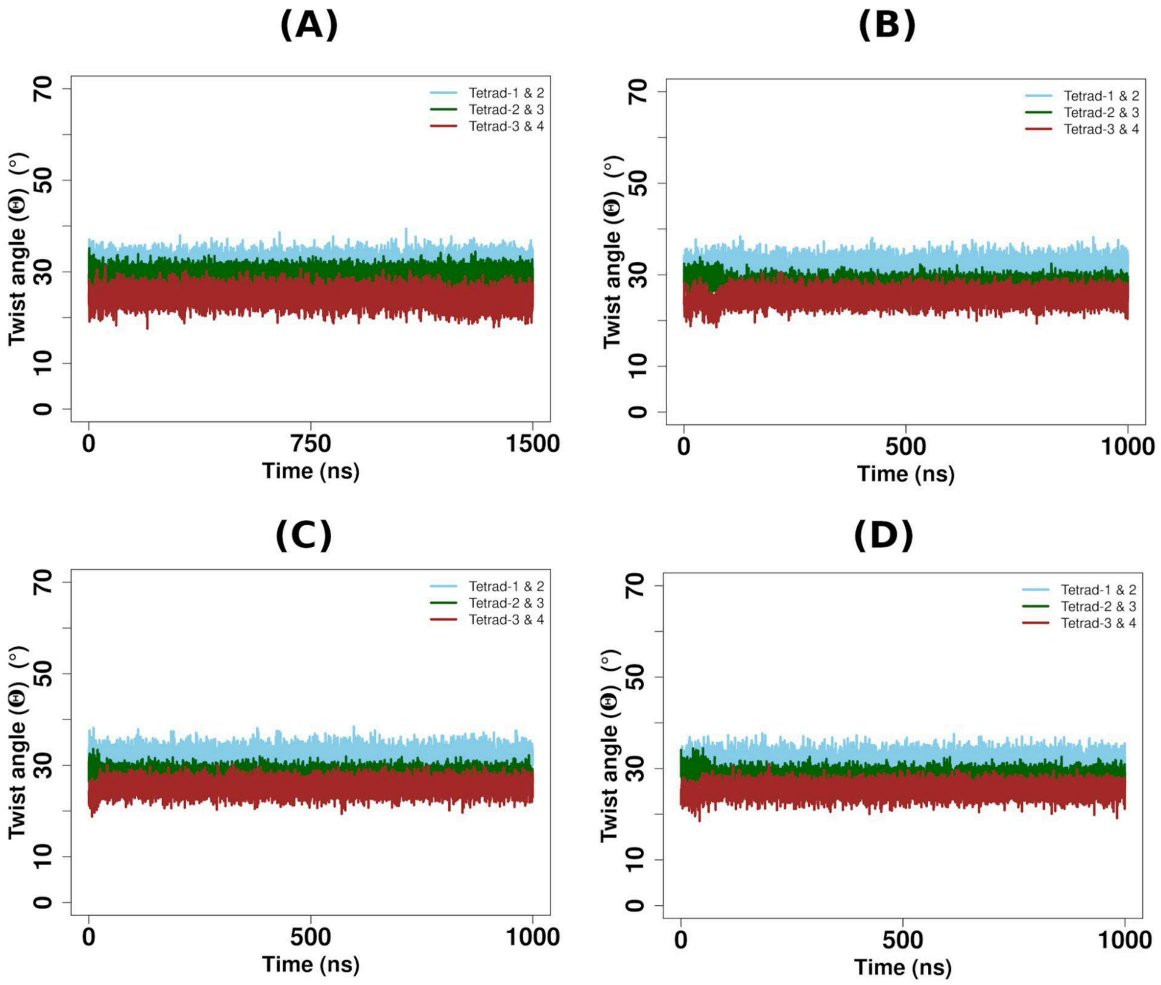


**Figure S44.** Time evolution of the twist angles (**Θ**) between the tetrads-1 and 2; 2 and 3; and 3 and 4 respectively for the monomeric quadruplex (GQ) containing G-tetrad at position 2 (Quad-G4) corresponding to the four simulation sets (A-D).

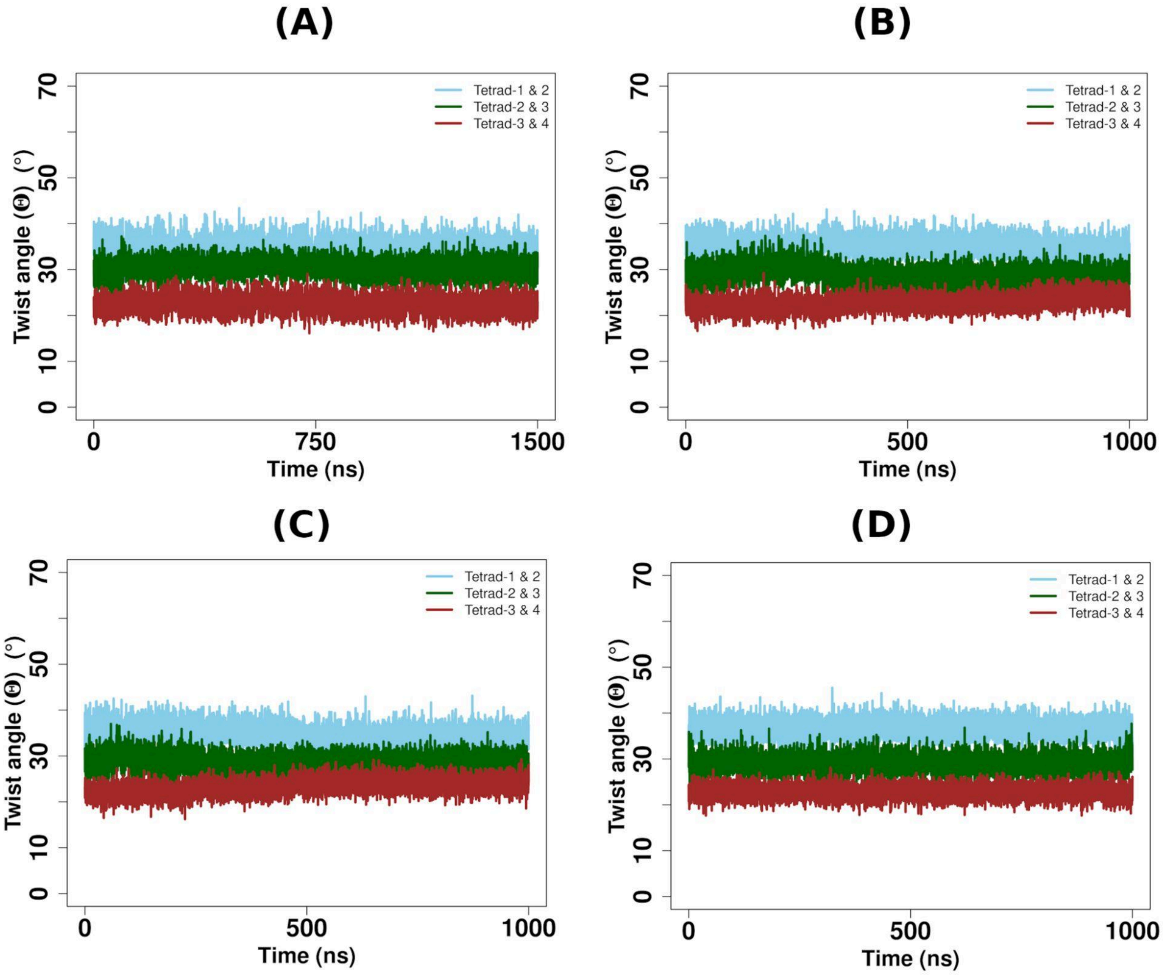


**Figure S45.** Time evolution of the twist angles (**Θ**) between the tetrads-1 and 2; 2 and 3; and 3 and 4 respectively for the monomeric quadruplex (GQ) containing Gm-tetrad at position 2 (Quad-Gm4) corresponding to the four simulation sets (A-D).

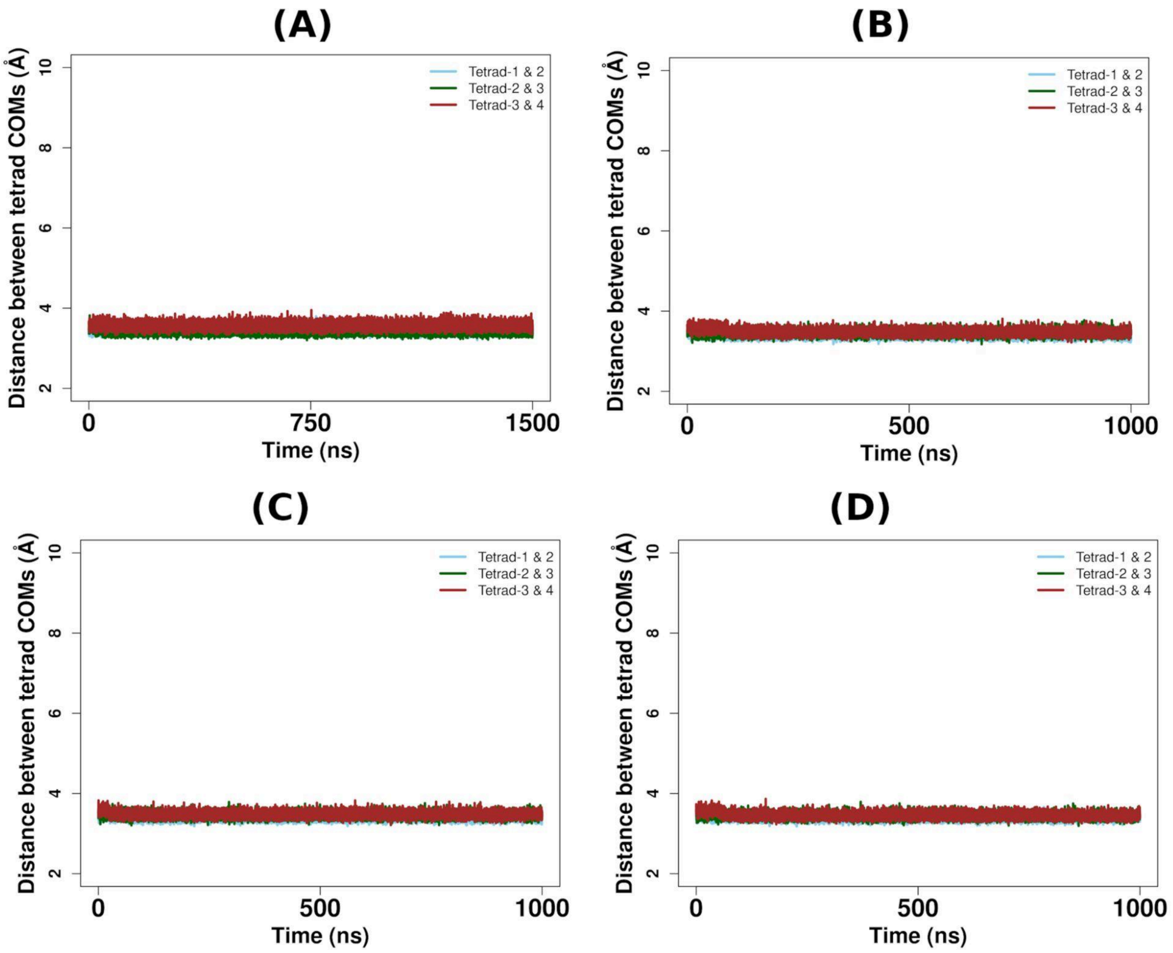


**Figure S46.** Time evolution of the distance between the COMs of the tetrads-1 and 2; 2 and 3; and 3 and 4 respectively for the monomeric quadruplex (GQ) containing G-tetrad at position 2 (Quad-G4) corresponding to the four simulation sets (A-D).

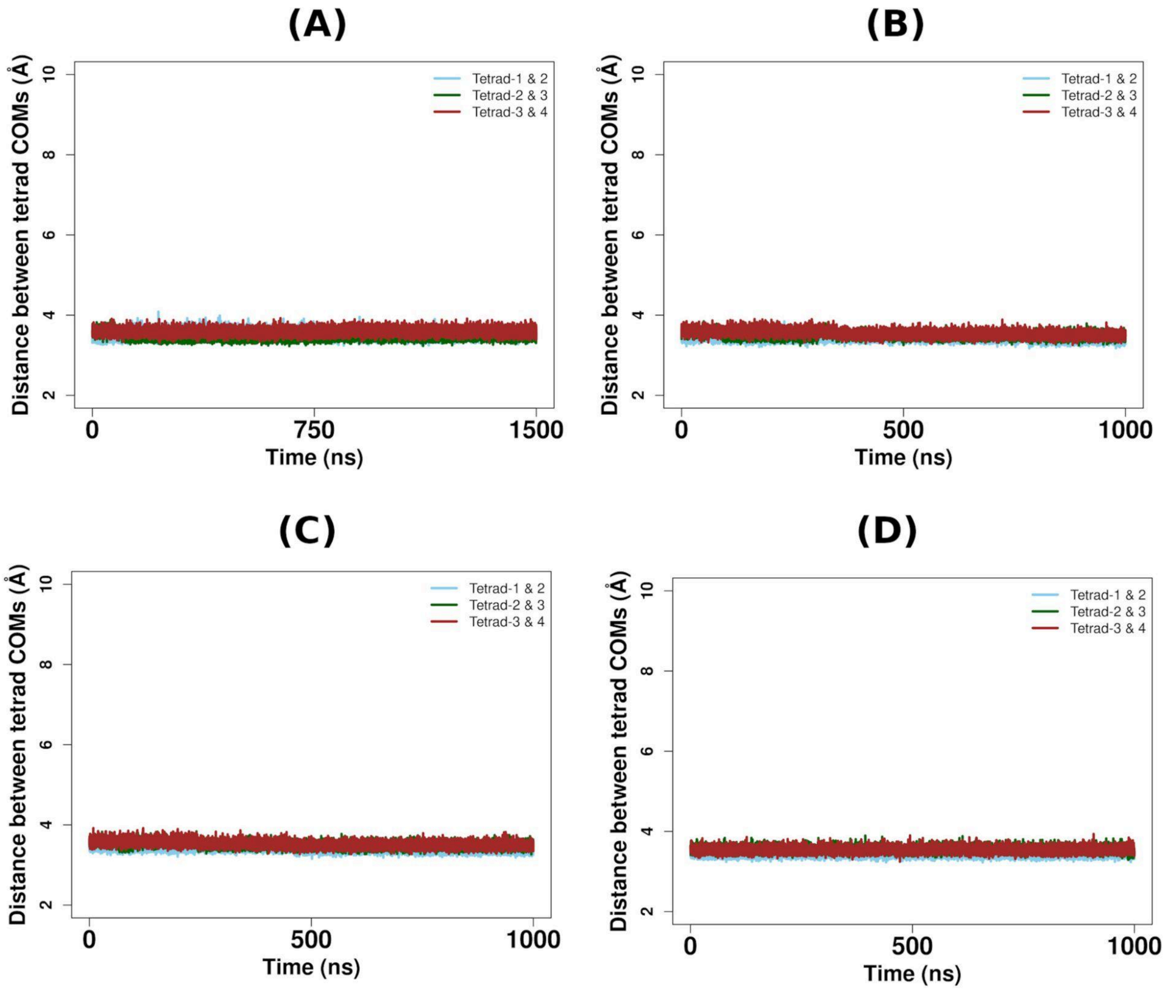


**Figure S47.** Time evolution of the distance between the COMs of the tetrads-1 and 2; 2 and 3; and 3 and 4 respectively for the monomeric quadruplex (GQ) containing Gm-tetrad at position 2 (Quad-Gm4) corresponding to the four simulation sets (A-D).

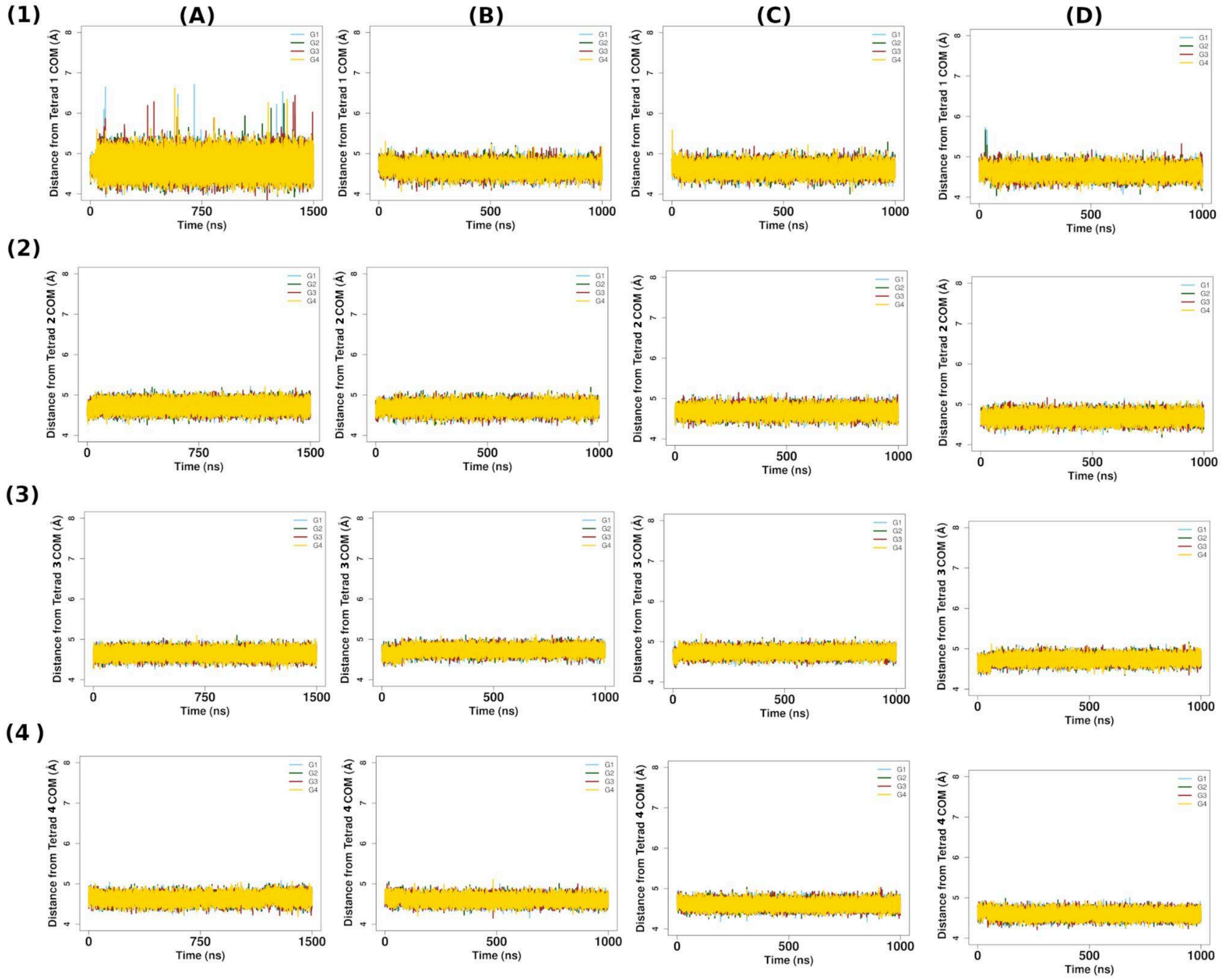


**Figure S48.** Time evolution of the distance between the tetrad-COM (for each of the tetrads 1-4 respectively) and COM each of the constituent G residues respectively for the monomeric quadruplex containing G-tetrad at position 2 (Quad-G4) corresponding to the four simulation sets (A-D).

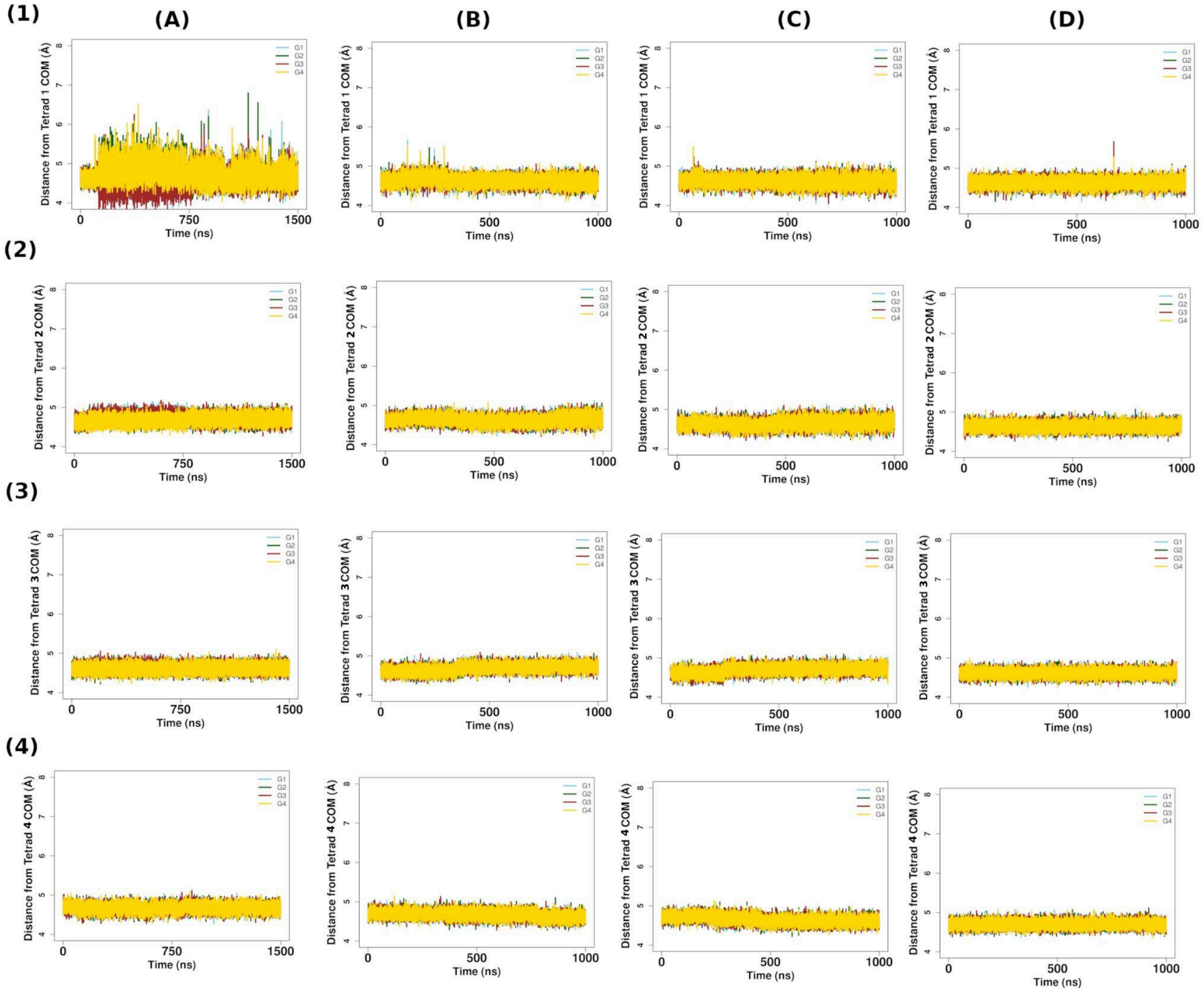


**Figure S49.** Time evolution of the distance between the tetrad-COM (for each of the tetrads 1-4 respectively) and COM each of the constituent G/Gm residues respectively for the monomeric quadruplex containing Gm-tetrad at position 2 (Quad-Gm4) corresponding to the four simulation sets (A-D).

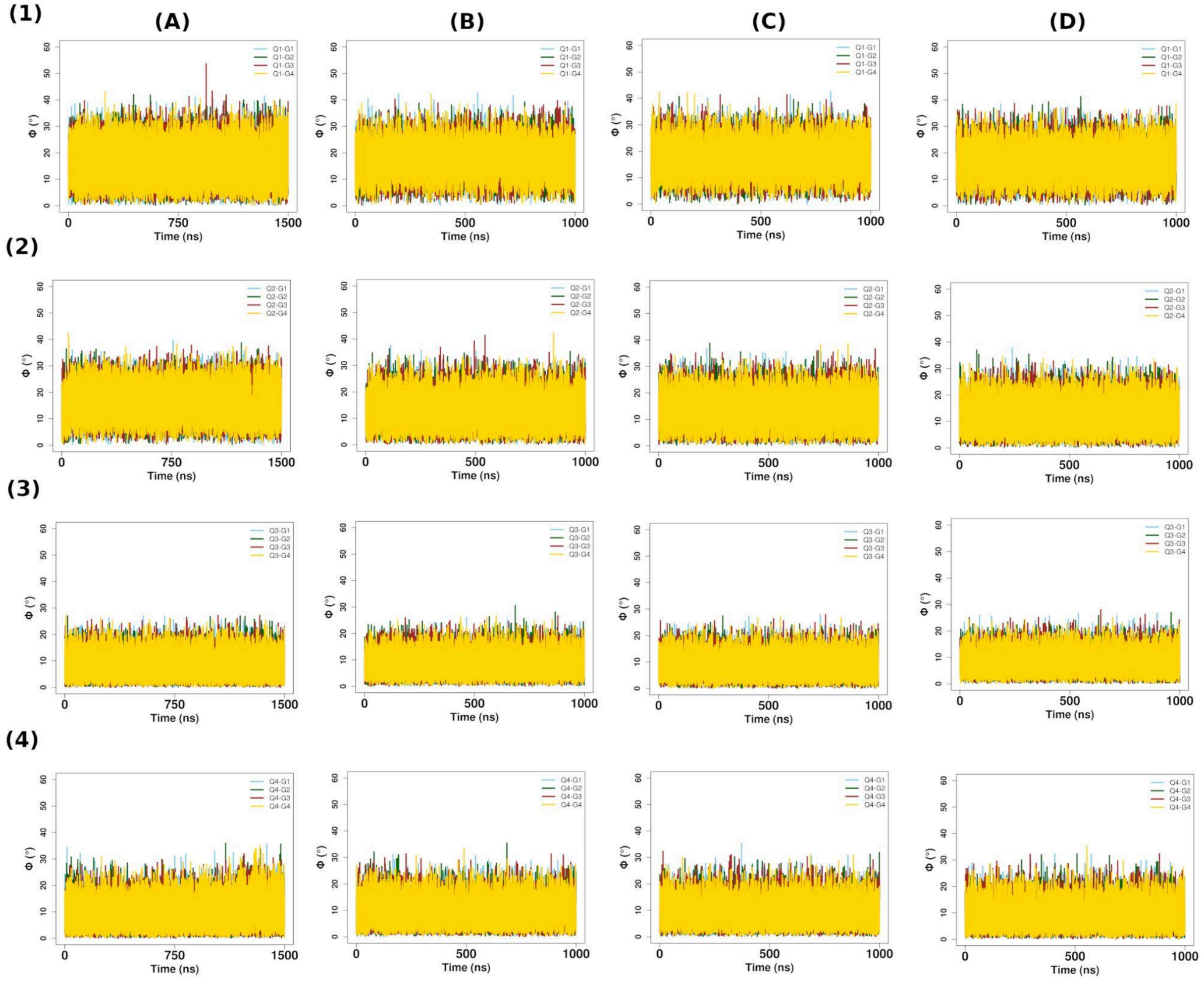


**Figure S50.** Time evolution of the angles (ϕ) between the normals to the G-planes in a tetrad and the axis of the tetrad (for each of the tetrads 1-4 respectively) and COM of each of the constituent G residues respectively for the monomeric quadruplex containing G-tetrad at position 2 (Quad-G4) corresponding to the four simulation sets (A-D).

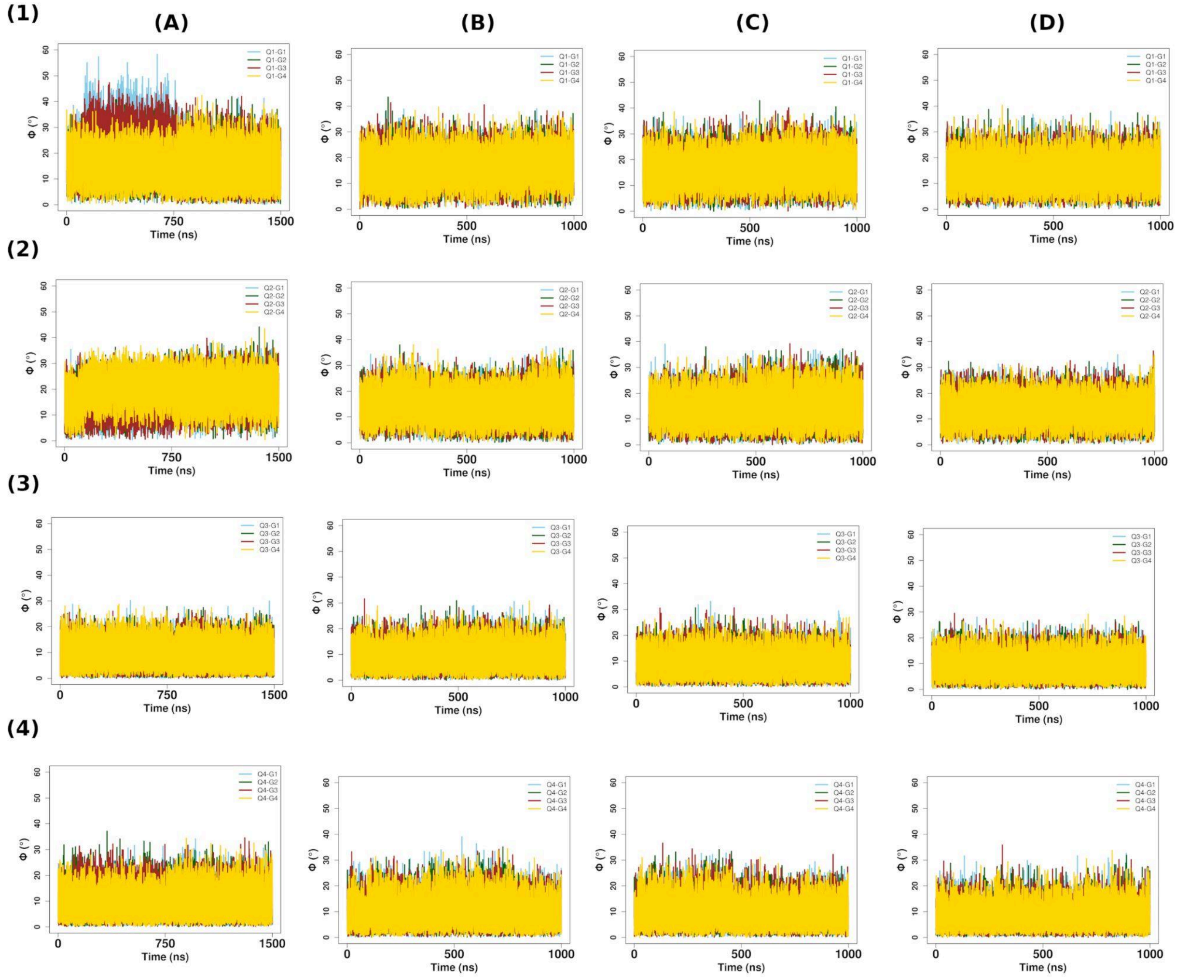


**Figure S51.** Time evolution of the angles (ϕ) between the normals to the G/Gm-planes in a tetrad and the axis of the tetrad (for each of the tetrads 1-4 respectively) and COM of each of the constituent G/Gm residues respectively for the monomeric quadruplex containing Gm-tetrad at position 2 (Quad-Gm4) corresponding to the four simulation sets (A-D).

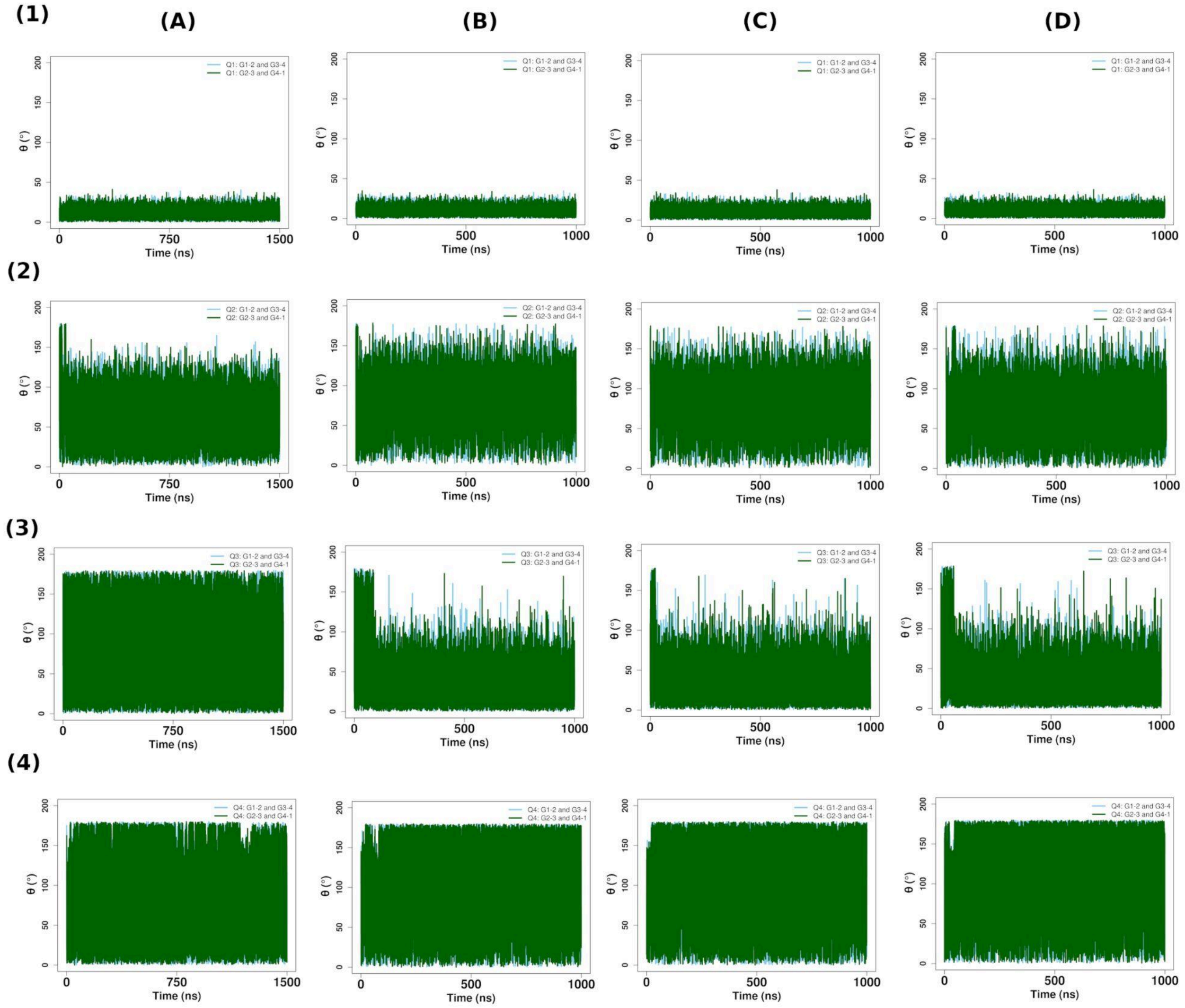


**Figure S52.** Time evolution of the angles (**θ**) between nominal planes formed by the G-pairs (for each of the tetrads 1-4 respectively) representing lengthwise tetrad-bending properties for the monomeric quadruplex containing G-tetrad at position 2 (Quad-G4) corresponding to the four simulation sets (A-D).

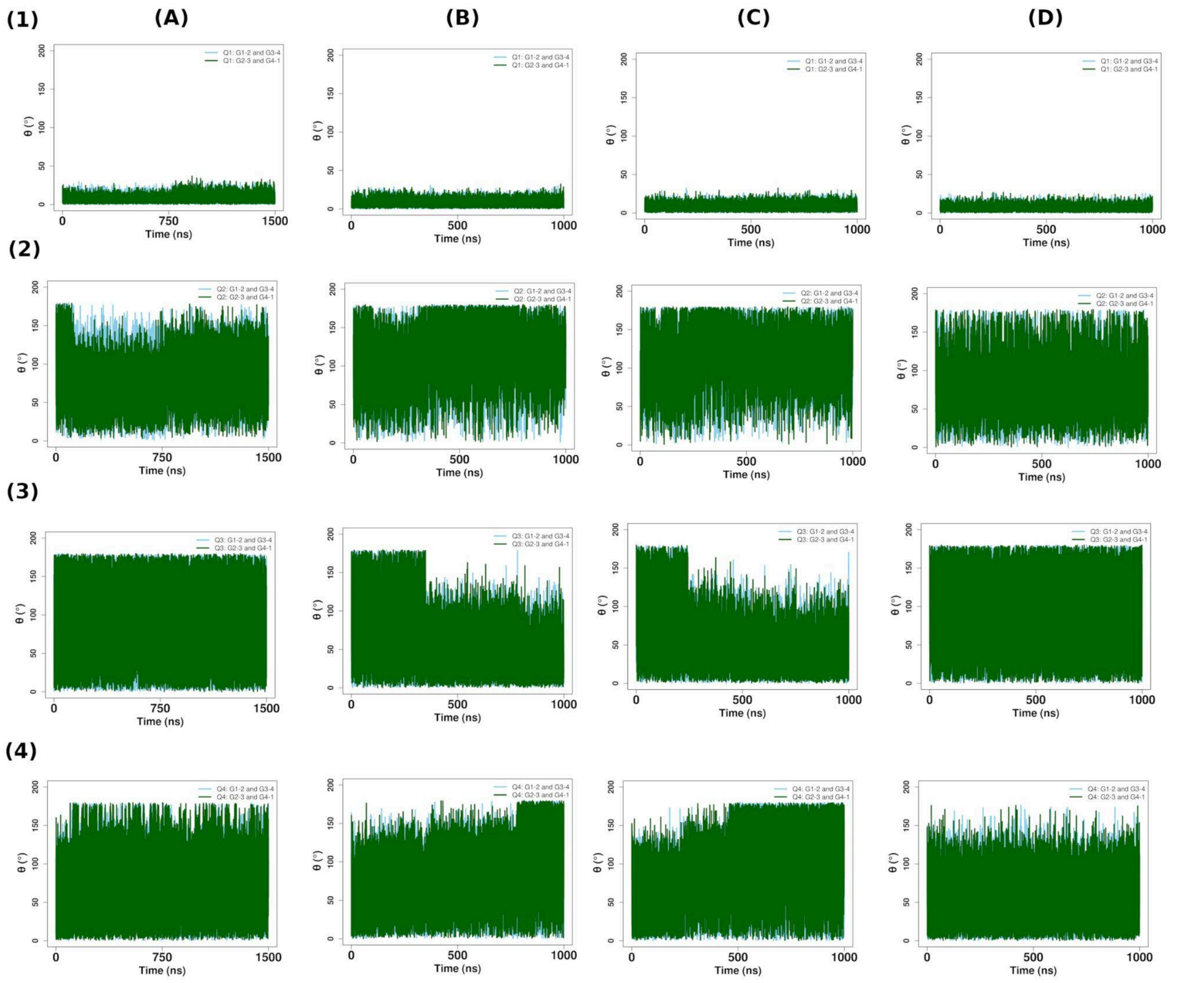

**Figure S53.** Time evolution of the angles (**θ**) between nominal planes formed by the G/Gm-pairs (for each of the tetrads 1-4 respectively) representing lengthwise tetrad-bending properties for the monomeric quadruplex containing Gm-tetrad at position 2 (Quad-Gm4) corresponding to the four simulation sets (A-D).

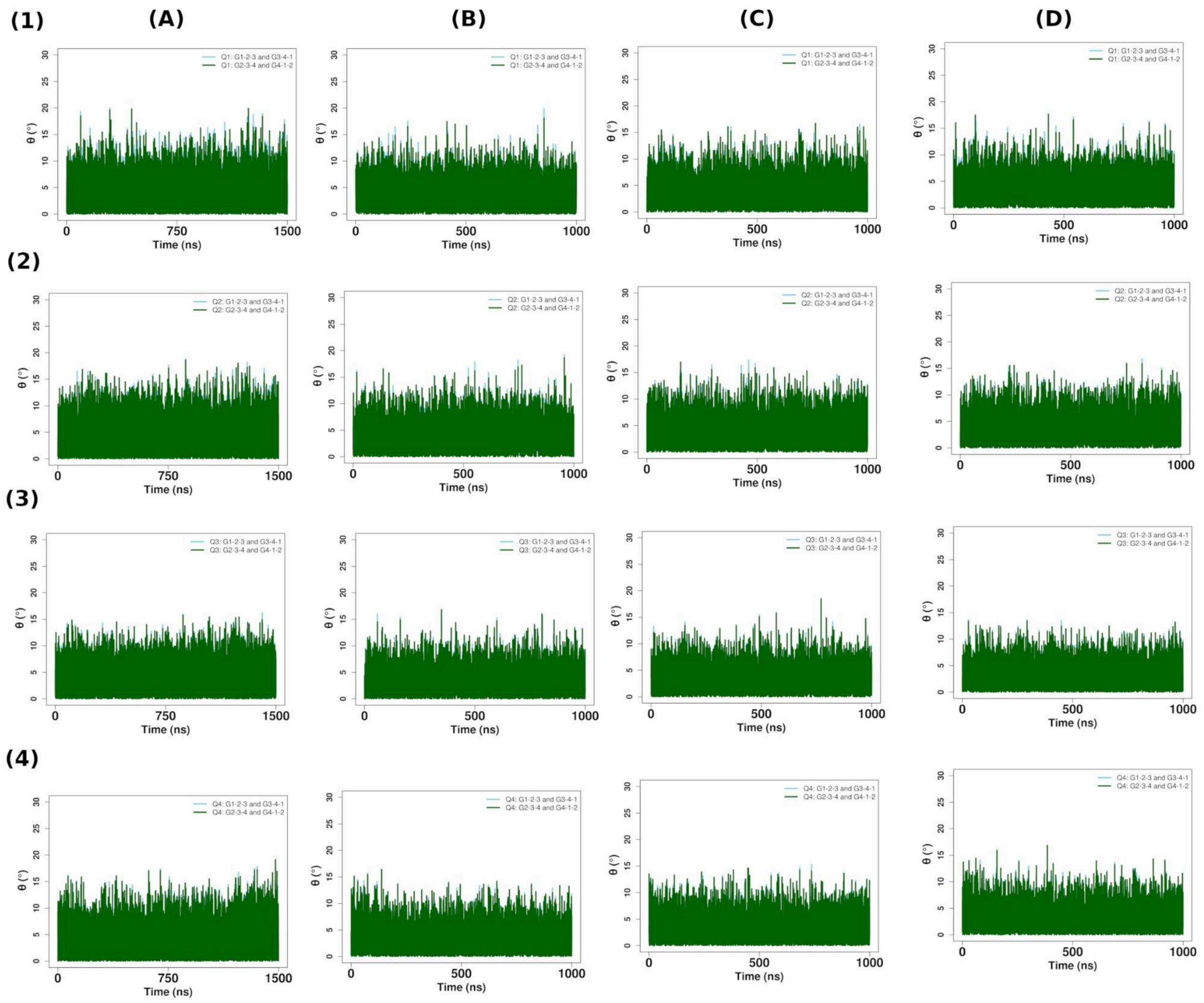


**Figure S54.** Time evolution of the angles (**θ**) between nominal planes formed by the G-triads (for each of the tetrads 1-4 respectively) representing diagonal tetrad-bending properties for the monomeric quadruplex containing G-tetrad at position 2 (Quad-G4) corresponding to the four simulation sets (A-D).

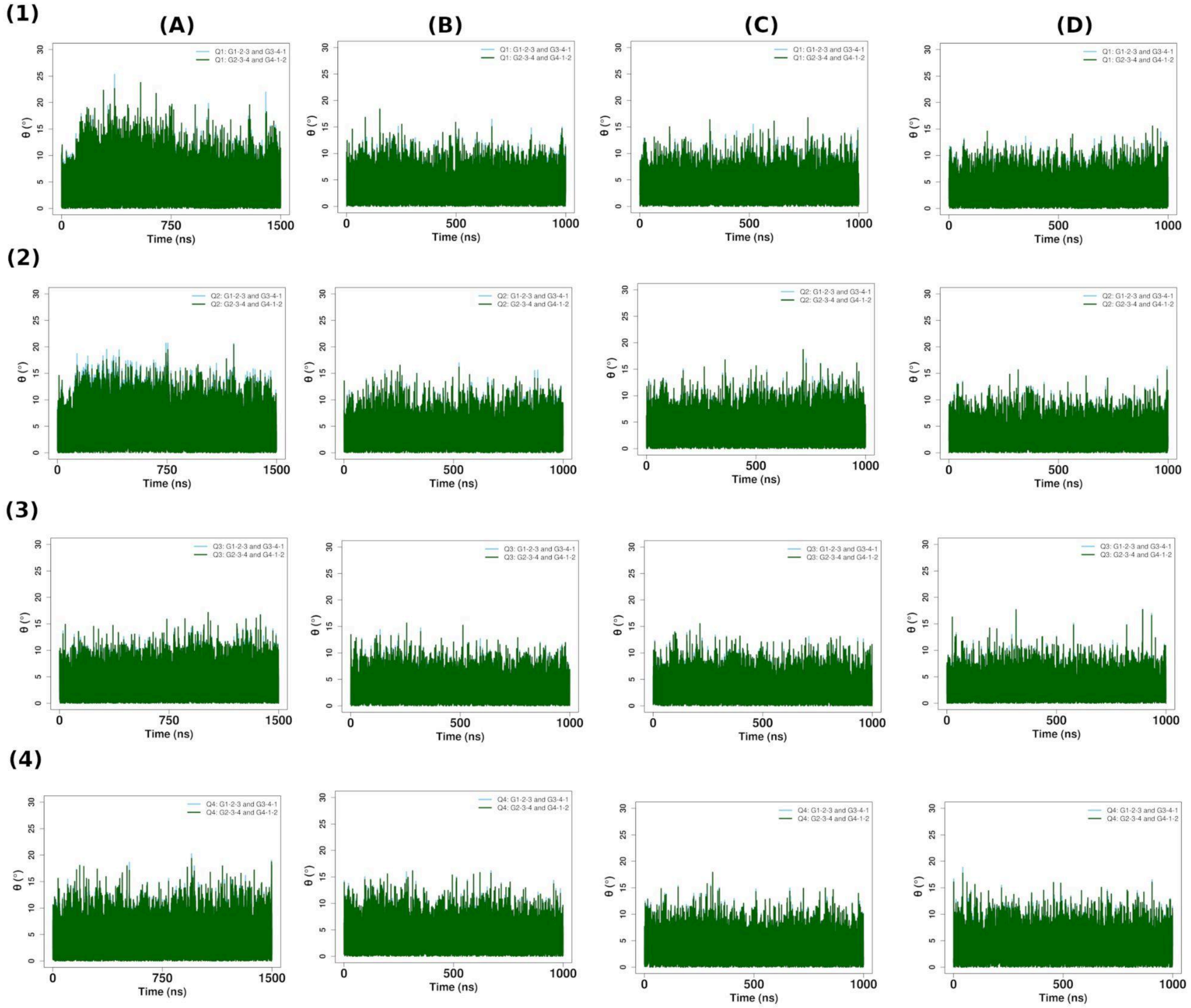


**Figure S55.** Time evolution of the angles (**θ**) between nominal planes formed by the G/Gm-triads (for each of the tetrads 1-4 respectively) representing diagonal tetrad-bending properties for the monomeric quadruplex containing Gm-tetrad at position 2 (Quad-Gm4) corresponding to the four simulation sets (A-D).

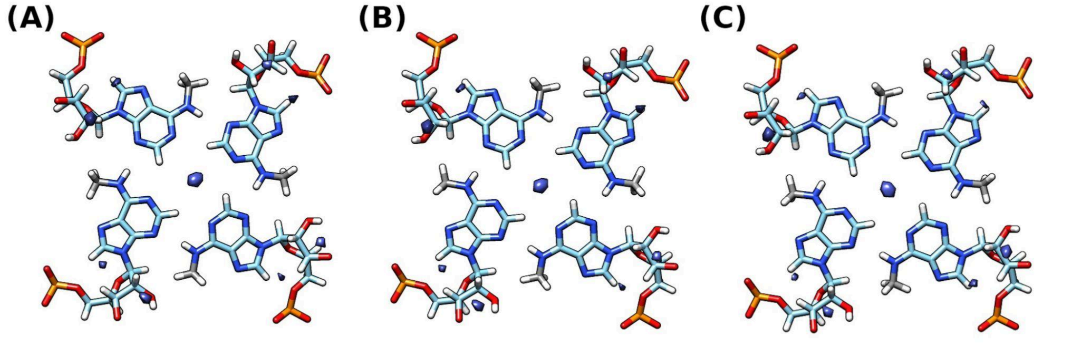


**Figure S56.** Water occupancy maps for m$^6$A-tetrad respectively for the (A) first (B) second and (C) third simulation sets (around the average structures).

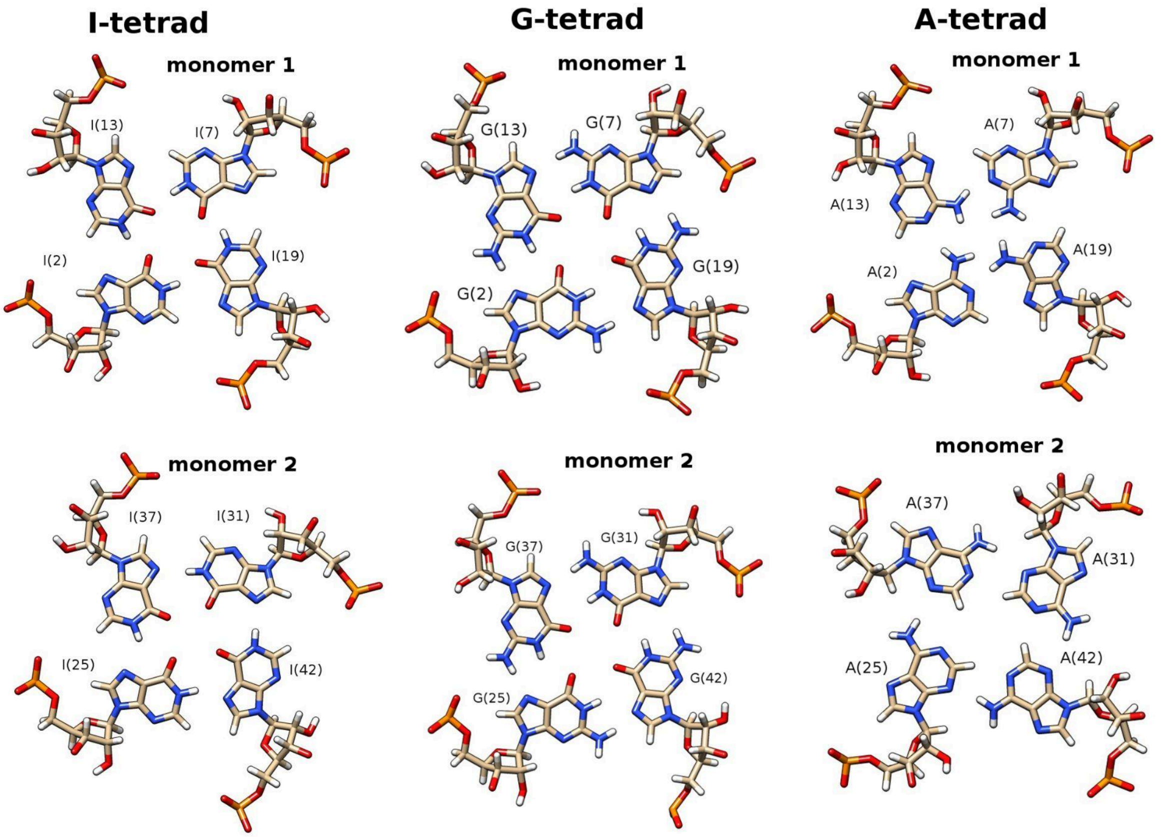


**Figure S57.** Different geometries observed for the three purine tetrads at position 2 (of each monomeric quadruplex within the quadruplex-dimer). The N1(H)-O6 (top) and N7-(H)N1 (bottom) geometries of I-tetrad observed in the fourth simulation set; the geometry of G-tetrad observed for each monomer in the fourth simulation set and the N6(H)-N1 (top) and N6(H)-N3 (bottom) geometries of A-tetrad observed in the second simulation set are shown.

(1)

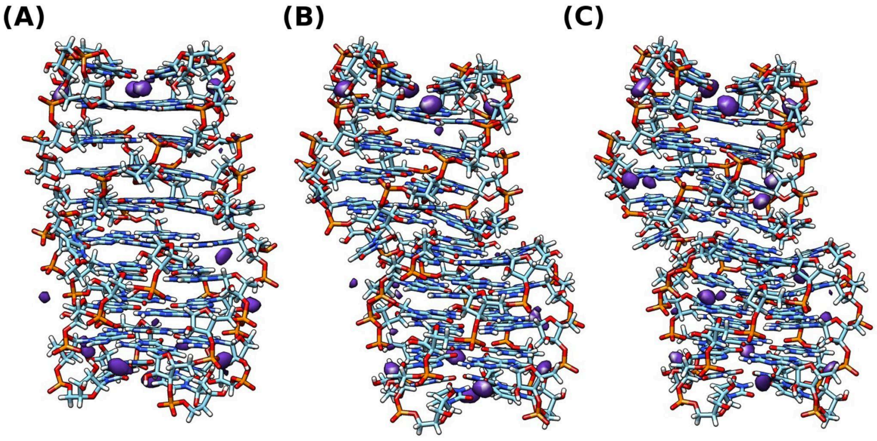

*(2)*

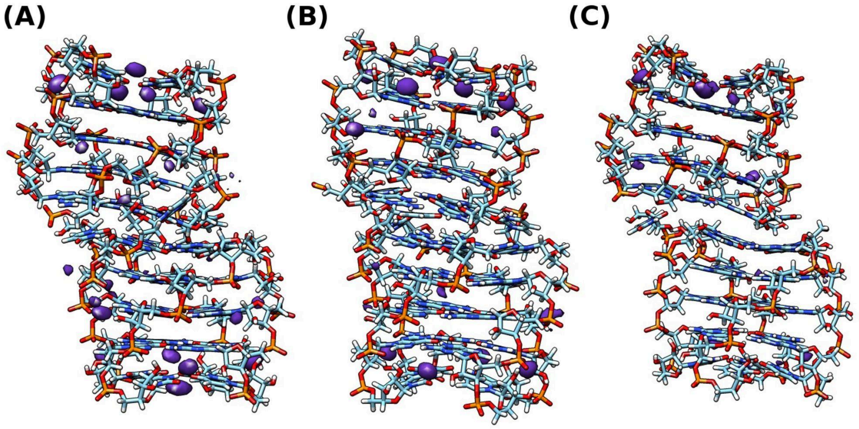


*(3)*

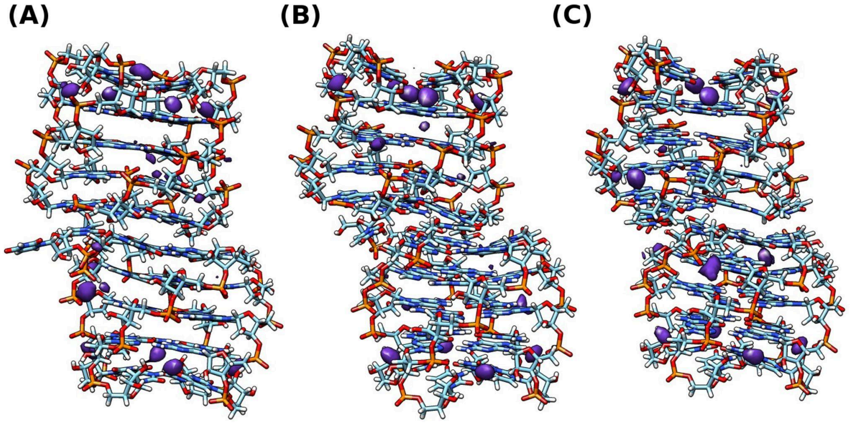


**Figure S58.** Occupancy maps of $K^+$ ions for the quadruplex dimers in this study from the first three sets (Sets-1, 2 and 3) of simulations (A) Quad-I4 (containing I(2) tetrads); (B) Quad-G4 (containing G(2) tetrads) and (C) Quad-A4 (containing A(2) tetrads). $K^+$ ion binding pocket is observed around the reversed U(5)-tetrads for each monomer.
